# Transcription factor motifs associated with anterior insula gene-expression underlying mood disorder phenotypes

**DOI:** 10.1101/864900

**Authors:** Dhivya Arasappan, Simon B. Eickhoff, Charles B Nemeroff, Hans A. Hofmann, Mbemba Jabbi

**Affiliations:** Center for Biomedical Research Support, University of Texas at Austin; Institute of Systems Neuroscience, Heinrich Heine University Düsseldorf, Germany; Institute of Neuroscience and Medicine (INM-7), Research Centre Jülich, Germany; Department of Psychiatry, Dell Medical School, University of Texas at Austin; The Mulva Clinic for Neurosciences, Dell Medical School, University of Texas at Austin; Institute of Early Life Adversity Research; Institute of Neuroscience, University of Texas at Austin; Department of Integrative Biology, the University of Texas at Austin; Department of Psychology, University of Texas at Austin

**Keywords:** Brain, gene expression, transcription factors, behavior, mood disorders, RNA-sequencing

## Abstract

**Background:** Mood disorders represent a major cause of morbidity and mortality worldwide but the brain-related molecular pathophysiology in mood disorders remains largely undefined.

**Methods:** Because the anterior insula is reduced in volume in patients with mood disorders, RNA was extracted from postmortem mood disorder samples and compared with unaffected control samples for RNA-sequencing identification of differentially expressed genes (DEGs) in *a*) bipolar disorder (BD; n=37) versus (vs.) controls (n=33), and *b*) major depressive disorder (MDD n=30) vs controls, and *c*) low vs. high Axis-I comorbidity (a measure of cumulative psychiatric disease burden). Given the regulatory role of transcription factors (TFs) in gene expression via specific-DNA-binding domains (motifs), we used JASPAR TF binding database to identify TF-motifs.

**Results:** We found that DEGs in BD vs. controls, MDD vs. controls, and high vs. low Axis-I comorbidity were associated with TF-motifs that are known to regulate expression of toll-like receptor genes, cellular homeostatic-control genes, and genes involved in embryonic, cellular/organ and brain development.

**Discussion:** *Robust imaging-guided transcriptomics* (i.e., using meta-analytic imaging results to guide independent post-mortem dissection for RNA-sequencing) was applied by targeting the gray matter volume reduction in the anterior insula in mood disorders, to guide independent postmortem identification of TF motifs regulating DEG. TF motifs were identified for immune, cellular, embryonic and neurodevelopmental processes.

**Conclusion:** Our findings of TF-motifs that regulate the expression of immune, cellular homeostatic-control, and developmental genes provides novel information about the hierarchical relationship between gene regulatory networks, the TFs that control them, and proximate underlying neuroanatomical phenotypes in mood disorders.

## INTRODUCTION

Adaptive behavior is, in part, governed by genes, especially their coding regions, through gene-mediated molecular processes that are critical for brain development and function. These gene mediated processes, through changes in gene expression, can give rise to complex behavioral phenotypes. Transcription factors (TFs) are sequence-specific DNA-binding proteins that are also known as regulatory proteins. TFs regulate the expression of genes by recognizing and binding to specific DNA regulatory elements called DNA binding domains in the promoter region of genes (Changeaux 2017; Lambert et al. 2018). This attribute enables TFs to hierarchically regulate the expression of genes by controlling (i.e. promoting/activating or blocking/repressing) transcription of the adjacent coding regions into mRNA, and subsequent translation into proteins (Changeaux 2017; Lambert et al. 2018). TFs are therefore in a functionally elevated status in the hierarchy of gene expression repertoires because they are able to cooperatively, or synergistically regulate genes encoding other TFs (Lambert et al. 2018). Consequently, gene regulatory networks underlie essential biological processes such as brain development, synaptic formations and emergent behaviors like learning and memory that depend on neurodevelopmental processes (Kerszberg & Changeaux 1998; Tsigelny et al. 2013).

Based on the complex role of thousands of transcription factors in controlling gene regulation, dysregulation of TF-mediated gene expression programs has been hypothesized to contribute to a broad range of diseases (Lee & Young 2013), including neuropsychiatric disorders (Lee & Young 2013; Nord AS, Pattabiraman K, Visel A & Rubinstein JLR 2015; Changeaux 2017; Torre-Ubieta et al. 2018). However, the putative role of TF networks in mood disorder pathogenesis is not well understood. Adult brain gray matter volumetric (GMV) reductions in the anterior part of the insula, a region of the cerebral cortex folded deep within the lateral sulcus, have been consistently identified in mood disorders (Goodkind et al. 2015; Wise et al. 2016). Such reduced GMV are found to predict cognitive impairment and affective dysfunction in both MDD and BD (McTeague et al. 2017). Functional magnetic resonance imaging (fMRI) of the right anterior insula region has been reported to selectively predict the therapeutic response to psychotherapy and pharmacotherapy (McGrath et al. 2013). Because mood disorders are the most common neuropsychiatric syndromes (Yizhar 2012), and they collectively account for a high global burden of disease (Collins et al. 2010), the current study examined the role of TFs in anterior insula gene expression profiles associated with affective dysfunction. Specifically, we used robust imaging-guided transcriptomics, a method that performs meta-analyses of neuroimaging results of gray matter changes associated with a disease phenotype (i.e., mood disorder diagnosis), to guide independent post-mortem dissection of the identified regional gray matter change for RNA-sequencing studies of gene expression profiles for the disease phenotype in question (i.e., mood disorders). We tested the hypothesis that TF motifs will be associated with differentially expressed genes (DEGs) in the postmortem anterior insula cortex of mood disorder (i.e., MDD & BD) donors relative to controls. This robust imaging-guided transcriptomics methods enabled the goal of realizing a more anatomically precise RNA-seq study of the putative pathological tissue associated with mood disorder diagnoses. The results indicate that relative to controls, DEGs in postmortem mood disorder tissue are associated with TF motifs known to regulate expression of i) toll-like receptor signaling/immune and inflammatory processing genes, *b*) cellular homeostatic control genes, and *c*) genes involved in cellular and brain developmental processes.

## METHODS

### Localization of brain gray matter loss in mood disorders

To identify the brain’s most anatomically proximate regional involvement in mood pathology, with the goal of targeting the identified pathological sites for postmortem transcriptomics, we first performed a large scale meta-analysis of voxel based morphometry studies of gray matter loss in mood disorders using the anatomical likelihood estimation approach. The anatomical likelihood estimation approach models the spatial uncertainty associated with each reported location for significant between-group differences (Eickhoff et al. 2009), and further compute the convergence across all included experiments by the union of the ensuing probabilistic model relative to a null-distribution and thereby reflecting a random spatial association between the findings of different experiments (Eickhoff et al. 2012). The identified brain region exhibiting the most extensive gray matter loss in mood disorder brain imaging cohorts (**Figure 1A**) was used for our robust imaging-guided transcriptomics by first generating native space reconstruction of the reduced anatomical sub-region (**Figure 1 B & C**), and then using the reconstructed images were to guide postmortem tissue section in independent samples (**Figure 1 C**). Specifically, the identified gray matter loss was demarcated in 3-D space to guide the dissection of the targeted region in the independent postmortem mood disorder cohorts.

**Figure 1.**
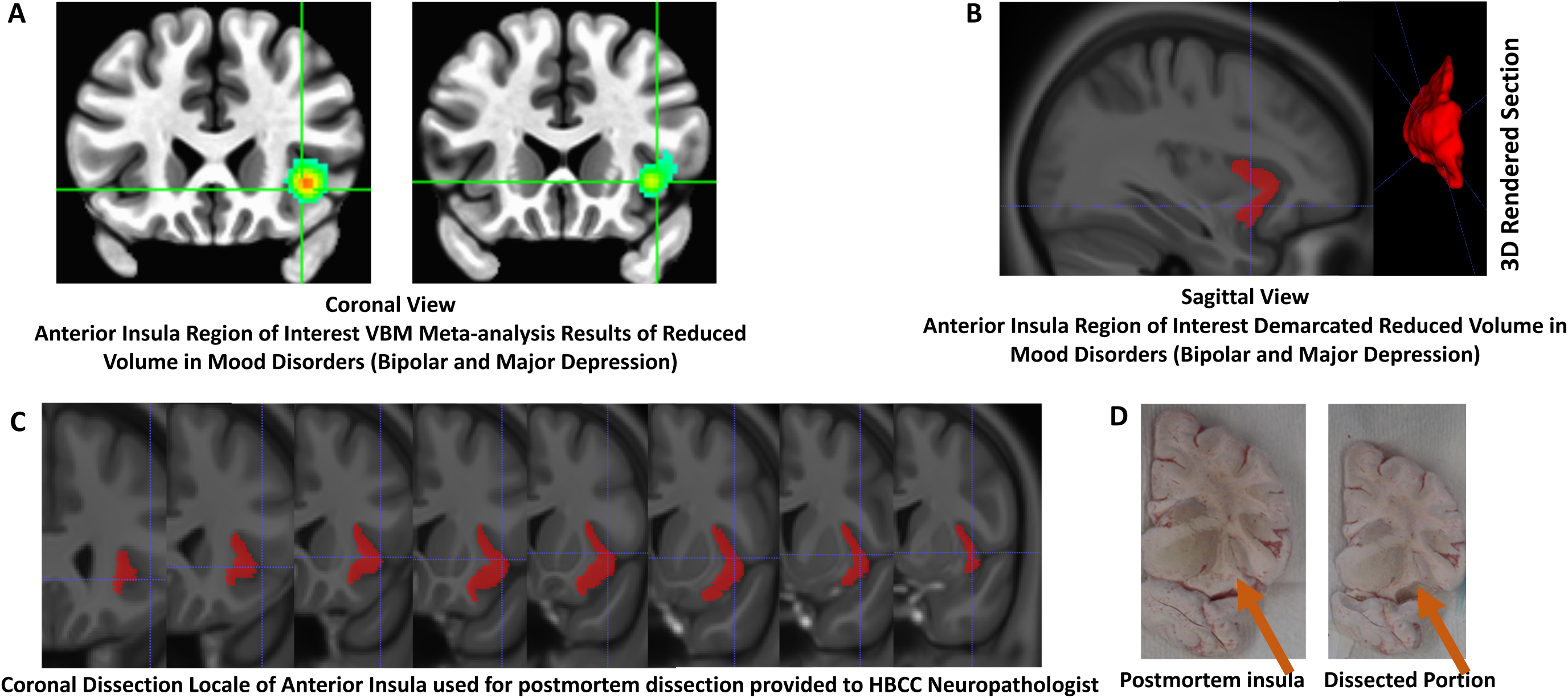
A-B. Gray matter loss in anterior insula cortex of BD and MDD vs. controls as identified with largescale voxel-based morphometry imaging meta-analysis. A) illustrates localized anterior insula gray matter reduced area associated with mood disorder diagnoses on a coronal section; and B) illustrates the sagittal reconstruction of the reduced volume in A as well as the 3-dimentional view of the reduced volume. The anatomical information in A and B illustrated slide by slide on coronal sections in C) and these images are used to guide postmortem dissection of anterior insula tissue from an independent sample in D.

### Postmortem variable factor-analysis

To explore morbidity related gene expression profiles beyond the conventional DSM diagnosis-centered case vs. control statistical comparisons, we conducted an exploratory factor analysis, a data reduction technique, to identify higher-order composite variables included with the postmortem data. For each postmortem sample, the donor data includes specific mood disorder and comorbid lifetime-Axis-I comorbidity (i.e., number of lifetime-Axis-I diagnostic occurrences, e.g. (poly)-substance use disorders, psychosis, anxiety, eating disorders, etc. alongside the primary mood disorder diagnosis of BD or MDD); comorbid lifetime-Axis-III diagnoses (i.e. number of lifetime-Axis-III medical conditions such as diabetes, cancer, cardiovascular disease, etc.); cause of death (i.e. death by suicide, homicides, accidents or natural death due to comorbid Axis-III medical conditions); and as specified by the medical examiner reports (e.g. blunt force trauma to the chest, gunshot, motor vehicle accident, drowning, hanging, etc.); demographics (race, age at death, sex, years of education, number of children/fecundity, and marital records); technical variables (brain-weight, postmortem interval ‘i.e. the time that has elapsed since a person has died’, pH, and RNA integrity number (RIN)); and toxicology (blood alcohol/blood narcotics levels) (see **Table 1** for diagnostics relations with demographics, suicide and other manners of death, positive toxicology and postmortem qualitative data; & **Supplementary Table 1** for diagnostic comorbidity on axis-I/co-occurring lifetime mental illness,). Principal Axis Factoring (Oblimin Rotation with Kaizer Normalization) (Costello & Osborne 2005) was applied to identify higher-order factors explaining the differences in postmortem variables and included those variables with communalities of ≥ 0.5. Given our focus on identifying TF motifs associated with mood disorder comorbidity burden, we first conducted DEG analysis for bipolar disorder vs. controls, MDD vs. controls, and high vs. low Axis-I comorbidity by conducting a split-half comparison of the lower half vs. the higher scoring donors on this higher-order variable representing Axis-I comorbidity.

**Table1.**
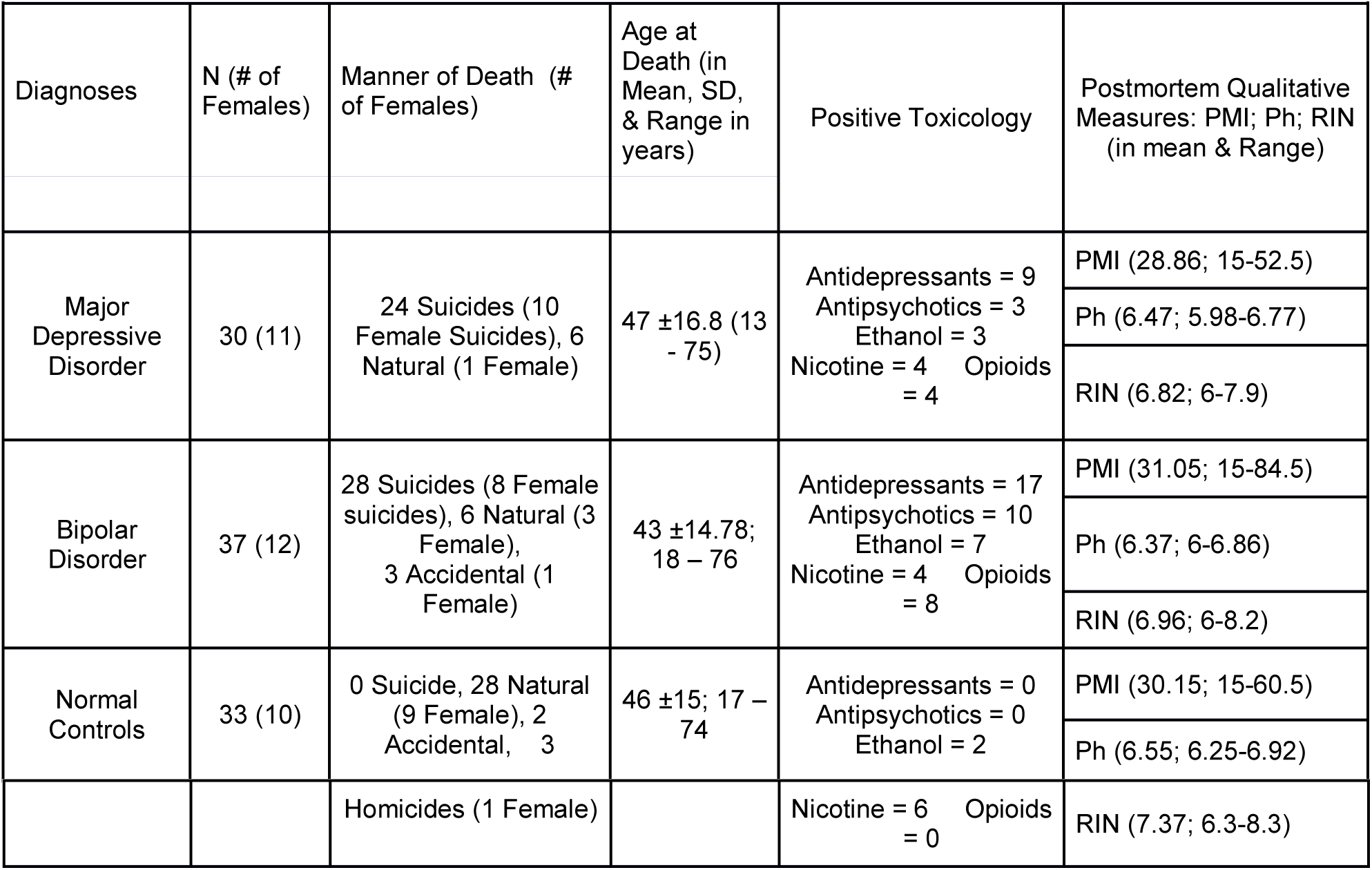
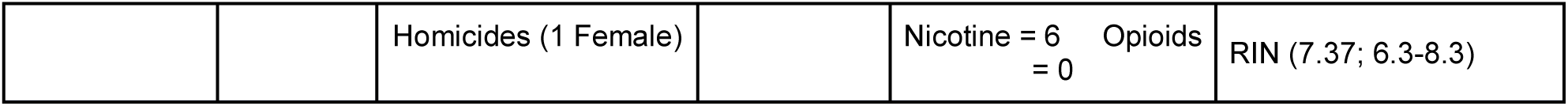
Anterior Insula sample diagnostics, suicide, demographics, toxicology and *postmortem* quality data

### Brain Dissection

The NIMH Human Brain Collection Core (HBCC) provided the *Postmortem* samples for which informed consents are acquired according to NIH IRB guidelines. Clinical characterization, neuropathology screening, and toxicology analyses followed previous protocols (Martin et al. 2006). We applied *robust imaging-guided transcriptomics* as follows: first, the region of interest targeted for dissection was defined as portion of right AIC encompassing the identified in a meta-analysis of imaging studies to harbor the most reduced GMV subsection of the entire brain in mood disorders (See **Figure 1A**). Electronic image slide of the imaging-defined GMV loss volumes (**Figure 1 B & C**) were then shared with the HBCC neuropathologist who used these images to guide dissection of coronal tissue slabs of each postmortem donor brain (see **Figure 1D**) at the NIH clinical center.

### RNA-Extraction

All dissected tissues were separately pulverized and 50 mg aliquoted from each sample for standardized total RNA processing. Specifically, RNeasy Lipid Tissue Mini Kit (50) was used for RNA purification using the 50 RNeasy Mini Spin Columns, Collection Tubes (1.5 ml and 2 ml), QIAzol Lysis Reagent, RNase-free Reagents and Buffers kit from Qiagen. DNase treatment was applied to the purified RNA using Qiagen RNase-Free DNase Set (50) kit consisting of 1500 Kunitz units RNase-free DNase I, RNase-free Buffer RDD, and RNase-free water for 50 RNA minipreps. After DNAse treatment, the purified RNA from the pulverized AIC tissue sections were used to determine RNA quality as measured in RNA integrity number (RIN) values using Agilent 6000 RNA Nano Kit consisting of the microfluidic chips, Agilent 6000 RNA Nano ladder and reagents on Agilent 2100 Bioanalyzer. Samples with RIN < 6 were excluded and the 100 samples meeting inclusion were shipped directly from the NIMH HBCC core to the Genome Sequencing and Analysis Facility (GSAF: https://wikis.utexas.edu/display/GSAF/Home+Page) at the University of Texas, Austin, USA for RNA-sequencing.

### Illumina-Sequencing, Read-Mapping and Gene-Quantification

Total RNA was extracted and only samples with RNA integrity numbers (RIN values) greater than 6 as confirmed using the Agilent Bioanalyzer were used for library preparation. First. Ribosomal RNA was depleted using RiboMinus Eukaryote kit from Life Technologies (Foster City, CA, USA) for RNA-Seq and confirmed using an Agilent Technologies’ Bioanalyzer (Santa Clara, CA, USA). mRNA selection was completed using the Poly(A) purist kit from Thermofisher and paired-end libraries with average insert sizes of 200bp were obtained using NEBNext Ultra II Directional RNAs Library Prep kit from New England BioLabs. All 100 samples were processed and then sequenced on the Illumina HiSeq 4000, PE150, at the Genome Sequencing and Analysis Facility at UT Austin, USA.

30 million paired-end reads per sample (150 base pairs in length) were generated by sequencing runs of 4 samples per lane of the sequencer. Sequenced reads were assessed for quality with Fastqc (Andrews 2010) to specifically assess sequencing reads for median base quality, average base quality, sequence duplication, over-represented sequences and adapter contamination.

### Differential Gene Expression Analysis

The reads were pseudo-aligned to the human reference transcriptome (GRCh38-gencode) using kallisto (Bray 2016), and transcript-level abundances were obtained. The transcript-level counts were aggregated to gene-level using tximport in R. The abundances were normalized using DESeq2 (Anders & Huber 2010; Love et al. 2014), and transformed with variance stabilizing transformation (defined here as a transformation technique that seeks to create more homoscedasticity, thereby having a closer to constant variance in the dataset regardless of the mean expression value (Love et al. 2014)). We performed differential expression analysis based on the negative binomial distribution for modeled gene counts using DESeq2. RIN-values were included in the DESeq2 design matrix as a covariate to control for the potentially confounding effects of RNA quality. The analysis controlled for group factors as well as possible individual outliers by removing genes with expression values of 0 in 80% or more of the samples. We performed differential expression analysis to identify DEGs in the following comparisons: BD vs. controls; MDD vs. controls, and in a pooled cohort of BD & MDD individuals with high vs. low Axis-I comorbidity at absolute fold change >=2 and adjusted p-values <=0.1 were selected as significantly differentially expressed genes. The DESeq2 package by default calculates false discovery rate adjusted p-values according to Benjamini and Hochberg (Benjamin & Hochberg 1995). We used an adjusted p-value <=.1 as a cut-off to balance type-1 and type-2 error rates and allow more inclusive capture of regulatory elements/TFs associated with DEGs in for mood disorders across bipolar disorder and unipolar depression.

### JASPAR 2018 TFs binding profiling

Given the role of transcription factors (TFs) in regulating gene expression via specific DNA-binding domains (motifs) in the gene promoters, we followed-up on our previous study (Jabbi et al. 2020) that identified whole transcriptome gene expression profiles in mood disorders, with the aim to explore in more detail, mood disorder transcriptomics by identifying TF motifs. To this goal, we used JASPAR TF binding database to identify motifs that were associated with DEGs in BD vs. controls; MDD vs. controls, and in pooled mood disorder individuals with high vs. low Axis-I comorbidity.

JASPAR is an open access database of non-redundant, manually curated TF-binding profiles provided as a publicly available web framework (Khan A et al. 2018). The JASPAR 2018 version has an updated list of 579 non-redundant TF-binding profiles of the vertebrate taxonomy that are systematically stored as position frequency matrices (PFMs), which summarizes experimentally determined DNA sequences bound by an individual TFs by counting the number of occurrences of each nucleotide at each position within aligned TF-binding sites (Khan A et al. 2018). This JASPAR 2018 CORE vertebrate database of 579 PFMs was first used to predict TF-binding sites in the human genome, and then made available to the scientific community through the UCSC Genome Browser track data hub (http://jaspar.genereg.net/genome-tracks/) for use to identify specific TF-binding profiles.

*EnrichR* (Kuleshov et al. 2016) was used to identify TF motifs that were associated with our DEGs. We focused on the top 10 most significant TF motifs found in the database of 579 PFMs associated with the DEGs observed in our DESeq2 results dataset for each of the 3 analytical contrasts (i.e. BD vs. controls; MDD vs. controls; pooled mood disorder cohort with high Axis-I comorbidity vs. pooled mood disorder cohort with low Axis-I comorbidity). TF motifs within the JASPAR 2018 that were found to be associated with DEGs at adjusted p-value cutoff of 0.1 (i.e. using the false discovery rate Benjamini and Hochberg multiple comparison method (Benjamini & Hochberg 1995) were identified as identified TFs associated with each of the 3 DEG contrasts (i.e. BD vs. controls; MDD vs. controls; low vs. high Axis-I comorbidity) we selected the top 10 TF motifs (see **Figures 2&3**).

**Figure 2.**
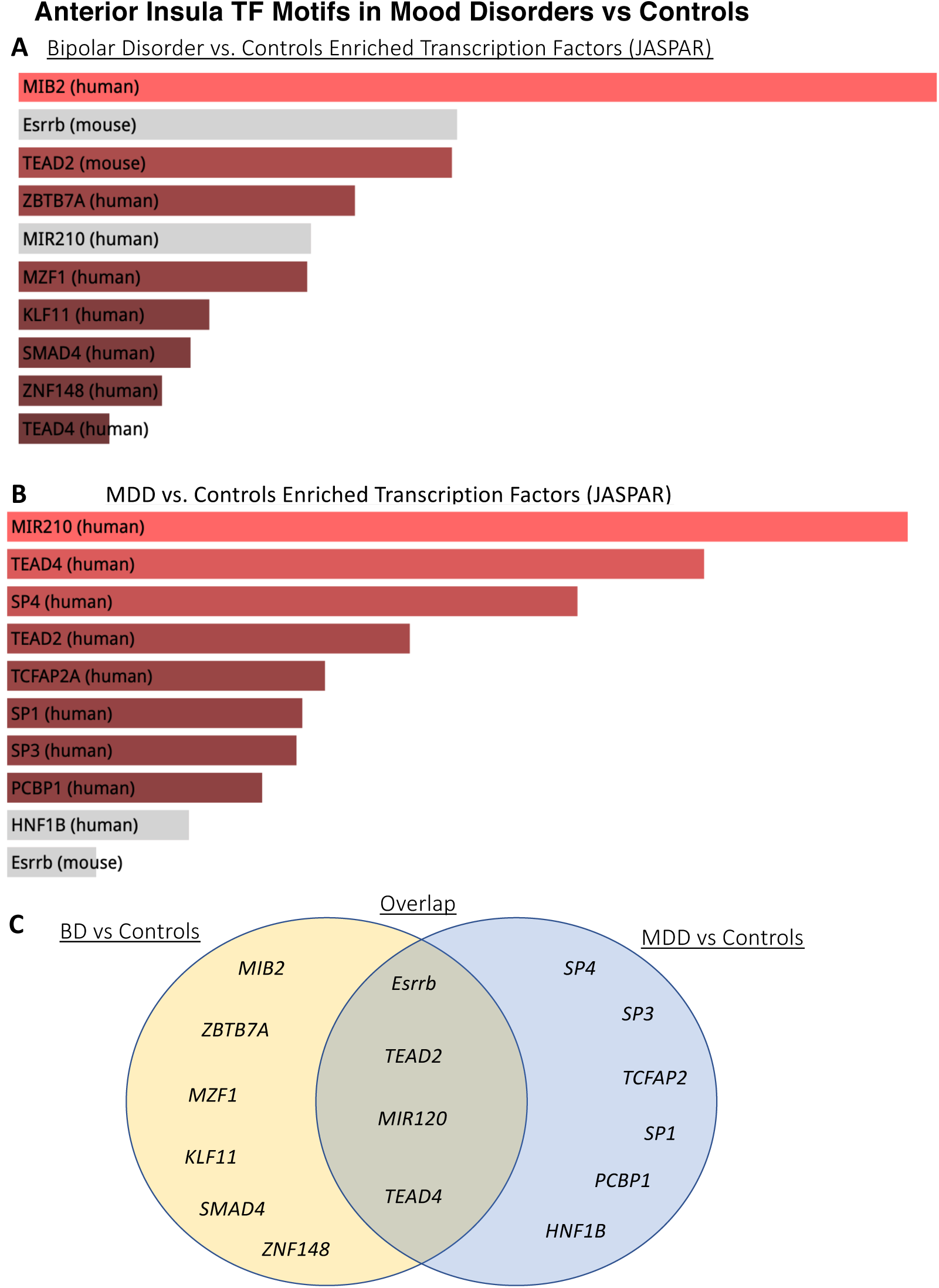
Anterior Insula postmortem TF motifs for gene expression profiles in BD and MDD relative to controls. A) Gene expression profiles in BD vs. Controls are illustrated to be associated with the top ten TF motifs for genes that are implicated in regulating inflammatory and immune responses, early embryonic and cellular development, and in post transcriptional gene expression. B) Gene expression profiles in MDD vs. Controls shown to be associated with the top ten TF motifs known to regulate inflammatory and toll-like receptor signaling, cellular development and peripheral homeostatic/hormonal signaling and post transcriptional regulation/DNA methylation. C) Venn diagram for TF motifs specific for BD vs. controls, MDD vs. controls and the overlapping TFs motifs for which were found for both BD vs. controls and MDD vs. controls.

## RESULTS

### Robust Imaging-guided Transcriptomics

Using the anatomical likelihood estimation meta-analysis (Eickhoff et al. 2009), we previously identified the right anterior insula as the brain region with the most extensive GMV reduction in mood disorders relative to healthy controls (***Figure. 1A***) (Jabbi et al. 2020). We manually reconstructed this region in ITKSnap (http://www.itksnap.org/pmwiki/pmwiki.php) and provided this 3-D volume information to the pathologist at the brain bank to guide postmortem dissection of anterior insula tissue from each sample for subsequent RNA extraction and RNA-sequencing. The dissected regional volume that captured the reduced GMV corresponded to the anterior portion of the insula, where the caudate and putamen are approximately equal in size (***Figure. 1A-D***).

### Postmortem Variability

Factor analysis identified two higher-order factors representing 1) number of Axis-1 diagnostic comorbidity and suicide completion (e.g. substance use disorders or abuse/psychosis/anxiety, and whether the donors died by suicide), and 2) RNA integrity number (RIN) values which were subsequently included in all differential gene expression analyses as covariates. The identified higher order factor Axis-I diagnostic comorbidity and suicide completion were used for the comparison of high versus low scores of each of these variables to determine differential gene expression and their regulatory TF motifs.

Apart from race, no other demographic variables differed across groups [MDD & BD samples had more Caucasian donors whereas controls had more African American donors (p<0.0001, F=12)]. Covarying for race in subsequent ANOVAs, *Axis-I comorbidity burden*, which clustered with mood disorder donor sample *suicide completion rate* in the explorative factor analysis, was different across groups (p<0.0001, F=30); showing Bonferroni corrected pairwise-comparison differences between MDD > controls (p<0.0001); BD > controls (p<0.0001); and BD > MDD (p = 0.004).

### DEGs and JASPAR TF motif identification of group-related differential gene expression

We found 456 differentially expressed genes for the bipolar vs. controls (**Supplementary Table 2**) and 2722 differentially expressed genes for MDD vs. controls (**Supplementary Table 3**). Further analysis that includes DEG results from more than one brain region in the same mood disorder samples will help explain if our observed higher number of DEGs in MDD vs. control samples relative to bipolar disorder vs. controls is pathophysiologically relevant in a region-specific manner. By looking upstream of BD vs. control DEGs for enriched TF motifs, we found that DEGs had motifs of *MIB2, Esrrb, TEAD2, ZBTB7A, MIR210, MZF1, KLF11, SMAD4, ZNF148*, and *TEAD4* TFs (***Figure. 2A***). Similar analysis of MDD vs. controls DEGs identified associated motifs of *MIR210, TEAD4, SP4, TEAD2, TCFAP2A, SP1, SP3, PCBP1, HNF1B*, and *Esrrb* TFs (***Figure. 2B***).

Given our approach to examine the hierarchical relationship between TF motifs and DEGs profiles associated with the degree of psychiatric morbidity/Axis-I comorbidity in the two mood disorder cohorts, we first used principal axis factoring to identify *Axis I comorbidity (i*.*e. total number of psychiatric diagnoses)*, and *manner of death by suicide or non-suicide* (together comprising the factor we refer to here as the Axis-I comorbidity burden) to compositely explain variability in (i) <u>Axis-I comorbidity burden</u>, and (ii) <u>suicide completion</u>. We included this factor in our analysis using a split half approach of high vs. low Axis-I comorbidity burden to identify TF motifs for DEG profiles for Axis-I comorbidity burden (see **Supplementary Table 4 for complete DEGs**). We found DEGs in the pooled BD & MDD donor samples (i.e. without control donors) and these DEGs were associated with TF motifs of *NFATC2, GABPA, HMAGA1, NR3C1, GTF2I, IRF2, POU1F1, SAMD9L, SNAI1*, and *CBEPB* (***Figure. 3***). Collectively, these TFs are known to regulate expression of toll-like receptor signaling genes, cellular homeostatic control genes, and genes involved in embryonic, cellular and neurodevelopmental processes.

**Figure 3.**
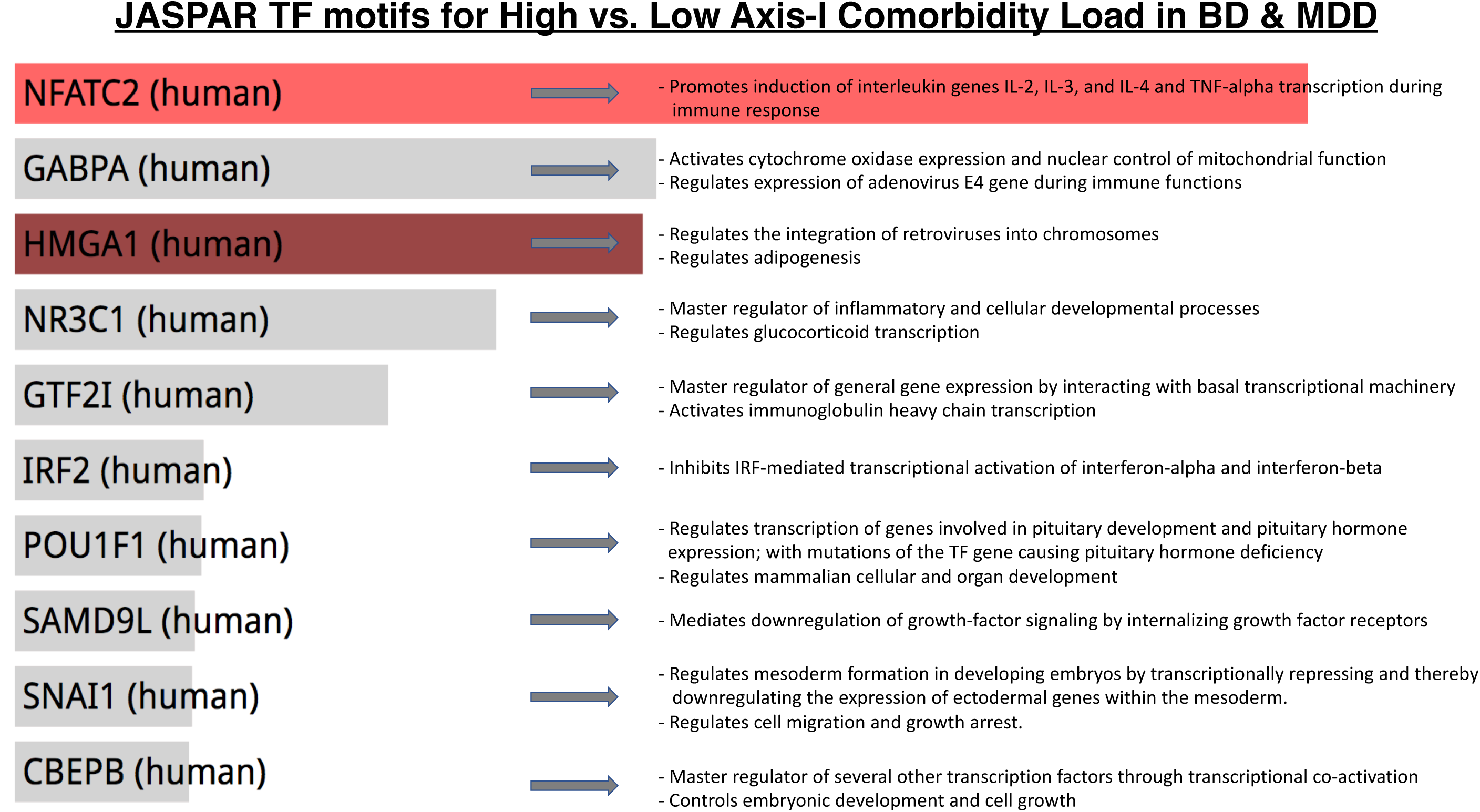
List of TF genes showing JASPAR 2018 identified associated motifs for gene expression profiles for high compared to low Axis-I comorbidity burden in BD and MDD. The identified TFs include putative master transcriptional regulators (NR3C1, GTF2I and CBEPB) as well as regulators of cytokine gene expression (NFATC2 and IRF2), viral mediated functional immune-related gene expression (GABPA and HMGA1), and in regulating early embryonic (SNAI1), cellular (SAMD9L) and hormonal development (POU1F1).

Specifically, comparing BD vs. controls revealed differentially expressed genes with associated motifs for TFs involved in regulating *a*) immune response and antigen processing (i.e. *MIB2*), *b*) master transcriptional repression and activation of a wide range of genes (i.e. *ZBTB7A*), *c*) Zinc Finger transcriptional control and cellular developmental processes and apoptosis (i.e. *ZBTB7A, MZF1, KLF11, ZNF148*), *d*) post-translational modification of gene expression and ion channel transporter expression (i.e. *SMAD4, MIR210, Esrrb*) and *e*) Hippo-signaling pathway, organ size control and regulation of cell proliferation and apoptosis (i.e. *TEAD2, TEAD4*) (***Figure 2A***). Comparing MDD vs. controls revealed differentially expressed genes with associated motifs for TFs involved in regulating *a*) post-translational modification of gene expression and ion channel transporter expression (i.e. *MIR210, Esrrb*) and *b*) Hippo-signaling pathway, organ size control and regulation of cell proliferation and apoptosis (i.e. *TEAD2, TEAD4*), *c*) Zinc Finger G Protein-Coupled receptor binding TF activity (i.e. *SP4, SP3, SP1*) implicated in mood disorders (Zhou et al. 2009; Shi et al. 20011; Shyn et al. 2011), *d*) TF super families involved in embryonic development, activation or repression of transcriptional activity of different sets of genes and RNA-binding (i.e. *TFAP2A, PCBP1, HNF1B*) (***Figure 2B***).

Restricting our analysis to comparing mood disorder donors with high Axis-I comorbidity vs. mood disorder donors scoring low on Axis-I comorbidity, we found differentially expressed genes with associated motifs for TFs involved in regulating *a*) inflammatory and immune response and toll-like receptor signaling, *b*) embryonic and cellular developmental processes, and *c*) cellular and peripheral homeostatic control (see ***Figure 3*** for specific regulatory functions for each of the identified TFs). When comparing DEGs in high vs. low Axis-I comorbidity in mood disorders, we found 4 motifs for master TFs implicated in regulation an array of functions including general transcriptional regulation through interaction with basal transcriptional machinery (i.e. *G2FTI*), master transcriptional co-activation of several other TFs (i.e. *CREBBP*), and a master TF that regulates inflammatory, cellular developmental, and glucocorticoid gene expression processes (i.e. *NR3C1* and *POU1F1*) (see ***Figure 3*** for specific regulatory functions for each of the identified TFs). Specifically, the *GTF2i* TF gene has been genetically associated with the regulation of brain-mediated affective, anxiety and other neurocognitive functions (Jabbi et al 2015; Procyshyn et al. 2017; Stein et al. 2017; Roy 2017; Jabbi et al. 2019) and in embryonic and neurodevelopment (Enkmandakh et al. 2009; Roy 2017; Barak et al. 2019). *CREBBP* has been earlier found to be involved in depressive illness and treatment response (Young et al. 2002; Cristafulli et al 2012; Cho et al. 2019), whereas *POU1F1* (Gerritsen et al. 2017), and especially *NR3C1* have been implicated in an array of regulatory functions and genetically associated with glucocorticoid functions, and mediating the interplay between early environmental adversity experience and the etiology of mood illnesses (Kundakovic et al. 2007; Lewis et al. 2012; Mandelli & Serretti 2013; Perroud et al. 2014; Smart et al. 2015; Roy et al. 2017; Keller et al. 2017; Farrell et al. 2018; Peng et al. 2018; Duffy et al. 2019).

## DISCUSSION

Understanding the molecular substrates for regional brain abnormalities underlying major mental illness such as mood disorders, which affect over 10% of the population across the lifetime in the US (Collins et a. 2011), will not only contribute to better mechanistic understanding of the neurobiological bases for behavioral pathologies (Nestler 2015; Sahin & Sur Science 2015; Torre-Ubieta et al. 2018; Jabbi & Nemeroff 2019), but will also be essential for novel drug design (Changeaux 2017).

In the present study, we applied *robust imaging-guided transcriptomics* to target neuroimaging identified GMV reductions in the anterior insula associated with mood and depressive disorder diagnoses (Goodkind et al. 2015; Jabbi et al. 2020) for dissection and subsequent RNA-seq analysis in postmortem donors. This novel approach allows the precise molecular targeting of a localized anterior insula region that has been found to exhibit the strongest degree of gray matter loss in mood disorders (Jabbi et al. 2020). This *robust imaging-guided transcriptomics* study therefore integrated RNA-seq characterization of gene expression abnormalities in an anatomically abnormal region in mood disorders and allowed testing of the hypothesis that transcription factor motifs will be associated with differential gene expression profiles in the postmortem anterior insula cortex of mood disorder subjects relative to controls.

We found that comparing BD vs. controls revealed DEGs with associated motifs for TFs involved in regulating immune response and antigen processing, master transcriptional repression/activation of a wide range of genes, Zinc Finger transcriptional control of cellular developmental processes, post-translational modification of gene expression and ion channel transporter expression and Hippo-signaling pathway involved in organ size control and regulation of cellular development. Comparing MDD vs. controls revealed DEGs with associated motifs for TFs involved in regulating post-translational modification of gene expression and ion channel transporter expression and Hippo-signaling pathway, Zinc Finger G Protein-Coupled receptor binding TF activity that has been implicated in mood disorders (Zhou et al. 2009; Shi et al. 20011; Shyn et al. 2011), and TF super families involved in embryonic and cellular development. DEGs in high vs. low Axis-I comorbidity revealed motifs for master TFs implicated in the regulation of an array of functions including general transcriptional regulation through interaction with basal transcriptional machinery, and in inflammatory, cellular developmental, and glucocorticoid gene expression processes. Of interest, the two comparisons between BD vs. controls and MDD vs. controls identified TF motifs of similar pathways including four TFs (i.e. *MIR120, TEAD2, TEAD4* and *Esrrb*) suggesting possible specific as well as overlapping TF motifs for BD & MDD. If replicated, the identified TF motifs associated with DEGs in mood disorder diagnoses and overall disease morbidity will likely reveal a vulnerability in the general transcriptional pathway mechanism in mood disorder disease states.

The identified DEG-profile associated TF motifs are known to regulate expression of toll-like receptor signaling genes, cellular homeostatic control genes, and embryonic and cellular including neurodevelopmental gene networks found to be differentially expressed in BD vs. controls, MDD vs. controls, and in high Axis-I comorbid mood disorder individuals vs. low Axis-I comorbidity mood disorder individuals. We found putative hierarchical TF regulatory involvement in the gene expression landscapes associated with BD and MDD diagnoses, such that enrichments of master transcription factors were predominantly associated with the gene expression landscape in elevated mood disorder morbidity, thereby echoing previous work (Tsigelny et al. 2013; Lee & Young 2013; Changeaux 2017; Lambert et al. 2018; Chen C et al. 2019). It is important to note that a number of the TFs we have associated with anterior insula gene expression in bipolar disorder vs. controls (i.e., MIB2, Esrrb, MZF1, and ZNF148) and in major depressives vs. controls (i.e., PCP1, HNF1B, and Esrrb) are novel in that they have not been previously been identified in any genetic screens. Interestingly, many of the TFs we have identified have been implicated in neural development: TEAD2 and TEAD4 appear to play a role in cortical development (Mukhtar et al. 2020). Furthermore, KLF11 (Duncan et al. 2012; Duncan et al. 2015; Harris et al. 2015; Kollert et al. 2020), SP1 (Brewer et al. 2004; Tamayo et al. 2009; Pinacho et al. 2011; Fuste et al. 2013; Saucedo-Uribe et al. 2019; Hung et al. 2020), and SP4 (Zou et al. 2009; Fuste et al. 2013; Young et al. 2015) have been found to be involved in the genetic regulation of cell-fate and apoptosis as well as in mood and related psychiatric disorders. And SMAD2 has been linked to antidepressant activity (Dow et al. 2005) in addition to neuronal development (Zhang et al. 2013). Finally, our results are in line with recent postmortem studies of other brain areas in mood disorder morbidity that found disease-related gene expression changes in immune and inflammatory signaling (Pantazatos et al. 2007; Darby et al 2016; Wohleb et al. 2016; Pacifico et al. 2017; Pandey et al. 2017; Nagy et al. 2020), cellular homeostatic and synaptic signaling (Pantazatos et al 2007; Sequeira et al. 2007; Sequeira et al. 2009; Kang et al. 2012; Li et al. 2013; Gray et al. 2015; Yin et al. 2016; Labonte et al. 2017; Zhao et al. 2018), and in cellular and neurodevelopmental signaling (Paterson et al. 2017).

Conventional analyses of differential gene expression differences related to a specific phenotype of interest does not account for the hierarchical nature of gene expression regulation, e.g., that TFs regulate other gene classes. Our current identification of mood disorder DEG associated TF motifs known to regulate the expression of toll-like receptor signaling pathway genes, cellular homeostatic control genes, and embryonic as well as cellular and neurodevelopment pathway genes in mood disorder diagnostic phenotypes, suggests a hierarchical involvement of the identified TFs in the neurobiological abnormalities underlying mood disorder phenotypes.

Despite its obvious strengths, our use of novel *imaging-guided transcriptomics* approach to identify TF motifs associated with gene expression in mood disorders has several limitations. *First*, we provide statistical associations rather than direct testing of the causal role of the identified TF motifs in *1*) the developmental or disease-driven emergence of the anatomical gray matter loss identified in the anterior insula of mood disorder individuals, and *2*) the observed postmortem differential gene expression landscape measured within the anterior insula in mood disorder morbidity. *Second*, anatomical abnormalities in the anterior insula are not specific to mood disorder diagnoses *per se*, but involved in several neuropsychiatric diagnoses including anxiety, psychosis, substance use and eating disorders (Goodkind et al. 2016; Janiri et al. 2019), suggesting the importance of including other more mood disorder-specific regional anatomical brain abnormalities in future transcriptomic studies. However, we specifically targeted the anterior insula subsection found to be the most reduced in GMV measures across the entire brain in mood disorders in our meta-analysis with the expectation that this locus of anatomical abnormality likely harbors important transcriptomic clues to mood disorder molecular abnormalities. *Third*, our approach does not allow us to confirm whether the postmortem mood disorder samples we examined also showed a reduction in anterior insula GMV. Future studies need to find a way to estimate both anatomical and gene expression abnormalities in the same postmortem brain samples across brain regions to directly link anatomical abnormalities with underlying molecular genetic regulatory abnormalities in the same postmortem brains. *Fourth*, in light of the limited number of samples for a study of complex biological variables, and the fact that the TF data are secondary derivations from the differential gene expression analyses, our results could be confounded by factors between cases and controls, despite our attempts to control for such factors. *Fifth*, bulk tissue RNA-seq cannot account for the role of specific cell-types in defining TF regulatory involvements in mood disorder transcriptomics. Single cell RNA-seq analyses in specific brain regions, ideally across the lifespan, will greatly advance our understanding of the cell-type specific role of the identified TFs in the pathogenesis of mood disorders. *Finally*, including a comparison psychiatric cohort without mood symptoms such as schizophrenia in future studies will help address the specificity of the identified TF targets for mood disorder therapeutics. These limitations notwithstanding, our current focus on the anterior insula given its identification as the region exhibiting the most extensive gray matter loss associated with mood disorder diagnoses, provides a framework for integrative translational studies of anatomical and molecular abnormalities underlying prevalent brain diseases.

In conclusion, we applied *robust imaging-guided transcriptomics* to characterize the roles of specific TFs in mood disorders. Our study provides important initial insights into the molecular pathways, and the relevant TFs that may be regulating them, in the context of mood disorders and related psychiatric diseases. Our results suggest that the postmortem gene expression patterns we observed in the anterior insula of mood disorder donors are associated with (*1*) the magnitude of mood disorder morbidity in a putative anatomically compromised brain region in mood disorders; and (*2*) specific TFs known to regulate broad transcriptional processes, immune response, cellular homeostasis, embryonic and neuronal development, and the etiology of mood disorders. Together, these findings illustrate that studies of gene regulatory networks have the potential to elucidate the hierarchical organizational principles of the gene expression landscapes driving major psychiatric disorders, and thereby accelerate novel pharmacological target discovery.

## Author Contributions

MJ conceived and designed the studies, and acquired postmortem material from the NIMH HBCC. DA, SBE, and MJ performed the experiments and analyzed the data and results. DA and MJ drafted the manuscript and SBE, CBN, and HH contributed critically and substantially to the content/writing of the manuscript and interpretation of the findings.

## Acknowledgements

The NIMH Human Brain Collection Core provided RNA-samples for donors and we thank the NIMH and Drs. Barbara Lipska, Stefano Marenco, Pavan Auluck and HBCC colleagues for the studied samples. We thank Wade Weber of Dell Medical School Psychiatry Department, UT Austin for assistance in preparing the manuscript, Dr. Mark Bond of Dell Medical School Psychiatry Department, UT Austin for statistical reviews, and Jessica Podnar and several GSAF colleagues for RNA-seq support.

## Competing Interests

Mbemba Jabbi, none.

Dhivya Arasappan, none.

Simon Eickhoff none.

Hans Hofmann, none.

Charles B Nemeroff:

Research/Grants: National Institutes of Health (NIH)

Consulting (last year): Taisho Pharmaceutical, Inc., Signant Health, Sunovion Pharmaceuticals, Inc., Janssen Research & Development LLC, Magstim, Inc., Navitor Pharmaceuticals, Inc., Intra-Cellular Therapies, Inc., EMA Wellness, Acadia Pharmaceuticals, Axsome, Sage, BioXcel Therapeutics, Silo Pharma, Aditum Bio

Stockholder: Xhale, Seattle Genetics, Antares, BI Gen Holdings, Inc., Corcept Therapeutics Pharmaceuticals Company, EMA Wellness

Scientific Advisory Boards: Brain and Behavior Research Foundation (BBRF), Anxiety Disorders Association of America (ADAA), Skyland Trail, Signant Health, Laureate Institute for Brain Research (LIBR), Inc., Magnolia CNS

Board of Directors: Gratitude America, ADAA, Xhale Smart, Inc.

Income sources or equity of $10,000 or more: American Psychiatric Publishing, Xhale, Signant Health, CME Outfitters, Takeda, Intra-Cellular Therapies, Inc., EMA Wellness

Patents: Method and devices for transdermal delivery of lithium (US 6,375,990B1) Method of assessing antidepressant drug therapy via transport inhibition of monoamine neurotransmitters by ex vivo assay (US 7,148,027B2)

Speakers Bureau: None

## Table 1 Legend

Abbreviations: MDD, major depressive disorder; GMV, gray matter volume; AIC, anterior insula cortex; PMI, postmortem index; Ph, measure of acidity; RIN, RNA integrity number which is a measure of RNA quality.

## Supplementary Table 2-4 Legend

Abbreviations: baseMean, normalized read counts of all samples; log2Foldchange, effect size estimate; lfcSE, standard error of the log2foldchange; stat, Wald statistical test values; pvalue, uncorrected p-value; p-adj, corrected p-value. Fold change is calculated as the ratio of mean expression in high psychiatric morbidity & suicide mortality score to the mean expression in low psychiatric morbidity & suicide mortality score.

