## Supplementary Materials for "Transcription factor motifs associated with anterior insula gene-expression underlying mood disorder phenotypes"

**Supplementary Tables**

**Supplementary Table 1 Diagnoses**

|  | **Primary Diagnoses** |  |  |
| --- | --- | --- | --- |
| **Axis-I Load** | BD, n=37 | MDD, n=30 | Controls, n=33 |
| Polysubstance Use Disorder/Abuse | 12 | 10 | 0 |
| Alcohol Use Disorder/Abuse | 15 | 3 | 0 |
| Psychosis/Psychotic | 14 | 6 | 0 |
| Most Recent Episode = Depressed/MDD Recurrent | 21 | 26 | 0 |
| Specified/unspecified anxiety disorders | 5 | 4 | 0 |

**Supplementary Table 2 RNA-seq in BD vs Controls**

| Gene Ensemble ID | baseMean | log2FoldChange | lfcSE | stat | pvalue | padj |
| --- | --- | --- | --- | --- | --- | --- |
| ENSG00000124440.15 | 618.8567348 | 0.963640201 | 0.1918 | 5.02358 | 5.07E-07 | 0.00088 |
| ENSG00000187193.8 | 681.5193475 | 1.761186273 | 0.3466 | 5.08204 | 3.73E-07 | 0.00088 |
| ENSG00000225972.1 | 2253.039098 | -2.832747638 | 0.5483 | -5.1664 | 2.39E-07 | 0.00088 |
| ENSG00000183688.4 | 252.081524 | 0.538349335 | 0.1146 | 4.69667 | 2.64E-06 | 0.00345 |
| ENSG00000155380.11 | 741.7046199 | 0.618686889 | 0.1409 | 4.39053 | 1.13E-05 | 0.00739 |
| ENSG00000160801.13 | 213.937388 | 0.506308291 | 0.1148 | 4.4097 | 1.04E-05 | 0.00739 |
| ENSG00000167772.11 | 436.9239165 | 1.728915437 | 0.3902 | 4.43077 | 9.39E-06 | 0.00739 |
| ENSG00000198886.2 | 482968.542 | 0.707747893 | 0.1578 | 4.48619 | 7.25E-06 | 0.00739 |
| ENSG00000239282.7 | 134.2616841 | 0.668837237 | 0.1536 | 4.35549 | 1.33E-05 | 0.00771 |
| ENSG00000117394.19 | 1729.789884 | 0.486160403 | 0.1129 | 4.30723 | 1.65E-05 | 0.00785 |
| ENSG00000248050.1 | 117.9609801 | -0.508028662 | 0.1178 | -4.3137 | 1.61E-05 | 0.00785 |
| ENSG00000168394.10 | 316.7067126 | 0.598704662 | 0.1414 | 4.23538 | 2.28E-05 | 0.00994 |
| ENSG00000135750.14 | 1775.107969 | -0.423532959 | 0.1025 | -4.1329 | 3.58E-05 | 0.0144 |
| ENSG00000133048.12 | 428.5115721 | 1.141468155 | 0.2783 | 4.10214 | 4.09E-05 | 0.01476 |
| ENSG00000138772.12 | 160.1495313 | 0.655455502 | 0.1607 | 4.07917 | 4.52E-05 | 0.01476 |
| ENSG00000163884.3 | 513.5237656 | 0.501424979 | 0.1227 | 4.08562 | 4.40E-05 | 0.01476 |
| ENSG00000107551.20 | 1312.906855 | 0.342789897 | 0.0853 | 4.01731 | 5.89E-05 | 0.0181 |
| ENSG00000111077.17 | 1246.912152 | 0.386044021 | 0.0982 | 3.93051 | 8.48E-05 | 0.02332 |
| ENSG00000164292.12 | 2138.928979 | 0.612705959 | 0.1559 | 3.93064 | 8.47E-05 | 0.02332 |
| ENSG00000102935.11 | 528.2981002 | 0.391030572 | 0.101 | 3.87223 | 0.000108 | 0.02418 |
| ENSG00000112312.9 | 156.2709052 | 0.490640748 | 0.1276 | 3.84548 | 0.00012 | 0.02418 |
| ENSG00000115255.10 | 259.0543362 | 0.345281346 | 0.0887 | 3.8931 | 9.90E-05 | 0.02418 |
| ENSG00000160766.14 | 425.3136999 | -0.421247379 | 0.1081 | -3.8964 | 9.76E-05 | 0.02418 |
| ENSG00000168209.4 | 726.3332552 | 0.722729665 | 0.1871 | 3.86222 | 0.000112 | 0.02418 |
| ENSG00000213563.6 | 381.188925 | 0.321367962 | 0.0835 | 3.84972 | 0.000118 | 0.02418 |
| ENSG00000214456.8 | 173.7735687 | 0.621219435 | 0.1611 | 3.85708 | 0.000115 | 0.02418 |
| ENSG00000111275.12 | 5514.54366 | 0.372225112 | 0.0981 | 3.79568 | 0.000147 | 0.02482 |
| ENSG00000125144.13 | 515.0288621 | 0.839840643 | 0.221 | 3.79942 | 0.000145 | 0.02482 |
| ENSG00000163395.16 | 154.6436183 | 0.765116394 | 0.2019 | 3.78885 | 0.000151 | 0.02482 |
| ENSG00000163702.18 | 305.7218214 | 0.423557752 | 0.1115 | 3.79849 | 0.000146 | 0.02482 |
| ENSG00000184154.13 | 274.0279939 | 0.238282685 | 0.0629 | 3.7878 | 0.000152 | 0.02482 |
| ENSG00000204396.10 | 294.6973179 | -0.456159354 | 0.1196 | -3.8151 | 0.000136 | 0.02482 |
| ENSG00000134508.12 | 1844.148261 | 0.553796298 | 0.1481 | 3.73837 | 0.000185 | 0.02933 |
| ENSG00000100106.19 | 399.6725301 | 0.332240245 | 0.0893 | 3.72241 | 0.000197 | 0.03004 |
| ENSG00000142733.14 | 221.915313 | 0.484414692 | 0.1306 | 3.7103 | 0.000207 | 0.03004 |
| ENSG00000196136.16 | 139.1643261 | 2.19111007 | 0.5896 | 3.71652 | 0.000202 | 0.03004 |
| ENSG00000205364.3 | 165.180011 | 1.084882983 | 0.2929 | 3.70348 | 0.000213 | 0.03004 |
| ENSG00000185745.9 | 988.2475227 | -0.261896671 | 0.0711 | -3.6845 | 0.000229 | 0.03097 |
| ENSG00000259291.2 | 1757.61328 | 0.28164587 | 0.0765 | 3.68229 | 0.000231 | 0.03097 |
| ENSG00000134115.12 | 344.7228445 | -0.446208669 | 0.1217 | -3.6675 | 0.000245 | 0.032 |
| ENSG00000254245.2 | 217.1909948 | 0.716294576 | 0.1961 | 3.65293 | 0.000259 | 0.03305 |
| ENSG00000136235.15 | 571.7448926 | 0.743646179 | 0.2051 | 3.62517 | 0.000289 | 0.03593 |
| ENSG00000088826.17 | 676.5691445 | 0.503253705 | 0.1391 | 3.61823 | 0.000297 | 0.03605 |
| ENSG00000225630.1 | 518.8572599 | 0.916401344 | 0.2545 | 3.60042 | 0.000318 | 0.03773 |
| ENSG00000078401.6 | 207.0370193 | 0.809790005 | 0.2271 | 3.56559 | 0.000363 | 0.03887 |
| ENSG00000099840.13 | 185.7621813 | 0.370272161 | 0.1034 | 3.58032 | 0.000343 | 0.03887 |
| ENSG00000121207.11 | 163.1211115 | 0.470276814 | 0.1317 | 3.57047 | 0.000356 | 0.03887 |
| ENSG00000176244.6 | 700.4203421 | 0.562162627 | 0.1577 | 3.56456 | 0.000364 | 0.03887 |
| ENSG00000259823.5 | 182.6148087 | -0.501280269 | 0.14 | -3.5803 | 0.000343 | 0.03887 |
| ENSG00000043355.11 | 231.8283677 | 0.507563503 | 0.1436 | 3.53521 | 0.000407 | 0.04074 |
| ENSG00000103742.11 | 211.9033035 | 0.493144986 | 0.1396 | 3.53154 | 0.000413 | 0.04074 |
| ENSG00000132541.10 | 675.6970467 | 0.371373549 | 0.105 | 3.53728 | 0.000404 | 0.04074 |
| ENSG00000198888.2 | 329578.949 | 0.485108269 | 0.137 | 3.54117 | 0.000398 | 0.04074 |
| ENSG00000105499.13 | 783.0291304 | -0.285481662 | 0.0812 | -3.5154 | 0.000439 | 0.04098 |
| ENSG00000157502.13 | 462.7888844 | -0.460142954 | 0.1309 | -3.5154 | 0.000439 | 0.04098 |
| ENSG00000212907.2 | 39665.08564 | 0.608061415 | 0.1729 | 3.51671 | 0.000437 | 0.04098 |
| ENSG00000185220.11 | 346.4447315 | 0.297865031 | 0.0854 | 3.48716 | 0.000488 | 0.04476 |
| ENSG00000185201.16 | 241.0811026 | 0.840441354 | 0.2429 | 3.46068 | 0.000539 | 0.04855 |
| ENSG00000197121.14 | 2179.163626 | -0.331653165 | 0.0962 | -3.4491 | 0.000562 | 0.04981 |
| ENSG00000135245.9 | 258.5582349 | 0.931965886 | 0.271 | 3.43908 | 0.000584 | 0.05084 |
| ENSG00000099998.17 | 237.366125 | 0.503758258 | 0.1477 | 3.41178 | 0.000645 | 0.05155 |
| ENSG00000106003.12 | 233.8560245 | 0.628800375 | 0.1842 | 3.41443 | 0.000639 | 0.05155 |
| ENSG00000175287.18 | 226.9126389 | 0.580292789 | 0.1695 | 3.42389 | 0.000617 | 0.05155 |
| ENSG00000185596.16 | 273.741632 | 0.359185616 | 0.1054 | 3.40943 | 0.000651 | 0.05155 |
| ENSG00000198899.2 | 163086.25 | 0.46863008 | 0.1372 | 3.41571 | 0.000636 | 0.05155 |
| ENSG00000279457.3 | 320.2683895 | 0.320832374 | 0.0938 | 3.42003 | 0.000626 | 0.05155 |
| ENSG00000146535.13 | 1983.366475 | 0.374478172 | 0.1101 | 3.40007 | 0.000674 | 0.05208 |
| ENSG00000150054.18 | 452.2068091 | -0.294008327 | 0.0865 | -3.3985 | 0.000678 | 0.05208 |
| ENSG00000070882.12 | 796.762601 | -0.314757713 | 0.0929 | -3.3885 | 0.000703 | 0.05323 |
| ENSG00000096060.14 | 713.0093842 | 0.746671091 | 0.2235 | 3.3404 | 0.000837 | 0.05465 |
| ENSG00000106477.18 | 596.6628089 | -0.232902607 | 0.0691 | -3.3708 | 0.00075 | 0.05465 |
| ENSG00000144649.8 | 119.7836749 | 0.538198242 | 0.161 | 3.34324 | 0.000828 | 0.05465 |
| ENSG00000145632.14 | 2836.925791 | -0.277907273 | 0.0832 | -3.342 | 0.000832 | 0.05465 |
| ENSG00000159459.11 | 1185.200581 | -0.188446867 | 0.0563 | -3.3485 | 0.000812 | 0.05465 |
| ENSG00000162496.8 | 331.1849718 | 0.384516845 | 0.1141 | 3.36901 | 0.000754 | 0.05465 |
| ENSG00000163449.10 | 260.7193655 | -0.350893525 | 0.1048 | -3.3477 | 0.000815 | 0.05465 |
| ENSG00000169203.16 | 1185.179393 | 0.421863657 | 0.1253 | 3.36809 | 0.000757 | 0.05465 |
| ENSG00000189221.9 | 962.2013909 | 0.288573298 | 0.0861 | 3.35119 | 0.000805 | 0.05465 |
| ENSG00000213366.12 | 1105.371028 | 0.294676082 | 0.0876 | 3.36318 | 0.000771 | 0.05465 |
| ENSG00000280234.1 | 255.7622779 | 0.332303207 | 0.0992 | 3.34857 | 0.000812 | 0.05465 |
| ENSG00000116962.14 | 204.9825137 | 0.582264456 | 0.1749 | 3.32897 | 0.000872 | 0.0552 |
| ENSG00000161958.10 | 196.5969166 | 0.463021659 | 0.1392 | 3.32711 | 0.000878 | 0.0552 |
| ENSG00000176974.18 | 150.9680719 | 0.308871787 | 0.0929 | 3.32382 | 0.000888 | 0.0552 |
| ENSG00000179820.15 | 1528.672329 | -0.242094512 | 0.0729 | -3.3207 | 0.000898 | 0.0552 |
| ENSG00000185033.14 | 724.1276432 | 0.343981661 | 0.1035 | 3.32444 | 0.000886 | 0.0552 |
| ENSG00000004799.7 | 770.3405685 | 0.925337047 | 0.2801 | 3.30341 | 0.000955 | 0.05705 |
| ENSG00000075073.14 | 127.5637483 | -0.407898217 | 0.1255 | -3.2504 | 0.001153 | 0.05705 |
| ENSG00000103319.11 | 759.7983042 | 0.301734976 | 0.0929 | 3.24922 | 0.001157 | 0.05705 |
| ENSG00000113161.15 | 2933.177234 | -0.345293149 | 0.1061 | -3.2538 | 0.001139 | 0.05705 |
| ENSG00000118242.15 | 269.2096791 | -0.333783385 | 0.1019 | -3.2755 | 0.001055 | 0.05705 |
| ENSG00000119711.12 | 3581.785212 | 0.460427413 | 0.1414 | 3.2573 | 0.001125 | 0.05705 |
| ENSG00000129353.14 | 2056.90083 | 0.318590614 | 0.0972 | 3.27637 | 0.001052 | 0.05705 |
| ENSG00000138604.9 | 1079.392739 | -0.317200201 | 0.0969 | -3.2741 | 0.00106 | 0.05705 |
| ENSG00000140905.9 | 947.4464287 | 0.255007972 | 0.0777 | 3.28394 | 0.001024 | 0.05705 |
| ENSG00000141736.13 | 387.5378855 | 0.480859373 | 0.1467 | 3.27687 | 0.00105 | 0.05705 |
| ENSG00000148411.7 | 3360.923118 | 0.37883269 | 0.1166 | 3.24955 | 0.001156 | 0.05705 |
| ENSG00000156097.12 | 388.8113414 | -0.399141387 | 0.121 | -3.2999 | 0.000967 | 0.05705 |
| ENSG00000165458.13 | 969.0932357 | 0.312742225 | 0.096 | 3.25632 | 0.001129 | 0.05705 |
| ENSG00000166839.16 | 652.0832585 | 0.275827695 | 0.0835 | 3.30431 | 0.000952 | 0.05705 |
| ENSG00000167315.17 | 613.8193974 | 0.423991166 | 0.1299 | 3.26331 | 0.001101 | 0.05705 |
| ENSG00000183580.9 | 373.112041 | 0.473234442 | 0.1451 | 3.26196 | 0.001106 | 0.05705 |
| ENSG00000196091.13 | 625.0857273 | 0.453599778 | 0.138 | 3.28641 | 0.001015 | 0.05705 |
| ENSG00000196756.11 | 135.4470966 | -0.357316623 | 0.1088 | -3.2832 | 0.001026 | 0.05705 |
| ENSG00000198695.2 | 26338.86319 | 0.575553857 | 0.1762 | 3.26584 | 0.001091 | 0.05705 |
| ENSG00000216775.2 | 150.5820377 | -0.324271904 | 0.0993 | -3.2653 | 0.001094 | 0.05705 |
| ENSG00000228109.1 | 116.9289105 | 0.299217007 | 0.092 | 3.25262 | 0.001143 | 0.05705 |
| ENSG00000075234.16 | 143.7534385 | 0.433444697 | 0.1348 | 3.2161 | 0.001299 | 0.05811 |
| ENSG00000076351.12 | 478.146586 | 0.215822647 | 0.0668 | 3.23235 | 0.001228 | 0.05811 |
| ENSG00000103335.20 | 667.7085261 | 0.348481089 | 0.1077 | 3.23702 | 0.001208 | 0.05811 |
| ENSG00000105552.14 | 305.6231345 | 0.265961908 | 0.0833 | 3.19213 | 0.001412 | 0.05811 |
| ENSG00000108604.15 | 280.2442816 | 0.256710143 | 0.0796 | 3.22502 | 0.00126 | 0.05811 |
| ENSG00000112333.11 | 468.4648489 | 0.513640288 | 0.1594 | 3.22187 | 0.001274 | 0.05811 |
| ENSG00000122971.8 | 220.0896329 | 0.418038871 | 0.13 | 3.21691 | 0.001296 | 0.05811 |
| ENSG00000129128.12 | 2411.292722 | -0.253638856 | 0.0789 | -3.2135 | 0.001311 | 0.05811 |
| ENSG00000132164.9 | 871.67119 | 0.403107038 | 0.1252 | 3.2198 | 0.001283 | 0.05811 |
| ENSG00000153234.13 | 1047.746983 | -0.518575882 | 0.16 | -3.2405 | 0.001193 | 0.05811 |
| ENSG00000166033.11 | 4072.375741 | 0.411115378 | 0.1275 | 3.224 | 0.001264 | 0.05811 |
| ENSG00000168884.14 | 157.4169547 | 0.358262825 | 0.1122 | 3.19446 | 0.001401 | 0.05811 |
| ENSG00000169715.14 | 697.4279137 | 0.711426615 | 0.2225 | 3.19729 | 0.001387 | 0.05811 |
| ENSG00000180914.10 | 193.4193108 | 0.509236426 | 0.1595 | 3.19263 | 0.00141 | 0.05811 |
| ENSG00000182902.13 | 1873.474604 | 0.531625228 | 0.1665 | 3.19309 | 0.001408 | 0.05811 |
| ENSG00000198157.10 | 257.6019508 | 0.242082877 | 0.0758 | 3.19456 | 0.0014 | 0.05811 |
| ENSG00000198786.2 | 164068.398 | 0.457014407 | 0.1429 | 3.19854 | 0.001381 | 0.05811 |
| ENSG00000204219.9 | 123.3731876 | 0.311140745 | 0.0974 | 3.19326 | 0.001407 | 0.05811 |
| ENSG00000224877.3 | 502.5295091 | 0.29589187 | 0.092 | 3.21732 | 0.001294 | 0.05811 |
| ENSG00000271447.5 | 276.1379582 | 0.497418316 | 0.1558 | 3.19354 | 0.001405 | 0.05811 |
| ENSG00000279672.1 | 163.3355491 | 0.54835873 | 0.1708 | 3.21099 | 0.001323 | 0.05811 |
| ENSG00000072858.10 | 913.0443961 | -0.253127446 | 0.0799 | -3.1687 | 0.001531 | 0.05842 |
| ENSG00000106689.10 | 1357.862805 | 0.32634786 | 0.1026 | 3.1813 | 0.001466 | 0.05842 |
| ENSG00000123570.3 | 793.3744933 | -0.245981344 | 0.0772 | -3.1869 | 0.001438 | 0.05842 |
| ENSG00000139800.8 | 119.7312918 | 0.408403792 | 0.1289 | 3.16863 | 0.001532 | 0.05842 |
| ENSG00000165475.13 | 800.610704 | 0.255567759 | 0.0806 | 3.17211 | 0.001513 | 0.05842 |
| ENSG00000167676.4 | 142.6468004 | 0.763192184 | 0.2403 | 3.17658 | 0.00149 | 0.05842 |
| ENSG00000168913.6 | 2485.284178 | 0.360429578 | 0.1134 | 3.17843 | 0.001481 | 0.05842 |
| ENSG00000170571.11 | 180.606981 | -0.419422386 | 0.1322 | -3.1717 | 0.001516 | 0.05842 |
| ENSG00000186994.11 | 160.7712968 | 0.356450435 | 0.1124 | 3.17083 | 0.00152 | 0.05842 |
| ENSG00000204264.8 | 132.3089305 | 0.386997309 | 0.1217 | 3.18123 | 0.001467 | 0.05842 |
| ENSG00000143545.8 | 384.2742479 | 0.451266646 | 0.1426 | 3.16471 | 0.001552 | 0.05879 |
| ENSG00000184313.19 | 375.1122858 | 0.325345574 | 0.1031 | 3.15511 | 0.001604 | 0.06032 |
| ENSG00000111321.10 | 170.6177297 | 0.405723137 | 0.1294 | 3.13517 | 0.001718 | 0.06081 |
| ENSG00000115252.18 | 2221.961333 | -0.313893075 | 0.0999 | -3.1419 | 0.001678 | 0.06081 |
| ENSG00000121742.16 | 824.4811766 | 0.806329681 | 0.2574 | 3.13252 | 0.001733 | 0.06081 |
| ENSG00000134201.10 | 517.5464543 | 0.69830488 | 0.2233 | 3.12738 | 0.001764 | 0.06081 |
| ENSG00000147852.15 | 913.2434299 | -0.22392206 | 0.0717 | -3.1227 | 0.001792 | 0.06081 |
| ENSG00000148498.15 | 312.5271537 | 0.43909375 | 0.1399 | 3.13776 | 0.001702 | 0.06081 |
| ENSG00000149403.11 | 632.0271387 | -0.242075921 | 0.0772 | -3.1366 | 0.001709 | 0.06081 |
| ENSG00000162878.12 | 587.5188408 | -0.292940768 | 0.0931 | -3.1457 | 0.001657 | 0.06081 |
| ENSG00000174498.13 | 146.0108927 | 0.248207131 | 0.0789 | 3.1442 | 0.001665 | 0.06081 |
| ENSG00000184113.9 | 911.1033866 | 0.473131489 | 0.1514 | 3.12584 | 0.001773 | 0.06081 |
| ENSG00000186594.13 | 275.4230067 | -0.323356216 | 0.103 | -3.1389 | 0.001696 | 0.06081 |
| ENSG00000186815.12 | 1269.772668 | 0.269187291 | 0.086 | 3.13008 | 0.001748 | 0.06081 |
| ENSG00000198804.2 | 886768.7381 | 0.446621636 | 0.1428 | 3.12847 | 0.001757 | 0.06081 |
| ENSG00000261604.1 | 165.1236074 | -0.393260426 | 0.125 | -3.1469 | 0.00165 | 0.06081 |
| ENSG00000271430.1 | 331.2626411 | -0.295246035 | 0.0945 | -3.1242 | 0.001783 | 0.06081 |
| ENSG00000087250.8 | 5732.91548 | 0.399554209 | 0.1284 | 3.1128 | 0.001853 | 0.06248 |
| ENSG00000085978.21 | 874.54499 | -0.260538142 | 0.0838 | -3.1104 | 0.001868 | 0.06258 |
| ENSG00000138759.17 | 781.1836725 | -0.367415046 | 0.1183 | -3.1066 | 0.001893 | 0.063 |
| ENSG00000156395.12 | 945.8704865 | -0.281759273 | 0.0909 | -3.1008 | 0.00193 | 0.06343 |
| ENSG00000162836.11 | 264.9424589 | 0.341846421 | 0.1102 | 3.1009 | 0.001929 | 0.06343 |
| ENSG00000159423.16 | 1260.5157 | 0.533704253 | 0.1722 | 3.0986 | 0.001944 | 0.06351 |
| ENSG00000169871.12 | 620.1988045 | 0.353633133 | 0.1143 | 3.09439 | 0.001972 | 0.06368 |
| ENSG00000173227.13 | 1383.109318 | -0.296802444 | 0.0959 | -3.0941 | 0.001974 | 0.06368 |
| ENSG00000175874.9 | 7714.560052 | -0.342803097 | 0.111 | -3.0888 | 0.00201 | 0.06444 |
| ENSG00000105854.12 | 2294.572254 | 0.472857175 | 0.1534 | 3.08319 | 0.002048 | 0.06526 |
| ENSG00000068615.17 | 2402.527791 | -0.283656126 | 0.0922 | -3.0754 | 0.002102 | 0.06579 |
| ENSG00000198727.2 | 258086.4497 | 0.397324379 | 0.1291 | 3.07682 | 0.002092 | 0.06579 |
| ENSG00000198763.3 | 246691.1654 | 0.412509305 | 0.1341 | 3.07632 | 0.002096 | 0.06579 |
| ENSG00000101187.15 | 291.5612067 | 0.669232289 | 0.2179 | 3.07106 | 0.002133 | 0.06606 |
| ENSG00000177133.10 | 640.0626687 | 0.573877139 | 0.1869 | 3.07058 | 0.002136 | 0.06606 |
| ENSG00000257434.1 | 686.3547618 | -0.450400705 | 0.1468 | -3.0689 | 0.002149 | 0.06606 |
| ENSG00000228253.1 | 7343.578608 | 0.507163047 | 0.1654 | 3.06651 | 0.002166 | 0.06619 |
| ENSG00000147119.3 | 205.3545967 | 0.457828203 | 0.1499 | 3.05475 | 0.002252 | 0.06804 |
| ENSG00000183773.15 | 1332.415106 | 0.384051657 | 0.1257 | 3.05483 | 0.002252 | 0.06804 |
| ENSG00000179388.8 | 2404.875323 | -0.315452825 | 0.1033 | -3.0528 | 0.002267 | 0.06808 |
| ENSG00000120738.7 | 2494.47022 | -0.641724676 | 0.2106 | -3.0472 | 0.00231 | 0.06819 |
| ENSG00000124570.17 | 749.190428 | 0.259729301 | 0.0852 | 3.04787 | 0.002305 | 0.06819 |
| ENSG00000170340.10 | 503.8812784 | -0.283338232 | 0.093 | -3.0478 | 0.002305 | 0.06819 |
| ENSG00000093010.12 | 992.5632863 | 0.238889118 | 0.0788 | 3.03239 | 0.002426 | 0.06898 |
| ENSG00000105143.12 | 628.97225 | -0.269081998 | 0.0886 | -3.0366 | 0.002393 | 0.06898 |
| ENSG00000142089.15 | 1188.421832 | 0.669663255 | 0.2208 | 3.03257 | 0.002425 | 0.06898 |
| ENSG00000144908.13 | 1214.86038 | 0.617141012 | 0.2036 | 3.03161 | 0.002433 | 0.06898 |
| ENSG00000149499.11 | 428.2224056 | 0.348895181 | 0.115 | 3.03372 | 0.002416 | 0.06898 |
| ENSG00000173166.17 | 1627.728747 | -0.251200757 | 0.0826 | -3.042 | 0.00235 | 0.06898 |
| ENSG00000182175.13 | 1583.109304 | 0.292891771 | 0.0967 | 3.03043 | 0.002442 | 0.06898 |
| ENSG00000183779.6 | 638.4409784 | 0.2820598 | 0.0929 | 3.03557 | 0.002401 | 0.06898 |
| ENSG00000157890.17 | 310.1371444 | 0.342577484 | 0.1132 | 3.0263 | 0.002476 | 0.06956 |
| ENSG00000206149.10 | 1535.084353 | 0.335945667 | 0.1112 | 3.02063 | 0.002523 | 0.0705 |
| ENSG00000107738.19 | 624.7326888 | 0.501107028 | 0.1662 | 3.01461 | 0.002573 | 0.07128 |
| ENSG00000134317.17 | 183.8521123 | -0.270434641 | 0.0897 | -3.014 | 0.002578 | 0.07128 |
| ENSG00000051620.10 | 264.860968 | 0.326862184 | 0.109 | 2.99805 | 0.002717 | 0.07359 |
| ENSG00000072952.18 | 1472.400334 | 0.630527525 | 0.211 | 2.98821 | 0.002806 | 0.07359 |
| ENSG00000076067.11 | 502.5091858 | 0.328906841 | 0.1101 | 2.98623 | 0.002824 | 0.07359 |
| ENSG00000088280.18 | 483.7216946 | 0.33549393 | 0.1125 | 2.98099 | 0.002873 | 0.07359 |
| ENSG00000101439.8 | 15415.99929 | 0.460757793 | 0.1549 | 2.97436 | 0.002936 | 0.07359 |
| ENSG00000109971.13 | 20086.37083 | -0.30951823 | 0.104 | -2.9763 | 0.002918 | 0.07359 |
| ENSG00000111907.20 | 874.6681372 | 0.445863501 | 0.149 | 2.99297 | 0.002763 | 0.07359 |
| ENSG00000112146.16 | 2155.542578 | -0.287495438 | 0.0967 | -2.9717 | 0.002962 | 0.07359 |
| ENSG00000112149.9 | 610.206279 | -0.315206339 | 0.1061 | -2.9707 | 0.002971 | 0.07359 |
| ENSG00000114805.16 | 902.1127984 | -0.385446804 | 0.1294 | -2.9785 | 0.002897 | 0.07359 |
| ENSG00000115594.11 | 196.4845161 | 0.714234124 | 0.2386 | 2.9934 | 0.002759 | 0.07359 |
| ENSG00000134343.12 | 1540.157367 | -0.306543156 | 0.1031 | -2.9737 | 0.002942 | 0.07359 |
| ENSG00000148925.10 | 1787.031245 | -0.25974832 | 0.0869 | -2.9877 | 0.00281 | 0.07359 |
| ENSG00000157927.16 | 174.8877553 | 0.262658226 | 0.0882 | 2.97767 | 0.002904 | 0.07359 |
| ENSG00000164089.8 | 2146.461798 | 0.693084588 | 0.2312 | 2.99799 | 0.002718 | 0.07359 |
| ENSG00000164600.6 | 1228.604796 | -0.366711505 | 0.1232 | -2.9772 | 0.002909 | 0.07359 |
| ENSG00000168497.4 | 365.6482181 | 0.611709704 | 0.2048 | 2.98744 | 0.002813 | 0.07359 |
| ENSG00000172638.12 | 474.8193945 | 0.332621739 | 0.1112 | 2.99081 | 0.002782 | 0.07359 |
| ENSG00000177189.12 | 1461.659653 | -0.18528858 | 0.0623 | -2.9729 | 0.00295 | 0.07359 |
| ENSG00000184985.16 | 852.1381292 | 0.290228247 | 0.0975 | 2.97694 | 0.002911 | 0.07359 |
| ENSG00000197872.11 | 5169.314158 | -0.280481293 | 0.0935 | -2.9987 | 0.002712 | 0.07359 |
| ENSG00000250366.2 | 934.1576287 | -0.341229908 | 0.1141 | -2.9896 | 0.002793 | 0.07359 |
| ENSG00000104356.10 | 140.0864253 | -0.221781545 | 0.0749 | -2.9628 | 0.003048 | 0.07459 |
| ENSG00000153208.16 | 478.3280191 | 0.49864434 | 0.1683 | 2.96293 | 0.003047 | 0.07459 |
| ENSG00000154556.17 | 3791.333614 | -0.199877152 | 0.0675 | -2.9612 | 0.003065 | 0.07459 |
| ENSG00000166925.8 | 2514.292348 | 0.396997356 | 0.1341 | 2.96077 | 0.003069 | 0.07459 |
| ENSG00000169738.7 | 776.7509568 | 0.306974848 | 0.1038 | 2.95832 | 0.003093 | 0.07484 |
| ENSG00000196218.12 | 1227.713726 | 0.248679018 | 0.0842 | 2.95362 | 0.003141 | 0.07529 |
| ENSG00000204186.7 | 1675.38811 | -0.214222646 | 0.0725 | -2.9545 | 0.003132 | 0.07529 |
| ENSG00000077942.18 | 518.4552193 | 0.36250142 | 0.1228 | 2.95077 | 0.00317 | 0.0753 |
| ENSG00000169676.5 | 188.5763176 | -0.42853305 | 0.1452 | -2.9511 | 0.003166 | 0.0753 |
| ENSG00000135063.17 | 306.7529546 | 0.70304373 | 0.2389 | 2.94296 | 0.003251 | 0.07687 |
| ENSG00000154134.14 | 384.9520282 | 0.289379652 | 0.0984 | 2.94025 | 0.003279 | 0.0772 |
| ENSG00000121310.16 | 819.0154594 | 0.293141747 | 0.0997 | 2.93886 | 0.003294 | 0.0772 |
| ENSG00000115641.18 | 476.3020352 | -0.288483738 | 0.0984 | -2.9323 | 0.003365 | 0.07851 |
| ENSG00000147872.9 | 161.8748221 | 0.44030825 | 0.1503 | 2.93034 | 0.003386 | 0.07864 |
| ENSG00000075391.16 | 2241.684017 | -0.199136413 | 0.0681 | -2.9229 | 0.003468 | 0.07984 |
| ENSG00000104325.6 | 570.8537227 | 0.263473617 | 0.0902 | 2.92152 | 0.003483 | 0.07984 |
| ENSG00000128602.9 | 164.8396393 | 0.49761521 | 0.1702 | 2.92336 | 0.003463 | 0.07984 |
| ENSG00000007168.12 | 11642.24485 | -0.272993385 | 0.0949 | -2.8771 | 0.004013 | 0.08005 |
| ENSG00000016391.10 | 565.9001823 | 0.433928809 | 0.1509 | 2.87496 | 0.004041 | 0.08005 |
| ENSG00000088888.17 | 1843.979386 | 0.214011929 | 0.0738 | 2.89881 | 0.003746 | 0.08005 |
| ENSG00000100243.20 | 2554.74795 | 0.243395856 | 0.0836 | 2.91108 | 0.003602 | 0.08005 |
| ENSG00000102760.12 | 663.1660043 | 0.398816303 | 0.1386 | 2.87841 | 0.003997 | 0.08005 |
| ENSG00000105355.8 | 230.6019854 | 0.471228661 | 0.1631 | 2.88985 | 0.003854 | 0.08005 |
| ENSG00000107902.13 | 1112.412696 | 0.257328811 | 0.0886 | 2.90512 | 0.003671 | 0.08005 |
| ENSG00000112972.14 | 4521.763611 | -0.299278246 | 0.1039 | -2.8808 | 0.003967 | 0.08005 |
| ENSG00000116016.13 | 3222.829377 | 0.340523567 | 0.1179 | 2.88824 | 0.003874 | 0.08005 |
| ENSG00000126603.8 | 259.921486 | 0.300831685 | 0.1033 | 2.91183 | 0.003593 | 0.08005 |
| ENSG00000128203.6 | 775.4944467 | -0.284521903 | 0.0982 | -2.8971 | 0.003766 | 0.08005 |
| ENSG00000128311.13 | 172.824971 | 0.41874846 | 0.1443 | 2.90098 | 0.00372 | 0.08005 |
| ENSG00000130304.16 | 1026.621665 | 0.225430286 | 0.0777 | 2.89966 | 0.003736 | 0.08005 |
| ENSG00000134533.6 | 254.1833868 | 0.297806301 | 0.103 | 2.89035 | 0.003848 | 0.08005 |
| ENSG00000137959.15 | 368.2626357 | -0.47782559 | 0.165 | -2.896 | 0.003779 | 0.08005 |
| ENSG00000138018.17 | 2145.445253 | -0.276086113 | 0.0952 | -2.8998 | 0.003734 | 0.08005 |
| ENSG00000144485.10 | 404.1359102 | 0.282220376 | 0.0983 | 2.87217 | 0.004077 | 0.08005 |
| ENSG00000145817.16 | 791.8763876 | -0.244078795 | 0.0843 | -2.8939 | 0.003805 | 0.08005 |
| ENSG00000146648.16 | 656.7642037 | 0.439351772 | 0.1521 | 2.8877 | 0.003881 | 0.08005 |
| ENSG00000149782.11 | 248.1808879 | 0.358480524 | 0.1242 | 2.88543 | 0.003909 | 0.08005 |
| ENSG00000155097.11 | 6263.033015 | -0.256301379 | 0.0887 | -2.8886 | 0.003869 | 0.08005 |
| ENSG00000156076.9 | 1182.923618 | 0.673404473 | 0.2308 | 2.91804 | 0.003522 | 0.08005 |
| ENSG00000156140.9 | 276.9708009 | -0.325786459 | 0.1118 | -2.915 | 0.003557 | 0.08005 |
| ENSG00000157833.12 | 749.3141043 | 0.475988137 | 0.1632 | 2.91643 | 0.003541 | 0.08005 |
| ENSG00000160678.11 | 1430.290838 | 0.455239936 | 0.156 | 2.91769 | 0.003526 | 0.08005 |
| ENSG00000164211.12 | 1044.420009 | -0.297262565 | 0.1027 | -2.8942 | 0.003801 | 0.08005 |
| ENSG00000165813.17 | 911.2351531 | -0.18733774 | 0.065 | -2.8842 | 0.003924 | 0.08005 |
| ENSG00000168710.17 | 6851.098043 | 0.352212984 | 0.1227 | 2.87115 | 0.00409 | 0.08005 |
| ENSG00000170214.3 | 548.1964334 | -0.322114647 | 0.1106 | -2.9117 | 0.003595 | 0.08005 |
| ENSG00000173597.8 | 191.0643906 | 0.379549245 | 0.1321 | 2.87257 | 0.004071 | 0.08005 |
| ENSG00000176697.18 | 534.8480073 | -0.35156929 | 0.1219 | -2.8833 | 0.003935 | 0.08005 |
| ENSG00000183579.15 | 696.5368873 | 0.462918255 | 0.1609 | 2.87653 | 0.004021 | 0.08005 |
| ENSG00000184845.3 | 459.4959179 | -0.335529107 | 0.1168 | -2.8716 | 0.004084 | 0.08005 |
| ENSG00000184925.11 | 126.7995421 | 0.404523934 | 0.1392 | 2.90636 | 0.003657 | 0.08005 |
| ENSG00000198417.6 | 268.3169056 | 0.511631786 | 0.1776 | 2.88133 | 0.00396 | 0.08005 |
| ENSG00000204136.10 | 228.7744671 | 0.313087879 | 0.1085 | 2.88656 | 0.003895 | 0.08005 |
| ENSG00000213853.9 | 512.0022274 | 0.335015262 | 0.1153 | 2.90646 | 0.003655 | 0.08005 |
| ENSG00000214353.7 | 122.9753443 | 0.464817663 | 0.1609 | 2.88936 | 0.00386 | 0.08005 |
| ENSG00000266964.5 | 1000.586071 | 0.404638982 | 0.1405 | 2.87965 | 0.003981 | 0.08005 |
| ENSG00000163659.12 | 470.2137556 | 0.552809289 | 0.1927 | 2.86941 | 0.004112 | 0.08019 |
| ENSG00000139044.10 | 384.8481593 | -0.270260179 | 0.0943 | -2.8662 | 0.004155 | 0.08042 |
| ENSG00000163138.18 | 394.8204951 | -0.154023291 | 0.0537 | -2.8668 | 0.004147 | 0.08042 |
| ENSG00000204580.12 | 2341.688718 | 0.363263151 | 0.127 | 2.86052 | 0.004229 | 0.08156 |
| ENSG00000018625.14 | 16613.41505 | 0.497613998 | 0.1742 | 2.85685 | 0.004279 | 0.08191 |
| ENSG00000125148.6 | 1840.241547 | 0.754375362 | 0.264 | 2.857 | 0.004277 | 0.08191 |
| ENSG00000070961.15 | 11528.80746 | -0.29679056 | 0.104 | -2.8545 | 0.00431 | 0.08207 |
| ENSG00000161509.13 | 1025.03731 | 0.571724694 | 0.2003 | 2.85391 | 0.004318 | 0.08207 |
| ENSG00000132470.13 | 543.1865289 | 0.674492153 | 0.2366 | 2.85032 | 0.004367 | 0.0823 |
| ENSG00000145428.14 | 608.6145029 | -0.233567713 | 0.082 | -2.8495 | 0.004378 | 0.0823 |
| ENSG00000182054.9 | 2630.449212 | 0.231032705 | 0.081 | 2.85095 | 0.004359 | 0.0823 |
| ENSG00000198721.12 | 508.1021196 | 0.23887378 | 0.0839 | 2.84841 | 0.004394 | 0.0823 |
| ENSG00000104870.12 | 385.3654324 | 0.344810436 | 0.1212 | 2.84527 | 0.004437 | 0.08245 |
| ENSG00000132825.6 | 178.7886282 | 0.317986697 | 0.1117 | 2.84565 | 0.004432 | 0.08245 |
| ENSG00000160284.14 | 343.756055 | 0.425942472 | 0.1497 | 2.84442 | 0.004449 | 0.08245 |
| ENSG00000170091.10 | 10472.82852 | -0.255295982 | 0.0898 | -2.8429 | 0.004471 | 0.08255 |
| ENSG00000196305.17 | 2484.525669 | -0.238581338 | 0.084 | -2.8418 | 0.004486 | 0.08255 |
| ENSG00000080824.18 | 35389.8096 | -0.229621288 | 0.0809 | -2.8379 | 0.004541 | 0.08265 |
| ENSG00000101871.14 | 240.1075518 | 0.349326019 | 0.1233 | 2.83331 | 0.004607 | 0.08265 |
| ENSG00000107317.11 | 10474.67469 | 0.373363914 | 0.1318 | 2.83274 | 0.004615 | 0.08265 |
| ENSG00000139211.6 | 427.8926208 | -0.306598355 | 0.1082 | -2.8343 | 0.004592 | 0.08265 |
| ENSG00000147416.10 | 10692.88082 | -0.265878308 | 0.0937 | -2.8373 | 0.00455 | 0.08265 |
| ENSG00000148175.12 | 2173.348321 | 0.450269244 | 0.1588 | 2.83492 | 0.004584 | 0.08265 |
| ENSG00000170370.11 | 672.5168641 | 0.482869247 | 0.1705 | 2.83271 | 0.004616 | 0.08265 |
| ENSG00000180530.10 | 1074.299164 | -0.240909959 | 0.0851 | -2.8322 | 0.004623 | 0.08265 |
| ENSG00000198732.10 | 714.9656215 | 0.369211471 | 0.1304 | 2.83144 | 0.004634 | 0.08265 |
| ENSG00000076248.10 | 449.1387736 | 0.255643784 | 0.0904 | 2.8292 | 0.004666 | 0.08267 |
| ENSG00000161714.11 | 1289.922243 | 0.271573316 | 0.096 | 2.82944 | 0.004663 | 0.08267 |
| ENSG00000067182.7 | 360.2707029 | 0.474003356 | 0.1685 | 2.81235 | 0.004918 | 0.08356 |
| ENSG00000103591.12 | 1129.202327 | -0.217785234 | 0.0773 | -2.8178 | 0.004835 | 0.08356 |
| ENSG00000107249.21 | 229.4086268 | 0.425040925 | 0.1506 | 2.82174 | 0.004776 | 0.08356 |
| ENSG00000110799.13 | 1371.475488 | 0.448617738 | 0.1593 | 2.81594 | 0.004863 | 0.08356 |
| ENSG00000154856.12 | 912.4126843 | 0.404316134 | 0.1438 | 2.8119 | 0.004925 | 0.08356 |
| ENSG00000164904.16 | 2011.47367 | 0.3327244 | 0.1181 | 2.8177 | 0.004837 | 0.08356 |
| ENSG00000168385.17 | 5111.762616 | 0.256147147 | 0.0908 | 2.8225 | 0.004765 | 0.08356 |
| ENSG00000173611.17 | 2916.439848 | -0.249859521 | 0.0887 | -2.817 | 0.004848 | 0.08356 |
| ENSG00000179520.10 | 127.1057205 | -0.634980899 | 0.2258 | -2.8125 | 0.004915 | 0.08356 |
| ENSG00000248874.5 | 133.1598579 | 0.27334104 | 0.0971 | 2.81435 | 0.004888 | 0.08356 |
| ENSG00000257151.1 | 6714.777149 | -0.235084873 | 0.0833 | -2.8218 | 0.004776 | 0.08356 |
| ENSG00000271533.1 | 428.9588366 | -0.299399578 | 0.1063 | -2.8162 | 0.00486 | 0.08356 |
| ENSG00000273259.2 | 225.0005272 | 2.02873173 | 0.721 | 2.81359 | 0.004899 | 0.08356 |
| ENSG00000109107.13 | 11398.95909 | 0.296261563 | 0.1055 | 2.80756 | 0.004992 | 0.08415 |
| ENSG00000165424.6 | 2240.659601 | 0.356179803 | 0.1268 | 2.80825 | 0.004981 | 0.08415 |
| ENSG00000177917.10 | 163.3555815 | 0.252485304 | 0.09 | 2.80588 | 0.005018 | 0.08432 |
| ENSG00000254685.6 | 190.7509631 | -0.214548126 | 0.0765 | -2.8045 | 0.00504 | 0.08441 |
| ENSG00000100095.18 | 5913.885229 | -0.233435405 | 0.0833 | -2.8022 | 0.005076 | 0.08448 |
| ENSG00000144619.14 | 1110.794057 | -0.249784187 | 0.0891 | -2.8029 | 0.005064 | 0.08448 |
| ENSG00000118946.11 | 3038.96428 | -0.266192641 | 0.0952 | -2.7963 | 0.00517 | 0.08492 |
| ENSG00000145536.15 | 127.883223 | -0.291571053 | 0.1042 | -2.799 | 0.005126 | 0.08492 |
| ENSG00000179774.8 | 140.0403799 | -0.313302802 | 0.1121 | -2.7944 | 0.0052 | 0.08492 |
| ENSG00000187091.13 | 297.4094244 | 0.445055167 | 0.1592 | 2.79611 | 0.005172 | 0.08492 |
| ENSG00000196353.11 | 1780.906643 | -0.317015154 | 0.1134 | -2.7952 | 0.005188 | 0.08492 |
| ENSG00000198301.11 | 942.7054727 | -0.247732074 | 0.0886 | -2.7946 | 0.005197 | 0.08492 |
| ENSG00000086544.2 | 280.3822862 | 0.383521453 | 0.1375 | 2.78991 | 0.005272 | 0.08546 |
| ENSG00000089472.16 | 457.8082042 | 0.505607875 | 0.1813 | 2.78932 | 0.005282 | 0.08546 |
| ENSG00000130158.13 | 342.2132089 | 0.322564753 | 0.1156 | 2.78976 | 0.005275 | 0.08546 |
| ENSG00000148672.8 | 8733.236213 | 0.333747672 | 0.1197 | 2.78815 | 0.005301 | 0.0855 |
| ENSG00000000003.14 | 272.9838746 | 0.225892943 | 0.0811 | 2.78667 | 0.005325 | 0.08563 |
| ENSG00000265972.5 | 1308.493071 | 0.506853949 | 0.182 | 2.78535 | 0.005347 | 0.08572 |
| ENSG00000122863.5 | 327.5611876 | 0.601845869 | 0.2165 | 2.77971 | 0.005441 | 0.08695 |
| ENSG00000198938.2 | 379375.5292 | 0.366746337 | 0.132 | 2.7775 | 0.005478 | 0.08728 |
| ENSG00000087266.15 | 646.6670266 | 0.383020675 | 0.138 | 2.77506 | 0.005519 | 0.08767 |
| ENSG00000144791.9 | 312.2169508 | 0.202816534 | 0.0732 | 2.76996 | 0.005606 | 0.08826 |
| ENSG00000146674.14 | 540.9833178 | 0.450635422 | 0.1626 | 2.7709 | 0.00559 | 0.08826 |
| ENSG00000167543.15 | 213.7188088 | 0.261622001 | 0.0945 | 2.7699 | 0.005607 | 0.08826 |
| ENSG00000141639.11 | 2083.837805 | 0.256636563 | 0.0928 | 2.765 | 0.005692 | 0.0892 |
| ENSG00000169439.11 | 936.9060493 | 0.458760135 | 0.1659 | 2.76451 | 0.005701 | 0.0892 |
| ENSG00000065559.14 | 3985.891609 | -0.260947335 | 0.0946 | -2.7593 | 0.005793 | 0.08957 |
| ENSG00000110218.8 | 415.774422 | -0.274580424 | 0.0997 | -2.7536 | 0.005895 | 0.08957 |
| ENSG00000128309.16 | 283.6321715 | 0.250563692 | 0.0909 | 2.75712 | 0.005831 | 0.08957 |
| ENSG00000132872.11 | 5481.111737 | -0.365958721 | 0.1327 | -2.7581 | 0.005814 | 0.08957 |
| ENSG00000135821.17 | 32357.9099 | 0.522897878 | 0.1896 | 2.75756 | 0.005823 | 0.08957 |
| ENSG00000137285.9 | 5150.337321 | 0.499590094 | 0.1811 | 2.75926 | 0.005793 | 0.08957 |
| ENSG00000144057.15 | 945.1980634 | -0.228118602 | 0.0828 | -2.754 | 0.005888 | 0.08957 |
| ENSG00000165795.22 | 17198.77313 | 0.386729215 | 0.1404 | 2.7535 | 0.005896 | 0.08957 |
| ENSG00000171100.14 | 325.1753851 | 0.248308335 | 0.0901 | 2.75565 | 0.005858 | 0.08957 |
| ENSG00000184557.4 | 231.4315575 | 1.255515329 | 0.4554 | 2.75696 | 0.005834 | 0.08957 |
| ENSG00000124145.6 | 857.3049763 | 0.740383441 | 0.2698 | 2.74421 | 0.006066 | 0.09005 |
| ENSG00000169246.16 | 1526.773438 | 0.304585238 | 0.1109 | 2.74748 | 0.006006 | 0.09005 |
| ENSG00000183054.11 | 2985.283085 | -0.29130751 | 0.1061 | -2.7462 | 0.006029 | 0.09005 |
| ENSG00000186998.15 | 647.0837226 | 0.285000188 | 0.1037 | 2.74702 | 0.006014 | 0.09005 |
| ENSG00000196843.15 | 323.0079413 | 0.464084488 | 0.1691 | 2.74443 | 0.006062 | 0.09005 |
| ENSG00000204054.13 | 603.4918518 | 0.223932522 | 0.0815 | 2.74661 | 0.006021 | 0.09005 |
| ENSG00000204128.5 | 1603.901446 | 0.178442369 | 0.065 | 2.74647 | 0.006024 | 0.09005 |
| ENSG00000280165.1 | 1074.307497 | -0.277054642 | 0.1009 | -2.7456 | 0.006039 | 0.09005 |
| ENSG00000114738.10 | 209.5500706 | 0.408353527 | 0.1492 | 2.73652 | 0.006209 | 0.09154 |
| ENSG00000137040.9 | 1218.641106 | -0.21995843 | 0.0804 | -2.7372 | 0.006196 | 0.09154 |
| ENSG00000198300.12 | 10506.90381 | -0.262736343 | 0.096 | -2.736 | 0.006218 | 0.09154 |
| ENSG00000134917.9 | 655.9500979 | -0.254357163 | 0.0932 | -2.7301 | 0.006331 | 0.0919 |
| ENSG00000145808.8 | 137.9670236 | -0.254227433 | 0.0931 | -2.731 | 0.006313 | 0.0919 |
| ENSG00000149489.8 | 210.9022552 | 0.386896942 | 0.1416 | 2.73304 | 0.006275 | 0.0919 |
| ENSG00000156017.12 | 440.4657099 | -0.212197433 | 0.0777 | -2.7324 | 0.006287 | 0.0919 |
| ENSG00000198712.1 | 260136.0079 | 0.361072779 | 0.1322 | 2.73044 | 0.006325 | 0.0919 |
| ENSG00000034239.10 | 176.9190825 | -0.440153747 | 0.1613 | -2.7287 | 0.006358 | 0.09204 |
| ENSG00000135324.5 | 565.5405437 | -0.326995808 | 0.1199 | -2.7276 | 0.00638 | 0.0921 |
| ENSG00000183943.5 | 275.606306 | 0.290366548 | 0.1065 | 2.72577 | 0.006415 | 0.09236 |
| ENSG00000113790.10 | 145.5238976 | 0.238468269 | 0.0876 | 2.72307 | 0.006468 | 0.09255 |
| ENSG00000135622.12 | 849.5563261 | -0.239309066 | 0.0879 | -2.7215 | 0.006499 | 0.09255 |
| ENSG00000143416.20 | 580.1501695 | 0.429159588 | 0.1577 | 2.72157 | 0.006497 | 0.09255 |
| ENSG00000179399.14 | 338.4003353 | 0.459356513 | 0.1688 | 2.72202 | 0.006488 | 0.09255 |
| ENSG00000027001.9 | 160.3703248 | -0.177761002 | 0.0655 | -2.7156 | 0.006616 | 0.0927 |
| ENSG00000076555.15 | 670.3796883 | 0.421647051 | 0.1551 | 2.7185 | 0.006558 | 0.0927 |
| ENSG00000137877.9 | 127.4116522 | -0.222644254 | 0.0819 | -2.7177 | 0.006573 | 0.0927 |
| ENSG00000167614.13 | 5598.152543 | 0.286591041 | 0.1055 | 2.71558 | 0.006616 | 0.0927 |
| ENSG00000170425.3 | 147.2930502 | 0.448598333 | 0.165 | 2.71841 | 0.00656 | 0.0927 |
| ENSG00000270580.5 | 123.2052522 | -0.333879634 | 0.1229 | -2.716 | 0.006608 | 0.0927 |
| ENSG00000144730.16 | 512.3725107 | 0.238970938 | 0.0881 | 2.71376 | 0.006652 | 0.09296 |
| ENSG00000140988.15 | 3283.666653 | 0.244887707 | 0.0903 | 2.712 | 0.006688 | 0.09313 |
| ENSG00000198898.12 | 2806.717231 | -0.200597804 | 0.074 | -2.7114 | 0.0067 | 0.09313 |
| ENSG00000069431.11 | 617.8905408 | 0.300181533 | 0.1108 | 2.71028 | 0.006723 | 0.09319 |
| ENSG00000006756.15 | 279.6406576 | 0.217214733 | 0.0802 | 2.70763 | 0.006776 | 0.09353 |
| ENSG00000144909.7 | 437.6453202 | 0.299521456 | 0.1106 | 2.7073 | 0.006783 | 0.09353 |
| ENSG00000111058.7 | 534.665924 | 0.43109181 | 0.1594 | 2.70384 | 0.006854 | 0.09383 |
| ENSG00000119699.7 | 346.4571297 | 0.366383307 | 0.1355 | 2.70384 | 0.006854 | 0.09383 |
| ENSG00000145623.12 | 233.6367238 | 0.748131153 | 0.2767 | 2.70364 | 0.006859 | 0.09383 |
| ENSG00000029153.14 | 816.3399365 | -0.213387256 | 0.079 | -2.7026 | 0.00688 | 0.09388 |
| ENSG00000168461.12 | 2744.078645 | 0.349881336 | 0.1296 | 2.69894 | 0.006956 | 0.09463 |
| ENSG00000186951.16 | 599.4039075 | 0.341297196 | 0.1265 | 2.69822 | 0.006971 | 0.09463 |
| ENSG00000184307.13 | 1079.102218 | -0.290451138 | 0.1078 | -2.6954 | 0.007031 | 0.09519 |
| ENSG00000168309.16 | 20192.21484 | 0.406533821 | 0.1509 | 2.69408 | 0.007058 | 0.09532 |
| ENSG00000254154.8 | 185.6334427 | -0.294421718 | 0.1094 | -2.6921 | 0.007101 | 0.09565 |
| ENSG00000101938.14 | 1085.894513 | 0.336275533 | 0.125 | 2.68937 | 0.007159 | 0.09617 |
| ENSG00000122756.14 | 987.8383211 | 0.306627008 | 0.1141 | 2.68766 | 0.007195 | 0.09642 |
| ENSG00000185187.12 | 206.6138424 | 0.305809939 | 0.1138 | 2.68661 | 0.007218 | 0.09648 |
| ENSG00000019505.7 | 5833.470928 | -0.268763452 | 0.1002 | -2.6825 | 0.007308 | 0.09656 |
| ENSG00000091157.13 | 3214.253404 | -0.239826919 | 0.0895 | -2.6804 | 0.007353 | 0.09656 |
| ENSG00000130876.11 | 206.1356424 | 0.530085574 | 0.1976 | 2.68318 | 0.007293 | 0.09656 |
| ENSG00000144339.11 | 1284.217714 | -0.274758398 | 0.1024 | -2.683 | 0.007297 | 0.09656 |
| ENSG00000198108.3 | 239.3702428 | -0.259291445 | 0.0967 | -2.6808 | 0.007345 | 0.09656 |
| ENSG00000278727.1 | 158.4589981 | -0.249742297 | 0.0931 | -2.6826 | 0.007305 | 0.09656 |
| ENSG00000281183.1 | 259.3414572 | -0.309123249 | 0.1153 | -2.6804 | 0.007353 | 0.09656 |
| ENSG00000111262.4 | 1214.55067 | -0.409662236 | 0.1531 | -2.676 | 0.00745 | 0.09709 |
| ENSG00000152503.9 | 1130.129317 | -0.245072647 | 0.0916 | -2.6768 | 0.007432 | 0.09709 |
| ENSG00000197147.12 | 3535.064753 | -0.245407193 | 0.0917 | -2.6768 | 0.007433 | 0.09709 |
| ENSG00000232164.1 | 125.9709992 | 0.336336208 | 0.1258 | 2.67317 | 0.007514 | 0.09768 |
| ENSG00000092621.11 | 1315.234003 | 0.39286478 | 0.1471 | 2.67016 | 0.007582 | 0.09805 |
| ENSG00000133019.11 | 2241.462133 | -0.27913124 | 0.1046 | -2.6698 | 0.00759 | 0.09805 |
| ENSG00000151322.18 | 1311.153771 | 0.32074738 | 0.1202 | 2.66857 | 0.007618 | 0.09805 |
| ENSG00000272933.1 | 242.8904415 | 0.264977135 | 0.0993 | 2.66908 | 0.007606 | 0.09805 |
| ENSG00000006715.15 | 3334.824128 | -0.201444547 | 0.0758 | -2.656 | 0.007907 | 0.09838 |
| ENSG00000081853.14 | 141.0320995 | 0.42145683 | 0.1588 | 2.65381 | 0.007959 | 0.09838 |
| ENSG00000100504.16 | 205.2718449 | 0.39564586 | 0.1491 | 2.65365 | 0.007963 | 0.09838 |
| ENSG00000105227.14 | 157.752722 | 0.390526994 | 0.1468 | 2.65967 | 0.007822 | 0.09838 |
| ENSG00000115419.12 | 9608.610268 | -0.231662763 | 0.0871 | -2.6599 | 0.007816 | 0.09838 |
| ENSG00000116017.10 | 170.4783262 | 0.17106977 | 0.0643 | 2.66007 | 0.007812 | 0.09838 |
| ENSG00000134121.9 | 9798.902337 | -0.285183121 | 0.1072 | -2.6613 | 0.007783 | 0.09838 |
| ENSG00000134569.9 | 2024.99386 | 0.468570625 | 0.1764 | 2.65573 | 0.007914 | 0.09838 |
| ENSG00000143842.14 | 383.4877943 | 0.263713916 | 0.0993 | 2.65609 | 0.007905 | 0.09838 |
| ENSG00000158987.20 | 900.1415563 | -0.164414788 | 0.0617 | -2.6658 | 0.007681 | 0.09838 |
| ENSG00000159082.17 | 5691.845409 | -0.281656238 | 0.106 | -2.6573 | 0.007877 | 0.09838 |
| ENSG00000163820.14 | 424.7679963 | 0.319028772 | 0.1197 | 2.66588 | 0.007679 | 0.09838 |
| ENSG00000182022.17 | 1229.927385 | -0.188840976 | 0.0711 | -2.6572 | 0.00788 | 0.09838 |
| ENSG00000182118.6 | 149.1604739 | 0.370862756 | 0.1397 | 2.65397 | 0.007955 | 0.09838 |
| ENSG00000198478.7 | 2127.750591 | -0.207167828 | 0.078 | -2.6558 | 0.007912 | 0.09838 |
| ENSG00000227827.3 | 237.5437372 | 1.233968209 | 0.4644 | 2.65694 | 0.007885 | 0.09838 |
| ENSG00000234899.9 | 124.5553491 | 0.259598743 | 0.0975 | 2.66145 | 0.00778 | 0.09838 |
| ENSG00000107105.14 | 2155.202955 | -0.240371221 | 0.0907 | -2.6495 | 0.008061 | 0.09857 |
| ENSG00000114573.9 | 9114.211783 | -0.278556287 | 0.1051 | -2.6509 | 0.008029 | 0.09857 |
| ENSG00000116514.16 | 632.7910882 | -0.242532833 | 0.0915 | -2.6513 | 0.008018 | 0.09857 |
| ENSG00000169006.6 | 511.8877771 | 0.556060234 | 0.21 | 2.64765 | 0.008105 | 0.09857 |
| ENSG00000180667.10 | 490.9186387 | -0.218100786 | 0.0824 | -2.6475 | 0.00811 | 0.09857 |
| ENSG00000198840.2 | 65313.42286 | 0.390169892 | 0.1472 | 2.64999 | 0.008049 | 0.09857 |
| ENSG00000245573.7 | 142.5051524 | 0.199994323 | 0.0755 | 2.64898 | 0.008073 | 0.09857 |
| ENSG00000100033.16 | 1479.904266 | 0.474477323 | 0.1803 | 2.63155 | 0.0085 | 0.09878 |
| ENSG00000100427.15 | 4127.417475 | 0.496973306 | 0.1889 | 2.63022 | 0.008533 | 0.09878 |
| ENSG00000102349.16 | 366.2941783 | -0.20299759 | 0.0771 | -2.6322 | 0.008484 | 0.09878 |
| ENSG00000103811.15 | 624.927613 | 0.382827999 | 0.1452 | 2.6364 | 0.008379 | 0.09878 |
| ENSG00000110492.15 | 157.7224995 | 0.293785061 | 0.1112 | 2.64245 | 0.008231 | 0.09878 |
| ENSG00000125851.9 | 2895.066206 | -0.272584173 | 0.1036 | -2.6314 | 0.008502 | 0.09878 |
| ENSG00000134321.11 | 173.4904795 | -0.275075635 | 0.1045 | -2.6316 | 0.008498 | 0.09878 |
| ENSG00000135094.10 | 132.3058703 | 0.547109162 | 0.2077 | 2.63372 | 0.008446 | 0.09878 |
| ENSG00000142875.19 | 12373.085 | -0.229418634 | 0.0868 | -2.6428 | 0.008222 | 0.09878 |
| ENSG00000144645.13 | 457.9152452 | -0.232543236 | 0.0883 | -2.6344 | 0.008429 | 0.09878 |
| ENSG00000148158.16 | 1742.128061 | -0.20199492 | 0.0767 | -2.6347 | 0.008422 | 0.09878 |
| ENSG00000152413.14 | 2875.851175 | -0.258366383 | 0.0978 | -2.6413 | 0.008258 | 0.09878 |
| ENSG00000154319.14 | 709.4091477 | 0.427838869 | 0.1626 | 2.63118 | 0.008509 | 0.09878 |
| ENSG00000154822.16 | 817.5715817 | -0.318738591 | 0.1211 | -2.6321 | 0.008486 | 0.09878 |
| ENSG00000163393.12 | 517.1145798 | -0.20165222 | 0.0763 | -2.6426 | 0.008228 | 0.09878 |
| ENSG00000163539.16 | 8839.681328 | -0.213333475 | 0.0811 | -2.6298 | 0.008543 | 0.09878 |
| ENSG00000166257.8 | 12722.07474 | -0.266440505 | 0.1013 | -2.6307 | 0.00852 | 0.09878 |
| ENSG00000173272.14 | 955.5773635 | 0.247469224 | 0.094 | 2.63156 | 0.008499 | 0.09878 |
| ENSG00000181444.12 | 204.2910894 | 0.264002384 | 0.0999 | 2.642 | 0.008242 | 0.09878 |
| ENSG00000210082.2 | 1273536.754 | 0.355386499 | 0.1348 | 2.63711 | 0.008361 | 0.09878 |
| ENSG00000237870.6 | 137.3220258 | 0.221585757 | 0.0839 | 2.64153 | 0.008253 | 0.09878 |
| ENSG00000253982.1 | 253.0773459 | 0.237816999 | 0.0903 | 2.63223 | 0.008483 | 0.09878 |
| ENSG00000128849.10 | 862.0865723 | 0.292541234 | 0.1113 | 2.62848 | 0.008577 | 0.09895 |
| ENSG00000109113.18 | 291.6791859 | 0.447929378 | 0.1705 | 2.62769 | 0.008597 | 0.09896 |
| ENSG00000153446.15 | 202.9942341 | 0.441551125 | 0.1681 | 2.6264 | 0.008629 | 0.09911 |
| ENSG00000074590.13 | 4316.722815 | -0.328966875 | 0.1254 | -2.6229 | 0.008719 | 0.09993 |

**Supplementary Table 3 RNA-seq in MDD vs Controls**

| Gene Ensemble ID | baseMean | log2FoldChange | lfcSE | stat | pvalue | padj |
| --- | --- | --- | --- | --- | --- | --- |
| ENSG00000121207.11 | 197.2274 | 0.912906057 | 0.1603 | 5.6943 | 1.24E-08 | 0.0001 |
| ENSG00000160606.10 | 69.5778 | 1.035594387 | 0.1983 | 5.2225 | 1.76E-07 | 0.0005 |
| ENSG00000175287.18 | 256.2094 | 0.864415678 | 0.1642 | 5.263 | 1.42E-07 | 0.0005 |
| ENSG00000185220.11 | 376.162 | 0.480757873 | 0.0912 | 5.2734 | 1.34E-07 | 0.0005 |
| ENSG00000176244.6 | 779.0974 | 0.812075787 | 0.1589 | 5.1113 | 3.20E-07 | 0.0005 |
| ENSG00000214456.8 | 187.0731 | 0.794324426 | 0.1552 | 5.1168 | 3.11E-07 | 0.0005 |
| ENSG00000124440.15 | 606.7692 | 0.941920116 | 0.1905 | 4.9442 | 7.65E-07 | 0.001 |
| ENSG00000132541.10 | 712.3851 | 0.481864623 | 0.0989 | 4.8743 | 1.09E-06 | 0.001 |
| ENSG00000147119.3 | 251.7566 | 0.931770988 | 0.191 | 4.8786 | 1.07E-06 | 0.001 |
| ENSG00000166292.11 | 99.84989 | 0.889665426 | 0.1808 | 4.9207 | 8.62E-07 | 0.001 |
| ENSG00000279103.1 | 99.94133 | 0.912942009 | 0.1865 | 4.8943 | 9.87E-07 | 0.001 |
| ENSG00000056487.15 | 125.6708 | 0.757104683 | 0.1607 | 4.71 | 2.48E-06 | 0.0011 |
| ENSG00000111087.9 | 36.37791 | 1.082827773 | 0.2278 | 4.7529 | 2.00E-06 | 0.0011 |
| ENSG00000111275.12 | 5859.444 | 0.502586334 | 0.1066 | 4.7166 | 2.40E-06 | 0.0011 |
| ENSG00000132164.9 | 965.8192 | 0.639572988 | 0.1342 | 4.7667 | 1.87E-06 | 0.0011 |
| ENSG00000138759.17 | 756.8359 | -0.621747945 | 0.1287 | -4.83 | 1.37E-06 | 0.0011 |
| ENSG00000139352.3 | 242.6982 | 0.476113497 | 0.1006 | 4.7315 | 2.23E-06 | 0.0011 |
| ENSG00000163884.3 | 531.648 | 0.573389399 | 0.1215 | 4.7176 | 2.39E-06 | 0.0011 |
| ENSG00000165478.6 | 3937.246 | 0.99043693 | 0.2044 | 4.8463 | 1.26E-06 | 0.0011 |
| ENSG00000168209.4 | 815.1293 | 0.992762628 | 0.2098 | 4.7331 | 2.21E-06 | 0.0011 |
| ENSG00000177133.10 | 738.3511 | 0.907612217 | 0.191 | 4.752 | 2.01E-06 | 0.0011 |
| ENSG00000246022.2 | 61.45793 | 0.761495297 | 0.1592 | 4.7835 | 1.72E-06 | 0.0011 |
| ENSG00000255690.2 | 1413.821 | 0.852295666 | 0.1799 | 4.7369 | 2.17E-06 | 0.0011 |
| ENSG00000280087.1 | 181.6095 | 0.784637368 | 0.1651 | 4.7539 | 2.00E-06 | 0.0011 |
| ENSG00000099840.13 | 198.6617 | 0.516895386 | 0.1109 | 4.6613 | 3.14E-06 | 0.0011 |
| ENSG00000105088.8 | 3423.557 | 0.565163702 | 0.1212 | 4.6648 | 3.09E-06 | 0.0011 |
| ENSG00000105552.14 | 322.1754 | 0.368299776 | 0.0791 | 4.6566 | 3.22E-06 | 0.0011 |
| ENSG00000107551.20 | 1334.856 | 0.35551548 | 0.0766 | 4.6439 | 3.42E-06 | 0.0011 |
| ENSG00000111405.8 | 50.70506 | 0.770791032 | 0.1672 | 4.61 | 4.03E-06 | 0.0011 |
| ENSG00000134508.12 | 1958.795 | 0.692949658 | 0.1503 | 4.6113 | 4.00E-06 | 0.0011 |
| ENSG00000137434.11 | 13.49299 | 0.734441682 | 0.1593 | 4.6115 | 4.00E-06 | 0.0011 |
| ENSG00000138670.17 | 399.6312 | -0.469341671 | 0.1012 | -4.638 | 3.52E-06 | 0.0011 |
| ENSG00000145020.15 | 239.1144 | 0.371286334 | 0.0804 | 4.6177 | 3.88E-06 | 0.0011 |
| ENSG00000174498.13 | 156.2136 | 0.389887357 | 0.0832 | 4.6839 | 2.81E-06 | 0.0011 |
| ENSG00000211592.8 | 16.70326 | -2.66188589 | 0.5701 | -4.669 | 3.02E-06 | 0.0011 |
| ENSG00000239282.7 | 138.4068 | 0.740203148 | 0.1604 | 4.6157 | 3.92E-06 | 0.0011 |
| ENSG00000259823.5 | 182.3106 | -0.669209741 | 0.1451 | -4.612 | 3.98E-06 | 0.0011 |
| ENSG00000278709.1 | 21.84242 | -0.666984434 | 0.144 | -4.632 | 3.62E-06 | 0.0011 |
| ENSG00000159423.16 | 1426.851 | 0.82323545 | 0.1793 | 4.5921 | 4.39E-06 | 0.0011 |
| ENSG00000213563.6 | 395.7833 | 0.387841808 | 0.0845 | 4.5892 | 4.45E-06 | 0.0011 |
| ENSG00000130304.16 | 1117.446 | 0.411477142 | 0.0898 | 4.5804 | 4.64E-06 | 0.0012 |
| ENSG00000129007.14 | 36.50058 | 1.018111508 | 0.2247 | 4.5303 | 5.89E-06 | 0.0013 |
| ENSG00000154930.14 | 1415.939 | 0.811596969 | 0.179 | 4.5333 | 5.81E-06 | 0.0013 |
| ENSG00000165795.22 | 19499.2 | 0.679472214 | 0.1501 | 4.5283 | 5.95E-06 | 0.0013 |
| ENSG00000167676.4 | 166.024 | 1.114639029 | 0.2459 | 4.5335 | 5.80E-06 | 0.0013 |
| ENSG00000183773.15 | 1468.881 | 0.607680351 | 0.134 | 4.5334 | 5.80E-06 | 0.0013 |
| ENSG00000271447.5 | 315.541 | 0.80838449 | 0.1783 | 4.5331 | 5.81E-06 | 0.0013 |
| ENSG00000121570.12 | 236.5686 | 1.378421234 | 0.3048 | 4.5222 | 6.12E-06 | 0.0013 |
| ENSG00000141401.11 | 95.37342 | 0.566464439 | 0.1255 | 4.513 | 6.39E-06 | 0.0013 |
| ENSG00000166033.11 | 4464.653 | 0.621604376 | 0.1378 | 4.5109 | 6.45E-06 | 0.0013 |
| ENSG00000165424.6 | 2425.954 | 0.533459548 | 0.1186 | 4.4963 | 6.92E-06 | 0.0014 |
| ENSG00000088826.17 | 724.598 | 0.659384469 | 0.1474 | 4.4732 | 7.71E-06 | 0.0015 |
| ENSG00000154856.12 | 1024.659 | 0.673911661 | 0.1507 | 4.4714 | 7.77E-06 | 0.0015 |
| ENSG00000167315.17 | 669.8439 | 0.62334091 | 0.1393 | 4.4759 | 7.61E-06 | 0.0015 |
| ENSG00000139800.8 | 131.7381 | 0.628041896 | 0.1406 | 4.4656 | 7.99E-06 | 0.0015 |
| ENSG00000181444.12 | 221.0688 | 0.433824157 | 0.0973 | 4.4603 | 8.19E-06 | 0.0015 |
| ENSG00000272189.1 | 176.4028 | 0.91382829 | 0.205 | 4.4584 | 8.26E-06 | 0.0015 |
| ENSG00000073756.11 | 512.9418 | -1.198423659 | 0.2692 | -4.452 | 8.50E-06 | 0.0015 |
| ENSG00000095917.13 | 12.24725 | 1.615858283 | 0.3636 | 4.4445 | 8.81E-06 | 0.0015 |
| ENSG00000122756.14 | 1110.902 | 0.578707271 | 0.1305 | 4.4361 | 9.16E-06 | 0.0015 |
| ENSG00000132692.18 | 5194.371 | 0.626566604 | 0.1412 | 4.4383 | 9.07E-06 | 0.0015 |
| ENSG00000157833.12 | 852.7857 | 0.778270948 | 0.1753 | 4.4386 | 9.05E-06 | 0.0015 |
| ENSG00000263165.1 | 75.94467 | 0.515228326 | 0.1161 | 4.4392 | 9.03E-06 | 0.0015 |
| ENSG00000187527.10 | 59.55576 | 0.494177245 | 0.1117 | 4.4252 | 9.64E-06 | 0.0015 |
| ENSG00000214353.7 | 138.2394 | 0.737203309 | 0.1666 | 4.4258 | 9.61E-06 | 0.0015 |
| ENSG00000177917.10 | 174.4994 | 0.38999313 | 0.0884 | 4.4123 | 1.02E-05 | 0.0016 |
| ENSG00000124171.8 | 282.9921 | 0.602099597 | 0.1369 | 4.3995 | 1.09E-05 | 0.0016 |
| ENSG00000263146.2 | 320.5808 | 0.593334648 | 0.1348 | 4.4008 | 1.08E-05 | 0.0016 |
| ENSG00000100033.16 | 1691.487 | 0.787924933 | 0.1804 | 4.3676 | 1.26E-05 | 0.0017 |
| ENSG00000103740.9 | 1488.833 | 0.861252384 | 0.1974 | 4.3627 | 1.28E-05 | 0.0017 |
| ENSG00000128311.13 | 189.609 | 0.629569794 | 0.1437 | 4.3824 | 1.17E-05 | 0.0017 |
| ENSG00000167614.13 | 6337.165 | 0.575142566 | 0.1317 | 4.366 | 1.27E-05 | 0.0017 |
| ENSG00000168913.6 | 2690.903 | 0.537939532 | 0.123 | 4.3733 | 1.22E-05 | 0.0017 |
| ENSG00000182054.9 | 2891.398 | 0.443153542 | 0.101 | 4.387 | 1.15E-05 | 0.0017 |
| ENSG00000184985.16 | 917.3183 | 0.449952308 | 0.1027 | 4.3826 | 1.17E-05 | 0.0017 |
| ENSG00000196071.4 | 86.23646 | -0.703781049 | 0.1607 | -4.379 | 1.19E-05 | 0.0017 |
| ENSG00000275294.4 | 11.26307 | 0.767843225 | 0.176 | 4.3628 | 1.28E-05 | 0.0017 |
| ENSG00000279175.1 | 174.6829 | 0.952260472 | 0.2184 | 4.3598 | 1.30E-05 | 0.0017 |
| ENSG00000279474.1 | 28.55098 | 0.794937356 | 0.1823 | 4.3601 | 1.30E-05 | 0.0017 |
| ENSG00000088888.17 | 1967.895 | 0.347616085 | 0.0799 | 4.3514 | 1.35E-05 | 0.0017 |
| ENSG00000204128.5 | 1737.721 | 0.350654822 | 0.0805 | 4.3563 | 1.32E-05 | 0.0017 |
| ENSG00000260084.1 | 26.09703 | 0.610429272 | 0.1403 | 4.3517 | 1.35E-05 | 0.0017 |
| ENSG00000160678.11 | 1588.498 | 0.698079895 | 0.1606 | 4.3468 | 1.38E-05 | 0.0017 |
| ENSG00000164691.16 | 67.40394 | -0.532085508 | 0.1225 | -4.342 | 1.41E-05 | 0.0017 |
| ENSG00000138696.10 | 709.7538 | 0.851444421 | 0.1962 | 4.3394 | 1.43E-05 | 0.0017 |
| ENSG00000225194.2 | 126.8389 | 0.650156918 | 0.1504 | 4.3236 | 1.54E-05 | 0.0018 |
| ENSG00000109107.13 | 12344.25 | 0.471810921 | 0.1094 | 4.314 | 1.60E-05 | 0.0018 |
| ENSG00000150281.6 | 62.7375 | 0.584810414 | 0.1355 | 4.3148 | 1.60E-05 | 0.0018 |
| ENSG00000164106.7 | 950.9248 | 0.486190915 | 0.1128 | 4.3105 | 1.63E-05 | 0.0018 |
| ENSG00000248050.1 | 123.055 | -0.507209673 | 0.1176 | -4.312 | 1.62E-05 | 0.0018 |
| ENSG00000251372.5 | 64.18166 | 0.726279915 | 0.1683 | 4.3147 | 1.60E-05 | 0.0018 |
| ENSG00000148672.8 | 9640.994 | 0.559949947 | 0.1302 | 4.2992 | 1.71E-05 | 0.0019 |
| ENSG00000134917.9 | 647.9403 | -0.412252336 | 0.0961 | -4.29 | 1.78E-05 | 0.002 |
| ENSG00000130876.11 | 242.5716 | 0.908811667 | 0.212 | 4.2863 | 1.82E-05 | 0.002 |
| ENSG00000164089.8 | 2420.395 | 0.9752965 | 0.228 | 4.2777 | 1.89E-05 | 0.002 |
| ENSG00000107902.13 | 1176.657 | 0.368918348 | 0.0863 | 4.2735 | 1.92E-05 | 0.0021 |
| ENSG00000163395.16 | 171.5377 | 1.01124833 | 0.2367 | 4.2714 | 1.94E-05 | 0.0021 |
| ENSG00000068078.17 | 3998.839 | 0.999833773 | 0.2342 | 4.2688 | 1.97E-05 | 0.0021 |
| ENSG00000016391.10 | 628.9317 | 0.678425059 | 0.1594 | 4.2563 | 2.08E-05 | 0.0021 |
| ENSG00000100583.4 | 86.72896 | -0.3346549 | 0.0787 | -4.255 | 2.09E-05 | 0.0021 |
| ENSG00000136197.12 | 198.6012 | -0.421934694 | 0.0991 | -4.259 | 2.06E-05 | 0.0021 |
| ENSG00000172638.12 | 514.5341 | 0.510109335 | 0.1199 | 4.253 | 2.11E-05 | 0.0021 |
| ENSG00000099139.13 | 842.8758 | 1.12900565 | 0.2658 | 4.2474 | 2.16E-05 | 0.0021 |
| ENSG00000164292.12 | 2192.048 | 0.668110501 | 0.1573 | 4.2476 | 2.16E-05 | 0.0021 |
| ENSG00000215440.11 | 779.7976 | 0.40347247 | 0.095 | 4.2452 | 2.18E-05 | 0.0021 |
| ENSG00000100427.15 | 4774.816 | 0.837386796 | 0.1975 | 4.2401 | 2.23E-05 | 0.0021 |
| ENSG00000143416.20 | 658.1303 | 0.723181446 | 0.1705 | 4.2407 | 2.23E-05 | 0.0021 |
| ENSG00000178814.16 | 198.8434 | 0.646229316 | 0.1525 | 4.2385 | 2.25E-05 | 0.0021 |
| ENSG00000119927.13 | 802.1382 | 0.807335851 | 0.1909 | 4.2281 | 2.36E-05 | 0.0022 |
| ENSG00000125285.5 | 345.8321 | 0.774295145 | 0.1832 | 4.227 | 2.37E-05 | 0.0022 |
| ENSG00000092621.11 | 1475.087 | 0.658608218 | 0.1561 | 4.2191 | 2.45E-05 | 0.0022 |
| ENSG00000115507.9 | 71.06444 | 0.738682353 | 0.1751 | 4.2198 | 2.45E-05 | 0.0022 |
| ENSG00000163702.18 | 314.8277 | 0.475861105 | 0.1128 | 4.2194 | 2.45E-05 | 0.0022 |
| ENSG00000181449.3 | 1684.84 | 0.77430641 | 0.1839 | 4.211 | 2.54E-05 | 0.0023 |
| ENSG00000253661.1 | 54.41791 | 0.658541428 | 0.1563 | 4.2127 | 2.52E-05 | 0.0023 |
| ENSG00000167601.11 | 1081.029 | 0.816203791 | 0.194 | 4.2068 | 2.59E-05 | 0.0023 |
| ENSG00000182103.4 | 580.9953 | 0.69016823 | 0.1641 | 4.205 | 2.61E-05 | 0.0023 |
| ENSG00000101144.12 | 795.3251 | 0.668741335 | 0.1596 | 4.1897 | 2.79E-05 | 0.0023 |
| ENSG00000122971.8 | 230.8269 | 0.516260879 | 0.1234 | 4.1848 | 2.85E-05 | 0.0023 |
| ENSG00000146535.13 | 2067.276 | 0.455277087 | 0.1085 | 4.1962 | 2.71E-05 | 0.0023 |
| ENSG00000153140.8 | 254.9191 | 0.262137336 | 0.0625 | 4.1956 | 2.72E-05 | 0.0023 |
| ENSG00000157890.17 | 323.2786 | 0.420121052 | 0.1004 | 4.1847 | 2.86E-05 | 0.0023 |
| ENSG00000168461.12 | 2984.66 | 0.538951214 | 0.1288 | 4.1848 | 2.85E-05 | 0.0023 |
| ENSG00000168710.17 | 7480.951 | 0.551376306 | 0.1317 | 4.1866 | 2.83E-05 | 0.0023 |
| ENSG00000184221.12 | 2455.222 | 0.511963977 | 0.1223 | 4.1861 | 2.84E-05 | 0.0023 |
| ENSG00000240184.6 | 7863.955 | 0.493385712 | 0.118 | 4.1822 | 2.89E-05 | 0.0023 |
| ENSG00000266964.5 | 1091.595 | 0.602743999 | 0.1441 | 4.1831 | 2.88E-05 | 0.0023 |
| ENSG00000002933.7 | 211.0701 | 0.743675905 | 0.1784 | 4.1687 | 3.06E-05 | 0.0023 |
| ENSG00000101400.5 | 2854.467 | 0.485115803 | 0.1163 | 4.1722 | 3.02E-05 | 0.0023 |
| ENSG00000119711.12 | 3819.452 | 0.604914193 | 0.1452 | 4.1661 | 3.10E-05 | 0.0023 |
| ENSG00000154319.14 | 798.029 | 0.701869151 | 0.1684 | 4.1682 | 3.07E-05 | 0.0023 |
| ENSG00000163285.7 | 1903.844 | 0.532060879 | 0.1274 | 4.1771 | 2.95E-05 | 0.0023 |
| ENSG00000198732.10 | 751.113 | 0.46922835 | 0.1126 | 4.1671 | 3.08E-05 | 0.0023 |
| ENSG00000203414.2 | 98.92333 | 1.108763349 | 0.2658 | 4.1712 | 3.03E-05 | 0.0023 |
| ENSG00000214290.7 | 65.38371 | 0.550811314 | 0.1318 | 4.1788 | 2.93E-05 | 0.0023 |
| ENSG00000255503.1 | 16.05253 | 0.648265311 | 0.1556 | 4.1672 | 3.08E-05 | 0.0023 |
| ENSG00000196218.12 | 1288.089 | 0.337805652 | 0.0813 | 4.1576 | 3.22E-05 | 0.0024 |
| ENSG00000161958.10 | 200.5203 | 0.493451261 | 0.1189 | 4.1518 | 3.30E-05 | 0.0025 |
| ENSG00000171517.5 | 63.01099 | 0.931041855 | 0.2243 | 4.1509 | 3.31E-05 | 0.0025 |
| ENSG00000186951.16 | 659.4588 | 0.559229201 | 0.135 | 4.1433 | 3.42E-05 | 0.0025 |
| ENSG00000104325.6 | 608.8082 | 0.398010731 | 0.0961 | 4.141 | 3.46E-05 | 0.0025 |
| ENSG00000186815.12 | 1368.809 | 0.432365358 | 0.1046 | 4.1318 | 3.60E-05 | 0.0026 |
| ENSG00000166839.16 | 676.4015 | 0.336762854 | 0.0817 | 4.1244 | 3.72E-05 | 0.0027 |
| ENSG00000147852.15 | 917.3519 | -0.316949381 | 0.0769 | -4.121 | 3.77E-05 | 0.0027 |
| ENSG00000271860.6 | 87.48412 | 0.441909378 | 0.1073 | 4.1171 | 3.84E-05 | 0.0027 |
| ENSG00000106003.12 | 245.7343 | 0.744894137 | 0.181 | 4.1145 | 3.88E-05 | 0.0027 |
| ENSG00000176399.3 | 13.16854 | 0.83833701 | 0.2039 | 4.1114 | 3.93E-05 | 0.0028 |
| ENSG00000165475.13 | 848.4398 | 0.372219897 | 0.0906 | 4.1091 | 3.97E-05 | 0.0028 |
| ENSG00000182902.13 | 1982.591 | 0.660328265 | 0.1608 | 4.1053 | 4.04E-05 | 0.0028 |
| ENSG00000185033.14 | 756.5435 | 0.42850694 | 0.1043 | 4.1065 | 4.02E-05 | 0.0028 |
| ENSG00000132613.14 | 10017.71 | 0.478360688 | 0.1167 | 4.0981 | 4.17E-05 | 0.0028 |
| ENSG00000166925.8 | 2721.067 | 0.574323275 | 0.1403 | 4.0924 | 4.27E-05 | 0.0029 |
| ENSG00000183580.9 | 396.0073 | 0.6059737 | 0.1481 | 4.0918 | 4.28E-05 | 0.0029 |
| ENSG00000269293.2 | 151.9808 | 0.315415319 | 0.0771 | 4.0934 | 4.25E-05 | 0.0029 |
| ENSG00000121742.16 | 897.2621 | 1.011184292 | 0.2473 | 4.0897 | 4.32E-05 | 0.0029 |
| ENSG00000130203.9 | 10421.98 | 0.888667423 | 0.2175 | 4.0854 | 4.40E-05 | 0.0029 |
| ENSG00000185960.13 | 63.91012 | 1.39584739 | 0.3417 | 4.0851 | 4.41E-05 | 0.0029 |
| ENSG00000185960.13_PAR_Y | 63.91012 | 1.39584739 | 0.3417 | 4.0851 | 4.41E-05 | 0.0029 |
| ENSG00000110900.14 | 227.1915 | 0.528204024 | 0.1294 | 4.0811 | 4.48E-05 | 0.0029 |
| ENSG00000226191.3 | 15.8543 | -0.883535025 | 0.2165 | -4.081 | 4.48E-05 | 0.0029 |
| ENSG00000228288.6 | 32.94073 | 0.607452047 | 0.1489 | 4.0783 | 4.54E-05 | 0.0029 |
| ENSG00000043355.11 | 247.2894 | 0.654853198 | 0.1612 | 4.0626 | 4.85E-05 | 0.0029 |
| ENSG00000105854.12 | 2458.66 | 0.629183686 | 0.1548 | 4.0639 | 4.83E-05 | 0.0029 |
| ENSG00000112333.11 | 501.4093 | 0.669098963 | 0.1646 | 4.0639 | 4.83E-05 | 0.0029 |
| ENSG00000126778.8 | 15.72251 | 0.914637626 | 0.2247 | 4.0698 | 4.70E-05 | 0.0029 |
| ENSG00000134533.6 | 270.5631 | 0.429042587 | 0.1054 | 4.0696 | 4.71E-05 | 0.0029 |
| ENSG00000148411.7 | 3536.6 | 0.484548939 | 0.1191 | 4.0689 | 4.72E-05 | 0.0029 |
| ENSG00000157551.17 | 110.5432 | 0.477018002 | 0.1175 | 4.0612 | 4.88E-05 | 0.0029 |
| ENSG00000179761.11 | 178.333 | 0.691766184 | 0.1701 | 4.0679 | 4.74E-05 | 0.0029 |
| ENSG00000180769.8 | 369.574 | 0.499783087 | 0.1231 | 4.0615 | 4.88E-05 | 0.0029 |
| ENSG00000254245.2 | 232.4474 | 0.879774821 | 0.2163 | 4.0679 | 4.74E-05 | 0.0029 |
| ENSG00000259065.1 | 15.44446 | 0.856526678 | 0.2106 | 4.0673 | 4.76E-05 | 0.0029 |
| ENSG00000102935.11 | 546.8663 | 0.454929959 | 0.1121 | 4.0584 | 4.94E-05 | 0.0029 |
| ENSG00000182175.13 | 1707.016 | 0.456966672 | 0.1126 | 4.0583 | 4.94E-05 | 0.0029 |
| ENSG00000144040.12 | 5098.027 | 0.577290044 | 0.1424 | 4.0533 | 5.05E-05 | 0.003 |
| ENSG00000281453.1 | 120.2169 | 0.597651977 | 0.1474 | 4.0539 | 5.04E-05 | 0.003 |
| ENSG00000166535.19 | 142.7777 | 0.571173243 | 0.141 | 4.0511 | 5.10E-05 | 0.003 |
| ENSG00000018625.14 | 18477.44 | 0.745858629 | 0.1843 | 4.0477 | 5.17E-05 | 0.003 |
| ENSG00000111907.20 | 949.1767 | 0.632779447 | 0.1565 | 4.0441 | 5.25E-05 | 0.003 |
| ENSG00000161509.13 | 1150.281 | 0.841509024 | 0.2081 | 4.043 | 5.28E-05 | 0.003 |
| ENSG00000197921.5 | 146.6547 | 1.016729392 | 0.2514 | 4.0439 | 5.26E-05 | 0.003 |
| ENSG00000198695.2 | 29121.03 | 0.810562992 | 0.2007 | 4.0396 | 5.35E-05 | 0.003 |
| ENSG00000146005.3 | 2580.435 | 0.497124849 | 0.1231 | 4.0383 | 5.38E-05 | 0.003 |
| ENSG00000101198.14 | 568.7849 | 0.698285287 | 0.1731 | 4.0347 | 5.47E-05 | 0.003 |
| ENSG00000154134.14 | 414.7111 | 0.45033489 | 0.1116 | 4.0357 | 5.44E-05 | 0.003 |
| ENSG00000181192.11 | 531.2712 | 0.408800065 | 0.1013 | 4.034 | 5.48E-05 | 0.003 |
| ENSG00000228672.3 | 118.5856 | 0.355503881 | 0.0882 | 4.0319 | 5.53E-05 | 0.003 |
| ENSG00000135094.10 | 151.5535 | 0.854189719 | 0.212 | 4.0295 | 5.59E-05 | 0.003 |
| ENSG00000164050.12 | 2508.085 | 0.657880416 | 0.1633 | 4.0292 | 5.60E-05 | 0.003 |
| ENSG00000146409.10 | 726.6475 | 0.444207668 | 0.1105 | 4.0216 | 5.78E-05 | 0.0031 |
| ENSG00000256196.1 | 53.77714 | -0.466857951 | 0.1161 | -4.02 | 5.82E-05 | 0.0031 |
| ENSG00000101439.8 | 17007.3 | 0.687486857 | 0.1713 | 4.0132 | 5.99E-05 | 0.0032 |
| ENSG00000187091.13 | 327.7333 | 0.667972733 | 0.1665 | 4.011 | 6.05E-05 | 0.0032 |
| ENSG00000256463.8 | 322.5545 | 0.722992939 | 0.1805 | 4.0053 | 6.19E-05 | 0.0033 |
| ENSG00000093010.12 | 1061.807 | 0.378968314 | 0.0947 | 4.0006 | 6.32E-05 | 0.0033 |
| ENSG00000150893.10 | 146.9877 | 0.831490123 | 0.2077 | 4.0024 | 6.27E-05 | 0.0033 |
| ENSG00000152990.13 | 1010.173 | 0.693787756 | 0.1734 | 4.0006 | 6.32E-05 | 0.0033 |
| ENSG00000135750.14 | 1823.057 | -0.469217257 | 0.1177 | -3.986 | 6.73E-05 | 0.0035 |
| ENSG00000223802.7 | 3984.033 | 0.463309297 | 0.1163 | 3.984 | 6.78E-05 | 0.0035 |
| ENSG00000128309.16 | 302.4299 | 0.382333374 | 0.096 | 3.9822 | 6.83E-05 | 0.0035 |
| ENSG00000106689.10 | 1413.444 | 0.400831006 | 0.1009 | 3.9713 | 7.15E-05 | 0.0037 |
| ENSG00000000003.14 | 290.5289 | 0.35234314 | 0.0889 | 3.9645 | 7.35E-05 | 0.0037 |
| ENSG00000183779.6 | 683.1492 | 0.425193153 | 0.1074 | 3.9606 | 7.48E-05 | 0.0038 |
| ENSG00000100979.14 | 1009.939 | 0.657428374 | 0.1662 | 3.9552 | 7.65E-05 | 0.0038 |
| ENSG00000121270.15 | 31.92215 | 0.580951024 | 0.1468 | 3.9568 | 7.60E-05 | 0.0038 |
| ENSG00000125462.16 | 6885.226 | 0.521271966 | 0.1318 | 3.9561 | 7.62E-05 | 0.0038 |
| ENSG00000129353.14 | 2159.652 | 0.414577766 | 0.1051 | 3.9441 | 8.01E-05 | 0.0038 |
| ENSG00000130055.13 | 135.9079 | 0.633690308 | 0.1608 | 3.942 | 8.08E-05 | 0.0038 |
| ENSG00000134569.9 | 2250.172 | 0.713490028 | 0.1806 | 3.9515 | 7.77E-05 | 0.0038 |
| ENSG00000157557.11 | 2818.806 | -0.337551602 | 0.0855 | -3.947 | 7.90E-05 | 0.0038 |
| ENSG00000170075.8 | 2953.557 | 0.841145294 | 0.2132 | 3.946 | 7.95E-05 | 0.0038 |
| ENSG00000170370.11 | 723.8971 | 0.65095723 | 0.165 | 3.9444 | 8.00E-05 | 0.0038 |
| ENSG00000186603.5 | 28.97122 | 0.553437142 | 0.1404 | 3.9426 | 8.06E-05 | 0.0038 |
| ENSG00000186998.15 | 708.2768 | 0.487040184 | 0.1235 | 3.9439 | 8.02E-05 | 0.0038 |
| ENSG00000196668.3 | 51.13421 | 0.519294414 | 0.1316 | 3.9453 | 7.97E-05 | 0.0038 |
| ENSG00000198886.2 | 451250.1 | 0.542290743 | 0.1374 | 3.9472 | 7.91E-05 | 0.0038 |
| ENSG00000107317.11 | 11635.76 | 0.615252526 | 0.1562 | 3.9387 | 8.19E-05 | 0.0039 |
| ENSG00000198157.10 | 264.0034 | 0.26807481 | 0.0681 | 3.9337 | 8.36E-05 | 0.0039 |
| ENSG00000279672.1 | 174.9132 | 0.706079058 | 0.1795 | 3.9338 | 8.36E-05 | 0.0039 |
| ENSG00000184154.13 | 280.6833 | 0.261988608 | 0.0667 | 3.9307 | 8.47E-05 | 0.004 |
| ENSG00000167210.16 | 43.70135 | -0.516979171 | 0.1317 | -3.927 | 8.61E-05 | 0.004 |
| ENSG00000181856.14 | 62.75413 | 0.828849976 | 0.2111 | 3.9258 | 8.64E-05 | 0.004 |
| ENSG00000248713.1 | 28.08973 | -0.89323771 | 0.2275 | -3.926 | 8.65E-05 | 0.004 |
| ENSG00000179627.9 | 18.69501 | 0.560025691 | 0.1427 | 3.9242 | 8.70E-05 | 0.004 |
| ENSG00000165474.5 | 32.47352 | 1.132234445 | 0.2887 | 3.922 | 8.78E-05 | 0.004 |
| ENSG00000165458.13 | 1029.338 | 0.438212805 | 0.1119 | 3.916 | 9.00E-05 | 0.0041 |
| ENSG00000122986.13 | 118.1383 | 0.527000501 | 0.1347 | 3.9113 | 9.18E-05 | 0.0041 |
| ENSG00000146250.6 | 180.7432 | 0.936059821 | 0.2393 | 3.9123 | 9.14E-05 | 0.0041 |
| ENSG00000089335.20 | 3003.436 | 0.678626042 | 0.1741 | 3.8971 | 9.74E-05 | 0.0044 |
| ENSG00000178537.9 | 174.5833 | 0.493003447 | 0.1265 | 3.8968 | 9.75E-05 | 0.0044 |
| ENSG00000236255.1 | 1130.596 | 1.605895886 | 0.4126 | 3.8917 | 9.95E-05 | 0.0044 |
| ENSG00000179399.14 | 368.3619 | 0.653915423 | 0.1682 | 3.8868 | 0.0001 | 0.0045 |
| ENSG00000267385.1 | 23.34444 | 1.027787327 | 0.2645 | 3.8861 | 0.0001 | 0.0045 |
| ENSG00000144908.13 | 1325.601 | 0.821804597 | 0.2116 | 3.8832 | 0.0001 | 0.0045 |
| ENSG00000153446.15 | 231.096 | 0.743443572 | 0.1916 | 3.8805 | 0.0001 | 0.0046 |
| ENSG00000227640.2 | 164.5085 | 0.656893763 | 0.1694 | 3.8788 | 0.0001 | 0.0046 |
| ENSG00000177045.7 | 70.26145 | 0.810146887 | 0.2092 | 3.8728 | 0.00011 | 0.0047 |
| ENSG00000166123.13 | 2252.784 | 0.457709243 | 0.1183 | 3.8706 | 0.00011 | 0.0047 |
| ENSG00000128602.9 | 181.3846 | 0.718590834 | 0.1859 | 3.8648 | 0.00011 | 0.0048 |
| ENSG00000064655.18 | 152.3829 | 0.766589998 | 0.1988 | 3.8568 | 0.00011 | 0.0049 |
| ENSG00000169006.6 | 578.5476 | 0.841871133 | 0.2184 | 3.8545 | 0.00012 | 0.0049 |
| ENSG00000204580.12 | 2531.971 | 0.536956665 | 0.1393 | 3.8544 | 0.00012 | 0.0049 |
| ENSG00000178974.9 | 1349.209 | -0.406271125 | 0.1054 | -3.853 | 0.00012 | 0.0049 |
| ENSG00000103742.11 | 219.1352 | 0.560753313 | 0.1458 | 3.8458 | 0.00012 | 0.0051 |
| ENSG00000114315.3 | 358.0531 | 0.882259958 | 0.2294 | 3.8452 | 0.00012 | 0.0051 |
| ENSG00000179388.8 | 2443.505 | -0.382976867 | 0.0996 | -3.845 | 0.00012 | 0.0051 |
| ENSG00000136114.15 | 235.2386 | 0.533758255 | 0.1389 | 3.8422 | 0.00012 | 0.0051 |
| ENSG00000126785.12 | 332.9438 | 0.579846687 | 0.151 | 3.8412 | 0.00012 | 0.0051 |
| ENSG00000134376.14 | 393.6164 | 0.490821323 | 0.1279 | 3.8385 | 0.00012 | 0.0051 |
| ENSG00000149782.11 | 272.6127 | 0.571078128 | 0.1488 | 3.8368 | 0.00012 | 0.0051 |
| ENSG00000204396.10 | 310.4841 | -0.414675757 | 0.1081 | -3.837 | 0.00012 | 0.0051 |
| ENSG00000105852.10 | 30.10673 | 0.749960838 | 0.1957 | 3.8329 | 0.00013 | 0.0051 |
| ENSG00000189221.9 | 993.439 | 0.338213625 | 0.0883 | 3.8316 | 0.00013 | 0.0051 |
| ENSG00000196091.13 | 646.0416 | 0.518218192 | 0.1353 | 3.8313 | 0.00013 | 0.0051 |
| ENSG00000276649.1 | 140.0724 | 0.338434058 | 0.0883 | 3.8323 | 0.00013 | 0.0051 |
| ENSG00000100767.15 | 535.6679 | 0.775702264 | 0.2026 | 3.8292 | 0.00013 | 0.0052 |
| ENSG00000274956.2 | 1748.869 | 0.650802388 | 0.17 | 3.829 | 0.00013 | 0.0052 |
| ENSG00000115380.19 | 886.5195 | 0.921200039 | 0.2407 | 3.8278 | 0.00013 | 0.0052 |
| ENSG00000204347.3 | 59.81885 | 0.787335669 | 0.2058 | 3.8261 | 0.00013 | 0.0052 |
| ENSG00000197961.11 | 1072.735 | 0.412099459 | 0.1077 | 3.825 | 0.00013 | 0.0052 |
| ENSG00000100106.19 | 411.1776 | 0.374552375 | 0.098 | 3.821 | 0.00013 | 0.0052 |
| ENSG00000261026.1 | 23.76471 | -0.679090259 | 0.1778 | -3.819 | 0.00013 | 0.0053 |
| ENSG00000076351.12 | 493.8901 | 0.259642145 | 0.0681 | 3.8111 | 0.00014 | 0.0053 |
| ENSG00000114738.10 | 226.4984 | 0.582936399 | 0.1531 | 3.8087 | 0.00014 | 0.0053 |
| ENSG00000123080.10 | 117.3642 | 0.538258734 | 0.1413 | 3.8094 | 0.00014 | 0.0053 |
| ENSG00000168385.17 | 5382.716 | 0.356046048 | 0.0934 | 3.8123 | 0.00014 | 0.0053 |
| ENSG00000180537.12 | 432.9882 | 0.524322336 | 0.1376 | 3.8095 | 0.00014 | 0.0053 |
| ENSG00000234899.9 | 132.078 | 0.378283932 | 0.0992 | 3.8138 | 0.00014 | 0.0053 |
| ENSG00000243069.7 | 446.8998 | 0.437721802 | 0.1149 | 3.8105 | 0.00014 | 0.0053 |
| ENSG00000134595.8 | 21.3872 | 0.716620057 | 0.1883 | 3.8052 | 0.00014 | 0.0054 |
| ENSG00000125144.13 | 493.4402 | 0.751133524 | 0.1976 | 3.8015 | 0.00014 | 0.0054 |
| ENSG00000149090.11 | 680.6047 | 0.64464161 | 0.1695 | 3.8022 | 0.00014 | 0.0054 |
| ENSG00000129244.8 | 9124.93 | 0.696102341 | 0.1833 | 3.7967 | 0.00015 | 0.0055 |
| ENSG00000270804.1 | 75.70724 | 0.378300021 | 0.0996 | 3.7967 | 0.00015 | 0.0055 |
| ENSG00000137285.9 | 5706.905 | 0.737859184 | 0.1945 | 3.7931 | 0.00015 | 0.0056 |
| ENSG00000177189.12 | 1466.781 | -0.275717717 | 0.0727 | -3.79 | 0.00015 | 0.0056 |
| ENSG00000021300.13 | 8237.922 | 0.431833258 | 0.114 | 3.7869 | 0.00015 | 0.0056 |
| ENSG00000140022.9 | 789.838 | 0.729272727 | 0.1926 | 3.7865 | 0.00015 | 0.0056 |
| ENSG00000146648.16 | 723.0516 | 0.661790817 | 0.1748 | 3.7862 | 0.00015 | 0.0056 |
| ENSG00000148498.15 | 327.7184 | 0.539117043 | 0.1426 | 3.7815 | 0.00016 | 0.0057 |
| ENSG00000089472.16 | 496.1221 | 0.690815589 | 0.1828 | 3.7792 | 0.00016 | 0.0057 |
| ENSG00000109062.10 | 1045.581 | 0.701924738 | 0.186 | 3.7748 | 0.00016 | 0.0057 |
| ENSG00000135821.17 | 35484.82 | 0.737413768 | 0.1954 | 3.7748 | 0.00016 | 0.0057 |
| ENSG00000138029.13 | 1739.677 | 0.476794572 | 0.1263 | 3.7763 | 0.00016 | 0.0057 |
| ENSG00000156219.16 | 194.4457 | -0.40395722 | 0.107 | -3.776 | 0.00016 | 0.0057 |
| ENSG00000171388.11 | 430.5822 | 0.556214934 | 0.1473 | 3.7762 | 0.00016 | 0.0057 |
| ENSG00000233850.1 | 12.06498 | 0.696103577 | 0.1843 | 3.7768 | 0.00016 | 0.0057 |
| ENSG00000079337.15 | 1215.326 | 0.406285106 | 0.1077 | 3.7725 | 0.00016 | 0.0057 |
| ENSG00000117592.8 | 2062.079 | 0.360840423 | 0.0957 | 3.7699 | 0.00016 | 0.0057 |
| ENSG00000169738.7 | 814.1316 | 0.397174675 | 0.1054 | 3.7697 | 0.00016 | 0.0057 |
| ENSG00000184313.19 | 393.2561 | 0.417756848 | 0.1108 | 3.7693 | 0.00016 | 0.0057 |
| ENSG00000272338.2 | 39.11133 | 0.585317571 | 0.1552 | 3.7725 | 0.00016 | 0.0057 |
| ENSG00000273091.1 | 19.30018 | 0.651998032 | 0.1729 | 3.7709 | 0.00016 | 0.0057 |
| ENSG00000081248.10 | 51.15722 | -0.601551476 | 0.16 | -3.759 | 0.00017 | 0.0059 |
| ENSG00000114115.9 | 262.926 | 0.490259942 | 0.1304 | 3.7591 | 0.00017 | 0.0059 |
| ENSG00000116580.18 | 2250.78 | 0.562858195 | 0.1499 | 3.7547 | 0.00017 | 0.006 |
| ENSG00000075073.14 | 129.2261 | -0.499599413 | 0.1333 | -3.747 | 0.00018 | 0.0062 |
| ENSG00000156042.17 | 267.5233 | 0.382033502 | 0.102 | 3.7472 | 0.00018 | 0.0062 |
| ENSG00000170425.3 | 160.9039 | 0.650389852 | 0.1736 | 3.7473 | 0.00018 | 0.0062 |
| ENSG00000068976.13 | 544.8239 | 0.716892538 | 0.1916 | 3.7421 | 0.00018 | 0.0062 |
| ENSG00000082196.20 | 181.7521 | 0.460360269 | 0.123 | 3.7421 | 0.00018 | 0.0062 |
| ENSG00000127249.14 | 568.963 | 0.739629186 | 0.1976 | 3.7438 | 0.00018 | 0.0062 |
| ENSG00000163083.5 | 117.9638 | 0.5810006 | 0.1552 | 3.7435 | 0.00018 | 0.0062 |
| ENSG00000163346.16 | 2517.992 | 0.722684516 | 0.193 | 3.7453 | 0.00018 | 0.0062 |
| ENSG00000145555.14 | 2400.925 | 0.550233837 | 0.1472 | 3.7374 | 0.00019 | 0.0063 |
| ENSG00000151322.18 | 1413.516 | 0.486088444 | 0.1301 | 3.7366 | 0.00019 | 0.0063 |
| ENSG00000185432.11 | 2388.743 | 0.672365031 | 0.1799 | 3.7374 | 0.00019 | 0.0063 |
| ENSG00000088280.18 | 498.7201 | 0.383843574 | 0.1029 | 3.7307 | 0.00019 | 0.0063 |
| ENSG00000136235.15 | 568.0418 | 0.730863019 | 0.1959 | 3.7314 | 0.00019 | 0.0063 |
| ENSG00000170989.8 | 1279.364 | 0.770215123 | 0.2064 | 3.7318 | 0.00019 | 0.0063 |
| ENSG00000267014.5 | 34.81448 | 0.676846983 | 0.1813 | 3.7326 | 0.00019 | 0.0063 |
| ENSG00000169439.11 | 1026.253 | 0.668147382 | 0.1792 | 3.7292 | 0.00019 | 0.0063 |
| ENSG00000238230.1 | 20.42442 | 1.009220528 | 0.271 | 3.7247 | 0.0002 | 0.0064 |
| ENSG00000166682.10 | 168.7252 | 0.531170475 | 0.1428 | 3.7202 | 0.0002 | 0.0065 |
| ENSG00000157184.6 | 245.8209 | 0.359440872 | 0.0968 | 3.715 | 0.0002 | 0.0066 |
| ENSG00000168309.16 | 21561 | 0.551161903 | 0.1483 | 3.7161 | 0.0002 | 0.0066 |
| ENSG00000258010.3 | 54.26339 | 0.564293151 | 0.1519 | 3.7156 | 0.0002 | 0.0066 |
| ENSG00000101542.9 | 655.4343 | 0.424531084 | 0.1144 | 3.7098 | 0.00021 | 0.0067 |
| ENSG00000261177.1 | 24.78307 | 0.767677592 | 0.2072 | 3.7041 | 0.00021 | 0.0068 |
| ENSG00000110042.7 | 1345.151 | -0.272015474 | 0.0735 | -3.702 | 0.00021 | 0.0069 |
| ENSG00000119471.14 | 1120.676 | 0.480566677 | 0.13 | 3.6981 | 0.00022 | 0.007 |
| ENSG00000162738.5 | 288.2174 | 0.408520377 | 0.1105 | 3.6968 | 0.00022 | 0.007 |
| ENSG00000279118.1 | 466.5558 | 0.647297381 | 0.1752 | 3.6953 | 0.00022 | 0.007 |
| ENSG00000103260.8 | 2531.822 | 0.530698606 | 0.1438 | 3.6901 | 0.00022 | 0.0071 |
| ENSG00000136205.16 | 2962.642 | 0.601070702 | 0.1628 | 3.6916 | 0.00022 | 0.0071 |
| ENSG00000281183.1 | 254.2663 | -0.50425214 | 0.1366 | -3.69 | 0.00022 | 0.0071 |
| ENSG00000143819.12 | 1609.618 | 0.608374645 | 0.165 | 3.6864 | 0.00023 | 0.0071 |
| ENSG00000162407.8 | 2750.831 | 0.805592095 | 0.2186 | 3.685 | 0.00023 | 0.0071 |
| ENSG00000183579.15 | 744.6444 | 0.613809728 | 0.1665 | 3.6873 | 0.00023 | 0.0071 |
| ENSG00000184925.11 | 136.034 | 0.55929202 | 0.1518 | 3.6854 | 0.00023 | 0.0071 |
| ENSG00000203396.3 | 170.1716 | 1.314275206 | 0.3567 | 3.6847 | 0.00023 | 0.0071 |
| ENSG00000213626.11 | 573.5175 | -0.371827843 | 0.1009 | -3.686 | 0.00023 | 0.0071 |
| ENSG00000100968.13 | 335.8641 | 0.633103094 | 0.1719 | 3.6835 | 0.00023 | 0.0071 |
| ENSG00000271430.1 | 336.1798 | -0.363399839 | 0.0987 | -3.681 | 0.00023 | 0.0072 |
| ENSG00000204099.11 | 183.044 | 0.594732386 | 0.1616 | 3.6799 | 0.00023 | 0.0072 |
| ENSG00000105894.11 | 3203.992 | 0.543652305 | 0.1478 | 3.6789 | 0.00023 | 0.0072 |
| ENSG00000215915.9 | 93.24601 | 0.467868096 | 0.1272 | 3.6772 | 0.00024 | 0.0072 |
| ENSG00000127418.14 | 601.9529 | 0.708273108 | 0.1932 | 3.6669 | 0.00025 | 0.0074 |
| ENSG00000171903.16 | 129.3399 | 0.745310493 | 0.2032 | 3.6671 | 0.00025 | 0.0074 |
| ENSG00000197496.5 | 114.8715 | 0.702967648 | 0.1917 | 3.6667 | 0.00025 | 0.0074 |
| ENSG00000189171.14 | 680.9856 | 0.524945208 | 0.1434 | 3.6607 | 0.00025 | 0.0076 |
| ENSG00000144485.10 | 431.4361 | 0.419026223 | 0.1145 | 3.6586 | 0.00025 | 0.0076 |
| ENSG00000185338.4 | 9.129556 | 0.988950786 | 0.2703 | 3.6594 | 0.00025 | 0.0076 |
| ENSG00000101850.12 | 77.92988 | 0.649758547 | 0.1777 | 3.6575 | 0.00025 | 0.0076 |
| ENSG00000230839.1 | 31.75432 | 0.51097764 | 0.1397 | 3.6564 | 0.00026 | 0.0076 |
| ENSG00000170340.10 | 507.2153 | -0.378780495 | 0.1037 | -3.654 | 0.00026 | 0.0077 |
| ENSG00000168003.16 | 1849.312 | 0.34591413 | 0.0947 | 3.6516 | 0.00026 | 0.0077 |
| ENSG00000165584.15 | 9.342034 | 0.749016226 | 0.2053 | 3.6488 | 0.00026 | 0.0077 |
| ENSG00000213366.12 | 1133.058 | 0.324686816 | 0.089 | 3.6487 | 0.00026 | 0.0077 |
| ENSG00000262655.3 | 1319.474 | 0.58235329 | 0.1596 | 3.6494 | 0.00026 | 0.0077 |
| ENSG00000104760.16 | 17.32124 | 0.862722525 | 0.2365 | 3.6475 | 0.00026 | 0.0078 |
| ENSG00000237887.1 | 40.07037 | 0.632151217 | 0.1733 | 3.647 | 0.00027 | 0.0078 |
| ENSG00000164904.16 | 2135.145 | 0.458328515 | 0.1258 | 3.6446 | 0.00027 | 0.0078 |
| ENSG00000165730.14 | 676.1466 | 0.490207773 | 0.1345 | 3.6447 | 0.00027 | 0.0078 |
| ENSG00000204556.4 | 31.78005 | 0.572225465 | 0.157 | 3.6448 | 0.00027 | 0.0078 |
| ENSG00000092820.17 | 2683.501 | 0.646632447 | 0.1775 | 3.6429 | 0.00027 | 0.0078 |
| ENSG00000166183.15 | 22.95042 | 0.630665087 | 0.1734 | 3.6381 | 0.00027 | 0.0079 |
| ENSG00000117834.12 | 18.65488 | 0.680105689 | 0.187 | 3.6361 | 0.00028 | 0.008 |
| ENSG00000074047.21 | 55.00242 | 0.910099872 | 0.2504 | 3.634 | 0.00028 | 0.008 |
| ENSG00000254584.1 | 151.9211 | 1.122493986 | 0.3092 | 3.6301 | 0.00028 | 0.0081 |
| ENSG00000141756.18 | 378.8 | 0.641283891 | 0.1767 | 3.6292 | 0.00028 | 0.0081 |
| ENSG00000282851.1 | 77.08729 | 0.336963266 | 0.093 | 3.6235 | 0.00029 | 0.0083 |
| ENSG00000188493.14 | 95.04198 | 0.400458977 | 0.1106 | 3.6219 | 0.00029 | 0.0083 |
| ENSG00000016402.13 | 72.5259 | 0.64615279 | 0.1785 | 3.6208 | 0.00029 | 0.0083 |
| ENSG00000160781.15 | 4116.292 | 0.53035417 | 0.1465 | 3.6195 | 0.0003 | 0.0083 |
| ENSG00000176887.6 | 242.3897 | 0.402992107 | 0.1114 | 3.6163 | 0.0003 | 0.0084 |
| ENSG00000138604.9 | 1096.086 | -0.387256418 | 0.1072 | -3.614 | 0.0003 | 0.0085 |
| ENSG00000167107.12 | 203.0387 | 0.510911286 | 0.1415 | 3.6109 | 0.00031 | 0.0085 |
| ENSG00000072858.10 | 919.583 | -0.342010249 | 0.095 | -3.601 | 0.00032 | 0.0085 |
| ENSG00000113161.15 | 3010.163 | -0.383398729 | 0.1064 | -3.604 | 0.00031 | 0.0085 |
| ENSG00000133048.12 | 372.8535 | 0.864076082 | 0.2396 | 3.6064 | 0.00031 | 0.0085 |
| ENSG00000134873.9 | 550.956 | 0.763896807 | 0.2121 | 3.601 | 0.00032 | 0.0085 |
| ENSG00000135898.9 | 23.23787 | -0.514766538 | 0.1429 | -3.601 | 0.00032 | 0.0085 |
| ENSG00000173548.8 | 374.0284 | 0.501444979 | 0.1391 | 3.606 | 0.00031 | 0.0085 |
| ENSG00000176928.5 | 396.2692 | -0.508995563 | 0.1414 | -3.601 | 0.00032 | 0.0085 |
| ENSG00000185942.11 | 67.47485 | 0.578738864 | 0.1607 | 3.6016 | 0.00032 | 0.0085 |
| ENSG00000210082.2 | 1337848 | 0.455980192 | 0.1266 | 3.6031 | 0.00031 | 0.0085 |
| ENSG00000230062.5 | 15.63646 | -0.707932879 | 0.1965 | -3.602 | 0.00032 | 0.0085 |
| ENSG00000238099.2 | 12.21722 | 0.858871956 | 0.2384 | 3.602 | 0.00032 | 0.0085 |
| ENSG00000257261.5 | 17.05076 | 0.57705202 | 0.1602 | 3.6021 | 0.00032 | 0.0085 |
| ENSG00000267534.2 | 294.2147 | 1.101074242 | 0.3055 | 3.6041 | 0.00031 | 0.0085 |
| ENSG00000279457.3 | 328.7888 | 0.358553669 | 0.0995 | 3.6032 | 0.00031 | 0.0085 |
| ENSG00000178075.19 | 500.202 | 0.523968694 | 0.1456 | 3.5989 | 0.00032 | 0.0086 |
| ENSG00000006756.15 | 294.574 | 0.314741351 | 0.0875 | 3.5951 | 0.00032 | 0.0086 |
| ENSG00000103811.15 | 681.5277 | 0.5792423 | 0.1612 | 3.5931 | 0.00033 | 0.0086 |
| ENSG00000106571.12 | 241.3095 | 0.833036944 | 0.2316 | 3.5975 | 0.00032 | 0.0086 |
| ENSG00000116711.9 | 114.2159 | -0.421179442 | 0.1171 | -3.597 | 0.00032 | 0.0086 |
| ENSG00000156076.9 | 1289.598 | 0.87841316 | 0.2443 | 3.5956 | 0.00032 | 0.0086 |
| ENSG00000166819.11 | 77.02226 | 0.659851856 | 0.1837 | 3.5929 | 0.00033 | 0.0086 |
| ENSG00000169203.16 | 1191.861 | 0.412616382 | 0.1148 | 3.5938 | 0.00033 | 0.0086 |
| ENSG00000196873.15 | 881.5314 | 0.464165599 | 0.1291 | 3.5944 | 0.00033 | 0.0086 |
| ENSG00000235989.3 | 39.58766 | 0.732352774 | 0.2037 | 3.5951 | 0.00032 | 0.0086 |
| ENSG00000108604.15 | 287.7159 | 0.28933074 | 0.0806 | 3.5896 | 0.00033 | 0.0086 |
| ENSG00000198786.2 | 169124.6 | 0.515568804 | 0.1436 | 3.591 | 0.00033 | 0.0086 |
| ENSG00000236790.5 | 72.07537 | 0.572401292 | 0.1594 | 3.5901 | 0.00033 | 0.0086 |
| ENSG00000268038.1 | 36.3263 | 0.917431736 | 0.2556 | 3.5894 | 0.00033 | 0.0086 |
| ENSG00000072201.13 | 1719.376 | -0.372896937 | 0.104 | -3.587 | 0.00033 | 0.0087 |
| ENSG00000135424.15 | 1108.925 | 0.452927624 | 0.1264 | 3.5843 | 0.00034 | 0.0087 |
| ENSG00000087076.8 | 251.0114 | 0.445289146 | 0.1243 | 3.5826 | 0.00034 | 0.0088 |
| ENSG00000252202.1 | 11.54209 | 0.580140611 | 0.1622 | 3.5758 | 0.00035 | 0.009 |
| ENSG00000025423.11 | 241.2882 | 0.655639082 | 0.1834 | 3.5745 | 0.00035 | 0.009 |
| ENSG00000076248.10 | 469.188 | 0.333864909 | 0.0935 | 3.5714 | 0.00036 | 0.009 |
| ENSG00000135069.13 | 1887.846 | 0.556641387 | 0.1559 | 3.5707 | 0.00036 | 0.009 |
| ENSG00000160194.17 | 1459.732 | 0.443896919 | 0.1243 | 3.5701 | 0.00036 | 0.009 |
| ENSG00000167543.15 | 230.2532 | 0.421215218 | 0.1179 | 3.572 | 0.00035 | 0.009 |
| ENSG00000259479.6 | 42.88381 | -1.413301746 | 0.3958 | -3.571 | 0.00036 | 0.009 |
| ENSG00000080573.6 | 2518.775 | 0.504985064 | 0.1415 | 3.5682 | 0.00036 | 0.0091 |
| ENSG00000103202.12 | 806.0529 | 0.299543626 | 0.084 | 3.5677 | 0.00036 | 0.0091 |
| ENSG00000110436.11 | 59030.34 | 0.799737892 | 0.2242 | 3.567 | 0.00036 | 0.0091 |
| ENSG00000198417.6 | 280.0516 | 0.604014981 | 0.1692 | 3.5691 | 0.00036 | 0.0091 |
| ENSG00000250366.2 | 943.7866 | -0.433837448 | 0.1217 | -3.566 | 0.00036 | 0.0091 |
| ENSG00000262528.2 | 40.91312 | 0.466695055 | 0.1309 | 3.5663 | 0.00036 | 0.0091 |
| ENSG00000248508.6 | 113.6354 | 0.397181861 | 0.1115 | 3.5629 | 0.00037 | 0.0091 |
| ENSG00000278224.4 | 25.76837 | 0.491008188 | 0.1378 | 3.5632 | 0.00037 | 0.0091 |
| ENSG00000135744.7 | 2616.248 | 0.716406042 | 0.2013 | 3.5596 | 0.00037 | 0.0092 |
| ENSG00000147853.16 | 1288.71 | 0.281752117 | 0.0792 | 3.5565 | 0.00038 | 0.0093 |
| ENSG00000170412.16 | 165.4244 | 0.653971789 | 0.1839 | 3.556 | 0.00038 | 0.0093 |
| ENSG00000235711.4 | 178.3638 | -0.575659722 | 0.1619 | -3.556 | 0.00038 | 0.0093 |
| ENSG00000115641.18 | 486.509 | -0.33331899 | 0.0938 | -3.555 | 0.00038 | 0.0093 |
| ENSG00000109113.18 | 318.8599 | 0.6513732 | 0.1833 | 3.5531 | 0.00038 | 0.0093 |
| ENSG00000140459.17 | 36.72118 | 0.70594151 | 0.1987 | 3.5529 | 0.00038 | 0.0093 |
| ENSG00000261402.1 | 36.00217 | -0.725223919 | 0.2041 | -3.553 | 0.00038 | 0.0093 |
| ENSG00000126603.8 | 276.8082 | 0.430546061 | 0.1212 | 3.5512 | 0.00038 | 0.0093 |
| ENSG00000164794.8 | 2051.749 | -0.481995258 | 0.1358 | -3.549 | 0.00039 | 0.0094 |
| ENSG00000109917.10 | 2061.587 | 0.503593063 | 0.142 | 3.5453 | 0.00039 | 0.0094 |
| ENSG00000112183.14 | 427.0623 | -0.295200559 | 0.0833 | -3.546 | 0.00039 | 0.0094 |
| ENSG00000231064.6 | 179.722 | 0.506141442 | 0.1427 | 3.5465 | 0.00039 | 0.0094 |
| ENSG00000153902.13 | 1669.481 | 0.326622043 | 0.0922 | 3.5408 | 0.0004 | 0.0096 |
| ENSG00000163406.10 | 940.068 | 0.575430953 | 0.1625 | 3.5404 | 0.0004 | 0.0096 |
| ENSG00000259291.2 | 1800.067 | 0.309447431 | 0.0874 | 3.5405 | 0.0004 | 0.0096 |
| ENSG00000198804.2 | 918059.9 | 0.515013295 | 0.1455 | 3.5398 | 0.0004 | 0.0096 |
| ENSG00000141736.13 | 398.8014 | 0.535299402 | 0.1513 | 3.5373 | 0.0004 | 0.0096 |
| ENSG00000134042.12 | 1277.187 | 0.60457224 | 0.171 | 3.5362 | 0.00041 | 0.0096 |
| ENSG00000278962.1 | 83.48361 | -0.651837156 | 0.1844 | -3.535 | 0.00041 | 0.0097 |
| ENSG00000104369.4 | 770.0397 | -0.317658931 | 0.0899 | -3.534 | 0.00041 | 0.0097 |
| ENSG00000261829.1 | 50.49612 | 0.48587618 | 0.1375 | 3.5324 | 0.00041 | 0.0097 |
| ENSG00000125398.5 | 1523.246 | 0.758175862 | 0.2149 | 3.5285 | 0.00042 | 0.0098 |
| ENSG00000238260.1 | 40.31723 | 0.479199051 | 0.1358 | 3.5284 | 0.00042 | 0.0098 |
| ENSG00000125820.5 | 142.8645 | 0.597050072 | 0.1695 | 3.5233 | 0.00043 | 0.01 |
| ENSG00000056736.9 | 437.8211 | 0.437757313 | 0.1245 | 3.5167 | 0.00044 | 0.0101 |
| ENSG00000141485.16 | 339.4684 | 0.717993935 | 0.2042 | 3.5154 | 0.00044 | 0.0101 |
| ENSG00000147576.15 | 435.6637 | 0.506190097 | 0.1439 | 3.5174 | 0.00044 | 0.0101 |
| ENSG00000150054.18 | 452.1762 | -0.414487054 | 0.1179 | -3.517 | 0.00044 | 0.0101 |
| ENSG00000152661.7 | 6850.649 | 0.850352892 | 0.2419 | 3.5153 | 0.00044 | 0.0101 |
| ENSG00000180340.6 | 22.23337 | 0.761934971 | 0.2168 | 3.5148 | 0.00044 | 0.0101 |
| ENSG00000225174.1 | 35.50946 | 0.762370025 | 0.2167 | 3.518 | 0.00043 | 0.0101 |
| ENSG00000276509.1 | 31.16459 | 0.58040458 | 0.1651 | 3.5155 | 0.00044 | 0.0101 |
| ENSG00000149489.8 | 220.6648 | 0.477455031 | 0.1359 | 3.5141 | 0.00044 | 0.0101 |
| ENSG00000243989.8 | 138.6106 | 0.378037446 | 0.1076 | 3.5135 | 0.00044 | 0.0101 |
| ENSG00000169302.14 | 208.5537 | 0.338321661 | 0.0964 | 3.511 | 0.00045 | 0.0102 |
| ENSG00000156097.12 | 395.1335 | -0.479835599 | 0.1367 | -3.509 | 0.00045 | 0.0103 |
| ENSG00000156140.9 | 281.1584 | -0.398462194 | 0.1136 | -3.507 | 0.00045 | 0.0103 |
| ENSG00000157150.4 | 103.1805 | 0.423757069 | 0.1208 | 3.5074 | 0.00045 | 0.0103 |
| ENSG00000117650.12 | 76.28448 | -0.571060993 | 0.1629 | -3.505 | 0.00046 | 0.0104 |
| ENSG00000115255.10 | 262.0289 | 0.343887748 | 0.0981 | 3.504 | 0.00046 | 0.0104 |
| ENSG00000158748.3 | 70.99315 | -0.531983493 | 0.1519 | -3.503 | 0.00046 | 0.0104 |
| ENSG00000161955.16 | 430.8287 | 0.631261836 | 0.1803 | 3.5014 | 0.00046 | 0.0104 |
| ENSG00000079215.13 | 13631.38 | 0.823374261 | 0.2352 | 3.5004 | 0.00046 | 0.0104 |
| ENSG00000131941.7 | 229.932 | 0.580826851 | 0.166 | 3.4986 | 0.00047 | 0.0105 |
| ENSG00000175130.6 | 3623.195 | 0.358967552 | 0.1026 | 3.4986 | 0.00047 | 0.0105 |
| ENSG00000072310.16 | 1553.233 | 0.4899166 | 0.1401 | 3.498 | 0.00047 | 0.0105 |
| ENSG00000068305.17 | 3306.549 | -0.332890342 | 0.0952 | -3.496 | 0.00047 | 0.0105 |
| ENSG00000133433.10 | 297.6652 | -0.679755824 | 0.1944 | -3.496 | 0.00047 | 0.0105 |
| ENSG00000164976.8 | 2055.755 | 0.614518558 | 0.1757 | 3.497 | 0.00047 | 0.0105 |
| ENSG00000241839.9 | 665.4163 | 0.708607736 | 0.2027 | 3.4964 | 0.00047 | 0.0105 |
| ENSG00000090776.5 | 196.9365 | 0.348332769 | 0.0997 | 3.4941 | 0.00048 | 0.0105 |
| ENSG00000114790.12 | 791.5309 | 0.588255609 | 0.1686 | 3.4896 | 0.00048 | 0.0106 |
| ENSG00000168016.13 | 1519.51 | -0.273851897 | 0.0785 | -3.49 | 0.00048 | 0.0106 |
| ENSG00000169860.6 | 299.9168 | 0.708550246 | 0.203 | 3.4906 | 0.00048 | 0.0106 |
| ENSG00000184232.8 | 657.7468 | 0.661321787 | 0.1895 | 3.4898 | 0.00048 | 0.0106 |
| ENSG00000196139.13 | 144.3517 | 0.52026633 | 0.1494 | 3.4835 | 0.00049 | 0.0108 |
| ENSG00000162836.11 | 275.7251 | 0.416303809 | 0.1196 | 3.4807 | 0.0005 | 0.0109 |
| ENSG00000173511.9 | 1543.358 | 0.322208031 | 0.0926 | 3.4796 | 0.0005 | 0.0109 |
| ENSG00000168394.10 | 300.7263 | 0.460908459 | 0.1325 | 3.4786 | 0.0005 | 0.0109 |
| ENSG00000124145.6 | 932.3735 | 0.940391012 | 0.2705 | 3.4769 | 0.00051 | 0.011 |
| ENSG00000149499.11 | 448.7009 | 0.440628227 | 0.1267 | 3.4771 | 0.00051 | 0.011 |
| ENSG00000142611.16 | 352.0445 | 0.578394327 | 0.1664 | 3.4758 | 0.00051 | 0.011 |
| ENSG00000163380.15 | 45.97178 | 0.501811209 | 0.1444 | 3.4758 | 0.00051 | 0.011 |
| ENSG00000136383.6 | 220.519 | -0.407607458 | 0.1173 | -3.474 | 0.00051 | 0.011 |
| ENSG00000135540.11 | 465.2531 | 0.636690313 | 0.1834 | 3.4716 | 0.00052 | 0.0111 |
| ENSG00000113749.7 | 839.2904 | -0.30669795 | 0.0884 | -3.47 | 0.00052 | 0.0112 |
| ENSG00000204136.10 | 233.5136 | 0.333285408 | 0.0961 | 3.469 | 0.00052 | 0.0112 |
| ENSG00000164049.14 | 84.94397 | 0.380988088 | 0.1099 | 3.4672 | 0.00053 | 0.0112 |
| ENSG00000178602.7 | 11.83954 | 0.935318439 | 0.2698 | 3.4662 | 0.00053 | 0.0112 |
| ENSG00000178796.12 | 84.48379 | -0.372256691 | 0.1074 | -3.466 | 0.00053 | 0.0112 |
| ENSG00000198888.2 | 325582.8 | 0.436063733 | 0.1258 | 3.4669 | 0.00053 | 0.0112 |
| ENSG00000166148.3 | 43.29561 | -0.748151118 | 0.216 | -3.464 | 0.00053 | 0.0112 |
| ENSG00000129151.8 | 241.9193 | 0.695004429 | 0.2007 | 3.4635 | 0.00053 | 0.0112 |
| ENSG00000257829.1 | 22.28032 | -0.581683271 | 0.168 | -3.463 | 0.00053 | 0.0112 |
| ENSG00000156966.6 | 199.9362 | 0.511391147 | 0.1478 | 3.4602 | 0.00054 | 0.0113 |
| ENSG00000248394.1 | 12.01657 | 0.597105765 | 0.1726 | 3.4592 | 0.00054 | 0.0114 |
| ENSG00000087903.12 | 222.3047 | 0.47332608 | 0.137 | 3.4552 | 0.00055 | 0.0115 |
| ENSG00000220575.7 | 349.9783 | -0.400568128 | 0.1159 | -3.455 | 0.00055 | 0.0115 |
| ENSG00000260641.1 | 165.5298 | -0.344526771 | 0.0997 | -3.456 | 0.00055 | 0.0115 |
| ENSG00000140470.13 | 101.7125 | 0.462917665 | 0.1341 | 3.4526 | 0.00056 | 0.0115 |
| ENSG00000236197.3 | 116.844 | 0.50204936 | 0.1454 | 3.4531 | 0.00055 | 0.0115 |
| ENSG00000188394.6 | 45.23381 | -0.414422172 | 0.1201 | -3.452 | 0.00056 | 0.0115 |
| ENSG00000160602.13 | 59.95308 | 0.344567342 | 0.0998 | 3.4514 | 0.00056 | 0.0115 |
| ENSG00000283538.1 | 39.54512 | -0.474431364 | 0.1375 | -3.45 | 0.00056 | 0.0116 |
| ENSG00000227195.9 | 1721.395 | 0.476059971 | 0.1381 | 3.4484 | 0.00056 | 0.0116 |
| ENSG00000144649.8 | 118.3932 | 0.494417632 | 0.1434 | 3.4475 | 0.00057 | 0.0116 |
| ENSG00000144730.16 | 531.2024 | 0.295807217 | 0.0859 | 3.4456 | 0.00057 | 0.0116 |
| ENSG00000162599.15 | 2298.176 | 0.43022748 | 0.1248 | 3.4469 | 0.00057 | 0.0116 |
| ENSG00000219607.3 | 150.6593 | 0.638188752 | 0.1852 | 3.4458 | 0.00057 | 0.0116 |
| ENSG00000264049.1 | 24.01484 | 0.437296052 | 0.1269 | 3.4453 | 0.00057 | 0.0116 |
| ENSG00000177303.9 | 860.2189 | 0.42849758 | 0.1244 | 3.4435 | 0.00057 | 0.0117 |
| ENSG00000132329.10 | 786.1248 | 0.439248165 | 0.1276 | 3.4417 | 0.00058 | 0.0117 |
| ENSG00000188338.14 | 360.941 | 0.488486218 | 0.1419 | 3.4421 | 0.00058 | 0.0117 |
| ENSG00000250602.5 | 34.01684 | 0.629337301 | 0.1828 | 3.4428 | 0.00058 | 0.0117 |
| ENSG00000101846.6 | 1138.596 | -0.394808615 | 0.1149 | -3.437 | 0.00059 | 0.0119 |
| ENSG00000121281.12 | 304.9349 | -0.405046144 | 0.1179 | -3.437 | 0.00059 | 0.0119 |
| ENSG00000214954.8 | 213.912 | 1.128905008 | 0.3286 | 3.4358 | 0.00059 | 0.0119 |
| ENSG00000130787.13 | 2891.168 | 0.274845449 | 0.0801 | 3.4313 | 0.0006 | 0.012 |
| ENSG00000165175.15 | 1168.234 | 0.377485866 | 0.11 | 3.4324 | 0.0006 | 0.012 |
| ENSG00000178235.7 | 1859.243 | -0.405728934 | 0.1183 | -3.43 | 0.0006 | 0.012 |
| ENSG00000205213.13 | 633.9643 | 0.407331379 | 0.1187 | 3.431 | 0.0006 | 0.012 |
| ENSG00000225630.1 | 2626.084 | 1.916272249 | 0.5584 | 3.4317 | 0.0006 | 0.012 |
| ENSG00000253305.2 | 171.1901 | 0.575025349 | 0.1676 | 3.4305 | 0.0006 | 0.012 |
| ENSG00000253762.1 | 17.28613 | -0.584115324 | 0.1702 | -3.432 | 0.0006 | 0.012 |
| ENSG00000145632.14 | 2920.783 | -0.295488501 | 0.0864 | -3.422 | 0.00062 | 0.0123 |
| ENSG00000149503.12 | 517.9161 | -0.361705325 | 0.1057 | -3.422 | 0.00062 | 0.0123 |
| ENSG00000054179.11 | 275.4627 | 0.671268286 | 0.1962 | 3.4205 | 0.00062 | 0.0123 |
| ENSG00000100439.10 | 300.6539 | 0.449687228 | 0.1315 | 3.4195 | 0.00063 | 0.0123 |
| ENSG00000155324.9 | 1094.314 | 0.596748973 | 0.1745 | 3.4193 | 0.00063 | 0.0123 |
| ENSG00000144230.16 | 404.346 | 0.648402422 | 0.1899 | 3.4152 | 0.00064 | 0.0125 |
| ENSG00000132825.6 | 189.0581 | 0.432957634 | 0.1268 | 3.4141 | 0.00064 | 0.0125 |
| ENSG00000173905.8 | 1458.035 | 0.445879815 | 0.1306 | 3.4142 | 0.00064 | 0.0125 |
| ENSG00000106278.11 | 5952.247 | 0.575628283 | 0.1687 | 3.4129 | 0.00064 | 0.0125 |
| ENSG00000184608.8 | 10.51318 | 0.792761574 | 0.2323 | 3.4134 | 0.00064 | 0.0125 |
| ENSG00000128641.18 | 732.7911 | -0.31408473 | 0.0921 | -3.412 | 0.00064 | 0.0125 |
| ENSG00000265179.6 | 109.936 | -0.440463186 | 0.1292 | -3.41 | 0.00065 | 0.0126 |
| ENSG00000145832.12 | 199.1432 | 0.531461843 | 0.1561 | 3.4056 | 0.00066 | 0.0128 |
| ENSG00000177283.6 | 300.1315 | 0.527405924 | 0.1549 | 3.4041 | 0.00066 | 0.0128 |
| ENSG00000188763.4 | 104.4552 | 0.418017828 | 0.1228 | 3.4043 | 0.00066 | 0.0128 |
| ENSG00000135916.15 | 15080.71 | 0.312870618 | 0.0919 | 3.4031 | 0.00067 | 0.0128 |
| ENSG00000218227.3 | 86.39331 | 0.51478958 | 0.1514 | 3.4009 | 0.00067 | 0.0129 |
| ENSG00000148925.10 | 1798.617 | -0.351879847 | 0.1035 | -3.399 | 0.00068 | 0.0129 |
| ENSG00000261386.2 | 19.68503 | 0.492671701 | 0.1449 | 3.3994 | 0.00068 | 0.0129 |
| ENSG00000136237.18 | 3663.294 | -0.388799648 | 0.1145 | -3.397 | 0.00068 | 0.013 |
| ENSG00000120885.20 | 48646.56 | 0.39967189 | 0.1178 | 3.3939 | 0.00069 | 0.0131 |
| ENSG00000128203.6 | 780.112 | -0.382670769 | 0.1128 | -3.394 | 0.00069 | 0.0131 |
| ENSG00000135312.6 | 81.22858 | -0.512116062 | 0.1509 | -3.394 | 0.00069 | 0.0131 |
| ENSG00000174136.11 | 1993.795 | -0.347013111 | 0.1023 | -3.392 | 0.00069 | 0.0132 |
| ENSG00000112972.14 | 4608.79 | -0.354488761 | 0.1045 | -3.391 | 0.0007 | 0.0132 |
| ENSG00000180613.10 | 8.932334 | 0.824539855 | 0.2433 | 3.389 | 0.0007 | 0.0132 |
| ENSG00000257434.1 | 683.9654 | -0.613439962 | 0.1811 | -3.387 | 0.00071 | 0.0133 |
| ENSG00000143149.12 | 1413.996 | 0.360552799 | 0.1065 | 3.3859 | 0.00071 | 0.0133 |
| ENSG00000077585.13 | 649.4413 | 0.40844735 | 0.1207 | 3.3842 | 0.00071 | 0.0134 |
| ENSG00000182154.7 | 2911.111 | 0.313437656 | 0.0926 | 3.3846 | 0.00071 | 0.0134 |
| ENSG00000106538.9 | 236.313 | 0.427374961 | 0.1263 | 3.3835 | 0.00072 | 0.0134 |
| ENSG00000169856.8 | 10.27343 | 0.641010734 | 0.1895 | 3.3826 | 0.00072 | 0.0134 |
| ENSG00000126016.14 | 1805.144 | 0.457995207 | 0.1355 | 3.3811 | 0.00072 | 0.0135 |
| ENSG00000067177.14 | 299.1036 | 0.583609247 | 0.1728 | 3.3771 | 0.00073 | 0.0136 |
| ENSG00000106565.17 | 210.651 | 0.654778108 | 0.194 | 3.3743 | 0.00074 | 0.0137 |
| ENSG00000213390.10 | 192.5034 | 0.364848895 | 0.1083 | 3.369 | 0.00075 | 0.014 |
| ENSG00000104870.12 | 408.8568 | 0.468222295 | 0.1391 | 3.3661 | 0.00076 | 0.0141 |
| ENSG00000144579.7 | 552.3538 | 0.41610238 | 0.1236 | 3.3669 | 0.00076 | 0.0141 |
| ENSG00000224058.2 | 109.3861 | 0.350101349 | 0.104 | 3.3665 | 0.00076 | 0.0141 |
| ENSG00000162621.6 | 78.76877 | -0.616398993 | 0.1832 | -3.364 | 0.00077 | 0.0141 |
| ENSG00000236953.1 | 59.97226 | 1.096655002 | 0.326 | 3.3644 | 0.00077 | 0.0141 |
| ENSG00000272953.1 | 21.76376 | 0.512653318 | 0.1524 | 3.3634 | 0.00077 | 0.0141 |
| ENSG00000167799.9 | 45.42938 | 0.407399564 | 0.1212 | 3.3627 | 0.00077 | 0.0141 |
| ENSG00000059804.15 | 2556.235 | -0.319958477 | 0.0952 | -3.362 | 0.00077 | 0.0141 |
| ENSG00000180543.4 | 1094.988 | -0.326530198 | 0.0971 | -3.361 | 0.00078 | 0.0142 |
| ENSG00000103335.20 | 692.6909 | 0.414429305 | 0.1234 | 3.3597 | 0.00078 | 0.0142 |
| ENSG00000166257.8 | 12702.98 | -0.387955893 | 0.1155 | -3.36 | 0.00078 | 0.0142 |
| ENSG00000259984.1 | 11.80163 | -0.61262572 | 0.1824 | -3.359 | 0.00078 | 0.0142 |
| ENSG00000162482.4 | 40.92003 | 0.429487905 | 0.1279 | 3.3582 | 0.00078 | 0.0142 |
| ENSG00000079482.12 | 1750.697 | 0.289079747 | 0.0862 | 3.355 | 0.00079 | 0.0143 |
| ENSG00000230651.7 | 142.8785 | 0.337946078 | 0.1007 | 3.3555 | 0.00079 | 0.0143 |
| ENSG00000112146.16 | 2181.327 | -0.365327758 | 0.1089 | -3.354 | 0.0008 | 0.0144 |
| ENSG00000179604.9 | 1578.771 | 0.612035961 | 0.1826 | 3.3526 | 0.0008 | 0.0144 |
| ENSG00000172819.16 | 161.0418 | 0.378535448 | 0.113 | 3.3497 | 0.00081 | 0.0145 |
| ENSG00000272990.1 | 85.06588 | 0.399701173 | 0.1194 | 3.3489 | 0.00081 | 0.0145 |
| ENSG00000164188.8 | 802.7233 | 0.729068456 | 0.2178 | 3.3481 | 0.00081 | 0.0146 |
| ENSG00000065320.8 | 246.4283 | 0.481099448 | 0.1437 | 3.3474 | 0.00082 | 0.0146 |
| ENSG00000101871.14 | 250.0777 | 0.426571253 | 0.1276 | 3.3435 | 0.00083 | 0.0146 |
| ENSG00000102349.16 | 367.2042 | -0.299567571 | 0.0896 | -3.345 | 0.00082 | 0.0146 |
| ENSG00000120896.13 | 962.5599 | 0.450564514 | 0.1348 | 3.3427 | 0.00083 | 0.0146 |
| ENSG00000125520.13 | 475.6485 | 0.402329212 | 0.1203 | 3.3432 | 0.00083 | 0.0146 |
| ENSG00000145721.11 | 651.4216 | 0.455308092 | 0.1362 | 3.3439 | 0.00083 | 0.0146 |
| ENSG00000172201.11 | 1299.698 | 0.614682141 | 0.1837 | 3.3461 | 0.00082 | 0.0146 |
| ENSG00000204219.9 | 128.621 | 0.388053268 | 0.1161 | 3.3437 | 0.00083 | 0.0146 |
| ENSG00000253642.5 | 44.51611 | 0.365095181 | 0.1092 | 3.3424 | 0.00083 | 0.0146 |
| ENSG00000259583.2 | 334.5829 | -0.422489927 | 0.1264 | -3.343 | 0.00083 | 0.0146 |
| ENSG00000267714.1 | 15.58187 | 0.728583228 | 0.2178 | 3.3452 | 0.00082 | 0.0146 |
| ENSG00000185920.15 | 871.7207 | 0.366327807 | 0.1097 | 3.3391 | 0.00084 | 0.0147 |
| ENSG00000134250.18 | 1322.904 | 0.681038483 | 0.204 | 3.3381 | 0.00084 | 0.0147 |
| ENSG00000279302.2 | 49.36598 | 0.458628947 | 0.1374 | 3.3382 | 0.00084 | 0.0147 |
| ENSG00000143842.14 | 406.7884 | 0.382760149 | 0.1147 | 3.3373 | 0.00085 | 0.0147 |
| ENSG00000181284.2 | 17.71969 | 0.587120946 | 0.176 | 3.3368 | 0.00085 | 0.0147 |
| ENSG00000279875.1 | 20.24228 | -0.550898422 | 0.1651 | -3.337 | 0.00085 | 0.0147 |
| ENSG00000213977.7 | 336.8634 | 0.340407112 | 0.1021 | 3.335 | 0.00085 | 0.0148 |
| ENSG00000225472.1 | 54.64098 | 0.586551554 | 0.1761 | 3.3313 | 0.00086 | 0.0149 |
| ENSG00000242252.1 | 19.68025 | 0.620911435 | 0.1864 | 3.3314 | 0.00086 | 0.0149 |
| ENSG00000152785.6 | 77.12798 | -0.473752951 | 0.1424 | -3.328 | 0.00088 | 0.0151 |
| ENSG00000083454.21 | 435.2298 | -0.342755762 | 0.1031 | -3.324 | 0.00089 | 0.0152 |
| ENSG00000104313.17 | 145.2743 | 0.491633358 | 0.1479 | 3.3238 | 0.00089 | 0.0152 |
| ENSG00000136457.9 | 132.5045 | 0.518544709 | 0.156 | 3.325 | 0.00088 | 0.0152 |
| ENSG00000165617.14 | 323.2374 | -0.444598585 | 0.1338 | -3.324 | 0.00089 | 0.0152 |
| ENSG00000165929.12 | 267.2982 | -0.419536864 | 0.1262 | -3.324 | 0.00089 | 0.0152 |
| ENSG00000175938.6 | 136.9773 | 0.412863765 | 0.1242 | 3.325 | 0.00088 | 0.0152 |
| ENSG00000131721.5 | 12.43737 | 0.684837201 | 0.2062 | 3.3215 | 0.0009 | 0.0152 |
| ENSG00000144152.12 | 262.9949 | -0.544690358 | 0.164 | -3.321 | 0.0009 | 0.0152 |
| ENSG00000180815.14 | 33.88426 | -0.471551368 | 0.1419 | -3.322 | 0.00089 | 0.0152 |
| ENSG00000205336.11 | 3556.061 | 0.552949713 | 0.1664 | 3.3226 | 0.00089 | 0.0152 |
| ENSG00000051108.14 | 1257.271 | -0.304385248 | 0.0917 | -3.32 | 0.0009 | 0.0153 |
| ENSG00000120915.13 | 251.1957 | 0.41538142 | 0.1252 | 3.3171 | 0.00091 | 0.0154 |
| ENSG00000141404.15 | 2458.639 | -0.325907009 | 0.0983 | -3.317 | 0.00091 | 0.0154 |
| ENSG00000185187.12 | 219.0529 | 0.425776864 | 0.1284 | 3.3162 | 0.00091 | 0.0154 |
| ENSG00000198945.7 | 268.2231 | -0.228760245 | 0.069 | -3.313 | 0.00092 | 0.0155 |
| ENSG00000007372.20 | 692.8418 | 0.496094616 | 0.1498 | 3.3125 | 0.00092 | 0.0155 |
| ENSG00000174145.7 | 976.6838 | -0.437503611 | 0.1321 | -3.312 | 0.00093 | 0.0155 |
| ENSG00000106648.13 | 61.07629 | -0.454605099 | 0.1373 | -3.31 | 0.00093 | 0.0156 |
| ENSG00000164117.13 | 230.422 | 0.296541263 | 0.0896 | 3.3106 | 0.00093 | 0.0156 |
| ENSG00000123096.11 | 518.0691 | 0.561006958 | 0.1696 | 3.3086 | 0.00094 | 0.0156 |
| ENSG00000129128.12 | 2424.378 | -0.348418407 | 0.1053 | -3.309 | 0.00094 | 0.0156 |
| ENSG00000270181.1 | 150.4932 | -0.69450189 | 0.2099 | -3.309 | 0.00094 | 0.0156 |
| ENSG00000171444.17 | 724.7128 | 0.48473148 | 0.1466 | 3.3061 | 0.00095 | 0.0157 |
| ENSG00000085978.21 | 894.8742 | -0.296279582 | 0.0896 | -3.305 | 0.00095 | 0.0157 |
| ENSG00000136869.13 | 422.2386 | 0.595860179 | 0.1803 | 3.3049 | 0.00095 | 0.0157 |
| ENSG00000235092.5 | 52.25543 | 0.443206255 | 0.1341 | 3.3043 | 0.00095 | 0.0157 |
| ENSG00000270118.1 | 28.24746 | 0.848326334 | 0.2567 | 3.3041 | 0.00095 | 0.0157 |
| ENSG00000232987.1 | 10.5158 | 0.663922409 | 0.2011 | 3.3016 | 0.00096 | 0.0159 |
| ENSG00000279569.1 | 177.8871 | 0.567775515 | 0.1721 | 3.3 | 0.00097 | 0.0159 |
| ENSG00000087250.8 | 5754.354 | 0.383275663 | 0.1162 | 3.2976 | 0.00098 | 0.016 |
| ENSG00000134548.9 | 389.0764 | 0.495556013 | 0.1503 | 3.2977 | 0.00097 | 0.016 |
| ENSG00000149269.9 | 8612.607 | -0.394196778 | 0.1195 | -3.297 | 0.00098 | 0.016 |
| ENSG00000156675.15 | 136.4387 | -0.329957832 | 0.1001 | -3.297 | 0.00098 | 0.016 |
| ENSG00000173166.17 | 1653.529 | -0.310732057 | 0.0943 | -3.296 | 0.00098 | 0.016 |
| ENSG00000148204.11 | 521.2118 | 0.523298346 | 0.1588 | 3.2954 | 0.00098 | 0.016 |
| ENSG00000106025.8 | 141.8458 | 0.456021691 | 0.1384 | 3.2945 | 0.00099 | 0.016 |
| ENSG00000137727.12 | 961.7339 | -0.49930605 | 0.1516 | -3.293 | 0.00099 | 0.0161 |
| ENSG00000165912.15 | 230.0003 | 0.336026068 | 0.102 | 3.2936 | 0.00099 | 0.0161 |
| ENSG00000110675.12 | 2904.944 | -0.413317821 | 0.1256 | -3.291 | 0.001 | 0.0161 |
| ENSG00000275395.4 | 152.9578 | -0.805713367 | 0.2448 | -3.291 | 0.001 | 0.0161 |
| ENSG00000163576.17 | 58.88987 | 0.643330989 | 0.1955 | 3.2902 | 0.001 | 0.0162 |
| ENSG00000073969.18 | 12377.42 | -0.473065618 | 0.144 | -3.286 | 0.00102 | 0.0164 |
| ENSG00000148482.11 | 659.056 | 0.819634372 | 0.2495 | 3.2855 | 0.00102 | 0.0164 |
| ENSG00000159459.11 | 1209.133 | -0.224959289 | 0.0685 | -3.284 | 0.00102 | 0.0164 |
| ENSG00000143772.9 | 2529.253 | 0.638430515 | 0.1944 | 3.2836 | 0.00102 | 0.0164 |
| ENSG00000164100.8 | 659.1327 | -0.337778746 | 0.1029 | -3.284 | 0.00102 | 0.0164 |
| ENSG00000198300.12 | 10511.97 | -0.376862436 | 0.1148 | -3.283 | 0.00103 | 0.0165 |
| ENSG00000261604.1 | 171.8729 | -0.387098441 | 0.1179 | -3.282 | 0.00103 | 0.0165 |
| ENSG00000118242.15 | 274.6443 | -0.388380946 | 0.1184 | -3.28 | 0.00104 | 0.0165 |
| ENSG00000169515.5 | 102.5561 | 0.563640308 | 0.1718 | 3.2805 | 0.00104 | 0.0165 |
| ENSG00000188064.9 | 415.1808 | 0.567068403 | 0.1729 | 3.2802 | 0.00104 | 0.0165 |
| ENSG00000128849.10 | 909.2992 | 0.39958166 | 0.1218 | 3.2795 | 0.00104 | 0.0165 |
| ENSG00000102385.12 | 1560.81 | -0.331981756 | 0.1013 | -3.278 | 0.00105 | 0.0166 |
| ENSG00000160808.9 | 33.4386 | 0.531572889 | 0.1622 | 3.2779 | 0.00105 | 0.0166 |
| ENSG00000186594.13 | 283.8062 | -0.343476191 | 0.1049 | -3.275 | 0.00105 | 0.0167 |
| ENSG00000163947.11 | 1855.136 | -0.366135303 | 0.1118 | -3.274 | 0.00106 | 0.0167 |
| ENSG00000170571.11 | 184.3864 | -0.487957755 | 0.1491 | -3.272 | 0.00107 | 0.0168 |
| ENSG00000198721.12 | 525.883 | 0.29100636 | 0.0889 | 3.2716 | 0.00107 | 0.0168 |
| ENSG00000259895.1 | 520.2575 | 1.123641519 | 0.3435 | 3.2715 | 0.00107 | 0.0168 |
| ENSG00000243056.1 | 80.69863 | 0.3559821 | 0.1089 | 3.2699 | 0.00108 | 0.0169 |
| ENSG00000077782.19 | 1572.825 | 0.25862261 | 0.0791 | 3.2694 | 0.00108 | 0.0169 |
| ENSG00000151892.14 | 669.7291 | 0.3607935 | 0.1104 | 3.2681 | 0.00108 | 0.0169 |
| ENSG00000143603.18 | 1266.154 | 0.450145948 | 0.1378 | 3.266 | 0.00109 | 0.017 |
| ENSG00000154217.14 | 1780.316 | 0.330173409 | 0.1011 | 3.2673 | 0.00109 | 0.017 |
| ENSG00000176971.3 | 244.8153 | 0.518360372 | 0.1587 | 3.2663 | 0.00109 | 0.017 |
| ENSG00000203952.9 | 32.13866 | 0.40713529 | 0.1246 | 3.2664 | 0.00109 | 0.017 |
| ENSG00000131398.13 | 1423.712 | -0.314638729 | 0.0964 | -3.264 | 0.0011 | 0.0171 |
| ENSG00000136824.18 | 412.4169 | -0.315532382 | 0.0967 | -3.263 | 0.0011 | 0.0171 |
| ENSG00000164776.9 | 162.2127 | 0.510990238 | 0.1566 | 3.2626 | 0.0011 | 0.0171 |
| ENSG00000101849.15 | 639.5579 | 0.307041318 | 0.0942 | 3.2601 | 0.00111 | 0.0172 |
| ENSG00000218109.5 | 11.29593 | -0.76343552 | 0.2342 | -3.26 | 0.00111 | 0.0172 |
| ENSG00000228203.6 | 361.1056 | -0.314626067 | 0.0965 | -3.26 | 0.00111 | 0.0172 |
| ENSG00000259891.1 | 198.8354 | 0.305742943 | 0.0938 | 3.2596 | 0.00112 | 0.0172 |
| ENSG00000260645.1 | 38.56871 | 0.37614762 | 0.1155 | 3.2575 | 0.00112 | 0.0173 |
| ENSG00000215612.7 | 30.72959 | 0.607930977 | 0.1867 | 3.2569 | 0.00113 | 0.0173 |
| ENSG00000055070.16 | 798.6625 | 0.32304194 | 0.0992 | 3.2557 | 0.00113 | 0.0173 |
| ENSG00000151012.13 | 1785.418 | 0.618186401 | 0.1901 | 3.2526 | 0.00114 | 0.0175 |
| ENSG00000138623.9 | 745.1443 | -0.381807382 | 0.1175 | -3.251 | 0.00115 | 0.0176 |
| ENSG00000167880.7 | 33.09898 | -0.660487541 | 0.2033 | -3.248 | 0.00116 | 0.0177 |
| ENSG00000254910.1 | 12.11719 | 0.831050465 | 0.2558 | 3.2485 | 0.00116 | 0.0177 |
| ENSG00000135097.6 | 289.4694 | 0.380855195 | 0.1173 | 3.2474 | 0.00116 | 0.0177 |
| ENSG00000166349.9 | 15.74094 | -0.531159826 | 0.1637 | -3.246 | 0.00117 | 0.0178 |
| ENSG00000004776.11 | 397.8296 | 0.532938585 | 0.1643 | 3.2431 | 0.00118 | 0.0179 |
| ENSG00000134343.12 | 1570.336 | -0.360846303 | 0.1113 | -3.243 | 0.00118 | 0.0179 |
| ENSG00000166763.7 | 117.3471 | -0.394388557 | 0.1216 | -3.244 | 0.00118 | 0.0179 |
| ENSG00000100095.18 | 5979.443 | -0.306067994 | 0.0944 | -3.241 | 0.00119 | 0.0179 |
| ENSG00000102760.12 | 666.5825 | 0.386522668 | 0.1192 | 3.2424 | 0.00119 | 0.0179 |
| ENSG00000168306.12 | 69.98073 | 0.588580048 | 0.1816 | 3.2419 | 0.00119 | 0.0179 |
| ENSG00000275620.1 | 756.1592 | 0.614970829 | 0.1897 | 3.2413 | 0.00119 | 0.0179 |
| ENSG00000080493.14 | 3044.362 | 0.73794046 | 0.2278 | 3.2394 | 0.0012 | 0.0179 |
| ENSG00000126215.13 | 207.6693 | 0.261487176 | 0.0807 | 3.2397 | 0.0012 | 0.0179 |
| ENSG00000170921.14 | 3614.883 | -0.297650135 | 0.0919 | -3.238 | 0.0012 | 0.018 |
| ENSG00000184486.9 | 607.9465 | 0.280606461 | 0.0867 | 3.2365 | 0.00121 | 0.0181 |
| ENSG00000100243.20 | 2659.224 | 0.310632697 | 0.096 | 3.2344 | 0.00122 | 0.0181 |
| ENSG00000229847.8 | 1687.919 | 0.603900216 | 0.1867 | 3.2342 | 0.00122 | 0.0181 |
| ENSG00000236523.2 | 203.9503 | 1.198672785 | 0.3706 | 3.2348 | 0.00122 | 0.0181 |
| ENSG00000164708.5 | 47.2294 | 0.6488903 | 0.2007 | 3.2332 | 0.00122 | 0.0182 |
| ENSG00000204054.13 | 629.0159 | 0.294189499 | 0.0911 | 3.2306 | 0.00124 | 0.0183 |
| ENSG00000164039.14 | 447.2266 | 0.474839074 | 0.147 | 3.23 | 0.00124 | 0.0183 |
| ENSG00000116641.17 | 1298.564 | 0.255189514 | 0.0791 | 3.2277 | 0.00125 | 0.0184 |
| ENSG00000143545.8 | 385.6284 | 0.439018869 | 0.136 | 3.2273 | 0.00125 | 0.0184 |
| ENSG00000179820.15 | 1562.686 | -0.279285602 | 0.0865 | -3.227 | 0.00125 | 0.0184 |
| ENSG00000275763.3 | 12.63059 | 0.662941984 | 0.2054 | 3.227 | 0.00125 | 0.0184 |
| ENSG00000216285.5 | 11.46404 | -0.87028065 | 0.27 | -3.223 | 0.00127 | 0.0187 |
| ENSG00000174514.12 | 3783.377 | -0.35343098 | 0.1097 | -3.221 | 0.00128 | 0.0188 |
| ENSG00000172209.5 | 855.8508 | -0.341585834 | 0.1061 | -3.22 | 0.00128 | 0.0188 |
| ENSG00000121764.11 | 18.709 | -0.675953612 | 0.21 | -3.219 | 0.00129 | 0.0188 |
| ENSG00000271601.3 | 562.4643 | 0.307578185 | 0.0955 | 3.2193 | 0.00129 | 0.0188 |
| ENSG00000230042.1 | 24.56102 | 0.538666363 | 0.1674 | 3.218 | 0.00129 | 0.0188 |
| ENSG00000069696.6 | 58.98384 | 0.429455907 | 0.1335 | 3.2171 | 0.0013 | 0.0189 |
| ENSG00000123643.12 | 636.6743 | -0.467985173 | 0.1455 | -3.216 | 0.0013 | 0.0189 |
| ENSG00000182732.16 | 835.2163 | -0.337549251 | 0.1051 | -3.213 | 0.00131 | 0.0191 |
| ENSG00000108094.14 | 1101.949 | -0.248034557 | 0.0772 | -3.212 | 0.00132 | 0.0191 |
| ENSG00000128989.10 | 16801.79 | -0.318828122 | 0.0993 | -3.211 | 0.00132 | 0.0191 |
| ENSG00000129521.13 | 597.8335 | 0.476706097 | 0.1485 | 3.2105 | 0.00133 | 0.0192 |
| ENSG00000135426.15 | 3489.51 | -0.395372139 | 0.1232 | -3.208 | 0.00134 | 0.0193 |
| ENSG00000100889.11 | 123.5031 | 0.306506672 | 0.0956 | 3.2062 | 0.00134 | 0.0194 |
| ENSG00000091157.13 | 3234.206 | -0.330560186 | 0.1032 | -3.205 | 0.00135 | 0.0195 |
| ENSG00000072952.18 | 1491.412 | 0.657591892 | 0.2053 | 3.2026 | 0.00136 | 0.0195 |
| ENSG00000146950.12 | 829.45 | -0.271187421 | 0.0847 | -3.203 | 0.00136 | 0.0195 |
| ENSG00000151490.13 | 1211.454 | -0.389057106 | 0.1214 | -3.204 | 0.00135 | 0.0195 |
| ENSG00000255337.1 | 12.1649 | 0.627212759 | 0.1958 | 3.2034 | 0.00136 | 0.0195 |
| ENSG00000272449.1 | 94.00264 | 0.528642927 | 0.1651 | 3.2025 | 0.00136 | 0.0195 |
| ENSG00000278983.1 | 67.28963 | 0.498847321 | 0.1558 | 3.2019 | 0.00137 | 0.0195 |
| ENSG00000114547.9 | 56.86907 | 0.334931263 | 0.1046 | 3.2012 | 0.00137 | 0.0195 |
| ENSG00000123453.17 | 139.4362 | 0.414672171 | 0.1296 | 3.2 | 0.00137 | 0.0196 |
| ENSG00000162496.8 | 333.7295 | 0.376916326 | 0.1178 | 3.1992 | 0.00138 | 0.0196 |
| ENSG00000179841.8 | 2070.811 | -0.364381711 | 0.1139 | -3.198 | 0.00138 | 0.0196 |
| ENSG00000143514.16 | 1490.128 | 0.554264491 | 0.1734 | 3.197 | 0.00139 | 0.0197 |
| ENSG00000162391.11 | 32.78035 | 0.530815496 | 0.166 | 3.1969 | 0.00139 | 0.0197 |
| ENSG00000261195.1 | 16.47588 | 0.654278923 | 0.2047 | 3.1964 | 0.00139 | 0.0197 |
| ENSG00000099834.18 | 78.10796 | 0.372242086 | 0.1165 | 3.1947 | 0.0014 | 0.0198 |
| ENSG00000138311.15 | 3049.23 | -0.33025687 | 0.1034 | -3.195 | 0.0014 | 0.0198 |
| ENSG00000156535.13 | 158.473 | 0.397318067 | 0.1244 | 3.1939 | 0.0014 | 0.0198 |
| ENSG00000102996.4 | 275.4178 | 0.256472684 | 0.0804 | 3.1916 | 0.00142 | 0.0199 |
| ENSG00000114698.14 | 620.159 | 0.549557536 | 0.1723 | 3.1903 | 0.00142 | 0.0199 |
| ENSG00000144619.14 | 1120.341 | -0.333492863 | 0.1045 | -3.191 | 0.00142 | 0.0199 |
| ENSG00000234177.5 | 91.19249 | 0.36712093 | 0.1151 | 3.1904 | 0.00142 | 0.0199 |
| ENSG00000048991.16 | 5694.658 | -0.303794138 | 0.0952 | -3.189 | 0.00143 | 0.0199 |
| ENSG00000113296.14 | 543.6963 | 0.343256895 | 0.1076 | 3.1893 | 0.00143 | 0.0199 |
| ENSG00000153944.10 | 2401.101 | 0.42903319 | 0.1346 | 3.1883 | 0.00143 | 0.02 |
| ENSG00000198301.11 | 939.7991 | -0.371416768 | 0.1165 | -3.188 | 0.00143 | 0.02 |
| ENSG00000089199.9 | 4719.763 | -0.392568085 | 0.1232 | -3.187 | 0.00144 | 0.02 |
| ENSG00000162390.17 | 389.5163 | 0.322929352 | 0.1013 | 3.1864 | 0.00144 | 0.02 |
| ENSG00000224003.1 | 36.9414 | 0.459685825 | 0.1442 | 3.1871 | 0.00144 | 0.02 |
| ENSG00000256193.5 | 593.5002 | -0.49107621 | 0.1542 | -3.185 | 0.00145 | 0.02 |
| ENSG00000107249.21 | 237.7876 | 0.494736264 | 0.1554 | 3.1844 | 0.00145 | 0.0201 |
| ENSG00000119408.16 | 774.1783 | 0.393686973 | 0.1237 | 3.1836 | 0.00145 | 0.0201 |
| ENSG00000161642.17 | 1813.875 | 0.339095413 | 0.1065 | 3.1832 | 0.00146 | 0.0201 |
| ENSG00000164199.15 | 2954.916 | 0.60883477 | 0.1913 | 3.1833 | 0.00146 | 0.0201 |
| ENSG00000255443.1 | 102.819 | 0.339468741 | 0.1066 | 3.1844 | 0.00145 | 0.0201 |
| ENSG00000134824.13 | 2976.719 | 0.400853039 | 0.126 | 3.1825 | 0.00146 | 0.0201 |
| ENSG00000153208.16 | 493.649 | 0.562455456 | 0.1768 | 3.1805 | 0.00147 | 0.0201 |
| ENSG00000154928.16 | 494.3676 | 0.343329986 | 0.108 | 3.1795 | 0.00148 | 0.0201 |
| ENSG00000166816.13 | 446.6458 | 0.329030443 | 0.1035 | 3.1803 | 0.00147 | 0.0201 |
| ENSG00000186081.11 | 70.09056 | -0.64841967 | 0.2039 | -3.18 | 0.00147 | 0.0201 |
| ENSG00000227906.7 | 149.7486 | -0.293115758 | 0.0922 | -3.18 | 0.00147 | 0.0201 |
| ENSG00000256982.1 | 114.1551 | -0.369680742 | 0.1162 | -3.18 | 0.00147 | 0.0201 |
| ENSG00000043143.20 | 1998.592 | -0.329989261 | 0.1038 | -3.178 | 0.00148 | 0.0202 |
| ENSG00000058335.15 | 4584.098 | -0.396846333 | 0.1249 | -3.178 | 0.00148 | 0.0202 |
| ENSG00000076555.15 | 688.2989 | 0.467689871 | 0.1472 | 3.1776 | 0.00149 | 0.0202 |
| ENSG00000250166.2 | 18.19472 | 0.535980365 | 0.1687 | 3.1778 | 0.00148 | 0.0202 |
| ENSG00000184845.3 | 468.0227 | -0.39879997 | 0.1255 | -3.177 | 0.00149 | 0.0202 |
| ENSG00000068697.6 | 1330.974 | 0.26550068 | 0.0836 | 3.175 | 0.0015 | 0.0203 |
| ENSG00000075234.16 | 146.7405 | 0.462261082 | 0.1456 | 3.1744 | 0.0015 | 0.0203 |
| ENSG00000007384.15 | 210.2977 | 0.374357238 | 0.118 | 3.1733 | 0.00151 | 0.0203 |
| ENSG00000174437.16 | 15425.1 | -0.295785373 | 0.0932 | -3.174 | 0.00151 | 0.0203 |
| ENSG00000164715.5 | 3837.645 | -0.391562606 | 0.1235 | -3.171 | 0.00152 | 0.0203 |
| ENSG00000206384.10 | 36.33721 | -0.422380838 | 0.1332 | -3.172 | 0.00152 | 0.0203 |
| ENSG00000233273.1 | 8.861566 | -0.712405891 | 0.2246 | -3.172 | 0.00151 | 0.0203 |
| ENSG00000236754.5 | 48.84448 | 0.528111759 | 0.1665 | 3.1716 | 0.00152 | 0.0203 |
| ENSG00000005421.8 | 14.03385 | 0.571087548 | 0.1801 | 3.1709 | 0.00152 | 0.0204 |
| ENSG00000180914.10 | 193.0005 | 0.487897727 | 0.1539 | 3.1704 | 0.00152 | 0.0204 |
| ENSG00000140522.11 | 141.7856 | 0.568694092 | 0.1794 | 3.1692 | 0.00153 | 0.0204 |
| ENSG00000273483.1 | 19.37787 | 0.502761613 | 0.1588 | 3.1661 | 0.00154 | 0.0206 |
| ENSG00000162461.7 | 102.6948 | 0.278623261 | 0.088 | 3.1657 | 0.00155 | 0.0206 |
| ENSG00000250007.6 | 36.31531 | -0.385517594 | 0.1218 | -3.164 | 0.00156 | 0.0207 |
| ENSG00000256124.5 | 63.72311 | 0.430476212 | 0.1361 | 3.1637 | 0.00156 | 0.0207 |
| ENSG00000172345.13 | 82.77664 | -0.328039189 | 0.1037 | -3.163 | 0.00156 | 0.0208 |
| ENSG00000133863.6 | 70.69418 | 0.399439065 | 0.1264 | 3.1606 | 0.00157 | 0.0209 |
| ENSG00000112312.9 | 149.3795 | 0.355349517 | 0.1125 | 3.1592 | 0.00158 | 0.021 |
| ENSG00000156711.16 | 684.9918 | -0.281732382 | 0.0892 | -3.158 | 0.00159 | 0.021 |
| ENSG00000214659.4 | 40.71453 | -0.390801072 | 0.1237 | -3.158 | 0.00159 | 0.021 |
| ENSG00000075391.16 | 2270.475 | -0.260888657 | 0.0827 | -3.155 | 0.0016 | 0.021 |
| ENSG00000101203.16 | 556.413 | 0.450722251 | 0.1428 | 3.1561 | 0.0016 | 0.021 |
| ENSG00000129219.13 | 376.8147 | 0.417551819 | 0.1323 | 3.157 | 0.00159 | 0.021 |
| ENSG00000135454.13 | 2421.475 | -0.27796246 | 0.0881 | -3.156 | 0.0016 | 0.021 |
| ENSG00000189334.8 | 9.062269 | 0.749185889 | 0.2374 | 3.1554 | 0.0016 | 0.021 |
| ENSG00000241983.3 | 18.33978 | 0.523329955 | 0.1658 | 3.156 | 0.0016 | 0.021 |
| ENSG00000163673.7 | 145.9606 | -0.443265829 | 0.1405 | -3.154 | 0.00161 | 0.021 |
| ENSG00000276085.1 | 56.08571 | -0.782801701 | 0.2482 | -3.154 | 0.00161 | 0.021 |
| ENSG00000180525.11 | 127.4395 | 0.392330876 | 0.1244 | 3.1536 | 0.00161 | 0.0211 |
| ENSG00000125851.9 | 2899.011 | -0.385137476 | 0.1222 | -3.153 | 0.00162 | 0.0211 |
| ENSG00000168062.9 | 18.8845 | 0.560001988 | 0.1778 | 3.1498 | 0.00163 | 0.0213 |
| ENSG00000134716.9 | 247.5333 | 0.576147825 | 0.183 | 3.1482 | 0.00164 | 0.0214 |
| ENSG00000166341.7 | 814.527 | 0.323037798 | 0.1026 | 3.1477 | 0.00165 | 0.0214 |
| ENSG00000127241.16 | 210.0616 | 0.546690989 | 0.1739 | 3.1446 | 0.00166 | 0.0216 |
| ENSG00000078246.16 | 270.0743 | 0.366809551 | 0.1167 | 3.1419 | 0.00168 | 0.0216 |
| ENSG00000111058.7 | 549.7024 | 0.481695244 | 0.1533 | 3.143 | 0.00167 | 0.0216 |
| ENSG00000121957.12 | 778.6683 | 0.317349069 | 0.101 | 3.142 | 0.00168 | 0.0216 |
| ENSG00000160218.12 | 974.9696 | -0.363698479 | 0.1158 | -3.142 | 0.00168 | 0.0216 |
| ENSG00000197217.12 | 1957.184 | -0.286195917 | 0.0911 | -3.142 | 0.00168 | 0.0216 |
| ENSG00000104921.14 | 9.758611 | -0.93532449 | 0.2978 | -3.141 | 0.00169 | 0.0217 |
| ENSG00000148734.7 | 138.0853 | 0.415574962 | 0.1323 | 3.1408 | 0.00169 | 0.0217 |
| ENSG00000065559.14 | 4027.473 | -0.339909846 | 0.1083 | -3.139 | 0.00169 | 0.0217 |
| ENSG00000087266.15 | 666.1082 | 0.432965425 | 0.138 | 3.1385 | 0.0017 | 0.0217 |
| ENSG00000134769.21 | 6548.482 | 0.379181288 | 0.1208 | 3.1386 | 0.0017 | 0.0217 |
| ENSG00000136160.14 | 1638.146 | 0.758519289 | 0.2417 | 3.1382 | 0.0017 | 0.0217 |
| ENSG00000251165.5 | 25.44423 | 0.692751037 | 0.2207 | 3.1391 | 0.00169 | 0.0217 |
| ENSG00000061918.12 | 4101.445 | -0.35366404 | 0.1128 | -3.135 | 0.00172 | 0.0218 |
| ENSG00000111077.17 | 1226.343 | 0.313035111 | 0.0999 | 3.1349 | 0.00172 | 0.0218 |
| ENSG00000121310.16 | 846.0317 | 0.344850464 | 0.11 | 3.1363 | 0.00171 | 0.0218 |
| ENSG00000123612.15 | 324.6833 | -0.377657464 | 0.1204 | -3.136 | 0.00171 | 0.0218 |
| ENSG00000156395.12 | 969.4542 | -0.31543356 | 0.1006 | -3.135 | 0.00172 | 0.0218 |
| ENSG00000253982.1 | 260.5661 | 0.275754224 | 0.0879 | 3.1356 | 0.00172 | 0.0218 |
| ENSG00000164877.18 | 1041.924 | 0.311610873 | 0.0994 | 3.1342 | 0.00172 | 0.0218 |
| ENSG00000273032.1 | 625.4555 | -0.345746497 | 0.1103 | -3.134 | 0.00173 | 0.0218 |
| ENSG00000007129.17 | 8.958607 | -0.747872766 | 0.2387 | -3.133 | 0.00173 | 0.0219 |
| ENSG00000251432.6 | 29.22615 | 0.552565072 | 0.1764 | 3.1326 | 0.00173 | 0.0219 |
| ENSG00000171100.14 | 336.2271 | 0.298815043 | 0.0954 | 3.131 | 0.00174 | 0.022 |
| ENSG00000180900.17 | 2086.171 | 0.283100363 | 0.0905 | 3.1295 | 0.00175 | 0.0221 |
| ENSG00000189420.8 | 325.9817 | -0.315090496 | 0.1008 | -3.127 | 0.00177 | 0.0222 |
| ENSG00000221737.1 | 36.82231 | 0.364206284 | 0.1165 | 3.1269 | 0.00177 | 0.0222 |
| ENSG00000278996.1 | 33.18242 | 0.636764113 | 0.2037 | 3.1267 | 0.00177 | 0.0222 |
| ENSG00000280441.2 | 33.18242 | 0.636764113 | 0.2037 | 3.1267 | 0.00177 | 0.0222 |
| ENSG00000107104.18 | 1224.112 | 0.39803196 | 0.1273 | 3.1258 | 0.00177 | 0.0222 |
| ENSG00000162757.4 | 78.43365 | -0.372497802 | 0.1192 | -3.124 | 0.00178 | 0.0223 |
| ENSG00000079335.18 | 114.9396 | 0.562787155 | 0.1802 | 3.1226 | 0.00179 | 0.0224 |
| ENSG00000211459.2 | 333400.4 | 0.345622888 | 0.1107 | 3.1215 | 0.0018 | 0.0224 |
| ENSG00000176697.18 | 550.285 | -0.380841755 | 0.122 | -3.121 | 0.0018 | 0.0225 |
| ENSG00000147509.13 | 450.2442 | 0.387972162 | 0.1243 | 3.1203 | 0.00181 | 0.0225 |
| ENSG00000156642.16 | 7492.232 | -0.280563899 | 0.0899 | -3.12 | 0.00181 | 0.0225 |
| ENSG00000130368.5 | 348.7932 | -0.422118801 | 0.1354 | -3.118 | 0.00182 | 0.0225 |
| ENSG00000171016.11 | 311.2274 | 0.259982635 | 0.0834 | 3.1188 | 0.00182 | 0.0225 |
| ENSG00000279778.1 | 20.88642 | -0.627887216 | 0.2014 | -3.118 | 0.00182 | 0.0225 |
| ENSG00000148655.14 | 48.13792 | 0.298723395 | 0.0958 | 3.1177 | 0.00182 | 0.0225 |
| ENSG00000046889.18 | 1467.444 | 0.497602433 | 0.1597 | 3.1151 | 0.00184 | 0.0225 |
| ENSG00000147655.10 | 505.5763 | -0.385232243 | 0.1236 | -3.116 | 0.00184 | 0.0225 |
| ENSG00000197385.5 | 34.99324 | -0.467141399 | 0.1499 | -3.117 | 0.00183 | 0.0225 |
| ENSG00000214193.9 | 306.1267 | -0.258321195 | 0.0829 | -3.117 | 0.00183 | 0.0225 |
| ENSG00000228794.8 | 1252.796 | -0.283739373 | 0.091 | -3.116 | 0.00183 | 0.0225 |
| ENSG00000249087.6 | 173.9668 | 0.244081113 | 0.0784 | 3.1151 | 0.00184 | 0.0225 |
| ENSG00000257519.1 | 35.75395 | 0.493437151 | 0.1584 | 3.1156 | 0.00184 | 0.0225 |
| ENSG00000101079.20 | 4089.988 | -0.33309395 | 0.1069 | -3.115 | 0.00184 | 0.0226 |
| ENSG00000051128.18 | 304.5957 | 0.34537557 | 0.1109 | 3.113 | 0.00185 | 0.0226 |
| ENSG00000086159.12 | 39.16052 | 0.482270616 | 0.1549 | 3.1135 | 0.00185 | 0.0226 |
| ENSG00000114166.7 | 1697.523 | 0.426041141 | 0.1369 | 3.1128 | 0.00185 | 0.0226 |
| ENSG00000233139.1 | 231.6763 | 1.1757812 | 0.3778 | 3.1124 | 0.00186 | 0.0226 |
| ENSG00000244219.6 | 28.42193 | 0.373151695 | 0.1199 | 3.1112 | 0.00186 | 0.0227 |
| ENSG00000197147.12 | 3568.434 | -0.325119585 | 0.1045 | -3.11 | 0.00187 | 0.0227 |
| ENSG00000148053.15 | 23304.73 | 0.371146542 | 0.1194 | 3.1087 | 0.00188 | 0.0228 |
| ENSG00000274459.1 | 16.89942 | 0.550587985 | 0.1771 | 3.1086 | 0.00188 | 0.0228 |
| ENSG00000272158.1 | 14.29653 | 0.530168017 | 0.1706 | 3.1079 | 0.00188 | 0.0228 |
| ENSG00000145428.14 | 621.2943 | -0.272854496 | 0.0878 | -3.106 | 0.0019 | 0.0229 |
| ENSG00000270001.1 | 22.71474 | 0.817118568 | 0.2631 | 3.1062 | 0.0019 | 0.0229 |
| ENSG00000137561.4 | 88.00084 | 0.509948039 | 0.1643 | 3.1044 | 0.00191 | 0.023 |
| ENSG00000139174.10 | 1109.164 | -0.35139727 | 0.1132 | -3.103 | 0.00191 | 0.023 |
| ENSG00000146147.14 | 487.0592 | -0.305114947 | 0.0983 | -3.104 | 0.00191 | 0.023 |
| ENSG00000188818.12 | 367.856 | -0.600598619 | 0.1935 | -3.104 | 0.00191 | 0.023 |
| ENSG00000198075.9 | 121.8146 | 0.455024994 | 0.1466 | 3.1037 | 0.00191 | 0.023 |
| ENSG00000103569.9 | 64.77177 | -0.558144224 | 0.1799 | -3.103 | 0.00192 | 0.023 |
| ENSG00000228412.7 | 28.4685 | 0.60670058 | 0.1956 | 3.1023 | 0.00192 | 0.023 |
| ENSG00000259605.3 | 14.9859 | 0.707654083 | 0.2282 | 3.1008 | 0.00193 | 0.0231 |
| ENSG00000170848.15 | 13.79524 | 0.529457113 | 0.1708 | 3.1 | 0.00194 | 0.0231 |
| ENSG00000234722.3 | 163.1387 | 0.239198164 | 0.0772 | 3.1002 | 0.00193 | 0.0231 |
| ENSG00000139211.6 | 437.1581 | -0.355151493 | 0.1146 | -3.099 | 0.00194 | 0.0232 |
| ENSG00000066248.14 | 5376.992 | -0.360946459 | 0.1165 | -3.098 | 0.00195 | 0.0232 |
| ENSG00000078549.14 | 4030.989 | 0.320729218 | 0.1035 | 3.098 | 0.00195 | 0.0232 |
| ENSG00000225938.1 | 14.61604 | 0.793410671 | 0.2561 | 3.0978 | 0.00195 | 0.0232 |
| ENSG00000248222.5 | 34.42102 | -0.489693256 | 0.158 | -3.099 | 0.00194 | 0.0232 |
| ENSG00000130844.16 | 1029.352 | -0.295544859 | 0.0954 | -3.097 | 0.00196 | 0.0232 |
| ENSG00000103591.12 | 1141.838 | -0.287346096 | 0.0928 | -3.096 | 0.00196 | 0.0232 |
| ENSG00000154556.17 | 3835.872 | -0.265412571 | 0.0857 | -3.095 | 0.00196 | 0.0233 |
| ENSG00000116729.13 | 1437.094 | 0.374876432 | 0.1211 | 3.0946 | 0.00197 | 0.0233 |
| ENSG00000111110.11 | 2236.534 | -0.332937924 | 0.1076 | -3.094 | 0.00197 | 0.0233 |
| ENSG00000125084.11 | 56.05156 | -0.456959676 | 0.1477 | -3.093 | 0.00198 | 0.0234 |
| ENSG00000132824.13 | 6450.538 | -0.301299205 | 0.0974 | -3.092 | 0.00199 | 0.0234 |
| ENSG00000185742.6 | 3286.31 | -0.473458741 | 0.1531 | -3.092 | 0.00199 | 0.0234 |
| ENSG00000105479.15 | 32.07874 | 0.317920467 | 0.1028 | 3.0915 | 0.00199 | 0.0234 |
| ENSG00000237870.6 | 143.99 | 0.31004526 | 0.1003 | 3.0912 | 0.00199 | 0.0234 |
| ENSG00000125817.7 | 2001.959 | 0.26360644 | 0.0853 | 3.0905 | 0.002 | 0.0234 |
| ENSG00000137936.16 | 284.766 | 0.335708401 | 0.1087 | 3.0887 | 0.00201 | 0.0235 |
| ENSG00000147588.6 | 4856.896 | 0.491441344 | 0.1591 | 3.0888 | 0.00201 | 0.0235 |
| ENSG00000152034.10 | 285.1177 | -0.524577949 | 0.1698 | -3.089 | 0.00201 | 0.0235 |
| ENSG00000272944.1 | 86.6903 | -0.411716901 | 0.1333 | -3.088 | 0.00202 | 0.0236 |
| ENSG00000135622.12 | 859.6573 | -0.310298189 | 0.1005 | -3.087 | 0.00202 | 0.0236 |
| ENSG00000171126.7 | 156.0144 | -0.379868332 | 0.1231 | -3.086 | 0.00203 | 0.0236 |
| ENSG00000182890.4 | 48.33191 | -0.402565076 | 0.1305 | -3.086 | 0.00203 | 0.0236 |
| ENSG00000054277.13 | 745.8895 | -0.368552933 | 0.1195 | -3.085 | 0.00203 | 0.0236 |
| ENSG00000233435.2 | 17.35061 | -0.705222161 | 0.2286 | -3.084 | 0.00204 | 0.0237 |
| ENSG00000261739.2 | 44.92723 | -0.522360361 | 0.1694 | -3.084 | 0.00204 | 0.0237 |
| ENSG00000232035.1 | 1842.695 | 1.235235814 | 0.4006 | 3.0837 | 0.00204 | 0.0237 |
| ENSG00000242221.8 | 845.9836 | 1.51959177 | 0.4932 | 3.0812 | 0.00206 | 0.0239 |
| ENSG00000132938.19 | 665.1334 | -0.302361289 | 0.0982 | -3.08 | 0.00207 | 0.0239 |
| ENSG00000112851.14 | 2181.063 | 0.352733749 | 0.1146 | 3.0783 | 0.00208 | 0.024 |
| ENSG00000115419.12 | 9625.97 | -0.335740808 | 0.1091 | -3.077 | 0.00209 | 0.024 |
| ENSG00000116514.16 | 633.9849 | -0.348573827 | 0.1133 | -3.077 | 0.00209 | 0.024 |
| ENSG00000172348.14 | 6019.814 | -0.359410579 | 0.1168 | -3.077 | 0.00209 | 0.024 |
| ENSG00000182022.17 | 1233.906 | -0.280733032 | 0.0912 | -3.077 | 0.00209 | 0.024 |
| ENSG00000213722.8 | 493.373 | 0.384800162 | 0.1251 | 3.077 | 0.00209 | 0.024 |
| ENSG00000135912.10 | 314.4769 | 0.351247398 | 0.1142 | 3.075 | 0.0021 | 0.0241 |
| ENSG00000121691.4 | 510.1117 | 0.390878482 | 0.1271 | 3.0742 | 0.00211 | 0.0242 |
| ENSG00000136802.11 | 2454.322 | 0.309931827 | 0.1008 | 3.0739 | 0.00211 | 0.0242 |
| ENSG00000102468.10 | 1871.454 | -0.38742436 | 0.1261 | -3.072 | 0.00213 | 0.0243 |
| ENSG00000234377.7 | 227.3397 | 0.578274826 | 0.1883 | 3.0717 | 0.00213 | 0.0243 |
| ENSG00000282943.1 | 25.29166 | 0.461021414 | 0.1501 | 3.0718 | 0.00213 | 0.0243 |
| ENSG00000019505.7 | 5845.538 | -0.378180844 | 0.1232 | -3.071 | 0.00214 | 0.0243 |
| ENSG00000153956.15 | 4317.518 | -0.307784179 | 0.1002 | -3.07 | 0.00214 | 0.0243 |
| ENSG00000171612.6 | 641.7042 | 0.298102283 | 0.0971 | 3.0707 | 0.00214 | 0.0243 |
| ENSG00000187193.8 | 441.9892 | 0.930677626 | 0.3032 | 3.0696 | 0.00214 | 0.0243 |
| ENSG00000189195.10 | 2865.335 | -0.322402662 | 0.1051 | -3.069 | 0.00215 | 0.0244 |
| ENSG00000107758.15 | 7753.203 | -0.406547108 | 0.1325 | -3.068 | 0.00216 | 0.0244 |
| ENSG00000115902.10 | 2905.208 | 0.315117515 | 0.1028 | 3.0668 | 0.00216 | 0.0244 |
| ENSG00000148158.16 | 1748.713 | -0.294404077 | 0.096 | -3.067 | 0.00216 | 0.0244 |
| ENSG00000160716.5 | 1907.601 | -0.342529824 | 0.1117 | -3.067 | 0.00216 | 0.0244 |
| ENSG00000044574.7 | 4796.939 | -0.323787743 | 0.1056 | -3.066 | 0.00217 | 0.0245 |
| ENSG00000157653.11 | 15.40446 | 0.476611425 | 0.1555 | 3.0653 | 0.00217 | 0.0245 |
| ENSG00000069966.18 | 3167.611 | -0.289726611 | 0.0946 | -3.064 | 0.00219 | 0.0246 |
| ENSG00000101367.8 | 1199.39 | 0.269346069 | 0.088 | 3.0625 | 0.0022 | 0.0247 |
| ENSG00000118160.13 | 5367.377 | -0.328595041 | 0.1073 | -3.062 | 0.0022 | 0.0247 |
| ENSG00000163449.10 | 269.8136 | -0.359383909 | 0.1175 | -3.057 | 0.00223 | 0.025 |
| ENSG00000184731.5 | 307.7047 | -0.335374056 | 0.1097 | -3.057 | 0.00224 | 0.0251 |
| ENSG00000127472.10 | 82.20338 | 0.557088267 | 0.1823 | 3.0561 | 0.00224 | 0.0251 |
| ENSG00000186564.5 | 30.79721 | 0.559200883 | 0.183 | 3.0561 | 0.00224 | 0.0251 |
| ENSG00000176204.13 | 779.4848 | -0.333505939 | 0.1092 | -3.055 | 0.00225 | 0.0251 |
| ENSG00000115705.20 | 12.18984 | -0.565483004 | 0.1851 | -3.054 | 0.00226 | 0.0251 |
| ENSG00000148400.9 | 807.2814 | 0.398989471 | 0.1306 | 3.054 | 0.00226 | 0.0251 |
| ENSG00000248738.6 | 136.2528 | -0.476992738 | 0.1562 | -3.054 | 0.00226 | 0.0251 |
| ENSG00000135250.16 | 4681.435 | -0.305654122 | 0.1001 | -3.053 | 0.00226 | 0.0252 |
| ENSG00000205517.12 | 105.2172 | 0.38639086 | 0.1266 | 3.0509 | 0.00228 | 0.0253 |
| ENSG00000141577.13 | 289.9037 | 0.195802481 | 0.0642 | 3.0503 | 0.00229 | 0.0254 |
| ENSG00000233930.3 | 387.8518 | -0.369866903 | 0.1213 | -3.05 | 0.00229 | 0.0254 |
| ENSG00000119938.8 | 981.2739 | 0.704683057 | 0.2311 | 3.0491 | 0.00229 | 0.0254 |
| ENSG00000196132.11 | 306.6667 | 0.336594824 | 0.1104 | 3.0489 | 0.0023 | 0.0254 |
| ENSG00000123427.16 | 79.99033 | 0.315776027 | 0.1036 | 3.0477 | 0.00231 | 0.0255 |
| ENSG00000177807.7 | 2999.301 | 0.549609635 | 0.1804 | 3.0474 | 0.00231 | 0.0255 |
| ENSG00000164951.15 | 4131.954 | -0.340096979 | 0.1116 | -3.046 | 0.00232 | 0.0256 |
| ENSG00000224533.4 | 39.41842 | 0.303034459 | 0.0995 | 3.0459 | 0.00232 | 0.0256 |
| ENSG00000184414.2 | 123.1836 | 1.283680394 | 0.4217 | 3.0443 | 0.00233 | 0.0257 |
| ENSG00000111218.11 | 865.9493 | -0.302776232 | 0.0995 | -3.043 | 0.00235 | 0.0257 |
| ENSG00000197872.11 | 5259.344 | -0.338697681 | 0.1113 | -3.043 | 0.00234 | 0.0257 |
| ENSG00000198216.10 | 4534.162 | -0.37020281 | 0.1217 | -3.042 | 0.00235 | 0.0257 |
| ENSG00000277013.1 | 28.36848 | 0.457548317 | 0.1504 | 3.0428 | 0.00234 | 0.0257 |
| ENSG00000081853.14 | 145.2465 | 0.472944049 | 0.1555 | 3.0416 | 0.00235 | 0.0258 |
| ENSG00000123999.4 | 94.43974 | 0.346665935 | 0.1141 | 3.0391 | 0.00237 | 0.0258 |
| ENSG00000133026.12 | 6479.309 | -0.314613308 | 0.1035 | -3.04 | 0.00237 | 0.0258 |
| ENSG00000138741.10 | 89.08624 | -0.332862485 | 0.1095 | -3.039 | 0.00237 | 0.0258 |
| ENSG00000162944.10 | 713.4959 | 0.406869075 | 0.1339 | 3.0392 | 0.00237 | 0.0258 |
| ENSG00000196576.14 | 1555.195 | 0.243767477 | 0.0802 | 3.0403 | 0.00236 | 0.0258 |
| ENSG00000087077.12 | 296.6818 | 0.480668853 | 0.1582 | 3.0386 | 0.00238 | 0.0258 |
| ENSG00000204839.8 | 60.15731 | 0.376545904 | 0.124 | 3.0368 | 0.00239 | 0.026 |
| ENSG00000153317.14 | 2272.528 | -0.265817943 | 0.0875 | -3.036 | 0.00239 | 0.026 |
| ENSG00000077942.18 | 534.636 | 0.413791204 | 0.1363 | 3.0357 | 0.0024 | 0.026 |
| ENSG00000081842.17 | 327.7646 | -0.486238521 | 0.1603 | -3.034 | 0.00241 | 0.0261 |
| ENSG00000148798.10 | 6019.864 | -0.313574017 | 0.1033 | -3.034 | 0.00241 | 0.0261 |
| ENSG00000226758.1 | 23.35572 | 0.453885978 | 0.1496 | 3.0338 | 0.00241 | 0.0261 |
| ENSG00000272150.5 | 60.21588 | 0.380641959 | 0.1254 | 3.0342 | 0.00241 | 0.0261 |
| ENSG00000176320.2 | 82.67696 | 0.306503748 | 0.1011 | 3.0331 | 0.00242 | 0.0261 |
| ENSG00000243364.7 | 13.89845 | 0.530096796 | 0.1748 | 3.0324 | 0.00243 | 0.0261 |
| ENSG00000115762.16 | 6091.415 | -0.286087706 | 0.0944 | -3.031 | 0.00243 | 0.0262 |
| ENSG00000131389.16 | 671.737 | -0.243819377 | 0.0805 | -3.028 | 0.00246 | 0.0264 |
| ENSG00000267128.1 | 56.23651 | 0.347905349 | 0.1149 | 3.0274 | 0.00247 | 0.0265 |
| ENSG00000160282.13 | 137.9133 | 0.540419748 | 0.1785 | 3.0271 | 0.00247 | 0.0265 |
| ENSG00000165152.8 | 2676.286 | -0.374726749 | 0.1239 | -3.025 | 0.00249 | 0.0266 |
| ENSG00000168491.9 | 77.56262 | -0.395515359 | 0.1307 | -3.025 | 0.00248 | 0.0266 |
| ENSG00000279528.1 | 54.74584 | 0.38490744 | 0.1272 | 3.0252 | 0.00248 | 0.0266 |
| ENSG00000154822.16 | 824.9668 | -0.411955294 | 0.1363 | -3.022 | 0.00251 | 0.0268 |
| ENSG00000072163.19 | 732.8248 | 0.390598432 | 0.1294 | 3.0192 | 0.00253 | 0.0269 |
| ENSG00000111262.4 | 1240.236 | -0.474406409 | 0.157 | -3.021 | 0.00252 | 0.0269 |
| ENSG00000122012.13 | 339.5855 | -0.45940399 | 0.1521 | -3.02 | 0.00252 | 0.0269 |
| ENSG00000144668.11 | 388.2342 | -0.264257963 | 0.0875 | -3.02 | 0.00253 | 0.0269 |
| ENSG00000156486.7 | 399.886 | -0.503737659 | 0.1668 | -3.019 | 0.00253 | 0.0269 |
| ENSG00000170419.10 | 2245.409 | -0.361786005 | 0.1198 | -3.019 | 0.00253 | 0.0269 |
| ENSG00000224731.1 | 104.5684 | 0.356120408 | 0.1179 | 3.0196 | 0.00253 | 0.0269 |
| ENSG00000229953.1 | 31.23574 | 0.474218008 | 0.157 | 3.0203 | 0.00253 | 0.0269 |
| ENSG00000157654.17 | 525.5945 | -0.627594139 | 0.2079 | -3.019 | 0.00254 | 0.0269 |
| ENSG00000232528.3 | 50.64138 | 0.425491549 | 0.141 | 3.0186 | 0.00254 | 0.0269 |
| ENSG00000112379.8 | 4867.787 | -0.284938663 | 0.0944 | -3.018 | 0.00254 | 0.0269 |
| ENSG00000057468.6 | 24.27464 | -0.497071983 | 0.1647 | -3.018 | 0.00255 | 0.0269 |
| ENSG00000111801.15 | 224.4016 | -0.220829671 | 0.0732 | -3.017 | 0.00255 | 0.0269 |
| ENSG00000205089.7 | 284.459 | 0.35192547 | 0.1167 | 3.0168 | 0.00255 | 0.0269 |
| ENSG00000182197.10 | 483.7157 | -0.215972394 | 0.0716 | -3.016 | 0.00256 | 0.0269 |
| ENSG00000019991.15 | 180.1384 | 0.535292209 | 0.1776 | 3.0142 | 0.00258 | 0.027 |
| ENSG00000155097.11 | 6350.407 | -0.322807769 | 0.1071 | -3.015 | 0.00257 | 0.027 |
| ENSG00000210194.1 | 15.66046 | 0.926131114 | 0.3073 | 3.0143 | 0.00258 | 0.027 |
| ENSG00000226757.2 | 40.88455 | -0.559750492 | 0.1857 | -3.015 | 0.00257 | 0.027 |
| ENSG00000120833.13 | 198.6528 | -0.326477222 | 0.1083 | -3.013 | 0.00258 | 0.027 |
| ENSG00000282164.2 | 685.9577 | -0.3878015 | 0.1287 | -3.013 | 0.00259 | 0.027 |
| ENSG00000141433.12 | 711.6115 | -0.380841051 | 0.1264 | -3.012 | 0.00259 | 0.0271 |
| ENSG00000223774.5 | 20.77568 | 0.612563872 | 0.2033 | 3.0124 | 0.00259 | 0.0271 |
| ENSG00000138449.10 | 231.6915 | 0.312865235 | 0.1039 | 3.0109 | 0.0026 | 0.0271 |
| ENSG00000175874.9 | 7834.472 | -0.416951425 | 0.1385 | -3.011 | 0.00261 | 0.0271 |
| ENSG00000183117.18 | 1809.429 | -0.256692335 | 0.0853 | -3.011 | 0.0026 | 0.0271 |
| ENSG00000140718.19 | 2858.163 | -0.245791919 | 0.0817 | -3.01 | 0.00261 | 0.0272 |
| ENSG00000158156.7 | 113.57 | 0.30358517 | 0.1009 | 3.0091 | 0.00262 | 0.0272 |
| ENSG00000188662.6 | 11.98832 | 0.632337685 | 0.2102 | 3.0087 | 0.00262 | 0.0272 |
| ENSG00000154438.7 | 17.29303 | 0.498685464 | 0.1658 | 3.0081 | 0.00263 | 0.0272 |
| ENSG00000261553.5 | 214.7215 | -0.420277162 | 0.1398 | -3.007 | 0.00264 | 0.0273 |
| ENSG00000197977.3 | 576.1662 | 0.571546661 | 0.1901 | 3.0068 | 0.00264 | 0.0273 |
| ENSG00000153048.10 | 608.3462 | 0.328479871 | 0.1093 | 3.0056 | 0.00265 | 0.0274 |
| ENSG00000175077.5 | 116.6743 | -0.47978775 | 0.1597 | -3.005 | 0.00266 | 0.0274 |
| ENSG00000124570.17 | 757.305 | 0.248201352 | 0.0826 | 3.0041 | 0.00266 | 0.0274 |
| ENSG00000259446.5 | 10.01645 | 0.91348855 | 0.3041 | 3.0044 | 0.00266 | 0.0274 |
| ENSG00000007908.15 | 23.53905 | -1.147124276 | 0.382 | -3.003 | 0.00267 | 0.0275 |
| ENSG00000170027.6 | 29315.17 | -0.367795356 | 0.1225 | -3.003 | 0.00267 | 0.0275 |
| ENSG00000165868.13 | 5918.582 | -0.375668178 | 0.1251 | -3.003 | 0.00267 | 0.0275 |
| ENSG00000163347.5 | 67.3478 | 0.391488413 | 0.1304 | 3.002 | 0.00268 | 0.0275 |
| ENSG00000181019.12 | 411.8284 | 0.459060527 | 0.1529 | 3.0022 | 0.00268 | 0.0275 |
| ENSG00000128594.7 | 2040.888 | -0.280695508 | 0.0935 | -3.001 | 0.00269 | 0.0275 |
| ENSG00000279742.1 | 78.48605 | 0.323743952 | 0.1079 | 3.0014 | 0.00269 | 0.0275 |
| ENSG00000007168.12 | 11780.03 | -0.349372817 | 0.1164 | -3.001 | 0.00269 | 0.0275 |
| ENSG00000128285.4 | 369.0852 | -0.38900625 | 0.1297 | -3 | 0.0027 | 0.0275 |
| ENSG00000148700.14 | 5136.902 | 0.367237986 | 0.1224 | 3 | 0.0027 | 0.0275 |
| ENSG00000272031.2 | 1105.885 | -0.27156782 | 0.0905 | -3 | 0.0027 | 0.0275 |
| ENSG00000138757.14 | 6416.024 | -0.413392737 | 0.1378 | -2.999 | 0.00271 | 0.0276 |
| ENSG00000132840.9 | 97.33725 | 0.407267268 | 0.1358 | 2.9982 | 0.00272 | 0.0276 |
| ENSG00000250584.2 | 46.62702 | -0.468609853 | 0.1563 | -2.998 | 0.00271 | 0.0276 |
| ENSG00000138411.10 | 1367.006 | -0.319400525 | 0.1066 | -2.997 | 0.00272 | 0.0276 |
| ENSG00000144959.9 | 2112.156 | -0.329704757 | 0.11 | -2.997 | 0.00273 | 0.0276 |
| ENSG00000159082.17 | 5713.394 | -0.387197625 | 0.1292 | -2.997 | 0.00273 | 0.0276 |
| ENSG00000254122.2 | 268.8593 | 0.41323328 | 0.1379 | 2.9976 | 0.00272 | 0.0276 |
| ENSG00000254154.8 | 191.5654 | -0.306912356 | 0.1024 | -2.997 | 0.00273 | 0.0276 |
| ENSG00000014824.13 | 3553.431 | -0.31473741 | 0.1051 | -2.996 | 0.00274 | 0.0276 |
| ENSG00000157087.16 | 9966.794 | -0.432021018 | 0.1442 | -2.996 | 0.00274 | 0.0276 |
| ENSG00000198342.9 | 64.49148 | 0.427349475 | 0.1427 | 2.9956 | 0.00274 | 0.0276 |
| ENSG00000127955.15 | 2047.034 | -0.29181582 | 0.0975 | -2.994 | 0.00275 | 0.0277 |
| ENSG00000197324.8 | 764.4679 | 0.520834694 | 0.174 | 2.9939 | 0.00275 | 0.0277 |
| ENSG00000196305.17 | 2517.414 | -0.305142832 | 0.102 | -2.992 | 0.00277 | 0.0278 |
| ENSG00000119946.10 | 1395.257 | -0.269440194 | 0.0901 | -2.991 | 0.00278 | 0.0278 |
| ENSG00000143028.8 | 41.05993 | 0.539823783 | 0.1804 | 2.9916 | 0.00278 | 0.0278 |
| ENSG00000153822.13 | 402.6873 | 0.470831996 | 0.1574 | 2.9913 | 0.00278 | 0.0278 |
| ENSG00000177728.15 | 2751.043 | 0.297270117 | 0.0994 | 2.991 | 0.00278 | 0.0278 |
| ENSG00000122591.11 | 717.2108 | 0.283191562 | 0.0947 | 2.9907 | 0.00278 | 0.0278 |
| ENSG00000123570.3 | 810.4741 | -0.285052897 | 0.0954 | -2.988 | 0.00281 | 0.028 |
| ENSG00000132640.14 | 3745.961 | -0.306114225 | 0.1025 | -2.987 | 0.00282 | 0.028 |
| ENSG00000133019.11 | 2256.961 | -0.373403816 | 0.125 | -2.987 | 0.00282 | 0.028 |
| ENSG00000140961.12 | 88.59687 | 0.407878304 | 0.1365 | 2.9887 | 0.0028 | 0.028 |
| ENSG00000170579.15 | 7894.953 | -0.36472123 | 0.1221 | -2.987 | 0.00282 | 0.028 |
| ENSG00000248896.2 | 10.64533 | -0.529162543 | 0.1771 | -2.988 | 0.00281 | 0.028 |
| ENSG00000266017.1 | 18.05238 | 0.432187025 | 0.1447 | 2.9871 | 0.00282 | 0.028 |
| ENSG00000276029.1 | 18.05238 | 0.432187025 | 0.1447 | 2.9871 | 0.00282 | 0.028 |
| ENSG00000143851.15 | 35.886 | -0.667813933 | 0.2237 | -2.986 | 0.00283 | 0.028 |
| ENSG00000145817.16 | 797.4954 | -0.331448127 | 0.111 | -2.985 | 0.00283 | 0.0281 |
| ENSG00000068615.17 | 2454.816 | -0.327754578 | 0.1099 | -2.984 | 0.00285 | 0.0282 |
| ENSG00000137033.11 | 361.5383 | 0.514261279 | 0.1724 | 2.9831 | 0.00285 | 0.0282 |
| ENSG00000162174.12 | 1760.622 | 0.321654312 | 0.1078 | 2.9836 | 0.00285 | 0.0282 |
| ENSG00000249577.1 | 13.9846 | 0.582304597 | 0.1952 | 2.9833 | 0.00285 | 0.0282 |
| ENSG00000069702.10 | 326.6522 | 0.431312108 | 0.1447 | 2.9808 | 0.00287 | 0.0283 |
| ENSG00000182255.6 | 377.5583 | -0.279724338 | 0.0938 | -2.981 | 0.00287 | 0.0283 |
| ENSG00000088538.12 | 5093.303 | -0.3427494 | 0.115 | -2.98 | 0.00288 | 0.0283 |
| ENSG00000152127.8 | 2349.465 | -0.286301845 | 0.0961 | -2.98 | 0.00288 | 0.0283 |
| ENSG00000141510.16 | 180.8486 | 0.382829032 | 0.1285 | 2.9793 | 0.00289 | 0.0284 |
| ENSG00000140905.9 | 961.9018 | 0.255589386 | 0.0859 | 2.9762 | 0.00292 | 0.0286 |
| ENSG00000158560.14 | 3842.408 | -0.397787573 | 0.1336 | -2.976 | 0.00292 | 0.0286 |
| ENSG00000168077.13 | 1400.13 | 0.541161798 | 0.1818 | 2.9765 | 0.00292 | 0.0286 |
| ENSG00000175147.11 | 113.6637 | 0.37452246 | 0.1259 | 2.9757 | 0.00292 | 0.0286 |
| ENSG00000203737.3 | 41.79145 | -0.457389243 | 0.1537 | -2.976 | 0.00292 | 0.0286 |
| ENSG00000115808.11 | 1237.504 | -0.271622974 | 0.0913 | -2.974 | 0.00294 | 0.0287 |
| ENSG00000203857.9 | 41.90087 | -0.474097445 | 0.1595 | -2.973 | 0.00295 | 0.0288 |
| ENSG00000113327.14 | 4188.989 | -0.403512268 | 0.1358 | -2.972 | 0.00296 | 0.0288 |
| ENSG00000157191.19 | 407.8733 | 0.271216945 | 0.0913 | 2.9704 | 0.00297 | 0.0288 |
| ENSG00000205106.4 | 37.08522 | 0.469244832 | 0.158 | 2.9706 | 0.00297 | 0.0288 |
| ENSG00000224310.1 | 104.1505 | 0.350268389 | 0.1179 | 2.9708 | 0.00297 | 0.0288 |
| ENSG00000248874.5 | 134.34 | 0.261322524 | 0.088 | 2.971 | 0.00297 | 0.0288 |
| ENSG00000255390.1 | 28.87219 | 0.871677909 | 0.2934 | 2.9711 | 0.00297 | 0.0288 |
| ENSG00000082497.11 | 183.4942 | -0.454657612 | 0.1532 | -2.969 | 0.00299 | 0.0289 |
| ENSG00000087086.14 | 15374.71 | 0.28828037 | 0.0971 | 2.9681 | 0.003 | 0.0289 |
| ENSG00000100802.14 | 75.94294 | 0.256930485 | 0.0866 | 2.9684 | 0.00299 | 0.0289 |
| ENSG00000103876.11 | 183.8155 | 0.34615034 | 0.1166 | 2.9678 | 0.003 | 0.0289 |
| ENSG00000110514.19 | 5381.607 | -0.333781151 | 0.1125 | -2.968 | 0.003 | 0.0289 |
| ENSG00000163393.12 | 524.5655 | -0.258185023 | 0.087 | -2.968 | 0.00299 | 0.0289 |
| ENSG00000198932.12 | 6117.847 | -0.306899929 | 0.1034 | -2.969 | 0.00298 | 0.0289 |
| ENSG00000064490.13 | 251.3075 | 0.261741623 | 0.0883 | 2.9657 | 0.00302 | 0.029 |
| ENSG00000138018.17 | 2175.935 | -0.344816949 | 0.1163 | -2.966 | 0.00302 | 0.029 |
| ENSG00000270959.1 | 73.5895 | 0.382587898 | 0.129 | 2.9659 | 0.00302 | 0.029 |
| ENSG00000101888.11 | 134.5516 | 0.574780478 | 0.1939 | 2.9649 | 0.00303 | 0.0291 |
| ENSG00000091656.15 | 495.5503 | 0.465569038 | 0.1571 | 2.9643 | 0.00303 | 0.0291 |
| ENSG00000241231.1 | 71.5702 | 0.721509069 | 0.2434 | 2.9645 | 0.00303 | 0.0291 |
| ENSG00000275342.4 | 593.4778 | -0.296481507 | 0.1 | -2.964 | 0.00304 | 0.0291 |
| ENSG00000280434.1 | 125.414 | 0.295312705 | 0.0996 | 2.9635 | 0.00304 | 0.0291 |
| ENSG00000011405.13 | 1751.846 | 0.39543415 | 0.1335 | 2.9628 | 0.00305 | 0.0291 |
| ENSG00000038427.15 | 1252.574 | 0.445747274 | 0.1504 | 2.9629 | 0.00305 | 0.0291 |
| ENSG00000169297.7 | 14.48423 | 0.57873463 | 0.1954 | 2.9619 | 0.00306 | 0.0292 |
| ENSG00000082512.14 | 371.2622 | -0.24983821 | 0.0844 | -2.961 | 0.00306 | 0.0292 |
| ENSG00000164128.6 | 635.3891 | -0.279097539 | 0.0943 | -2.961 | 0.00307 | 0.0292 |
| ENSG00000143994.13 | 51.35056 | 0.358130358 | 0.121 | 2.9604 | 0.00307 | 0.0292 |
| ENSG00000154174.7 | 2960.152 | -0.295959323 | 0.1 | -2.96 | 0.00307 | 0.0292 |
| ENSG00000111432.4 | 12.83851 | 0.733537521 | 0.248 | 2.9584 | 0.00309 | 0.0293 |
| ENSG00000124782.19 | 179.6365 | 0.366549016 | 0.1239 | 2.9585 | 0.00309 | 0.0293 |
| ENSG00000205927.4 | 749.1283 | 0.410422645 | 0.1387 | 2.9589 | 0.00309 | 0.0293 |
| ENSG00000104723.20 | 1863.645 | -0.317804954 | 0.1074 | -2.958 | 0.0031 | 0.0293 |
| ENSG00000264617.1 | 8.871917 | -0.785687904 | 0.2657 | -2.957 | 0.00311 | 0.0293 |
| ENSG00000278727.1 | 158.1929 | -0.367562698 | 0.1243 | -2.957 | 0.0031 | 0.0293 |
| ENSG00000116299.16 | 1097.992 | -0.259703541 | 0.0878 | -2.957 | 0.00311 | 0.0293 |
| ENSG00000068354.15 | 725.2992 | -0.249537112 | 0.0844 | -2.956 | 0.00312 | 0.0294 |
| ENSG00000171617.13 | 35795.9 | -0.372046992 | 0.1258 | -2.956 | 0.00311 | 0.0294 |
| ENSG00000183242.11 | 134.5917 | 1.223879284 | 0.4141 | 2.9553 | 0.00312 | 0.0294 |
| ENSG00000183908.5 | 285.8419 | -0.477303664 | 0.1616 | -2.955 | 0.00313 | 0.0295 |
| ENSG00000165801.9 | 695.5584 | 0.433069631 | 0.1466 | 2.9537 | 0.00314 | 0.0295 |
| ENSG00000101680.13 | 198.3792 | 0.459422164 | 0.1556 | 2.9529 | 0.00315 | 0.0296 |
| ENSG00000134909.18 | 5819.726 | -0.279659892 | 0.0947 | -2.952 | 0.00316 | 0.0296 |
| ENSG00000225972.1 | 2785.404 | -1.835374633 | 0.6219 | -2.951 | 0.00317 | 0.0297 |
| ENSG00000177182.10 | 541.2677 | -0.333392641 | 0.113 | -2.95 | 0.00318 | 0.0297 |
| ENSG00000226308.1 | 22.82437 | 0.526373301 | 0.1784 | 2.9501 | 0.00318 | 0.0297 |
| ENSG00000249947.2 | 12.14868 | -0.534996835 | 0.1814 | -2.95 | 0.00318 | 0.0297 |
| ENSG00000188643.10 | 582.5649 | 0.484145209 | 0.1643 | 2.9471 | 0.00321 | 0.0299 |
| ENSG00000117152.13 | 11851.7 | -0.406905783 | 0.1382 | -2.944 | 0.00324 | 0.0302 |
| ENSG00000226124.6 | 76.89888 | -0.284065328 | 0.0965 | -2.943 | 0.00325 | 0.0303 |
| ENSG00000279082.3 | 38.51004 | 0.625326514 | 0.2125 | 2.9425 | 0.00326 | 0.0303 |
| ENSG00000188511.12 | 27.39646 | 0.424014732 | 0.1441 | 2.9417 | 0.00326 | 0.0304 |
| ENSG00000185760.15 | 2276.802 | -0.358591478 | 0.122 | -2.94 | 0.00328 | 0.0305 |
| ENSG00000167191.11 | 8235.828 | 0.469161154 | 0.1596 | 2.9392 | 0.00329 | 0.0306 |
| ENSG00000166908.17 | 1513.131 | -0.309426312 | 0.1053 | -2.938 | 0.0033 | 0.0306 |
| ENSG00000114948.12 | 2985.614 | -0.321472306 | 0.1095 | -2.937 | 0.00332 | 0.0308 |
| ENSG00000118733.16 | 1721.384 | -0.385327172 | 0.1312 | -2.936 | 0.00332 | 0.0308 |
| ENSG00000135018.13 | 4878.544 | -0.30978775 | 0.1055 | -2.936 | 0.00333 | 0.0308 |
| ENSG00000138769.10 | 496.5422 | -0.390752858 | 0.1332 | -2.934 | 0.00335 | 0.0309 |
| ENSG00000091664.7 | 430.336 | -0.317831603 | 0.1084 | -2.933 | 0.00336 | 0.031 |
| ENSG00000152463.14 | 10.16825 | 0.568756366 | 0.1939 | 2.9329 | 0.00336 | 0.031 |
| ENSG00000147145.12 | 738.5035 | 1.284273089 | 0.438 | 2.9324 | 0.00336 | 0.031 |
| ENSG00000173611.17 | 2970.691 | -0.299247291 | 0.1021 | -2.932 | 0.00337 | 0.031 |
| ENSG00000073464.11 | 3529.489 | -0.385683731 | 0.1316 | -2.932 | 0.00337 | 0.031 |
| ENSG00000170647.3 | 71.48345 | 0.401456124 | 0.137 | 2.9311 | 0.00338 | 0.0311 |
| ENSG00000251339.5 | 24.23247 | 0.456949308 | 0.1559 | 2.9307 | 0.00338 | 0.0311 |
| ENSG00000151025.9 | 2046.753 | -0.442226852 | 0.151 | -2.93 | 0.00339 | 0.0311 |
| ENSG00000151690.14 | 3282.77 | -0.368048106 | 0.1256 | -2.93 | 0.00339 | 0.0311 |
| ENSG00000174370.9 | 33.79622 | -0.438222102 | 0.1496 | -2.929 | 0.0034 | 0.0311 |
| ENSG00000275465.5 | 17.58412 | 0.574765665 | 0.1962 | 2.9295 | 0.0034 | 0.0311 |
| ENSG00000246859.2 | 186.9391 | -0.358355031 | 0.1224 | -2.928 | 0.00342 | 0.0312 |
| ENSG00000007237.18 | 14629.47 | -0.358760967 | 0.1226 | -2.927 | 0.00343 | 0.0313 |
| ENSG00000165490.12 | 18.71516 | -0.393439523 | 0.1344 | -2.926 | 0.00343 | 0.0313 |
| ENSG00000101972.18 | 1537.33 | 0.170743344 | 0.0584 | 2.9257 | 0.00344 | 0.0314 |
| ENSG00000278195.1 | 366.3124 | -0.338015544 | 0.1156 | -2.924 | 0.00346 | 0.0315 |
| ENSG00000171303.6 | 775.8246 | -0.305766872 | 0.1046 | -2.923 | 0.00346 | 0.0315 |
| ENSG00000270953.1 | 286.141 | -0.47191955 | 0.1614 | -2.923 | 0.00347 | 0.0315 |
| ENSG00000159792.9 | 281.7337 | 0.233658303 | 0.08 | 2.9218 | 0.00348 | 0.0316 |
| ENSG00000169105.7 | 72.68189 | 0.376167356 | 0.1288 | 2.9202 | 0.0035 | 0.0318 |
| ENSG00000132854.18 | 38.29814 | -0.382664769 | 0.1311 | -2.92 | 0.0035 | 0.0318 |
| ENSG00000164574.15 | 588.4803 | 0.298451648 | 0.1022 | 2.9189 | 0.00351 | 0.0318 |
| ENSG00000160145.15 | 7245.997 | -0.298659534 | 0.1023 | -2.918 | 0.00352 | 0.0319 |
| ENSG00000110090.12 | 783.2193 | 0.39983814 | 0.137 | 2.9181 | 0.00352 | 0.0319 |
| ENSG00000147475.14 | 682.9745 | 0.26474629 | 0.0908 | 2.9168 | 0.00354 | 0.032 |
| ENSG00000151729.10 | 3890.113 | -0.273853744 | 0.0939 | -2.916 | 0.00354 | 0.032 |
| ENSG00000082701.14 | 3310.822 | -0.284568125 | 0.0976 | -2.915 | 0.00355 | 0.032 |
| ENSG00000121454.5 | 71.33518 | -0.359901399 | 0.1235 | -2.915 | 0.00355 | 0.032 |
| ENSG00000260166.1 | 28.71595 | -0.513415957 | 0.1762 | -2.915 | 0.00356 | 0.0321 |
| ENSG00000198569.9 | 43.26902 | 0.409530466 | 0.1405 | 2.9142 | 0.00357 | 0.0321 |
| ENSG00000100068.11 | 64.22541 | 0.33131292 | 0.1137 | 2.9135 | 0.00357 | 0.0321 |
| ENSG00000163749.17 | 96.53861 | -0.322160063 | 0.1106 | -2.913 | 0.00358 | 0.0321 |
| ENSG00000109971.13 | 20385.95 | -0.381411487 | 0.131 | -2.912 | 0.00359 | 0.0322 |
| ENSG00000116117.17 | 210.5386 | 0.488742653 | 0.1679 | 2.9109 | 0.0036 | 0.0322 |
| ENSG00000163399.15 | 16164.18 | -0.375630224 | 0.129 | -2.911 | 0.00361 | 0.0322 |
| ENSG00000176907.4 | 207.8654 | -0.436412235 | 0.1499 | -2.911 | 0.0036 | 0.0322 |
| ENSG00000228775.7 | 24.53363 | 0.524547049 | 0.1802 | 2.9113 | 0.0036 | 0.0322 |
| ENSG00000266094.7 | 490.3818 | -0.35246196 | 0.1211 | -2.911 | 0.00361 | 0.0322 |
| ENSG00000272916.5 | 59.47819 | -0.422077647 | 0.145 | -2.91 | 0.00361 | 0.0322 |
| ENSG00000040933.15 | 3607.661 | -0.323045014 | 0.111 | -2.91 | 0.00362 | 0.0322 |
| ENSG00000072954.6 | 980.6888 | -0.330063436 | 0.1134 | -2.909 | 0.00362 | 0.0322 |
| ENSG00000086061.15 | 5839.432 | -0.318159319 | 0.1094 | -2.909 | 0.00362 | 0.0322 |
| ENSG00000272933.1 | 249.5145 | 0.296408233 | 0.1019 | 2.9095 | 0.00362 | 0.0322 |
| ENSG00000049239.12 | 661.0623 | 0.239247524 | 0.0822 | 2.909 | 0.00363 | 0.0322 |
| ENSG00000070961.15 | 11800.08 | -0.336602219 | 0.1157 | -2.908 | 0.00363 | 0.0323 |
| ENSG00000120251.18 | 12319.55 | -0.270206289 | 0.093 | -2.907 | 0.00365 | 0.0324 |
| ENSG00000129159.6 | 2007.107 | -0.329068693 | 0.1132 | -2.906 | 0.00366 | 0.0324 |
| ENSG00000145536.15 | 130.6585 | -0.340604784 | 0.1172 | -2.906 | 0.00366 | 0.0324 |
| ENSG00000198934.4 | 1929.511 | -0.29860629 | 0.1028 | -2.906 | 0.00366 | 0.0324 |
| ENSG00000101911.12 | 734.3412 | -0.347053646 | 0.1195 | -2.904 | 0.00368 | 0.0324 |
| ENSG00000111961.17 | 2456.611 | 0.359284581 | 0.1237 | 2.9045 | 0.00368 | 0.0324 |
| ENSG00000154310.16 | 2226.392 | 0.257832627 | 0.0888 | 2.9051 | 0.00367 | 0.0324 |
| ENSG00000162694.13 | 872.9556 | -0.290218471 | 0.0999 | -2.904 | 0.00368 | 0.0324 |
| ENSG00000169871.12 | 619.5536 | 0.320174045 | 0.1102 | 2.9054 | 0.00367 | 0.0324 |
| ENSG00000273328.5 | 11.1994 | 0.452376187 | 0.1558 | 2.9043 | 0.00368 | 0.0324 |
| ENSG00000116106.11 | 3962.491 | -0.391869513 | 0.135 | -2.903 | 0.00369 | 0.0325 |
| ENSG00000162337.11 | 208.1201 | 0.461969193 | 0.1592 | 2.9024 | 0.0037 | 0.0326 |
| ENSG00000170091.10 | 10649.33 | -0.311822337 | 0.1075 | -2.902 | 0.00371 | 0.0326 |
| ENSG00000211448.11 | 2231.427 | 0.512840853 | 0.1768 | 2.9011 | 0.00372 | 0.0326 |
| ENSG00000172349.17 | 74.12363 | -0.32876539 | 0.1134 | -2.9 | 0.00373 | 0.0327 |
| ENSG00000261840.2 | 28.09983 | 0.398315468 | 0.1373 | 2.9002 | 0.00373 | 0.0327 |
| ENSG00000160710.15 | 6944.278 | -0.227216593 | 0.0784 | -2.9 | 0.00374 | 0.0327 |
| ENSG00000259319.1 | 157.7032 | -0.303850796 | 0.1048 | -2.899 | 0.00374 | 0.0327 |
| ENSG00000100320.22 | 6172.891 | -0.278302464 | 0.096 | -2.899 | 0.00375 | 0.0328 |
| ENSG00000087095.12 | 3387.795 | -0.325596046 | 0.1124 | -2.897 | 0.00377 | 0.0329 |
| ENSG00000254634.4 | 21.56644 | -0.468283944 | 0.1617 | -2.897 | 0.00377 | 0.0329 |
| ENSG00000144057.15 | 974.7211 | -0.234385732 | 0.0809 | -2.896 | 0.00378 | 0.0329 |
| ENSG00000110786.17 | 3014.948 | -0.291712009 | 0.1007 | -2.896 | 0.00378 | 0.033 |
| ENSG00000262075.3 | 27.3833 | 0.387260086 | 0.1337 | 2.8955 | 0.00379 | 0.033 |
| ENSG00000101938.14 | 1119.552 | 0.386003811 | 0.1333 | 2.8949 | 0.00379 | 0.033 |
| ENSG00000183090.5 | 576.0309 | -0.55457361 | 0.1916 | -2.895 | 0.00379 | 0.033 |
| ENSG00000272780.5 | 369.42 | -0.341953154 | 0.1182 | -2.894 | 0.00381 | 0.033 |
| ENSG00000169032.9 | 4739.827 | -0.336077367 | 0.1162 | -2.893 | 0.00382 | 0.0331 |
| ENSG00000189350.12 | 258.8218 | 0.299550553 | 0.1035 | 2.8929 | 0.00382 | 0.0331 |
| ENSG00000279456.1 | 73.93146 | -0.390580656 | 0.135 | -2.893 | 0.00382 | 0.0331 |
| ENSG00000270136.5 | 66.16847 | 0.75121979 | 0.2599 | 2.8908 | 0.00384 | 0.0333 |
| ENSG00000109436.7 | 2243.115 | -0.338586759 | 0.1172 | -2.89 | 0.00385 | 0.0333 |
| ENSG00000150760.12 | 1318.443 | 0.362319478 | 0.1254 | 2.8898 | 0.00386 | 0.0333 |
| ENSG00000163110.14 | 1442.019 | 0.441203801 | 0.1527 | 2.8889 | 0.00387 | 0.0333 |
| ENSG00000168830.7 | 120.928 | -0.289319943 | 0.1002 | -2.888 | 0.00387 | 0.0333 |
| ENSG00000173452.13 | 605.4813 | -0.365452374 | 0.1265 | -2.888 | 0.00387 | 0.0333 |
| ENSG00000186792.16 | 143.4105 | -0.314523353 | 0.1089 | -2.889 | 0.00387 | 0.0333 |
| ENSG00000138685.12 | 890.0545 | 0.566666391 | 0.1963 | 2.8866 | 0.00389 | 0.0335 |
| ENSG00000197901.11 | 43.02357 | 0.639148189 | 0.2214 | 2.8869 | 0.00389 | 0.0335 |
| ENSG00000113790.10 | 150.8398 | 0.295120011 | 0.1023 | 2.8858 | 0.0039 | 0.0335 |
| ENSG00000132932.16 | 2762.853 | -0.387299511 | 0.1342 | -2.886 | 0.00391 | 0.0335 |
| ENSG00000162687.16 | 568.1079 | -0.312573034 | 0.1083 | -2.885 | 0.00391 | 0.0335 |
| ENSG00000073712.14 | 1395.703 | 0.43534768 | 0.1509 | 2.8844 | 0.00392 | 0.0336 |
| ENSG00000255284.1 | 72.20659 | 0.267269592 | 0.0927 | 2.8841 | 0.00393 | 0.0336 |
| ENSG00000082397.17 | 5415.513 | -0.279489477 | 0.097 | -2.883 | 0.00394 | 0.0337 |
| ENSG00000169933.12 | 1917.896 | -0.429734351 | 0.1491 | -2.883 | 0.00394 | 0.0337 |
| ENSG00000005075.15 | 492.7869 | 0.2481315 | 0.0861 | 2.8813 | 0.00396 | 0.0338 |
| ENSG00000144339.11 | 1300.841 | -0.347461366 | 0.1207 | -2.88 | 0.00398 | 0.0339 |
| ENSG00000212694.8 | 308.2588 | 0.275086177 | 0.0955 | 2.88 | 0.00398 | 0.0339 |
| ENSG00000102032.12 | 171.7778 | 0.326494466 | 0.1134 | 2.879 | 0.00399 | 0.034 |
| ENSG00000139977.13 | 1420.242 | -0.267720644 | 0.093 | -2.878 | 0.004 | 0.034 |
| ENSG00000076604.14 | 333.5392 | 0.427443621 | 0.1486 | 2.8772 | 0.00401 | 0.0341 |
| ENSG00000092853.13 | 52.69903 | -0.385627989 | 0.134 | -2.877 | 0.00401 | 0.0341 |
| ENSG00000054793.13 | 14575.6 | -0.319030182 | 0.1109 | -2.876 | 0.00402 | 0.0341 |
| ENSG00000158301.18 | 2579.324 | -0.310519929 | 0.1079 | -2.877 | 0.00402 | 0.0341 |
| ENSG00000196876.14 | 4588.915 | -0.399960169 | 0.1391 | -2.876 | 0.00403 | 0.0341 |
| ENSG00000103512.14 | 3335.447 | -0.315767763 | 0.1098 | -2.875 | 0.00404 | 0.0342 |
| ENSG00000255330.9 | 2340.348 | -0.349731604 | 0.1216 | -2.875 | 0.00403 | 0.0342 |
| ENSG00000136267.13 | 2375.683 | -0.357418352 | 0.1244 | -2.874 | 0.00405 | 0.0343 |
| ENSG00000010818.8 | 4215.54 | -0.366693009 | 0.1277 | -2.872 | 0.00408 | 0.0343 |
| ENSG00000075035.9 | 1053.302 | -0.346456627 | 0.1206 | -2.873 | 0.00406 | 0.0343 |
| ENSG00000118402.5 | 1129.497 | -0.302737623 | 0.1055 | -2.87 | 0.0041 | 0.0343 |
| ENSG00000136044.11 | 846.3919 | 0.292833617 | 0.1019 | 2.8729 | 0.00407 | 0.0343 |
| ENSG00000161180.10 | 9.166609 | 0.513044413 | 0.1786 | 2.8723 | 0.00408 | 0.0343 |
| ENSG00000163239.12 | 28.40356 | 0.498355006 | 0.1736 | 2.8709 | 0.00409 | 0.0343 |
| ENSG00000177144.6 | 62.36554 | -0.50040401 | 0.1743 | -2.87 | 0.0041 | 0.0343 |
| ENSG00000187416.11 | 315.8073 | 0.371920946 | 0.1295 | 2.8713 | 0.00409 | 0.0343 |
| ENSG00000188133.5 | 52.68972 | -0.609776207 | 0.2123 | -2.872 | 0.00408 | 0.0343 |
| ENSG00000223414.2 | 60.15996 | -0.461948525 | 0.1609 | -2.872 | 0.00408 | 0.0343 |
| ENSG00000257270.1 | 38.37306 | 0.338339531 | 0.1178 | 2.8719 | 0.00408 | 0.0343 |
| ENSG00000271121.2 | 62.36554 | -0.50040401 | 0.1743 | -2.87 | 0.0041 | 0.0343 |
| ENSG00000280138.1 | 600.6269 | 0.432598333 | 0.1507 | 2.8711 | 0.00409 | 0.0343 |
| ENSG00000171004.17 | 425.3986 | -0.357873893 | 0.1247 | -2.869 | 0.00411 | 0.0344 |
| ENSG00000122359.17 | 3101.931 | -0.178450007 | 0.0622 | -2.869 | 0.00412 | 0.0344 |
| ENSG00000155657.25 | 648.0138 | -0.192821174 | 0.0672 | -2.868 | 0.00413 | 0.0344 |
| ENSG00000115602.16 | 52.38158 | 1.289688681 | 0.4498 | 2.8674 | 0.00414 | 0.0345 |
| ENSG00000182013.17 | 5520.146 | -0.346187968 | 0.1207 | -2.867 | 0.00414 | 0.0345 |
| ENSG00000010704.18 | 56.82584 | 0.418207903 | 0.1459 | 2.8662 | 0.00415 | 0.0345 |
| ENSG00000138448.11 | 2472.714 | 0.339466319 | 0.1184 | 2.8663 | 0.00415 | 0.0345 |
| ENSG00000240207.6 | 197.7832 | -0.323281991 | 0.1128 | -2.866 | 0.00415 | 0.0345 |
| ENSG00000153234.13 | 1100.785 | -0.496102871 | 0.1731 | -2.865 | 0.00416 | 0.0346 |
| ENSG00000198513.11 | 3539.64 | -0.293927599 | 0.1026 | -2.865 | 0.00417 | 0.0346 |
| ENSG00000134853.11 | 1744.931 | 0.416627884 | 0.1455 | 2.8641 | 0.00418 | 0.0347 |
| ENSG00000116675.15 | 9799.347 | -0.376299856 | 0.1314 | -2.863 | 0.00419 | 0.0347 |
| ENSG00000110218.8 | 414.2727 | -0.405826155 | 0.1418 | -2.862 | 0.00421 | 0.0348 |
| ENSG00000171302.16 | 575.9467 | -0.247813937 | 0.0866 | -2.862 | 0.00421 | 0.0348 |
| ENSG00000184898.6 | 132.9952 | 0.276607121 | 0.0967 | 2.8613 | 0.00422 | 0.0349 |
| ENSG00000102362.15 | 186.7841 | 0.46992753 | 0.1643 | 2.8606 | 0.00423 | 0.0349 |
| ENSG00000150394.13 | 1618.654 | -0.36510681 | 0.1276 | -2.86 | 0.00423 | 0.0349 |
| ENSG00000131771.13 | 4235.129 | 0.339747499 | 0.1188 | 2.8598 | 0.00424 | 0.035 |
| ENSG00000164270.17 | 70.69774 | -0.407377925 | 0.1425 | -2.859 | 0.00425 | 0.035 |
| ENSG00000137040.9 | 1241.283 | -0.265384381 | 0.0928 | -2.858 | 0.00426 | 0.035 |
| ENSG00000152583.12 | 39185.79 | 0.255533426 | 0.0894 | 2.8583 | 0.00426 | 0.035 |
| ENSG00000180176.14 | 41.81897 | -0.55574151 | 0.1944 | -2.859 | 0.00426 | 0.035 |
| ENSG00000130208.9 | 163.2219 | 0.545539312 | 0.1909 | 2.8577 | 0.00427 | 0.035 |
| ENSG00000149575.5 | 3662.072 | -0.348364618 | 0.1219 | -2.858 | 0.00427 | 0.035 |
| ENSG00000022355.16 | 6281.338 | -0.424027371 | 0.1484 | -2.857 | 0.00427 | 0.035 |
| ENSG00000221823.10 | 12391.52 | -0.303141881 | 0.1061 | -2.857 | 0.00428 | 0.035 |
| ENSG00000129493.14 | 550.0915 | 0.315586426 | 0.1106 | 2.8535 | 0.00432 | 0.0354 |
| ENSG00000166173.10 | 1425 | -0.231690197 | 0.0812 | -2.853 | 0.00433 | 0.0354 |
| ENSG00000214455.4 | 61.38224 | 0.340871347 | 0.1195 | 2.8527 | 0.00433 | 0.0354 |
| ENSG00000281005.1 | 38.4392 | 0.291557132 | 0.1022 | 2.853 | 0.00433 | 0.0354 |
| ENSG00000104327.7 | 1422.394 | -0.355321487 | 0.1246 | -2.852 | 0.00435 | 0.0354 |
| ENSG00000120265.16 | 5215.603 | -0.327073387 | 0.1147 | -2.852 | 0.00435 | 0.0354 |
| ENSG00000153898.12 | 19.40372 | 0.401898521 | 0.141 | 2.8509 | 0.00436 | 0.0355 |
| ENSG00000164484.11 | 305.5149 | -0.420446001 | 0.1475 | -2.851 | 0.00436 | 0.0355 |
| ENSG00000261325.1 | 44.05721 | -0.621288135 | 0.218 | -2.85 | 0.00437 | 0.0355 |
| ENSG00000101333.16 | 534.1941 | -0.271908839 | 0.0954 | -2.849 | 0.00438 | 0.0356 |
| ENSG00000197632.8 | 10.36796 | -0.613650577 | 0.2154 | -2.849 | 0.00438 | 0.0356 |
| ENSG00000203867.7 | 100.0823 | -0.307144871 | 0.1078 | -2.849 | 0.00438 | 0.0356 |
| ENSG00000273064.1 | 24.74615 | 0.399907153 | 0.1404 | 2.8493 | 0.00438 | 0.0356 |
| ENSG00000145794.16 | 775.6262 | 0.463910444 | 0.1629 | 2.8471 | 0.00441 | 0.0357 |
| ENSG00000070718.11 | 2243.075 | -0.321747145 | 0.113 | -2.846 | 0.00443 | 0.0358 |
| ENSG00000187068.2 | 1088.547 | 0.272304371 | 0.0957 | 2.8462 | 0.00442 | 0.0358 |
| ENSG00000225177.5 | 127.0112 | 0.538188943 | 0.1891 | 2.8466 | 0.00442 | 0.0358 |
| ENSG00000277531.2 | 1874.613 | -0.270772454 | 0.0951 | -2.846 | 0.00443 | 0.0358 |
| ENSG00000138698.14 | 2755.392 | -0.308703116 | 0.1085 | -2.845 | 0.00444 | 0.0358 |
| ENSG00000282160.1 | 12.25399 | 0.584708504 | 0.2055 | 2.845 | 0.00444 | 0.0358 |
| ENSG00000132872.11 | 5562.578 | -0.446348345 | 0.1569 | -2.844 | 0.00445 | 0.0358 |
| ENSG00000144909.7 | 439.3541 | 0.273556345 | 0.0962 | 2.8438 | 0.00446 | 0.0359 |
| ENSG00000164924.17 | 30993.34 | -0.258852505 | 0.091 | -2.844 | 0.00446 | 0.0359 |
| ENSG00000101004.14 | 318.0116 | 0.246570414 | 0.0867 | 2.8425 | 0.00448 | 0.0359 |
| ENSG00000147416.10 | 10900.34 | -0.31578025 | 0.1111 | -2.843 | 0.00447 | 0.0359 |
| ENSG00000162456.9 | 70.39671 | -0.364904913 | 0.1284 | -2.842 | 0.00448 | 0.0359 |
| ENSG00000172728.15 | 226.4842 | 0.362077509 | 0.1274 | 2.8419 | 0.00448 | 0.0359 |
| ENSG00000188859.6 | 426.3147 | -0.274655521 | 0.0966 | -2.842 | 0.00448 | 0.0359 |
| ENSG00000230461.8 | 75.17554 | 0.245926156 | 0.0865 | 2.8421 | 0.00448 | 0.0359 |
| ENSG00000147872.9 | 168.5238 | 0.522798368 | 0.184 | 2.8406 | 0.0045 | 0.036 |
| ENSG00000257572.1 | 23.72957 | 0.452902151 | 0.1595 | 2.8398 | 0.00451 | 0.0361 |
| ENSG00000154144.12 | 2501.365 | -0.276991438 | 0.0976 | -2.839 | 0.00452 | 0.0361 |
| ENSG00000070193.4 | 34.74407 | -0.560200373 | 0.1974 | -2.838 | 0.00454 | 0.0362 |
| ENSG00000109066.13 | 300.787 | 0.271250123 | 0.0956 | 2.8382 | 0.00454 | 0.0362 |
| ENSG00000204406.12 | 849.0184 | -0.179069641 | 0.0631 | -2.838 | 0.00454 | 0.0362 |
| ENSG00000232043.1 | 27.04968 | -0.439362192 | 0.1548 | -2.837 | 0.00455 | 0.0362 |
| ENSG00000166106.3 | 43.08653 | -0.537724612 | 0.1896 | -2.837 | 0.00456 | 0.0363 |
| ENSG00000188039.13 | 557.5419 | 0.582080519 | 0.2053 | 2.8357 | 0.00457 | 0.0364 |
| ENSG00000172575.11 | 1588.11 | -0.512944057 | 0.181 | -2.834 | 0.00459 | 0.0364 |
| ENSG00000185352.8 | 3129.53 | -0.345179522 | 0.1218 | -2.835 | 0.00459 | 0.0364 |
| ENSG00000227141.2 | 15.24569 | -0.524446946 | 0.185 | -2.834 | 0.00459 | 0.0364 |
| ENSG00000197375.12 | 332.8003 | 0.227466643 | 0.0803 | 2.8338 | 0.0046 | 0.0365 |
| ENSG00000157335.20 | 17.00826 | 0.743135647 | 0.2623 | 2.8332 | 0.00461 | 0.0365 |
| ENSG00000106733.20 | 454.0894 | -0.274784396 | 0.097 | -2.833 | 0.00461 | 0.0365 |
| ENSG00000132470.13 | 537.8449 | 0.649716331 | 0.2294 | 2.8326 | 0.00462 | 0.0365 |
| ENSG00000162877.12 | 23.53562 | 0.742527888 | 0.2622 | 2.8317 | 0.00463 | 0.0366 |
| ENSG00000182400.14 | 2344.847 | -0.294185086 | 0.1039 | -2.831 | 0.00463 | 0.0366 |
| ENSG00000150051.13 | 187.5076 | -0.346137297 | 0.1223 | -2.83 | 0.00465 | 0.0367 |
| ENSG00000118849.9 | 49.94546 | -0.262795843 | 0.0929 | -2.829 | 0.00467 | 0.0368 |
| ENSG00000204588.5 | 1117.28 | -0.524562546 | 0.1855 | -2.829 | 0.00468 | 0.0368 |
| ENSG00000276043.4 | 70.09377 | 0.30482848 | 0.1078 | 2.8285 | 0.00468 | 0.0368 |
| ENSG00000078140.13 | 3847.818 | -0.278451928 | 0.0985 | -2.828 | 0.00468 | 0.0368 |
| ENSG00000163362.10 | 84.19338 | 0.375220025 | 0.1328 | 2.8251 | 0.00473 | 0.0372 |
| ENSG00000181754.6 | 1849.544 | -0.28045685 | 0.0993 | -2.825 | 0.00473 | 0.0372 |
| ENSG00000115252.18 | 2291.52 | -0.329712402 | 0.1167 | -2.824 | 0.00474 | 0.0372 |
| ENSG00000177098.8 | 1736.749 | -0.305717738 | 0.1083 | -2.824 | 0.00475 | 0.0372 |
| ENSG00000003987.13 | 1062.603 | -0.303303091 | 0.1075 | -2.822 | 0.00478 | 0.0374 |
| ENSG00000156687.10 | 1389.777 | -0.340561023 | 0.1207 | -2.821 | 0.00478 | 0.0374 |
| ENSG00000160307.9 | 3126.551 | 0.420429598 | 0.149 | 2.8214 | 0.00478 | 0.0374 |
| ENSG00000163053.10 | 789.748 | -0.282962204 | 0.1003 | -2.822 | 0.00478 | 0.0374 |
| ENSG00000279123.1 | 8.960787 | 0.679488495 | 0.2408 | 2.8218 | 0.00478 | 0.0374 |
| ENSG00000079691.17 | 823.3993 | 0.35577862 | 0.1261 | 2.8205 | 0.00479 | 0.0374 |
| ENSG00000184368.15 | 2566.71 | -0.317664797 | 0.1127 | -2.82 | 0.0048 | 0.0375 |
| ENSG00000129250.11 | 3379.12 | 0.33649177 | 0.1193 | 2.8195 | 0.00481 | 0.0375 |
| ENSG00000159164.9 | 9421.495 | -0.267677302 | 0.0949 | -2.819 | 0.00481 | 0.0375 |
| ENSG00000091137.11 | 486.506 | -0.276608221 | 0.0981 | -2.819 | 0.00482 | 0.0375 |
| ENSG00000184305.14 | 153.6653 | -0.252415091 | 0.0896 | -2.818 | 0.00483 | 0.0375 |
| ENSG00000105355.8 | 231.7836 | 0.465412662 | 0.1652 | 2.818 | 0.00483 | 0.0376 |
| ENSG00000104237.7 | 11.13738 | 0.464833504 | 0.165 | 2.8167 | 0.00485 | 0.0377 |
| ENSG00000118432.12 | 3332.445 | -0.32448867 | 0.1152 | -2.817 | 0.00485 | 0.0377 |
| ENSG00000070882.12 | 833.8614 | -0.281315012 | 0.0999 | -2.815 | 0.00488 | 0.0378 |
| ENSG00000136040.8 | 1618.457 | -0.285137878 | 0.1013 | -2.815 | 0.00488 | 0.0378 |
| ENSG00000178685.13 | 337.8502 | 0.315065673 | 0.1119 | 2.8148 | 0.00488 | 0.0378 |
| ENSG00000072041.16 | 1715.358 | -0.343325863 | 0.122 | -2.813 | 0.00491 | 0.038 |
| ENSG00000109738.10 | 2352.204 | -0.314490893 | 0.1118 | -2.812 | 0.00492 | 0.038 |
| ENSG00000054803.3 | 948.4981 | -0.315095565 | 0.1121 | -2.812 | 0.00493 | 0.038 |
| ENSG00000126950.7 | 1105.439 | -0.322538987 | 0.1147 | -2.812 | 0.00492 | 0.038 |
| ENSG00000260822.1 | 1891.33 | -0.361579024 | 0.1286 | -2.812 | 0.00493 | 0.038 |
| ENSG00000008517.16 | 78.23709 | 0.501657417 | 0.1787 | 2.8076 | 0.00499 | 0.0382 |
| ENSG00000100075.9 | 550.1583 | 0.248089255 | 0.0884 | 2.8077 | 0.00499 | 0.0382 |
| ENSG00000101349.16 | 998.2639 | -0.29762295 | 0.1059 | -2.81 | 0.00496 | 0.0382 |
| ENSG00000113319.11 | 3914.982 | -0.388699929 | 0.1384 | -2.808 | 0.00498 | 0.0382 |
| ENSG00000114446.4 | 743.1536 | -0.187196322 | 0.0666 | -2.809 | 0.00496 | 0.0382 |
| ENSG00000141668.9 | 1938.073 | -0.351141504 | 0.1251 | -2.808 | 0.00499 | 0.0382 |
| ENSG00000157502.13 | 484.9694 | -0.438210522 | 0.156 | -2.809 | 0.00497 | 0.0382 |
| ENSG00000175600.15 | 33.07025 | 0.432848974 | 0.154 | 2.81 | 0.00495 | 0.0382 |
| ENSG00000229206.3 | 49.34497 | 0.665791119 | 0.2371 | 2.8085 | 0.00498 | 0.0382 |
| ENSG00000245573.7 | 146.1848 | 0.225308222 | 0.0802 | 2.8085 | 0.00498 | 0.0382 |
| ENSG00000270112.3 | 100.3133 | -0.378247551 | 0.1347 | -2.808 | 0.00498 | 0.0382 |
| ENSG00000280224.1 | 27.72758 | 0.438974911 | 0.1562 | 2.8095 | 0.00496 | 0.0382 |
| ENSG00000050748.17 | 5754.877 | -0.295388237 | 0.1052 | -2.807 | 0.005 | 0.0382 |
| ENSG00000172292.14 | 2648.736 | -0.39491684 | 0.1407 | -2.806 | 0.00502 | 0.0382 |
| ENSG00000183688.4 | 232.8734 | 0.317001126 | 0.113 | 2.806 | 0.00502 | 0.0382 |
| ENSG00000231826.5 | 24.46868 | 0.52486516 | 0.187 | 2.8061 | 0.00501 | 0.0382 |
| ENSG00000277586.1 | 16833.76 | -0.412313235 | 0.1469 | -2.806 | 0.00501 | 0.0382 |
| ENSG00000176749.8 | 7968.684 | -0.258996538 | 0.0923 | -2.805 | 0.00503 | 0.0382 |
| ENSG00000133424.20 | 2942.537 | -0.3328543 | 0.1187 | -2.805 | 0.00503 | 0.0383 |
| ENSG00000177511.5 | 6095.927 | -0.345869641 | 0.1233 | -2.805 | 0.00504 | 0.0383 |
| ENSG00000064692.18 | 135.8383 | 0.278725925 | 0.0994 | 2.8038 | 0.00505 | 0.0383 |
| ENSG00000104415.13 | 65.16243 | -0.678099405 | 0.2419 | -2.804 | 0.00505 | 0.0383 |
| ENSG00000186654.20 | 431.9294 | 0.265204456 | 0.0946 | 2.8039 | 0.00505 | 0.0383 |
| ENSG00000139970.16 | 21988.94 | -0.317840317 | 0.1134 | -2.803 | 0.00506 | 0.0383 |
| ENSG00000115556.13 | 185.3125 | 0.320871593 | 0.1145 | 2.803 | 0.00506 | 0.0383 |
| ENSG00000159784.17 | 3154.759 | -0.32567458 | 0.1163 | -2.801 | 0.00509 | 0.0385 |
| ENSG00000228109.1 | 116.2364 | 0.246051211 | 0.0879 | 2.7999 | 0.00511 | 0.0386 |
| ENSG00000132704.15 | 32.40795 | 0.345348559 | 0.1234 | 2.7988 | 0.00513 | 0.0387 |
| ENSG00000142149.8 | 391.5962 | -0.299839851 | 0.1071 | -2.799 | 0.00513 | 0.0387 |
| ENSG00000197299.10 | 156.3641 | 0.254403524 | 0.0909 | 2.7991 | 0.00512 | 0.0387 |
| ENSG00000186212.3 | 332.3475 | -0.497359528 | 0.1778 | -2.798 | 0.00514 | 0.0388 |
| ENSG00000110931.18 | 8625.961 | -0.317916866 | 0.1137 | -2.796 | 0.00518 | 0.039 |
| ENSG00000145358.6 | 101.9925 | 0.583390361 | 0.2087 | 2.7952 | 0.00519 | 0.039 |
| ENSG00000170500.12 | 7699.679 | -0.264411212 | 0.0946 | -2.795 | 0.00519 | 0.039 |
| ENSG00000100744.14 | 938.7115 | -0.268443822 | 0.0961 | -2.793 | 0.00522 | 0.0391 |
| ENSG00000104611.11 | 44.1794 | 0.362987022 | 0.13 | 2.793 | 0.00522 | 0.0391 |
| ENSG00000155090.14 | 603.8539 | -0.347928751 | 0.1246 | -2.793 | 0.00522 | 0.0391 |
| ENSG00000161860.7 | 19.73163 | 0.419521407 | 0.1502 | 2.7926 | 0.00523 | 0.0391 |
| ENSG00000171522.5 | 65.11073 | -0.509317177 | 0.1824 | -2.793 | 0.00523 | 0.0391 |
| ENSG00000175175.5 | 1351.693 | -0.379942323 | 0.1361 | -2.793 | 0.00523 | 0.0391 |
| ENSG00000213853.9 | 512.3013 | 0.303782771 | 0.1087 | 2.7934 | 0.00522 | 0.0391 |
| ENSG00000187730.7 | 2203.259 | -0.314363823 | 0.1127 | -2.791 | 0.00526 | 0.0393 |
| ENSG00000187824.8 | 127.4857 | 0.307653512 | 0.1103 | 2.7901 | 0.00527 | 0.0394 |
| ENSG00000010282.14 | 845.2744 | 0.386883106 | 0.1387 | 2.7895 | 0.00528 | 0.0394 |
| ENSG00000114573.9 | 9204.563 | -0.362450215 | 0.13 | -2.789 | 0.00529 | 0.0395 |
| ENSG00000196277.15 | 575.4563 | -0.249666506 | 0.0895 | -2.789 | 0.00529 | 0.0395 |
| ENSG00000078328.19 | 6504.403 | -0.280918231 | 0.1008 | -2.787 | 0.00531 | 0.0395 |
| ENSG00000119401.10 | 419.5034 | -0.313419206 | 0.1124 | -2.788 | 0.00531 | 0.0395 |
| ENSG00000164114.18 | 1629.672 | -0.286380303 | 0.1027 | -2.788 | 0.00531 | 0.0395 |
| ENSG00000067842.17 | 2015.591 | -0.330937714 | 0.1187 | -2.787 | 0.00532 | 0.0395 |
| ENSG00000155368.16 | 1281.678 | 0.33523459 | 0.1203 | 2.7868 | 0.00532 | 0.0395 |
| ENSG00000096654.15 | 499.9186 | -0.279708622 | 0.1004 | -2.785 | 0.00535 | 0.0396 |
| ENSG00000155858.5 | 918.9062 | -0.28444205 | 0.1021 | -2.786 | 0.00534 | 0.0396 |
| ENSG00000164061.4 | 9979.11 | -0.394751201 | 0.1417 | -2.786 | 0.00534 | 0.0396 |
| ENSG00000255085.8 | 25.17758 | 0.407170923 | 0.1462 | 2.7853 | 0.00535 | 0.0396 |
| ENSG00000164841.4 | 90.77258 | -0.34436501 | 0.1237 | -2.785 | 0.00536 | 0.0396 |
| ENSG00000267056.2 | 19.05727 | 0.783891222 | 0.2815 | 2.7847 | 0.00536 | 0.0396 |
| ENSG00000237624.1 | 15.68698 | -0.534036878 | 0.1918 | -2.784 | 0.00536 | 0.0396 |
| ENSG00000261115.5 | 2623.736 | -0.336504324 | 0.1209 | -2.784 | 0.00537 | 0.0397 |
| ENSG00000144749.13 | 2538.995 | 0.502653074 | 0.1806 | 2.7833 | 0.00538 | 0.0397 |
| ENSG00000140945.16 | 3223.874 | -0.334610052 | 0.1203 | -2.782 | 0.0054 | 0.0398 |
| ENSG00000117600.12 | 5031.322 | -0.336450345 | 0.121 | -2.782 | 0.00541 | 0.0399 |
| ENSG00000073910.19 | 2609.334 | -0.27374321 | 0.0984 | -2.781 | 0.00543 | 0.0399 |
| ENSG00000077157.21 | 3564.972 | -0.287531345 | 0.1034 | -2.781 | 0.00542 | 0.0399 |
| ENSG00000148175.12 | 2201.294 | 0.462286817 | 0.1662 | 2.7811 | 0.00542 | 0.0399 |
| ENSG00000205710.3 | 196.3858 | -0.349483628 | 0.1257 | -2.781 | 0.00542 | 0.0399 |
| ENSG00000124067.16 | 455.1266 | 0.379651459 | 0.1366 | 2.7794 | 0.00545 | 0.0399 |
| ENSG00000129636.12 | 3837.704 | -0.320485446 | 0.1153 | -2.78 | 0.00544 | 0.0399 |
| ENSG00000204264.8 | 130.5336 | 0.32238037 | 0.116 | 2.7797 | 0.00544 | 0.0399 |
| ENSG00000233123.1 | 577.2406 | -0.453847973 | 0.1633 | -2.78 | 0.00544 | 0.0399 |
| ENSG00000274211.4 | 2102.891 | -0.254200739 | 0.0915 | -2.779 | 0.00545 | 0.0399 |
| ENSG00000068912.13 | 1993.595 | -0.265208488 | 0.0955 | -2.777 | 0.00549 | 0.0401 |
| ENSG00000078053.16 | 3496.077 | -0.379255548 | 0.1366 | -2.777 | 0.00548 | 0.0401 |
| ENSG00000082458.11 | 2821.667 | -0.328970741 | 0.1185 | -2.777 | 0.00549 | 0.0401 |
| ENSG00000119508.17 | 687.2884 | -0.469166682 | 0.1689 | -2.777 | 0.00548 | 0.0401 |
| ENSG00000162378.12 | 4031.297 | -0.313869519 | 0.113 | -2.778 | 0.00548 | 0.0401 |
| ENSG00000164306.10 | 165.9791 | 0.209192569 | 0.0753 | 2.7767 | 0.00549 | 0.0401 |
| ENSG00000149948.13 | 11.8454 | 0.462748161 | 0.1667 | 2.7754 | 0.00551 | 0.0401 |
| ENSG00000238123.1 | 11.01168 | 0.568840727 | 0.2049 | 2.7756 | 0.00551 | 0.0401 |
| ENSG00000170832.12 | 1784.286 | -0.349275623 | 0.1259 | -2.775 | 0.00552 | 0.0401 |
| ENSG00000196935.8 | 606.4811 | 0.255990949 | 0.0923 | 2.7748 | 0.00552 | 0.0402 |
| ENSG00000011677.12 | 1948.004 | -0.338385374 | 0.122 | -2.774 | 0.00553 | 0.0402 |
| ENSG00000263264.1 | 18.63969 | 0.465348761 | 0.1678 | 2.7736 | 0.00554 | 0.0403 |
| ENSG00000152952.11 | 543.821 | 0.391425232 | 0.1412 | 2.7727 | 0.00556 | 0.0403 |
| ENSG00000171132.13 | 4329.714 | -0.345259885 | 0.1245 | -2.773 | 0.00555 | 0.0403 |
| ENSG00000186260.16 | 8014.36 | -0.302872014 | 0.1092 | -2.773 | 0.00556 | 0.0403 |
| ENSG00000136297.14 | 677.139 | 0.405936436 | 0.1464 | 2.7724 | 0.00557 | 0.0403 |
| ENSG00000168268.10 | 270.0501 | 0.241971437 | 0.0873 | 2.7705 | 0.0056 | 0.0405 |
| ENSG00000123329.17 | 105.7023 | -0.407516192 | 0.1471 | -2.77 | 0.00561 | 0.0406 |
| ENSG00000162909.17 | 1830.994 | 0.25652209 | 0.0926 | 2.7692 | 0.00562 | 0.0406 |
| ENSG00000139155.8 | 742.5106 | 0.606623673 | 0.2191 | 2.7686 | 0.00563 | 0.0406 |
| ENSG00000142632.16 | 22.13775 | 0.449434065 | 0.1623 | 2.7685 | 0.00563 | 0.0406 |
| ENSG00000162852.13 | 1731.598 | -0.302581036 | 0.1093 | -2.768 | 0.00564 | 0.0407 |
| ENSG00000112280.15 | 382.6672 | 0.316328084 | 0.1143 | 2.7677 | 0.00565 | 0.0407 |
| ENSG00000265015.1 | 9.952432 | -0.586978492 | 0.2121 | -2.767 | 0.00565 | 0.0407 |
| ENSG00000173064.11 | 5843.883 | -0.302751455 | 0.1094 | -2.766 | 0.00567 | 0.0408 |
| ENSG00000197860.9 | 3665.261 | -0.289157644 | 0.1045 | -2.766 | 0.00567 | 0.0408 |
| ENSG00000169918.9 | 1654.152 | -0.202770384 | 0.0733 | -2.765 | 0.0057 | 0.0409 |
| ENSG00000211896.7 | 19.10356 | -1.322391834 | 0.4784 | -2.764 | 0.0057 | 0.0409 |
| ENSG00000173262.11 | 14.77577 | -0.430975785 | 0.1559 | -2.764 | 0.00571 | 0.041 |
| ENSG00000058091.16 | 3623.653 | -0.348983638 | 0.1264 | -2.761 | 0.00576 | 0.041 |
| ENSG00000107643.15 | 1770.883 | -0.25684848 | 0.093 | -2.763 | 0.00573 | 0.041 |
| ENSG00000111885.6 | 1324.077 | -0.340595829 | 0.1233 | -2.762 | 0.00575 | 0.041 |
| ENSG00000134115.12 | 362.9923 | -0.40619138 | 0.1471 | -2.762 | 0.00575 | 0.041 |
| ENSG00000143036.16 | 110.2302 | 0.511162538 | 0.1851 | 2.7622 | 0.00574 | 0.041 |
| ENSG00000152931.7 | 1241.629 | -0.478521777 | 0.1733 | -2.761 | 0.00576 | 0.041 |
| ENSG00000170684.8 | 39.20598 | 0.355453559 | 0.1287 | 2.7613 | 0.00576 | 0.041 |
| ENSG00000180611.6 | 267.9036 | -0.316421134 | 0.1146 | -2.762 | 0.00574 | 0.041 |
| ENSG00000186094.16 | 341.1279 | -0.32244238 | 0.1167 | -2.763 | 0.00573 | 0.041 |
| ENSG00000233452.6 | 400.2523 | -0.315056676 | 0.1141 | -2.762 | 0.00575 | 0.041 |
| ENSG00000276204.1 | 10.07987 | 0.56363882 | 0.204 | 2.7632 | 0.00572 | 0.041 |
| ENSG00000183196.8 | 282.9767 | 0.312577861 | 0.1133 | 2.7597 | 0.00578 | 0.0412 |
| ENSG00000154027.18 | 9911.635 | -0.327782115 | 0.1188 | -2.759 | 0.00579 | 0.0412 |
| ENSG00000074590.13 | 4404.485 | -0.384632801 | 0.1394 | -2.759 | 0.00581 | 0.0412 |
| ENSG00000155307.17 | 50.6888 | -0.759689906 | 0.2755 | -2.757 | 0.00583 | 0.0413 |
| ENSG00000158987.20 | 918.3903 | -0.196309756 | 0.0712 | -2.757 | 0.00583 | 0.0413 |
| ENSG00000260260.1 | 112.1894 | 0.309358797 | 0.1122 | 2.7576 | 0.00582 | 0.0413 |
| ENSG00000147724.11 | 774.0251 | -0.272161109 | 0.0987 | -2.757 | 0.00584 | 0.0414 |
| ENSG00000185924.6 | 943.2112 | -0.314757568 | 0.1142 | -2.756 | 0.00584 | 0.0414 |
| ENSG00000214043.7 | 44.84882 | -0.445059517 | 0.1615 | -2.756 | 0.00585 | 0.0414 |
| ENSG00000225210.9 | 47.18985 | -0.446196466 | 0.1619 | -2.755 | 0.00586 | 0.0415 |
| ENSG00000123684.12 | 3006.778 | -0.281746916 | 0.1023 | -2.755 | 0.00587 | 0.0415 |
| ENSG00000182508.13 | 42.31101 | 0.663092406 | 0.2407 | 2.7546 | 0.00588 | 0.0415 |
| ENSG00000125827.8 | 4161.937 | -0.290252066 | 0.1054 | -2.754 | 0.00589 | 0.0415 |
| ENSG00000151458.11 | 999.9703 | -0.260453468 | 0.0946 | -2.753 | 0.0059 | 0.0416 |
| ENSG00000115365.11 | 5788.546 | -0.29384937 | 0.1068 | -2.752 | 0.00593 | 0.0417 |
| ENSG00000173402.11 | 1427.157 | 0.237001508 | 0.0861 | 2.7513 | 0.00594 | 0.0417 |
| ENSG00000185291.11 | 94.69521 | 0.242232571 | 0.088 | 2.7519 | 0.00593 | 0.0417 |
| ENSG00000185291.11_PAR_Y | 94.69521 | 0.242232571 | 0.088 | 2.7519 | 0.00593 | 0.0417 |
| ENSG00000274922.1 | 16.18074 | 0.46096815 | 0.1675 | 2.7515 | 0.00593 | 0.0417 |
| ENSG00000157064.10 | 5510.539 | -0.351100422 | 0.1276 | -2.751 | 0.00595 | 0.0417 |
| ENSG00000174151.14 | 569.0612 | -0.239871866 | 0.0872 | -2.751 | 0.00594 | 0.0417 |
| ENSG00000244053.1 | 15.30157 | 0.51719677 | 0.188 | 2.7507 | 0.00595 | 0.0417 |
| ENSG00000147676.13 | 2691.81 | -0.342076254 | 0.1244 | -2.75 | 0.00596 | 0.0417 |
| ENSG00000107518.16 | 4810.946 | -0.355092499 | 0.1292 | -2.749 | 0.00597 | 0.0418 |
| ENSG00000107611.14 | 107.8701 | 0.286253449 | 0.1041 | 2.7494 | 0.00597 | 0.0418 |
| ENSG00000279364.1 | 513.2679 | 0.28798314 | 0.1048 | 2.749 | 0.00598 | 0.0418 |
| ENSG00000178904.18 | 1049.809 | 0.314935999 | 0.1146 | 2.7486 | 0.00599 | 0.0418 |
| ENSG00000253719.3 | 5020.182 | -0.272051228 | 0.099 | -2.748 | 0.00599 | 0.0418 |
| ENSG00000176046.8 | 413.2172 | 0.451801255 | 0.1644 | 2.748 | 0.006 | 0.0419 |
| ENSG00000134121.9 | 9984.834 | -0.338978659 | 0.1234 | -2.747 | 0.00601 | 0.0419 |
| ENSG00000149970.14 | 3933.63 | -0.270637004 | 0.0985 | -2.747 | 0.00601 | 0.0419 |
| ENSG00000204186.7 | 1702.948 | -0.266231966 | 0.0969 | -2.747 | 0.00601 | 0.0419 |
| ENSG00000149452.15 | 28.28743 | 0.453682356 | 0.1653 | 2.7454 | 0.00604 | 0.0421 |
| ENSG00000225963.7 | 198.2012 | -0.269684221 | 0.0983 | -2.745 | 0.00606 | 0.0421 |
| ENSG00000019102.11 | 61.32666 | -0.335742366 | 0.1223 | -2.744 | 0.00606 | 0.0421 |
| ENSG00000150768.15 | 1241.77 | -0.22367411 | 0.0815 | -2.744 | 0.00606 | 0.0421 |
| ENSG00000240590.1 | 454.6793 | 1.326889102 | 0.4837 | 2.7434 | 0.00608 | 0.0422 |
| ENSG00000205364.3 | 137.2433 | 0.688223906 | 0.251 | 2.7424 | 0.0061 | 0.0423 |
| ENSG00000110427.14 | 6862.421 | -0.339086102 | 0.1237 | -2.742 | 0.00612 | 0.0424 |
| ENSG00000110841.13 | 2145.531 | -0.221790334 | 0.0809 | -2.741 | 0.00612 | 0.0424 |
| ENSG00000157734.13 | 509.5747 | 0.27478138 | 0.1002 | 2.7411 | 0.00612 | 0.0424 |
| ENSG00000258932.5 | 26.46309 | 0.44110106 | 0.1609 | 2.7412 | 0.00612 | 0.0424 |
| ENSG00000133083.14 | 10588.37 | -0.395938556 | 0.1445 | -2.74 | 0.00614 | 0.0424 |
| ENSG00000277734.6 | 34.05832 | -0.641194994 | 0.234 | -2.74 | 0.00615 | 0.0425 |
| ENSG00000105929.15 | 18.86694 | 0.489443897 | 0.1787 | 2.7389 | 0.00616 | 0.0425 |
| ENSG00000271127.1 | 27.41451 | -0.630069954 | 0.23 | -2.739 | 0.00616 | 0.0425 |
| ENSG00000184347.14 | 1269.534 | -0.327377721 | 0.1195 | -2.739 | 0.00617 | 0.0426 |
| ENSG00000132967.9 | 62.47657 | -1.114175715 | 0.4069 | -2.738 | 0.00618 | 0.0426 |
| ENSG00000154429.10 | 3070.061 | -0.263645593 | 0.0963 | -2.737 | 0.0062 | 0.0427 |
| ENSG00000051382.8 | 1833.167 | -0.305944748 | 0.1118 | -2.736 | 0.00621 | 0.0427 |
| ENSG00000156345.17 | 755.7323 | -0.272020602 | 0.0994 | -2.736 | 0.00621 | 0.0427 |
| ENSG00000021645.18 | 4651.582 | -0.268608677 | 0.0982 | -2.736 | 0.00622 | 0.0428 |
| ENSG00000224877.3 | 505.0478 | 0.272632412 | 0.0997 | 2.7352 | 0.00623 | 0.0428 |
| ENSG00000067715.13 | 34759.88 | -0.375693548 | 0.1374 | -2.734 | 0.00626 | 0.0429 |
| ENSG00000163536.12 | 9189.729 | -0.307365538 | 0.1124 | -2.734 | 0.00625 | 0.0429 |
| ENSG00000178764.7 | 460.4856 | 0.370874858 | 0.1357 | 2.7341 | 0.00626 | 0.0429 |
| ENSG00000152495.10 | 3634.624 | -0.381403258 | 0.1395 | -2.733 | 0.00627 | 0.0429 |
| ENSG00000107105.14 | 2183.801 | -0.306804729 | 0.1123 | -2.733 | 0.00627 | 0.0429 |
| ENSG00000111674.8 | 17104.51 | -0.341109068 | 0.1249 | -2.732 | 0.0063 | 0.0431 |
| ENSG00000164211.12 | 1084.99 | -0.286555924 | 0.1049 | -2.731 | 0.00632 | 0.0432 |
| ENSG00000188636.3 | 2097.428 | -0.266418947 | 0.0976 | -2.731 | 0.00632 | 0.0432 |
| ENSG00000102452.15 | 1948.76 | -0.285772478 | 0.1047 | -2.73 | 0.00634 | 0.0432 |
| ENSG00000198908.11 | 1203.482 | -0.302042531 | 0.1106 | -2.73 | 0.00634 | 0.0432 |
| ENSG00000125871.13 | 113.3821 | 0.253259853 | 0.0928 | 2.7289 | 0.00636 | 0.0433 |
| ENSG00000196743.8 | 996.0916 | 0.236952625 | 0.0869 | 2.7282 | 0.00637 | 0.0434 |
| ENSG00000212719.10 | 1532.966 | -0.287087771 | 0.1053 | -2.727 | 0.00638 | 0.0435 |
| ENSG00000243444.7 | 615.6836 | -0.376518345 | 0.1381 | -2.727 | 0.0064 | 0.0435 |
| ENSG00000133627.17 | 1752.916 | -0.266016454 | 0.0976 | -2.725 | 0.00642 | 0.0437 |
| ENSG00000142875.19 | 12491.82 | -0.306386939 | 0.1124 | -2.725 | 0.00644 | 0.0437 |
| ENSG00000103852.12 | 178.9391 | 0.19366643 | 0.0711 | 2.724 | 0.00645 | 0.0438 |
| ENSG00000144407.9 | 108.9301 | -0.372096326 | 0.1366 | -2.724 | 0.00646 | 0.0438 |
| ENSG00000158270.11 | 428.7611 | 0.361834979 | 0.1329 | 2.7231 | 0.00647 | 0.0438 |
| ENSG00000175344.17 | 386.3303 | -0.276088799 | 0.1014 | -2.723 | 0.00646 | 0.0438 |
| ENSG00000247363.2 | 12.09292 | 0.507052653 | 0.1862 | 2.7233 | 0.00646 | 0.0438 |
| ENSG00000065609.14 | 9074.312 | -0.298744915 | 0.1097 | -2.723 | 0.00648 | 0.0438 |
| ENSG00000154162.14 | 340.2714 | -0.337476762 | 0.124 | -2.723 | 0.00648 | 0.0438 |
| ENSG00000152784.15 | 1430.776 | -0.266043735 | 0.0977 | -2.722 | 0.00649 | 0.0439 |
| ENSG00000232284.7 | 50.25524 | 0.300728381 | 0.1105 | 2.7212 | 0.00651 | 0.0439 |
| ENSG00000272321.1 | 250.3997 | -0.297801273 | 0.1094 | -2.721 | 0.00651 | 0.0439 |
| ENSG00000166342.18 | 1112.795 | -0.294723234 | 0.1083 | -2.72 | 0.00652 | 0.044 |
| ENSG00000140488.15 | 869.0299 | -0.293691382 | 0.108 | -2.72 | 0.00653 | 0.0441 |
| ENSG00000170915.8 | 2635.787 | 0.37807144 | 0.1391 | 2.7182 | 0.00656 | 0.0442 |
| ENSG00000185900.9 | 233.5897 | -0.28671201 | 0.1055 | -2.718 | 0.00656 | 0.0442 |
| ENSG00000256085.1 | 16.54514 | -0.485163941 | 0.1785 | -2.718 | 0.00657 | 0.0442 |
| ENSG00000145147.19 | 1343.475 | -0.305334025 | 0.1123 | -2.718 | 0.00657 | 0.0442 |
| ENSG00000165886.4 | 223.7135 | 0.283171541 | 0.1042 | 2.7173 | 0.00658 | 0.0442 |
| ENSG00000177181.14 | 1427.102 | -0.323799816 | 0.1192 | -2.717 | 0.00659 | 0.0442 |
| ENSG00000172987.12 | 180.5192 | 0.462873205 | 0.1704 | 2.7165 | 0.0066 | 0.0443 |
| ENSG00000125780.11 | 23.90855 | -0.443215827 | 0.1632 | -2.716 | 0.00661 | 0.0443 |
| ENSG00000226203.1 | 58.42486 | -0.373637904 | 0.1376 | -2.716 | 0.00661 | 0.0443 |
| ENSG00000133318.13 | 31622.54 | -0.272281506 | 0.1003 | -2.715 | 0.00662 | 0.0444 |
| ENSG00000173726.10 | 9716.509 | -0.280916854 | 0.1035 | -2.715 | 0.00663 | 0.0444 |
| ENSG00000160685.13 | 265.5271 | 0.326924509 | 0.1204 | 2.7145 | 0.00664 | 0.0444 |
| ENSG00000115257.15 | 137.0502 | 0.311467758 | 0.1148 | 2.7141 | 0.00665 | 0.0444 |
| ENSG00000131094.3 | 567.5869 | 0.36910248 | 0.136 | 2.7137 | 0.00665 | 0.0444 |
| ENSG00000269699.5 | 125.8223 | -0.307878709 | 0.1135 | -2.713 | 0.00667 | 0.0445 |
| ENSG00000143473.11 | 721.8879 | -0.557217652 | 0.2054 | -2.712 | 0.00668 | 0.0446 |
| ENSG00000108947.4 | 2300.913 | -0.316446133 | 0.1167 | -2.712 | 0.00669 | 0.0446 |
| ENSG00000135926.13 | 977.6748 | 0.510325446 | 0.1882 | 2.7118 | 0.00669 | 0.0446 |
| ENSG00000198125.12 | 9.333601 | 0.62255402 | 0.2296 | 2.7116 | 0.0067 | 0.0446 |
| ENSG00000143847.15 | 3375.97 | -0.270210775 | 0.0997 | -2.71 | 0.00673 | 0.0447 |
| ENSG00000172366.19 | 494.5907 | 0.23628899 | 0.0872 | 2.7102 | 0.00672 | 0.0447 |
| ENSG00000111783.12 | 485.964 | 0.578533951 | 0.2135 | 2.7092 | 0.00674 | 0.0448 |
| ENSG00000143067.4 | 374.833 | -0.311397508 | 0.1149 | -2.709 | 0.00675 | 0.0448 |
| ENSG00000272163.1 | 515.3696 | -0.315136224 | 0.1163 | -2.709 | 0.00675 | 0.0448 |
| ENSG00000114646.9 | 2973.75 | 0.259913773 | 0.096 | 2.7085 | 0.00676 | 0.0448 |
| ENSG00000147224.10 | 1067.527 | -0.263898435 | 0.0975 | -2.707 | 0.00679 | 0.0449 |
| ENSG00000157152.16 | 13284.38 | -0.357646693 | 0.1321 | -2.707 | 0.00679 | 0.0449 |
| ENSG00000166111.9 | 4608.286 | -0.375387169 | 0.1387 | -2.707 | 0.00679 | 0.0449 |
| ENSG00000173598.13 | 6075.53 | -0.276429353 | 0.1021 | -2.707 | 0.00679 | 0.0449 |
| ENSG00000186288.5 | 392.3794 | -0.290505216 | 0.1073 | -2.707 | 0.0068 | 0.0449 |
| ENSG00000164116.16 | 787.1449 | -0.316525311 | 0.117 | -2.706 | 0.0068 | 0.0449 |
| ENSG00000144366.15 | 509.043 | -0.317688233 | 0.1174 | -2.706 | 0.00682 | 0.0449 |
| ENSG00000152503.9 | 1155.865 | -0.279838796 | 0.1034 | -2.706 | 0.00681 | 0.0449 |
| ENSG00000213397.10 | 552.3991 | 0.199478083 | 0.0737 | 2.7054 | 0.00682 | 0.0449 |
| ENSG00000246731.2 | 95.82545 | 0.44088364 | 0.1629 | 2.7058 | 0.00681 | 0.0449 |
| ENSG00000272579.1 | 309.6013 | 0.245422768 | 0.0907 | 2.7056 | 0.00682 | 0.0449 |
| ENSG00000077264.14 | 5140.839 | -0.30990909 | 0.1146 | -2.705 | 0.00683 | 0.045 |
| ENSG00000075945.12 | 3997.236 | -0.286450806 | 0.1059 | -2.704 | 0.00685 | 0.045 |
| ENSG00000118785.13 | 1139.543 | -0.668326043 | 0.2471 | -2.704 | 0.00684 | 0.045 |
| ENSG00000132639.12 | 55447.57 | -0.282888215 | 0.1046 | -2.705 | 0.00684 | 0.045 |
| ENSG00000180447.6 | 99.16855 | 0.33064336 | 0.1223 | 2.7041 | 0.00685 | 0.045 |
| ENSG00000204071.10 | 1064.281 | -0.250146806 | 0.0925 | -2.703 | 0.00686 | 0.045 |
| ENSG00000120306.9 | 705.0209 | 0.223305252 | 0.0826 | 2.7029 | 0.00687 | 0.0451 |
| ENSG00000119714.10 | 203.1641 | -0.29093464 | 0.1077 | -2.702 | 0.0069 | 0.0452 |
| ENSG00000186193.8 | 314.0314 | 0.323052027 | 0.1196 | 2.7003 | 0.00693 | 0.0454 |
| ENSG00000135709.12 | 12627.32 | -0.269871505 | 0.1 | -2.699 | 0.00695 | 0.0455 |
| ENSG00000139973.16 | 4547.226 | -0.315802188 | 0.117 | -2.699 | 0.00695 | 0.0455 |
| ENSG00000183715.13 | 4851.859 | -0.308899679 | 0.1146 | -2.696 | 0.00702 | 0.0459 |
| ENSG00000272556.2 | 76.4195 | 0.320441948 | 0.1189 | 2.6959 | 0.00702 | 0.0459 |
| ENSG00000182118.6 | 150.4933 | 0.364869655 | 0.1354 | 2.6956 | 0.00703 | 0.0459 |
| ENSG00000163818.16 | 1200.262 | -0.26811095 | 0.0995 | -2.694 | 0.00705 | 0.046 |
| ENSG00000144460.12 | 147.711 | -0.369295879 | 0.1371 | -2.694 | 0.00706 | 0.046 |
| ENSG00000166446.14 | 1250.462 | -0.292946799 | 0.1088 | -2.693 | 0.00708 | 0.0461 |
| ENSG00000172915.18 | 3759.684 | -0.381299516 | 0.1416 | -2.693 | 0.00708 | 0.0461 |
| ENSG00000185149.5 | 74.34968 | -0.374570231 | 0.1391 | -2.693 | 0.00708 | 0.0461 |
| ENSG00000151952.15 | 1091.864 | -0.374688798 | 0.1392 | -2.692 | 0.0071 | 0.0462 |
| ENSG00000102144.13 | 6874.463 | -0.29014674 | 0.1078 | -2.691 | 0.00713 | 0.0462 |
| ENSG00000106976.20 | 24308.23 | -0.246323448 | 0.0915 | -2.691 | 0.00712 | 0.0462 |
| ENSG00000140105.17 | 1968.845 | -0.361796185 | 0.1345 | -2.691 | 0.00713 | 0.0462 |
| ENSG00000163138.18 | 402.8943 | -0.18226531 | 0.0677 | -2.691 | 0.00713 | 0.0462 |
| ENSG00000174021.10 | 253.2051 | 0.372997937 | 0.1386 | 2.6903 | 0.00714 | 0.0462 |
| ENSG00000179406.6 | 154.8789 | 0.234890023 | 0.0873 | 2.6905 | 0.00714 | 0.0462 |
| ENSG00000243710.7 | 73.03173 | -0.267474916 | 0.0994 | -2.691 | 0.00713 | 0.0462 |
| ENSG00000051596.9 | 505.6049 | -0.365873521 | 0.136 | -2.689 | 0.00716 | 0.0462 |
| ENSG00000115194.10 | 1729.416 | -0.318499788 | 0.1184 | -2.69 | 0.00715 | 0.0462 |
| ENSG00000175946.8 | 12.98671 | -0.513922025 | 0.1911 | -2.689 | 0.00716 | 0.0462 |
| ENSG00000256443.1 | 98.51719 | -0.424463134 | 0.1578 | -2.69 | 0.00716 | 0.0462 |
| ENSG00000185650.9 | 2176.905 | 0.523973732 | 0.1949 | 2.6883 | 0.00718 | 0.0464 |
| ENSG00000188613.6 | 153.133 | 0.218712155 | 0.0814 | 2.688 | 0.00719 | 0.0464 |
| ENSG00000141469.16 | 349.3142 | 0.820644021 | 0.3054 | 2.6874 | 0.0072 | 0.0464 |
| ENSG00000088812.17 | 3522.594 | -0.265848838 | 0.099 | -2.686 | 0.00723 | 0.0465 |
| ENSG00000165806.19 | 146.513 | 0.50185206 | 0.1868 | 2.6865 | 0.00722 | 0.0465 |
| ENSG00000203668.2 | 956.676 | -0.304063986 | 0.1132 | -2.686 | 0.00723 | 0.0465 |
| ENSG00000187773.8 | 249.3106 | 0.399911151 | 0.1489 | 2.6855 | 0.00724 | 0.0465 |
| ENSG00000262098.1 | 15.3945 | -0.508743533 | 0.1894 | -2.686 | 0.00724 | 0.0465 |
| ENSG00000130540.13 | 6361.035 | -0.324183027 | 0.1207 | -2.685 | 0.00726 | 0.0466 |
| ENSG00000146701.11 | 3373.57 | -0.252928689 | 0.0942 | -2.684 | 0.00728 | 0.0467 |
| ENSG00000169836.4 | 29.19556 | -0.472270631 | 0.176 | -2.684 | 0.00728 | 0.0467 |
| ENSG00000177614.10 | 4311.318 | -0.245238289 | 0.0914 | -2.683 | 0.00729 | 0.0468 |
| ENSG00000111912.19 | 3522.734 | -0.37308032 | 0.1391 | -2.682 | 0.00731 | 0.0468 |
| ENSG00000149305.6 | 138.7741 | -0.381378818 | 0.1422 | -2.682 | 0.00732 | 0.0468 |
| ENSG00000248367.1 | 58.65052 | 0.288934288 | 0.1077 | 2.6819 | 0.00732 | 0.0468 |
| ENSG00000151229.12 | 4133.652 | -0.31637739 | 0.118 | -2.681 | 0.00734 | 0.0469 |
| ENSG00000069431.11 | 619.9108 | 0.273672771 | 0.1021 | 2.6805 | 0.00735 | 0.047 |
| ENSG00000084093.15 | 303.4063 | 0.325296866 | 0.1214 | 2.6805 | 0.00735 | 0.047 |
| ENSG00000123104.11 | 1101.153 | 0.388885958 | 0.1451 | 2.6798 | 0.00737 | 0.047 |
| ENSG00000233936.1 | 14.76384 | -0.530886161 | 0.1981 | -2.679 | 0.00737 | 0.047 |
| ENSG00000156206.13 | 12.08365 | -0.450347991 | 0.1681 | -2.679 | 0.00739 | 0.0471 |
| ENSG00000103353.15 | 1633.146 | -0.223184487 | 0.0833 | -2.678 | 0.0074 | 0.0472 |
| ENSG00000112159.11 | 2282.752 | -0.16068457 | 0.06 | -2.678 | 0.00742 | 0.0472 |
| ENSG00000130477.15 | 10312.73 | -0.3318181 | 0.124 | -2.677 | 0.00743 | 0.0472 |
| ENSG00000152284.4 | 148.7418 | 0.378867218 | 0.1415 | 2.6773 | 0.00742 | 0.0472 |
| ENSG00000163995.19 | 2039.714 | -0.236876624 | 0.0885 | -2.677 | 0.00744 | 0.0472 |
| ENSG00000166912.16 | 1338.358 | 0.334265178 | 0.1249 | 2.6771 | 0.00743 | 0.0472 |
| ENSG00000198961.9 | 15411.79 | -0.271871236 | 0.1016 | -2.677 | 0.00743 | 0.0472 |
| ENSG00000002746.14 | 1720.097 | -0.355377214 | 0.1328 | -2.676 | 0.00746 | 0.0473 |
| ENSG00000009844.15 | 1370.991 | -0.236015632 | 0.0882 | -2.676 | 0.00746 | 0.0473 |
| ENSG00000113732.8 | 423.2405 | 0.298532654 | 0.1116 | 2.6757 | 0.00746 | 0.0473 |
| ENSG00000176903.4 | 5873.103 | -0.265028175 | 0.0991 | -2.674 | 0.00748 | 0.0474 |
| ENSG00000185710.9 | 76.5176 | -0.672377853 | 0.2514 | -2.674 | 0.00749 | 0.0474 |
| ENSG00000163082.9 | 395.4726 | -0.284267671 | 0.1064 | -2.673 | 0.00752 | 0.0476 |
| ENSG00000144320.13 | 1581.21 | -0.236536781 | 0.0885 | -2.673 | 0.00753 | 0.0476 |
| ENSG00000085433.15 | 4212.381 | -0.293861154 | 0.11 | -2.672 | 0.00755 | 0.0477 |
| ENSG00000111249.13 | 1854.229 | -0.351331713 | 0.1315 | -2.672 | 0.00755 | 0.0477 |
| ENSG00000279107.1 | 42.45956 | 0.367351207 | 0.1375 | 2.6714 | 0.00755 | 0.0477 |
| ENSG00000047056.14 | 1376.607 | -0.266220258 | 0.0997 | -2.67 | 0.00757 | 0.0477 |
| ENSG00000109756.8 | 4199.882 | -0.242552669 | 0.0908 | -2.67 | 0.00759 | 0.0478 |
| ENSG00000145675.14 | 7147.27 | -0.256404014 | 0.096 | -2.67 | 0.00759 | 0.0478 |
| ENSG00000118308.15 | 295.9066 | -0.307837019 | 0.1153 | -2.669 | 0.0076 | 0.0478 |
| ENSG00000173546.7 | 725.9955 | 0.426013437 | 0.1596 | 2.669 | 0.00761 | 0.0478 |
| ENSG00000266573.5 | 11.86523 | -0.593136623 | 0.2222 | -2.669 | 0.00761 | 0.0478 |
| ENSG00000140497.16 | 373.5817 | 0.297253252 | 0.1114 | 2.6681 | 0.00763 | 0.0479 |
| ENSG00000254537.1 | 14.09865 | 0.467548082 | 0.1753 | 2.6679 | 0.00763 | 0.0479 |
| ENSG00000137941.16 | 4302.171 | -0.29093355 | 0.1091 | -2.666 | 0.00767 | 0.0481 |
| ENSG00000151376.16 | 1105.465 | -0.241380582 | 0.0905 | -2.666 | 0.00767 | 0.0481 |
| ENSG00000182310.13 | 209.918 | 0.223342547 | 0.0838 | 2.6663 | 0.00767 | 0.0481 |
| ENSG00000114757.18 | 4661.456 | -0.291421782 | 0.1093 | -2.665 | 0.00769 | 0.0481 |
| ENSG00000165832.5 | 1126.433 | -0.272115118 | 0.1021 | -2.665 | 0.00769 | 0.0481 |
| ENSG00000271461.1 | 19.38614 | 0.419657093 | 0.1574 | 2.6655 | 0.00769 | 0.0481 |
| ENSG00000198010.12 | 1824.9 | -0.24008408 | 0.0901 | -2.665 | 0.0077 | 0.0481 |
| ENSG00000159450.12 | 67.19294 | -0.292245665 | 0.1097 | -2.664 | 0.00771 | 0.0482 |
| ENSG00000105499.13 | 817.9771 | -0.253680088 | 0.0953 | -2.663 | 0.00775 | 0.0484 |
| ENSG00000258647.5 | 9.949155 | -0.788107816 | 0.296 | -2.663 | 0.00775 | 0.0484 |
| ENSG00000060140.8 | 446.4303 | -0.370869779 | 0.1393 | -2.662 | 0.00776 | 0.0484 |
| ENSG00000138078.15 | 16597.83 | -0.291965472 | 0.1097 | -2.662 | 0.00776 | 0.0484 |
| ENSG00000162783.10 | 812.6421 | -0.357543207 | 0.1343 | -2.661 | 0.00778 | 0.0485 |
| ENSG00000100030.14 | 11332.6 | -0.235499329 | 0.0885 | -2.661 | 0.0078 | 0.0485 |
| ENSG00000231189.1 | 35.38055 | 0.37559908 | 0.1412 | 2.6605 | 0.0078 | 0.0485 |
| ENSG00000112531.16 | 9168.272 | 0.359113294 | 0.135 | 2.6598 | 0.00782 | 0.0486 |
| ENSG00000147432.6 | 26.21482 | -0.436868629 | 0.1642 | -2.66 | 0.00782 | 0.0486 |
| ENSG00000034239.10 | 182.6303 | -0.470731074 | 0.177 | -2.659 | 0.00783 | 0.0486 |
| ENSG00000179455.7 | 79.53111 | 0.216474019 | 0.0814 | 2.6593 | 0.00783 | 0.0486 |
| ENSG00000084731.14 | 10349.76 | -0.351774325 | 0.1324 | -2.658 | 0.00786 | 0.0488 |
| ENSG00000109158.10 | 2718.295 | -0.283012779 | 0.1065 | -2.656 | 0.0079 | 0.049 |
| ENSG00000108797.11 | 6002.472 | -0.287512417 | 0.1083 | -2.655 | 0.00793 | 0.0491 |
| ENSG00000176148.15 | 990.3336 | -0.354373459 | 0.1335 | -2.654 | 0.00795 | 0.0492 |
| ENSG00000198899.2 | 154903 | 0.312706568 | 0.1178 | 2.6537 | 0.00796 | 0.0493 |
| ENSG00000044524.10 | 315.5983 | -0.293814902 | 0.1108 | -2.653 | 0.00799 | 0.0493 |
| ENSG00000115598.9 | 58.85751 | -0.395087489 | 0.1489 | -2.653 | 0.00799 | 0.0493 |
| ENSG00000197959.13 | 4363.837 | -0.291089955 | 0.1097 | -2.653 | 0.00799 | 0.0493 |
| ENSG00000251450.1 | 65.32187 | -0.317959125 | 0.1199 | -2.652 | 0.008 | 0.0494 |
| ENSG00000163539.16 | 9015.807 | -0.253743938 | 0.0957 | -2.652 | 0.008 | 0.0494 |
| ENSG00000134245.17 | 424.7539 | -0.243156881 | 0.0917 | -2.651 | 0.00804 | 0.0495 |
| ENSG00000144406.18 | 5056.642 | -0.305259535 | 0.1152 | -2.651 | 0.00804 | 0.0495 |
| ENSG00000196090.12 | 3823.163 | -0.378338291 | 0.1427 | -2.651 | 0.00803 | 0.0495 |
| ENSG00000174640.12 | 135.2757 | -0.308993877 | 0.1166 | -2.65 | 0.00805 | 0.0495 |
| ENSG00000068366.19 | 2712.951 | -0.307415096 | 0.1161 | -2.647 | 0.00811 | 0.0497 |
| ENSG00000138641.15 | 1479.972 | -0.216886732 | 0.0819 | -2.649 | 0.00808 | 0.0497 |
| ENSG00000145087.12 | 2441.92 | -0.374894919 | 0.1416 | -2.648 | 0.00809 | 0.0497 |
| ENSG00000156381.8 | 1267.103 | 0.239995245 | 0.0906 | 2.6476 | 0.00811 | 0.0497 |
| ENSG00000174236.3 | 17.32447 | -0.415376434 | 0.1568 | -2.649 | 0.00808 | 0.0497 |
| ENSG00000185345.18 | 630.5421 | -0.307183637 | 0.116 | -2.648 | 0.0081 | 0.0497 |
| ENSG00000248751.6 | 19.27273 | 0.681644085 | 0.2574 | 2.6477 | 0.0081 | 0.0497 |
| ENSG00000273448.1 | 86.42105 | 0.392807884 | 0.1484 | 2.6475 | 0.00811 | 0.0497 |
| ENSG00000279631.1 | 56.72486 | -0.354702314 | 0.134 | -2.648 | 0.0081 | 0.0497 |
| ENSG00000090932.10 | 177.2106 | 0.289672533 | 0.1095 | 2.6463 | 0.00814 | 0.0498 |
| ENSG00000160200.17 | 868.1911 | 0.466121421 | 0.1762 | 2.6455 | 0.00816 | 0.0499 |
| ENSG00000008086.11 | 4638.131 | -0.353379138 | 0.1336 | -2.645 | 0.00818 | 0.05 |
| ENSG00000163618.17 | 5252.509 | -0.317919578 | 0.1203 | -2.643 | 0.00821 | 0.0501 |
| ENSG00000254187.1 | 37.61185 | -0.398835314 | 0.1509 | -2.643 | 0.00821 | 0.0501 |
| ENSG00000051620.10 | 264.6308 | 0.290880521 | 0.1101 | 2.6427 | 0.00822 | 0.0501 |
| ENSG00000089356.17 | 28.28914 | 0.518820146 | 0.1963 | 2.6428 | 0.00822 | 0.0501 |
| ENSG00000253488.1 | 632.3482 | 1.280792354 | 0.4847 | 2.6425 | 0.00823 | 0.0501 |
| ENSG00000124772.11 | 3216.515 | -0.302995971 | 0.1147 | -2.642 | 0.00825 | 0.0502 |
| ENSG00000196937.10 | 4030.691 | -0.276872956 | 0.1048 | -2.641 | 0.00826 | 0.0502 |
| ENSG00000100852.12 | 4151.907 | 0.228933066 | 0.0867 | 2.6406 | 0.00828 | 0.0503 |
| ENSG00000144645.13 | 466.546 | -0.279145506 | 0.1057 | -2.64 | 0.00828 | 0.0503 |
| ENSG00000172340.14 | 332.6028 | 0.391786095 | 0.1484 | 2.6406 | 0.00828 | 0.0503 |
| ENSG00000185100.10 | 105.119 | 0.233218069 | 0.0883 | 2.6407 | 0.00827 | 0.0503 |
| ENSG00000138642.14 | 583.6052 | -0.241710302 | 0.0915 | -2.64 | 0.00828 | 0.0503 |
| ENSG00000157219.3 | 429.2163 | -0.300445134 | 0.1139 | -2.639 | 0.00832 | 0.0504 |
| ENSG00000157077.14 | 1927.273 | -0.297110579 | 0.1126 | -2.638 | 0.00833 | 0.0504 |
| ENSG00000196950.13 | 5418.238 | -0.340119417 | 0.1289 | -2.638 | 0.00833 | 0.0504 |
| ENSG00000263873.1 | 1039.559 | -0.432462288 | 0.1639 | -2.638 | 0.00833 | 0.0504 |
| ENSG00000099308.10 | 9603.365 | -0.275182413 | 0.1043 | -2.637 | 0.00836 | 0.0505 |
| ENSG00000205622.9 | 12.34384 | -0.437419718 | 0.1659 | -2.637 | 0.00836 | 0.0505 |
| ENSG00000258881.6 | 30.80426 | 0.385338872 | 0.1462 | 2.636 | 0.00839 | 0.0507 |
| ENSG00000004660.14 | 3830.768 | -0.272152394 | 0.1033 | -2.635 | 0.00841 | 0.0507 |
| ENSG00000236901.5 | 1190.996 | -0.286915329 | 0.1089 | -2.635 | 0.00841 | 0.0507 |
| ENSG00000268496.1 | 144.3214 | 0.252815889 | 0.096 | 2.6349 | 0.00842 | 0.0508 |
| ENSG00000128266.8 | 2592.189 | -0.263618842 | 0.1001 | -2.634 | 0.00844 | 0.0508 |
| ENSG00000182621.17 | 4385.658 | -0.350311524 | 0.133 | -2.634 | 0.00844 | 0.0508 |
| ENSG00000144791.9 | 319.5966 | 0.222384238 | 0.0845 | 2.6329 | 0.00847 | 0.0509 |
| ENSG00000165943.4 | 5824.051 | -0.331668704 | 0.126 | -2.633 | 0.00846 | 0.0509 |
| ENSG00000134109.10 | 635.3243 | -0.176780077 | 0.0672 | -2.632 | 0.00849 | 0.051 |
| ENSG00000107295.9 | 6044.146 | -0.3124528 | 0.1187 | -2.632 | 0.00849 | 0.051 |
| ENSG00000222019.7 | 75.23472 | 0.324767517 | 0.1234 | 2.6314 | 0.0085 | 0.0511 |
| ENSG00000075426.11 | 1591.348 | -0.362308732 | 0.1377 | -2.63 | 0.00853 | 0.0511 |
| ENSG00000077327.15 | 45.58916 | -0.557065222 | 0.2118 | -2.63 | 0.00854 | 0.0511 |
| ENSG00000112541.13 | 438.7509 | -0.307705431 | 0.117 | -2.629 | 0.00856 | 0.0511 |
| ENSG00000116133.11 | 4801.893 | -0.371231016 | 0.1412 | -2.63 | 0.00855 | 0.0511 |
| ENSG00000116604.17 | 4192.058 | -0.308725787 | 0.1173 | -2.631 | 0.00852 | 0.0511 |
| ENSG00000184613.10 | 8213.319 | -0.380925453 | 0.1449 | -2.629 | 0.00856 | 0.0511 |
| ENSG00000198898.12 | 2857.238 | -0.245353808 | 0.0933 | -2.629 | 0.00856 | 0.0511 |
| ENSG00000248522.1 | 30.3164 | -0.395543331 | 0.1504 | -2.631 | 0.00852 | 0.0511 |
| ENSG00000254198.1 | 64.928 | -0.32818956 | 0.1248 | -2.629 | 0.00856 | 0.0511 |
| ENSG00000259985.1 | 91.87314 | -0.281207369 | 0.107 | -2.629 | 0.00856 | 0.0511 |
| ENSG00000273489.1 | 295.6016 | 0.359772499 | 0.1368 | 2.6302 | 0.00853 | 0.0511 |
| ENSG00000146242.8 | 326.3123 | -0.547768479 | 0.2084 | -2.628 | 0.00858 | 0.0512 |
| ENSG00000029153.14 | 828.1542 | -0.271412111 | 0.1033 | -2.628 | 0.0086 | 0.0512 |
| ENSG00000126733.20 | 844.3955 | -0.312845967 | 0.1191 | -2.627 | 0.00861 | 0.0513 |
| ENSG00000255153.1 | 36.32655 | 0.326988742 | 0.1245 | 2.6269 | 0.00862 | 0.0513 |
| ENSG00000166813.14 | 146.454 | 0.261862127 | 0.0997 | 2.6266 | 0.00862 | 0.0513 |
| ENSG00000081138.13 | 272.5098 | -0.329228537 | 0.1254 | -2.626 | 0.00865 | 0.0514 |
| ENSG00000237515.7 | 2051.595 | -0.249012122 | 0.0949 | -2.625 | 0.00868 | 0.0516 |
| ENSG00000070182.18 | 1956.312 | -0.3741584 | 0.1426 | -2.624 | 0.00869 | 0.0516 |
| ENSG00000131899.10 | 1132.187 | 0.262650988 | 0.1001 | 2.6241 | 0.00869 | 0.0516 |
| ENSG00000186487.17 | 5607.213 | -0.29899363 | 0.114 | -2.623 | 0.00871 | 0.0517 |
| ENSG00000129654.7 | 39.59872 | 0.763789398 | 0.2912 | 2.6225 | 0.00873 | 0.0518 |
| ENSG00000004961.14 | 444.6926 | -0.25243689 | 0.0963 | -2.621 | 0.00876 | 0.0519 |
| ENSG00000009709.11 | 102.9941 | -0.431917591 | 0.1648 | -2.621 | 0.00878 | 0.0519 |
| ENSG00000122035.6 | 56.28503 | -0.458558669 | 0.175 | -2.621 | 0.00877 | 0.0519 |
| ENSG00000162728.4 | 4320.349 | -0.265112985 | 0.1012 | -2.621 | 0.00877 | 0.0519 |
| ENSG00000176974.18 | 150.4005 | 0.264292428 | 0.1008 | 2.621 | 0.00877 | 0.0519 |
| ENSG00000236882.7 | 17.52942 | 0.569090099 | 0.2171 | 2.6207 | 0.00877 | 0.0519 |
| ENSG00000005243.9 | 74.25177 | 0.396693849 | 0.1515 | 2.6192 | 0.00881 | 0.052 |
| ENSG00000006210.6 | 4017.932 | -0.292508683 | 0.1117 | -2.618 | 0.00884 | 0.052 |
| ENSG00000011347.9 | 6719.748 | -0.282668841 | 0.108 | -2.618 | 0.00884 | 0.052 |
| ENSG00000060709.14 | 2931.832 | -0.255908783 | 0.0978 | -2.618 | 0.00886 | 0.052 |
| ENSG00000078900.14 | 19.71062 | 0.440664898 | 0.1684 | 2.6169 | 0.00887 | 0.052 |
| ENSG00000081913.13 | 1737.667 | 0.30720035 | 0.1173 | 2.618 | 0.00885 | 0.052 |
| ENSG00000132718.8 | 17438.09 | -0.264990832 | 0.1013 | -2.617 | 0.00888 | 0.052 |
| ENSG00000135324.5 | 578.5965 | -0.373376744 | 0.1427 | -2.617 | 0.00887 | 0.052 |
| ENSG00000138867.16 | 539.1559 | 0.195538956 | 0.0747 | 2.6171 | 0.00887 | 0.052 |
| ENSG00000143858.11 | 122.4622 | -0.766502421 | 0.2929 | -2.617 | 0.00886 | 0.052 |
| ENSG00000155366.16 | 1100.271 | 0.341274857 | 0.1303 | 2.619 | 0.00882 | 0.052 |
| ENSG00000155792.9 | 250.5486 | -0.257149559 | 0.0983 | -2.617 | 0.00887 | 0.052 |
| ENSG00000164442.9 | 914.4017 | -0.286018329 | 0.1092 | -2.619 | 0.00881 | 0.052 |
| ENSG00000169715.14 | 650.2828 | 0.540019229 | 0.2062 | 2.6186 | 0.00883 | 0.052 |
| ENSG00000232871.8 | 29.68466 | 0.396552216 | 0.1514 | 2.6189 | 0.00882 | 0.052 |
| ENSG00000272221.1 | 19.83591 | 0.449328109 | 0.1715 | 2.6195 | 0.00881 | 0.052 |
| ENSG00000049883.14 | 572.9764 | -0.205457041 | 0.0785 | -2.617 | 0.00888 | 0.052 |
| ENSG00000063177.12 | 3458.24 | 0.251838967 | 0.0963 | 2.6147 | 0.00893 | 0.0522 |
| ENSG00000161835.10 | 1417.993 | -0.239032225 | 0.0914 | -2.615 | 0.00893 | 0.0522 |
| ENSG00000110344.9 | 1728.512 | -0.272127353 | 0.1042 | -2.612 | 0.00899 | 0.0525 |
| ENSG00000173391.8 | 170.9727 | -0.762132123 | 0.2918 | -2.612 | 0.009 | 0.0525 |
| ENSG00000162585.16 | 654.7559 | 0.200334303 | 0.0767 | 2.6113 | 0.00902 | 0.0526 |
| ENSG00000218336.8 | 2182.419 | -0.362584943 | 0.1388 | -2.611 | 0.00902 | 0.0526 |
| ENSG00000187672.12 | 2271.244 | -0.328930275 | 0.126 | -2.611 | 0.00903 | 0.0526 |
| ENSG00000198739.10 | 1004.15 | -0.292736499 | 0.1121 | -2.611 | 0.00903 | 0.0526 |
| ENSG00000187605.15 | 605.5633 | -0.225522763 | 0.0864 | -2.611 | 0.00904 | 0.0526 |
| ENSG00000139278.9 | 391.0485 | -0.29617841 | 0.1135 | -2.61 | 0.00906 | 0.0527 |
| ENSG00000006740.16 | 2860.807 | -0.25970067 | 0.0995 | -2.61 | 0.00906 | 0.0527 |
| ENSG00000137198.9 | 134.1869 | 0.494620576 | 0.1896 | 2.6089 | 0.00908 | 0.0528 |
| ENSG00000134594.4 | 254.4719 | -0.297713072 | 0.1141 | -2.609 | 0.00909 | 0.0528 |
| ENSG00000142606.15 | 11.54411 | 0.702224385 | 0.2692 | 2.6085 | 0.0091 | 0.0528 |
| ENSG00000250107.1 | 33.99741 | -0.351430565 | 0.1348 | -2.608 | 0.00911 | 0.0529 |
| ENSG00000125814.17 | 17749.21 | -0.301970594 | 0.1158 | -2.607 | 0.00912 | 0.0529 |
| ENSG00000101213.6 | 45.85336 | 0.32662901 | 0.1253 | 2.607 | 0.00913 | 0.0529 |
| ENSG00000198712.1 | 259406.2 | 0.323610051 | 0.1241 | 2.607 | 0.00913 | 0.0529 |
| ENSG00000138083.4 | 17.54672 | 0.552588183 | 0.2121 | 2.6057 | 0.00917 | 0.053 |
| ENSG00000170396.7 | 380.8086 | -0.298024396 | 0.1144 | -2.606 | 0.00916 | 0.053 |
| ENSG00000179796.11 | 425.6009 | 0.351504759 | 0.1349 | 2.6058 | 0.00917 | 0.053 |
| ENSG00000248383.4 | 167.1638 | -0.35325405 | 0.1356 | -2.605 | 0.00918 | 0.0531 |
| ENSG00000160801.13 | 201.2987 | 0.32987359 | 0.1266 | 2.6047 | 0.0092 | 0.0531 |
| ENSG00000152910.18 | 1489.625 | -0.297064054 | 0.1141 | -2.604 | 0.00922 | 0.0532 |
| ENSG00000236850.4 | 184.4416 | -0.254855507 | 0.0979 | -2.604 | 0.00922 | 0.0532 |
| ENSG00000007944.14 | 406.0805 | -0.250488765 | 0.0962 | -2.603 | 0.00924 | 0.0533 |
| ENSG00000142609.17 | 251.5347 | -0.188304502 | 0.0723 | -2.603 | 0.00924 | 0.0533 |
| ENSG00000135929.8 | 335.6032 | 0.360900008 | 0.1387 | 2.6023 | 0.00926 | 0.0533 |
| ENSG00000224963.3 | 19.58221 | 0.455501984 | 0.175 | 2.6023 | 0.00926 | 0.0533 |
| ENSG00000229920.2 | 122.4255 | 0.273019929 | 0.1049 | 2.6026 | 0.00925 | 0.0533 |
| ENSG00000112149.9 | 637.1848 | -0.291322122 | 0.112 | -2.601 | 0.00928 | 0.0534 |
| ENSG00000072315.3 | 176.7783 | -0.503984365 | 0.1938 | -2.601 | 0.00931 | 0.0535 |
| ENSG00000122733.12 | 6778.353 | -0.303823364 | 0.1168 | -2.601 | 0.00931 | 0.0535 |
| ENSG00000185518.11 | 10643.16 | -0.386792826 | 0.1487 | -2.6 | 0.00931 | 0.0535 |
| ENSG00000031081.10 | 589.4073 | 0.400372783 | 0.154 | 2.5999 | 0.00933 | 0.0535 |
| ENSG00000111328.6 | 2187.721 | 0.216048734 | 0.0831 | 2.5999 | 0.00932 | 0.0535 |
| ENSG00000138134.11 | 461.9565 | -0.268910279 | 0.1035 | -2.599 | 0.00934 | 0.0536 |
| ENSG00000117505.12 | 1478.412 | -0.196360153 | 0.0756 | -2.599 | 0.00935 | 0.0536 |
| ENSG00000171791.12 | 686.1207 | 0.246466463 | 0.0949 | 2.5982 | 0.00937 | 0.0536 |
| ENSG00000260975.1 | 55.36427 | 0.358656238 | 0.138 | 2.5983 | 0.00937 | 0.0536 |
| ENSG00000283623.1 | 19.39094 | -1.161544461 | 0.447 | -2.598 | 0.00936 | 0.0536 |
| ENSG00000107165.12 | 88.12402 | -0.417252066 | 0.1607 | -2.597 | 0.00942 | 0.0538 |
| ENSG00000177025.3 | 20.35285 | 0.404127626 | 0.1557 | 2.5962 | 0.00943 | 0.0538 |
| ENSG00000181800.5 | 124.2678 | -0.468037066 | 0.1803 | -2.596 | 0.00943 | 0.0538 |
| ENSG00000064961.18 | 305.2035 | 0.311852783 | 0.1202 | 2.5951 | 0.00946 | 0.0539 |
| ENSG00000267322.2 | 134.2138 | 0.246780337 | 0.0951 | 2.5952 | 0.00945 | 0.0539 |
| ENSG00000116741.7 | 716.6536 | -0.444537849 | 0.1714 | -2.593 | 0.00951 | 0.0541 |
| ENSG00000124198.8 | 2297.922 | -0.270441123 | 0.1043 | -2.593 | 0.00951 | 0.0541 |
| ENSG00000152049.6 | 173.8912 | 0.416234935 | 0.1605 | 2.5934 | 0.0095 | 0.0541 |
| ENSG00000272056.1 | 62.80134 | 0.2613852 | 0.1008 | 2.5937 | 0.00949 | 0.0541 |
| ENSG00000136167.13 | 183.446 | -0.727244274 | 0.2805 | -2.593 | 0.00953 | 0.0542 |
| ENSG00000188211.8 | 560.1773 | -0.362110163 | 0.1397 | -2.593 | 0.00952 | 0.0542 |
| ENSG00000174460.3 | 1910.27 | -0.349621827 | 0.1349 | -2.592 | 0.00953 | 0.0542 |
| ENSG00000226695.1 | 14.2461 | -0.582856917 | 0.2249 | -2.592 | 0.00954 | 0.0542 |
| ENSG00000131089.14 | 7517.81 | -0.247534092 | 0.0955 | -2.592 | 0.00955 | 0.0542 |
| ENSG00000120162.9 | 501.1095 | -0.248105359 | 0.0958 | -2.59 | 0.00959 | 0.0544 |
| ENSG00000138772.12 | 140.3964 | 0.31578015 | 0.1219 | 2.5897 | 0.0096 | 0.0544 |
| ENSG00000152413.14 | 2972.841 | -0.259033393 | 0.1 | -2.59 | 0.0096 | 0.0544 |
| ENSG00000180044.4 | 861.0661 | -0.341754469 | 0.132 | -2.59 | 0.0096 | 0.0544 |
| ENSG00000181656.6 | 411.7677 | -0.357440478 | 0.138 | -2.59 | 0.00961 | 0.0544 |
| ENSG00000100987.14 | 58.2868 | 0.338854079 | 0.1309 | 2.5881 | 0.00965 | 0.0545 |
| ENSG00000154917.10 | 18142.33 | -0.305845128 | 0.1182 | -2.588 | 0.00964 | 0.0545 |
| ENSG00000224682.1 | 26.71799 | 0.4384594 | 0.1694 | 2.5879 | 0.00965 | 0.0545 |
| ENSG00000227695.5 | 22.74553 | 0.360270377 | 0.1392 | 2.5881 | 0.00965 | 0.0545 |
| ENSG00000128245.14 | 29668.18 | -0.242746717 | 0.0938 | -2.587 | 0.00967 | 0.0546 |
| ENSG00000246223.8 | 46.16822 | 0.27558082 | 0.1065 | 2.5874 | 0.00967 | 0.0546 |
| ENSG00000183850.13 | 17.72717 | 0.455850245 | 0.1762 | 2.5871 | 0.00968 | 0.0546 |
| ENSG00000113594.9 | 1924.354 | 0.343627612 | 0.1329 | 2.5854 | 0.00973 | 0.0548 |
| ENSG00000101605.12 | 333.3735 | 0.376946147 | 0.1458 | 2.585 | 0.00974 | 0.0548 |
| ENSG00000174473.15 | 293.7509 | -0.266193995 | 0.103 | -2.585 | 0.00974 | 0.0548 |
| ENSG00000157600.11 | 540.1462 | 0.18792394 | 0.0728 | 2.5829 | 0.0098 | 0.0551 |
| ENSG00000182389.19 | 3408.277 | -0.294286336 | 0.1139 | -2.583 | 0.0098 | 0.0551 |
| ENSG00000175395.15 | 2259.47 | -0.257796717 | 0.0998 | -2.583 | 0.0098 | 0.0551 |
| ENSG00000162733.16 | 789.0615 | 0.389055496 | 0.1507 | 2.5824 | 0.00981 | 0.0551 |
| ENSG00000198910.12 | 5541.18 | -0.30341379 | 0.1175 | -2.582 | 0.00983 | 0.0552 |
| ENSG00000183307.3 | 839.4144 | -0.328663299 | 0.1273 | -2.581 | 0.00985 | 0.0553 |
| ENSG00000272419.5 | 378.7016 | -0.18765326 | 0.0727 | -2.581 | 0.00986 | 0.0553 |
| ENSG00000140067.6 | 109.9859 | 0.406881873 | 0.1577 | 2.58 | 0.00988 | 0.0553 |
| ENSG00000233110.1 | 55.35554 | -0.328156159 | 0.1272 | -2.58 | 0.00988 | 0.0553 |
| ENSG00000261399.1 | 21.99447 | 0.38782359 | 0.1503 | 2.5796 | 0.00989 | 0.0554 |
| ENSG00000126500.3 | 670.3873 | -0.258308659 | 0.1002 | -2.579 | 0.0099 | 0.0554 |
| ENSG00000178171.10 | 789.2573 | -0.243738991 | 0.0945 | -2.579 | 0.00991 | 0.0554 |
| ENSG00000187049.9 | 132.1537 | 0.227917532 | 0.0884 | 2.5786 | 0.00992 | 0.0554 |
| ENSG00000048052.21 | 1457.534 | -0.251945022 | 0.0977 | -2.578 | 0.00994 | 0.0555 |
| ENSG00000100678.18 | 769.3332 | -0.241710203 | 0.0938 | -2.577 | 0.00996 | 0.0556 |
| ENSG00000158195.10 | 1278.742 | 0.337286555 | 0.1309 | 2.5772 | 0.00996 | 0.0556 |
| ENSG00000174306.21 | 2212.965 | 0.174428087 | 0.0677 | 2.5767 | 0.00998 | 0.0556 |
| ENSG00000198108.3 | 245.2149 | -0.291719963 | 0.1132 | -2.576 | 0.00999 | 0.0556 |
| ENSG00000228878.7 | 86.30966 | 0.235535941 | 0.0914 | 2.5763 | 0.00999 | 0.0556 |
| ENSG00000232748.3 | 79.8927 | 0.298520794 | 0.1158 | 2.5769 | 0.00997 | 0.0556 |
| ENSG00000260118.1 | 53.82472 | 0.35320691 | 0.1371 | 2.5763 | 0.00999 | 0.0556 |
| ENSG00000223949.6 | 155.693 | 0.26759502 | 0.1039 | 2.5758 | 0.01 | 0.0556 |
| ENSG00000270909.1 | 16.34258 | 0.510809868 | 0.1983 | 2.5759 | 0.01 | 0.0556 |
| ENSG00000189212.12 | 712.5601 | -0.264459064 | 0.1027 | -2.575 | 0.01001 | 0.0556 |
| ENSG00000204052.4 | 495.7131 | -0.322388398 | 0.1252 | -2.575 | 0.01002 | 0.0556 |
| ENSG00000166159.10 | 891.8399 | -0.281817533 | 0.1095 | -2.574 | 0.01004 | 0.0557 |
| ENSG00000243708.10 | 253.847 | 0.280264322 | 0.1089 | 2.5745 | 0.01004 | 0.0557 |
| ENSG00000164038.14 | 1230.117 | -0.282768823 | 0.1099 | -2.574 | 0.01006 | 0.0558 |
| ENSG00000171914.15 | 5527.786 | -0.315423258 | 0.1226 | -2.573 | 0.01008 | 0.0559 |
| ENSG00000235772.1 | 10.38002 | -0.648633591 | 0.2521 | -2.573 | 0.01009 | 0.0559 |
| ENSG00000164209.16 | 2312.301 | -0.262755958 | 0.1022 | -2.572 | 0.01011 | 0.0559 |
| ENSG00000006715.15 | 3407.791 | -0.233712342 | 0.0909 | -2.572 | 0.01012 | 0.056 |
| ENSG00000171766.15 | 3383.873 | 0.28837211 | 0.1121 | 2.5717 | 0.01012 | 0.056 |
| ENSG00000003989.16 | 463.9506 | 0.589639488 | 0.2294 | 2.5705 | 0.01016 | 0.0561 |
| ENSG00000065183.15 | 499.728 | -0.19467073 | 0.0757 | -2.57 | 0.01017 | 0.0561 |
| ENSG00000243232.4 | 1189.511 | -0.407252549 | 0.1585 | -2.57 | 0.01017 | 0.0561 |
| ENSG00000261959.1 | 13.59009 | 0.557637449 | 0.217 | 2.5698 | 0.01018 | 0.0561 |
| ENSG00000197653.15 | 233.2781 | -0.265087642 | 0.1032 | -2.569 | 0.0102 | 0.0562 |
| ENSG00000275400.1 | 39.73678 | 0.377129992 | 0.1468 | 2.5687 | 0.01021 | 0.0563 |
| ENSG00000135631.16 | 2098.449 | -0.280244916 | 0.1091 | -2.568 | 0.01021 | 0.0563 |
| ENSG00000124785.8 | 3930.764 | -0.253839732 | 0.0988 | -2.568 | 0.01023 | 0.0563 |
| ENSG00000165714.10 | 513.1283 | -0.20392287 | 0.0794 | -2.567 | 0.01025 | 0.0564 |
| ENSG00000102466.15 | 1510.351 | -0.221159687 | 0.0862 | -2.567 | 0.01027 | 0.0565 |
| ENSG00000165973.18 | 707.93 | -0.430373743 | 0.1677 | -2.566 | 0.01028 | 0.0565 |
| ENSG00000184307.13 | 1113.249 | -0.302130176 | 0.1178 | -2.565 | 0.01031 | 0.0567 |
| ENSG00000163444.11 | 2638.408 | -0.258273221 | 0.1007 | -2.564 | 0.01034 | 0.0568 |
| ENSG00000135063.17 | 286.4839 | 0.534742683 | 0.2086 | 2.563 | 0.01038 | 0.0569 |
| ENSG00000134375.10 | 1445.121 | -0.256072274 | 0.0999 | -2.563 | 0.01038 | 0.0569 |
| ENSG00000122778.9 | 1342.218 | -0.249549377 | 0.0974 | -2.562 | 0.0104 | 0.057 |
| ENSG00000100647.7 | 739.1302 | -0.246516097 | 0.0962 | -2.562 | 0.01042 | 0.0571 |
| ENSG00000260212.1 | 1087.284 | 1.398009873 | 0.546 | 2.5606 | 0.01045 | 0.0572 |
| ENSG00000122641.10 | 143.7325 | -0.573121152 | 0.2239 | -2.56 | 0.01047 | 0.0573 |
| ENSG00000077092.18 | 379.2991 | -0.281858154 | 0.1101 | -2.559 | 0.01049 | 0.0574 |
| ENSG00000171532.4 | 3255.58 | 0.264831794 | 0.1035 | 2.5592 | 0.01049 | 0.0574 |
| ENSG00000184408.9 | 1043.793 | -0.251999751 | 0.0985 | -2.559 | 0.01051 | 0.0574 |
| ENSG00000067064.10 | 2603.263 | -0.198371619 | 0.0776 | -2.557 | 0.01057 | 0.0577 |
| ENSG00000148408.12 | 3120.308 | -0.264625946 | 0.1035 | -2.556 | 0.01059 | 0.0577 |
| ENSG00000160191.17 | 255.7492 | 0.237140033 | 0.0928 | 2.5557 | 0.0106 | 0.0577 |
| ENSG00000164588.6 | 1951.22 | -0.308447682 | 0.1207 | -2.556 | 0.01058 | 0.0577 |
| ENSG00000181090.18 | 787.9081 | 0.151001788 | 0.0591 | 2.5557 | 0.0106 | 0.0577 |
| ENSG00000280850.1 | 13.68461 | 0.574240663 | 0.2246 | 2.5563 | 0.01058 | 0.0577 |
| ENSG00000120053.10 | 4765.394 | -0.301643159 | 0.118 | -2.555 | 0.01061 | 0.0577 |
| ENSG00000239779.6 | 1305.214 | 0.196152064 | 0.0768 | 2.5554 | 0.01061 | 0.0577 |
| ENSG00000049130.13 | 1012.553 | -0.290567963 | 0.1137 | -2.555 | 0.01063 | 0.0578 |
| ENSG00000107738.19 | 618.6765 | 0.458828039 | 0.1796 | 2.5546 | 0.01063 | 0.0578 |
| ENSG00000140323.5 | 1995.67 | -0.206157897 | 0.0807 | -2.554 | 0.01065 | 0.0578 |
| ENSG00000268869.5 | 14.08219 | 0.552874433 | 0.2165 | 2.5539 | 0.01065 | 0.0578 |
| ENSG00000008394.12 | 845.4622 | 0.394966898 | 0.1547 | 2.5529 | 0.01068 | 0.058 |
| ENSG00000177234.7 | 64.0933 | 0.295436969 | 0.1157 | 2.5524 | 0.0107 | 0.058 |
| ENSG00000255823.3 | 25.91352 | 0.635677026 | 0.249 | 2.5524 | 0.0107 | 0.058 |
| ENSG00000103449.11 | 498.0768 | 0.309296995 | 0.1212 | 2.552 | 0.01071 | 0.058 |
| ENSG00000135953.10 | 101.3057 | -0.203029903 | 0.0796 | -2.55 | 0.01076 | 0.0582 |
| ENSG00000150995.18 | 6571.974 | -0.347497863 | 0.1363 | -2.55 | 0.01077 | 0.0582 |
| ENSG00000156017.12 | 454.8406 | -0.210197215 | 0.0824 | -2.55 | 0.01076 | 0.0582 |
| ENSG00000181852.17 | 2194.494 | -0.240536386 | 0.0943 | -2.549 | 0.01079 | 0.0583 |
| ENSG00000197043.13 | 5556.206 | -0.313001665 | 0.1228 | -2.55 | 0.01079 | 0.0583 |
| ENSG00000130653.15 | 804.2752 | 0.194074712 | 0.0762 | 2.5483 | 0.01082 | 0.0585 |
| ENSG00000198668.10 | 87448.27 | -0.254303832 | 0.0998 | -2.548 | 0.01084 | 0.0585 |
| ENSG00000117016.9 | 11156.78 | -0.329789315 | 0.1295 | -2.548 | 0.01085 | 0.0585 |
| ENSG00000110492.15 | 160.8786 | 0.308377629 | 0.1211 | 2.5471 | 0.01086 | 0.0585 |
| ENSG00000162913.9 | 97.45754 | -0.297650811 | 0.1169 | -2.547 | 0.01086 | 0.0585 |
| ENSG00000171208.9 | 1214.624 | -0.256679268 | 0.1008 | -2.546 | 0.01091 | 0.0588 |
| ENSG00000006283.17 | 1419.544 | -0.289818333 | 0.1139 | -2.545 | 0.01092 | 0.0588 |
| ENSG00000213468.4 | 28.92783 | 0.358374251 | 0.1408 | 2.5453 | 0.01092 | 0.0588 |
| ENSG00000229425.2 | 445.4559 | -0.293420978 | 0.1153 | -2.544 | 0.01095 | 0.0589 |
| ENSG00000158201.9 | 211.0305 | 0.362907493 | 0.1427 | 2.5434 | 0.01098 | 0.059 |
| ENSG00000102096.9 | 448.9575 | -0.241323271 | 0.0949 | -2.543 | 0.01099 | 0.059 |
| ENSG00000175906.4 | 381.2271 | -0.332519514 | 0.1308 | -2.543 | 0.01099 | 0.059 |
| ENSG00000133985.2 | 1295.136 | -0.238427712 | 0.0938 | -2.542 | 0.01101 | 0.0591 |
| ENSG00000162636.15 | 2482.331 | -0.336121846 | 0.1322 | -2.542 | 0.01102 | 0.0591 |
| ENSG00000225792.1 | 55.3651 | 0.383723991 | 0.151 | 2.5416 | 0.01103 | 0.0592 |
| ENSG00000198718.12 | 1515.137 | -0.276202572 | 0.1087 | -2.541 | 0.01105 | 0.0592 |
| ENSG00000008277.14 | 3699.29 | -0.29330258 | 0.1154 | -2.541 | 0.01105 | 0.0592 |
| ENSG00000139182.13 | 8359.421 | -0.338175988 | 0.1331 | -2.54 | 0.01108 | 0.0592 |
| ENSG00000163932.13 | 1006.419 | -0.221652502 | 0.0873 | -2.54 | 0.01107 | 0.0592 |
| ENSG00000164830.18 | 5624.015 | -0.369655482 | 0.1455 | -2.54 | 0.01107 | 0.0592 |
| ENSG00000280165.1 | 1103.083 | -0.303926199 | 0.1197 | -2.54 | 0.01109 | 0.0593 |
| ENSG00000214688.4 | 75.13975 | 0.826216293 | 0.3253 | 2.5395 | 0.0111 | 0.0593 |
| ENSG00000233705.6 | 2523.854 | -0.297842694 | 0.1173 | -2.539 | 0.01111 | 0.0593 |
| ENSG00000181722.16 | 1292.604 | 0.362567876 | 0.1428 | 2.5386 | 0.01113 | 0.0594 |
| ENSG00000184185.9 | 399.7368 | -0.342715013 | 0.135 | -2.538 | 0.01115 | 0.0595 |
| ENSG00000178297.12 | 39.44485 | 0.380913522 | 0.1502 | 2.5367 | 0.01119 | 0.0597 |
| ENSG00000178081.12 | 10.72114 | 0.578868467 | 0.2283 | 2.536 | 0.01121 | 0.0597 |
| ENSG00000272414.5 | 151.2854 | -0.264413822 | 0.1043 | -2.536 | 0.01122 | 0.0597 |
| ENSG00000153814.11 | 1594.638 | -0.258709784 | 0.102 | -2.536 | 0.01123 | 0.0598 |
| ENSG00000172782.11 | 188.9571 | -0.290097248 | 0.1144 | -2.535 | 0.01125 | 0.0599 |
| ENSG00000198938.2 | 374522.2 | 0.302339834 | 0.1193 | 2.5345 | 0.01126 | 0.0599 |
| ENSG00000128335.13 | 1242.687 | -0.275974858 | 0.1089 | -2.534 | 0.01127 | 0.0599 |
| ENSG00000198825.12 | 6693.965 | -0.268264589 | 0.1059 | -2.534 | 0.01127 | 0.0599 |
| ENSG00000136143.14 | 2182.251 | -0.258330755 | 0.102 | -2.534 | 0.01128 | 0.0599 |
| ENSG00000254682.1 | 84.7931 | 0.402438261 | 0.1589 | 2.5333 | 0.0113 | 0.06 |
| ENSG00000170325.14 | 431.5805 | -0.221781026 | 0.0876 | -2.533 | 0.01131 | 0.06 |
| ENSG00000008853.16 | 4327.229 | -0.229922814 | 0.0908 | -2.532 | 0.01134 | 0.0601 |
| ENSG00000215483.9 | 56.38949 | 0.30933917 | 0.1222 | 2.5318 | 0.01135 | 0.0601 |
| ENSG00000144567.10 | 4758.889 | -0.246608239 | 0.0974 | -2.531 | 0.01136 | 0.0601 |
| ENSG00000184343.10 | 154.858 | 0.173094879 | 0.0684 | 2.5316 | 0.01135 | 0.0601 |
| ENSG00000197106.6 | 7274.827 | -0.326031577 | 0.1288 | -2.53 | 0.01139 | 0.0603 |
| ENSG00000204767.3 | 68.38254 | -0.343710759 | 0.1359 | -2.529 | 0.01144 | 0.0605 |
| ENSG00000171885.13 | 13423.46 | 0.554006732 | 0.2191 | 2.5285 | 0.01146 | 0.0605 |
| ENSG00000105409.16 | 23577.6 | -0.283937245 | 0.1123 | -2.527 | 0.01149 | 0.0607 |
| ENSG00000159248.4 | 124.4096 | -0.314196178 | 0.1244 | -2.527 | 0.01152 | 0.0607 |
| ENSG00000165023.6 | 7398.52 | -0.402507708 | 0.1593 | -2.526 | 0.01152 | 0.0607 |
| ENSG00000165548.10 | 1205.612 | -0.250637015 | 0.0992 | -2.527 | 0.01151 | 0.0607 |
| ENSG00000181804.14 | 208.5989 | 0.317193905 | 0.1256 | 2.5263 | 0.01153 | 0.0607 |
| ENSG00000270629.5 | 752.6729 | 0.369333564 | 0.1462 | 2.527 | 0.0115 | 0.0607 |
| ENSG00000279952.1 | 29.7371 | -0.364885149 | 0.1444 | -2.527 | 0.0115 | 0.0607 |
| ENSG00000283020.1 | 20.80242 | 0.411682335 | 0.163 | 2.5264 | 0.01152 | 0.0607 |
| ENSG00000178502.5 | 243.8253 | -0.399291861 | 0.1581 | -2.525 | 0.01156 | 0.0608 |
| ENSG00000093217.9 | 101.6038 | -0.212087377 | 0.084 | -2.525 | 0.01157 | 0.0609 |
| ENSG00000185008.17 | 1745.767 | -0.322476566 | 0.1277 | -2.524 | 0.01159 | 0.0609 |
| ENSG00000279232.1 | 88.45218 | -0.602866862 | 0.2389 | -2.524 | 0.0116 | 0.0609 |
| ENSG00000002586.18 | 376.3772 | 0.455093422 | 0.1804 | 2.5229 | 0.01164 | 0.061 |
| ENSG00000002586.18_PAR_Y | 376.3772 | 0.455093422 | 0.1804 | 2.5229 | 0.01164 | 0.061 |
| ENSG00000164659.14 | 703.4589 | -0.245790404 | 0.0974 | -2.523 | 0.01163 | 0.061 |
| ENSG00000189181.4 | 63.3728 | -0.449660949 | 0.1782 | -2.523 | 0.01164 | 0.061 |
| ENSG00000243384.1 | 11.12598 | 0.775091668 | 0.3072 | 2.5229 | 0.01164 | 0.061 |
| ENSG00000231187.2 | 341.4043 | -0.340839754 | 0.1351 | -2.523 | 0.01165 | 0.061 |
| ENSG00000138640.14 | 1147.132 | -0.174522266 | 0.0692 | -2.522 | 0.01165 | 0.061 |
| ENSG00000268182.5 | 288.0233 | -0.316522574 | 0.1255 | -2.522 | 0.01167 | 0.061 |
| ENSG00000160796.16 | 359.2405 | -0.213658854 | 0.0847 | -2.522 | 0.01167 | 0.061 |
| ENSG00000187094.11 | 4254.704 | -0.255030894 | 0.1011 | -2.522 | 0.01168 | 0.0611 |
| ENSG00000106236.3 | 1889.908 | -0.323913881 | 0.1285 | -2.521 | 0.0117 | 0.0611 |
| ENSG00000165434.7 | 7457.921 | -0.343544859 | 0.1364 | -2.519 | 0.01176 | 0.0614 |
| ENSG00000183336.8 | 253.2068 | -0.288692918 | 0.1146 | -2.519 | 0.01176 | 0.0614 |
| ENSG00000265972.5 | 1290.264 | 0.454512288 | 0.1804 | 2.5188 | 0.01177 | 0.0614 |
| ENSG00000074211.13 | 8506.622 | -0.229285239 | 0.0911 | -2.517 | 0.01183 | 0.0615 |
| ENSG00000105519.14 | 327.9454 | 0.3010829 | 0.1196 | 2.518 | 0.0118 | 0.0615 |
| ENSG00000116478.11 | 381.0081 | 0.256853197 | 0.1021 | 2.5166 | 0.01185 | 0.0615 |
| ENSG00000144290.16 | 5285.085 | -0.350285026 | 0.1391 | -2.518 | 0.01182 | 0.0615 |
| ENSG00000150551.10 | 775.9417 | 0.306513579 | 0.1218 | 2.5169 | 0.01184 | 0.0615 |
| ENSG00000156804.7 | 866.4405 | -0.268901743 | 0.1069 | -2.517 | 0.01185 | 0.0615 |
| ENSG00000157680.15 | 1804.223 | -0.320950785 | 0.1275 | -2.518 | 0.01181 | 0.0615 |
| ENSG00000172602.9 | 1323.576 | -0.313523739 | 0.1246 | -2.517 | 0.01185 | 0.0615 |
| ENSG00000272869.5 | 37.15016 | 0.551291831 | 0.219 | 2.5174 | 0.01182 | 0.0615 |
| ENSG00000145451.12 | 1274.623 | -0.345680107 | 0.1374 | -2.515 | 0.01189 | 0.0617 |
| ENSG00000151150.21 | 5940.425 | -0.277338055 | 0.1103 | -2.515 | 0.0119 | 0.0617 |
| ENSG00000105379.9 | 1239.421 | 0.209879911 | 0.0835 | 2.5139 | 0.01194 | 0.0619 |
| ENSG00000139514.12 | 1513.236 | -0.216797866 | 0.0862 | -2.514 | 0.01194 | 0.0619 |
| ENSG00000254221.2 | 174.8783 | 0.384373921 | 0.1529 | 2.5138 | 0.01194 | 0.0619 |
| ENSG00000186479.4 | 1945.043 | -0.521636926 | 0.2075 | -2.513 | 0.01196 | 0.0619 |
| ENSG00000113532.12 | 289.1442 | -0.239498191 | 0.0953 | -2.513 | 0.01197 | 0.0619 |
| ENSG00000232527.7 | 47.96938 | -0.308576327 | 0.1228 | -2.513 | 0.01197 | 0.0619 |
| ENSG00000137273.3 | 285.1203 | 0.409070007 | 0.1628 | 2.5121 | 0.012 | 0.062 |
| ENSG00000119280.16 | 1558.032 | 0.387135435 | 0.1541 | 2.5116 | 0.01202 | 0.0621 |
| ENSG00000235532.1 | 13.53722 | 0.451602297 | 0.1799 | 2.5109 | 0.01204 | 0.0621 |
| ENSG00000237720.1 | 49.80653 | -0.399451019 | 0.1591 | -2.511 | 0.01204 | 0.0621 |
| ENSG00000185745.9 | 1043.486 | -0.193434759 | 0.077 | -2.511 | 0.01205 | 0.0622 |
| ENSG00000155093.17 | 6891.529 | -0.276479096 | 0.1101 | -2.51 | 0.01206 | 0.0622 |
| ENSG00000143570.17 | 414.7033 | 0.25162335 | 0.1002 | 2.51 | 0.01207 | 0.0622 |
| ENSG00000135919.12 | 4779.37 | 0.245480787 | 0.0979 | 2.5086 | 0.01212 | 0.0623 |
| ENSG00000170113.15 | 2342.728 | -0.256258945 | 0.1021 | -2.509 | 0.01212 | 0.0623 |
| ENSG00000179520.10 | 134.2427 | -0.606923568 | 0.2419 | -2.509 | 0.01212 | 0.0623 |
| ENSG00000248668.2 | 26.87983 | 0.32256015 | 0.1286 | 2.5088 | 0.01211 | 0.0623 |
| ENSG00000101638.13 | 2947.252 | -0.294057725 | 0.1173 | -2.507 | 0.01218 | 0.0625 |
| ENSG00000139163.15 | 2653.063 | -0.225574434 | 0.09 | -2.507 | 0.01218 | 0.0625 |
| ENSG00000141314.12 | 435.5739 | 0.226885395 | 0.0905 | 2.5068 | 0.01218 | 0.0625 |
| ENSG00000249244.1 | 29.92091 | -0.349069246 | 0.1392 | -2.507 | 0.01217 | 0.0625 |
| ENSG00000274292.1 | 62.1389 | 0.290800617 | 0.116 | 2.5068 | 0.01218 | 0.0625 |
| ENSG00000150275.17 | 427.1135 | 0.293429876 | 0.1171 | 2.5066 | 0.01219 | 0.0625 |
| ENSG00000171729.13 | 90.80881 | 0.339073362 | 0.1353 | 2.5062 | 0.0122 | 0.0625 |
| ENSG00000154975.13 | 2626.723 | -0.253037262 | 0.101 | -2.505 | 0.01223 | 0.0626 |
| ENSG00000280920.1 | 9.573427 | 0.517658256 | 0.2066 | 2.5054 | 0.01223 | 0.0626 |
| ENSG00000183864.4 | 873.4002 | 0.296728282 | 0.1185 | 2.5047 | 0.01226 | 0.0627 |
| ENSG00000229676.2 | 25.88474 | 0.386882339 | 0.1545 | 2.5039 | 0.01228 | 0.0628 |
| ENSG00000145476.15 | 716.3473 | 0.332862445 | 0.133 | 2.5034 | 0.0123 | 0.0628 |
| ENSG00000267801.1 | 18.06239 | 0.461781744 | 0.1845 | 2.5034 | 0.0123 | 0.0628 |
| ENSG00000263647.1 | 21.60311 | -0.486005661 | 0.1942 | -2.503 | 0.01231 | 0.0629 |
| ENSG00000054356.13 | 7722.181 | -0.359393106 | 0.1436 | -2.503 | 0.01232 | 0.0629 |
| ENSG00000147010.17 | 1674.325 | -0.209298624 | 0.0837 | -2.502 | 0.01235 | 0.063 |
| ENSG00000185915.5 | 76.57008 | -0.376750737 | 0.1506 | -2.502 | 0.01235 | 0.063 |
| ENSG00000166501.12 | 17476.35 | -0.306234833 | 0.1224 | -2.502 | 0.01236 | 0.063 |
| ENSG00000204116.11 | 1565.204 | -0.2674794 | 0.107 | -2.501 | 0.01239 | 0.0631 |
| ENSG00000165209.18 | 1399.433 | -0.243717579 | 0.0975 | -2.5 | 0.01244 | 0.0633 |
| ENSG00000126368.5 | 1566.928 | 0.294043994 | 0.1177 | 2.4993 | 0.01244 | 0.0633 |
| ENSG00000230102.7 | 17.86923 | 0.354378977 | 0.1418 | 2.4989 | 0.01246 | 0.0634 |
| ENSG00000233297.4 | 25.40108 | -0.543713585 | 0.2177 | -2.498 | 0.01249 | 0.0635 |
| ENSG00000166960.16 | 18.36506 | -0.342918565 | 0.1373 | -2.498 | 0.01249 | 0.0635 |
| ENSG00000141655.15 | 246.1323 | -0.236817664 | 0.0948 | -2.497 | 0.01253 | 0.0636 |
| ENSG00000274118.1 | 10.75512 | 0.534943737 | 0.2143 | 2.4958 | 0.01257 | 0.0638 |
| ENSG00000039139.9 | 154.1238 | -0.279777645 | 0.1121 | -2.495 | 0.0126 | 0.0639 |
| ENSG00000115155.16 | 158.6361 | -0.483054579 | 0.1936 | -2.495 | 0.01261 | 0.0639 |
| ENSG00000179954.15 | 134.4545 | 0.331649399 | 0.133 | 2.4941 | 0.01263 | 0.064 |
| ENSG00000075340.22 | 7124.923 | -0.302972733 | 0.1215 | -2.493 | 0.01266 | 0.0641 |
| ENSG00000174013.7 | 1203.806 | -0.231740739 | 0.093 | -2.493 | 0.01268 | 0.0642 |
| ENSG00000174175.16 | 14.42156 | -0.882420971 | 0.354 | -2.493 | 0.01268 | 0.0642 |
| ENSG00000159176.13 | 6750.4 | 0.359802406 | 0.1444 | 2.4921 | 0.0127 | 0.0642 |
| ENSG00000239389.7 | 118.1467 | 0.510732388 | 0.2049 | 2.4922 | 0.0127 | 0.0642 |
| ENSG00000256092.2 | 29.29287 | 0.327931414 | 0.1316 | 2.4914 | 0.01272 | 0.0643 |
| ENSG00000273674.4 | 49.52931 | 0.258681604 | 0.1038 | 2.4913 | 0.01273 | 0.0643 |
| ENSG00000158050.4 | 267.6206 | -0.361923912 | 0.1453 | -2.491 | 0.01274 | 0.0643 |
| ENSG00000215512.9 | 41.60709 | 0.334451018 | 0.1343 | 2.4907 | 0.01275 | 0.0643 |
| ENSG00000135838.13 | 321.3377 | 0.305953465 | 0.1229 | 2.4903 | 0.01276 | 0.0643 |
| ENSG00000177272.8 | 123.9535 | -0.43287254 | 0.1738 | -2.49 | 0.01276 | 0.0643 |
| ENSG00000003137.8 | 695.3326 | -0.361924299 | 0.1453 | -2.49 | 0.01277 | 0.0643 |
| ENSG00000272279.1 | 11.50928 | 0.651037563 | 0.2615 | 2.4894 | 0.0128 | 0.0644 |
| ENSG00000165449.11 | 267.8298 | 0.498025651 | 0.2001 | 2.4888 | 0.01282 | 0.0645 |
| ENSG00000228492.2 | 83.70689 | -0.284568836 | 0.1143 | -2.489 | 0.01282 | 0.0645 |
| ENSG00000115232.13 | 67.80359 | -0.372116214 | 0.1496 | -2.488 | 0.01284 | 0.0646 |
| ENSG00000139112.10 | 9462.215 | -0.292610324 | 0.1176 | -2.488 | 0.01286 | 0.0646 |
| ENSG00000151789.10 | 1096.105 | -0.294718516 | 0.1185 | -2.488 | 0.01286 | 0.0646 |
| ENSG00000177674.15 | 104.0367 | 0.33747702 | 0.1357 | 2.4877 | 0.01286 | 0.0646 |
| ENSG00000168658.18 | 26.59291 | 0.446983306 | 0.1797 | 2.4869 | 0.01289 | 0.0647 |
| ENSG00000241399.6 | 219.394 | 0.408552991 | 0.1643 | 2.4866 | 0.0129 | 0.0647 |
| ENSG00000112715.20 | 918.0238 | 0.407041602 | 0.1637 | 2.4859 | 0.01292 | 0.0648 |
| ENSG00000283533.1 | 9.152893 | -0.509553914 | 0.205 | -2.486 | 0.01293 | 0.0648 |
| ENSG00000173805.15 | 520.0394 | 0.283885429 | 0.1142 | 2.4853 | 0.01295 | 0.0648 |
| ENSG00000227825.4 | 186.9359 | -0.339628646 | 0.1367 | -2.485 | 0.01295 | 0.0649 |
| ENSG00000089558.8 | 168.4202 | -0.388838178 | 0.1566 | -2.484 | 0.013 | 0.065 |
| ENSG00000130413.15 | 252.5896 | 0.224539772 | 0.0904 | 2.4839 | 0.01299 | 0.065 |
| ENSG00000110665.11 | 13.50867 | -0.511898452 | 0.2061 | -2.484 | 0.01301 | 0.065 |
| ENSG00000131981.15 | 497.6027 | 0.398577699 | 0.1605 | 2.4833 | 0.01302 | 0.065 |
| ENSG00000254510.1 | 35.03569 | -0.335648732 | 0.1352 | -2.483 | 0.01302 | 0.065 |
| ENSG00000119403.13 | 338.7389 | 0.254295461 | 0.1025 | 2.4821 | 0.01306 | 0.0652 |
| ENSG00000129194.7 | 155.3803 | 0.287944413 | 0.116 | 2.4818 | 0.01307 | 0.0652 |
| ENSG00000205810.8 | 13.36393 | 0.559141015 | 0.2254 | 2.4812 | 0.0131 | 0.0653 |
| ENSG00000240694.8 | 12466.22 | -0.269410623 | 0.1086 | -2.481 | 0.01311 | 0.0653 |
| ENSG00000167157.10 | 10.27713 | 0.682254467 | 0.2751 | 2.4804 | 0.01312 | 0.0653 |
| ENSG00000169851.15 | 2868.045 | -0.245236854 | 0.0989 | -2.48 | 0.01313 | 0.0653 |
| ENSG00000228918.3 | 12.4938 | 0.392334091 | 0.1582 | 2.4803 | 0.01313 | 0.0653 |
| ENSG00000100664.10 | 5926.358 | -0.22241455 | 0.0897 | -2.48 | 0.01314 | 0.0654 |
| ENSG00000105143.12 | 658.0097 | -0.232232454 | 0.0936 | -2.48 | 0.01314 | 0.0654 |
| ENSG00000074527.11 | 707.7831 | -0.230568964 | 0.093 | -2.478 | 0.01319 | 0.0654 |
| ENSG00000102471.13 | 4180.207 | -0.290694866 | 0.1173 | -2.478 | 0.0132 | 0.0654 |
| ENSG00000112706.11 | 70.34466 | -0.266307563 | 0.1074 | -2.479 | 0.01316 | 0.0654 |
| ENSG00000131437.15 | 5878.348 | -0.275642391 | 0.1112 | -2.479 | 0.01319 | 0.0654 |
| ENSG00000176194.17 | 58.05536 | -0.476083225 | 0.1921 | -2.479 | 0.01318 | 0.0654 |
| ENSG00000177707.10 | 622.5153 | -0.24549147 | 0.0991 | -2.478 | 0.0132 | 0.0654 |
| ENSG00000213859.4 | 80.81335 | 0.392994418 | 0.1585 | 2.479 | 0.01318 | 0.0654 |
| ENSG00000241935.8 | 77.4791 | 0.386871882 | 0.1561 | 2.4786 | 0.01319 | 0.0654 |
| ENSG00000118596.11 | 657.2712 | -0.271787161 | 0.1097 | -2.477 | 0.01324 | 0.0655 |
| ENSG00000236397.3 | 18.0971 | 0.48967004 | 0.1977 | 2.4768 | 0.01326 | 0.0656 |
| ENSG00000108395.13 | 4301.739 | -0.297364824 | 0.1201 | -2.476 | 0.01329 | 0.0656 |
| ENSG00000138663.8 | 1352.269 | -0.255187121 | 0.1031 | -2.476 | 0.01329 | 0.0656 |
| ENSG00000180720.7 | 179.0174 | -0.442822785 | 0.1788 | -2.476 | 0.01327 | 0.0656 |
| ENSG00000258430.1 | 44.57635 | 1.305414645 | 0.5272 | 2.4759 | 0.01329 | 0.0656 |
| ENSG00000082781.11 | 595.3066 | 0.366303217 | 0.148 | 2.4751 | 0.01332 | 0.0657 |
| ENSG00000260230.2 | 6149.644 | -0.303546945 | 0.1226 | -2.475 | 0.01331 | 0.0657 |
| ENSG00000278192.1 | 36.96929 | 0.320007236 | 0.1293 | 2.4744 | 0.01334 | 0.0658 |
| ENSG00000147027.3 | 1794.175 | 0.284130742 | 0.1148 | 2.474 | 0.01336 | 0.0659 |
| ENSG00000163132.6 | 235.0998 | 0.400851143 | 0.1621 | 2.4734 | 0.01338 | 0.0659 |
| ENSG00000283171.1 | 39.33396 | 0.529968214 | 0.2143 | 2.4732 | 0.01339 | 0.066 |
| ENSG00000168792.4 | 80.81217 | 0.334249153 | 0.1352 | 2.4726 | 0.01341 | 0.066 |
| ENSG00000260518.1 | 19.02341 | -0.406521412 | 0.1644 | -2.473 | 0.01341 | 0.066 |
| ENSG00000278445.1 | 11.03744 | 0.495020816 | 0.2002 | 2.4726 | 0.01341 | 0.066 |
| ENSG00000120437.8 | 837.1367 | -0.240887965 | 0.0975 | -2.471 | 0.01347 | 0.0662 |
| ENSG00000224165.5 | 131.9763 | 0.160370632 | 0.0649 | 2.4706 | 0.01349 | 0.0663 |
| ENSG00000078098.13 | 71.15699 | -0.313754651 | 0.127 | -2.47 | 0.0135 | 0.0663 |
| ENSG00000165731.17 | 243.77 | -0.286900357 | 0.1162 | -2.47 | 0.01351 | 0.0663 |
| ENSG00000196581.10 | 2497.522 | -0.275212819 | 0.1114 | -2.469 | 0.01353 | 0.0664 |
| ENSG00000226174.6 | 20.95686 | 0.342488063 | 0.1387 | 2.4686 | 0.01356 | 0.0665 |
| ENSG00000073792.15 | 34.38212 | 0.399376161 | 0.1618 | 2.468 | 0.01359 | 0.0666 |
| ENSG00000118972.1 | 3384.164 | 1.001075747 | 0.4056 | 2.4681 | 0.01358 | 0.0666 |
| ENSG00000086717.18 | 298.5459 | -0.326005854 | 0.1321 | -2.468 | 0.01359 | 0.0666 |
| ENSG00000158528.11 | 2731.545 | -0.243506906 | 0.0987 | -2.467 | 0.01361 | 0.0666 |
| ENSG00000129534.13 | 187.0496 | -0.236070167 | 0.0957 | -2.467 | 0.01362 | 0.0666 |
| ENSG00000105357.15 | 1260.253 | 0.224889639 | 0.0912 | 2.4662 | 0.01366 | 0.0668 |
| ENSG00000177875.4 | 1233.015 | -0.288063517 | 0.1168 | -2.466 | 0.01368 | 0.0668 |
| ENSG00000233822.4 | 39.05306 | -0.245884788 | 0.0997 | -2.465 | 0.01368 | 0.0669 |
| ENSG00000067141.16 | 1584.956 | 0.191396209 | 0.0777 | 2.4648 | 0.01371 | 0.0669 |
| ENSG00000127334.10 | 516.0407 | -0.268080572 | 0.1088 | -2.465 | 0.01371 | 0.0669 |
| ENSG00000132294.14 | 4003.046 | -0.269700063 | 0.1094 | -2.464 | 0.01372 | 0.067 |
| ENSG00000259726.1 | 137.1681 | -0.3628074 | 0.1472 | -2.464 | 0.01374 | 0.067 |
| ENSG00000166313.18 | 7561.9 | -0.247920992 | 0.1006 | -2.464 | 0.01375 | 0.067 |
| ENSG00000115468.11 | 1127.166 | 0.320073423 | 0.1299 | 2.4632 | 0.01377 | 0.0671 |
| ENSG00000148680.15 | 95.32125 | -0.388402195 | 0.1577 | -2.463 | 0.01377 | 0.0671 |
| ENSG00000089101.17 | 62.01655 | -0.295556045 | 0.12 | -2.463 | 0.01379 | 0.0671 |
| ENSG00000261796.1 | 287.7864 | -0.27001311 | 0.1097 | -2.462 | 0.01382 | 0.0672 |
| ENSG00000168418.7 | 29.35208 | 0.348620371 | 0.1416 | 2.4615 | 0.01383 | 0.0673 |
| ENSG00000172071.11 | 357.7722 | -0.211704698 | 0.086 | -2.461 | 0.01385 | 0.0673 |
| ENSG00000165923.15 | 12.46009 | 0.464538005 | 0.1888 | 2.4609 | 0.01386 | 0.0673 |
| ENSG00000153012.11 | 432.5475 | -0.371533064 | 0.151 | -2.461 | 0.01387 | 0.0673 |
| ENSG00000143061.17 | 543.7284 | -0.313467665 | 0.1274 | -2.46 | 0.01389 | 0.0674 |
| ENSG00000181544.13 | 13.30905 | -0.380955387 | 0.1548 | -2.46 | 0.01389 | 0.0674 |
| ENSG00000265452.1 | 73.06968 | 0.311586844 | 0.1267 | 2.4593 | 0.01392 | 0.0675 |
| ENSG00000072840.12 | 119.6432 | 0.44038962 | 0.1791 | 2.4588 | 0.01394 | 0.0676 |
| ENSG00000211574.1 | 544.0252 | -0.243053307 | 0.0989 | -2.458 | 0.01396 | 0.0676 |
| ENSG00000233024.7 | 1612.251 | 0.35281926 | 0.1436 | 2.4577 | 0.01398 | 0.0677 |
| ENSG00000198478.7 | 2159.147 | -0.263632409 | 0.1073 | -2.457 | 0.01401 | 0.0678 |
| ENSG00000172725.13 | 471.4906 | 0.220221134 | 0.0896 | 2.4567 | 0.01402 | 0.0678 |
| ENSG00000181215.13 | 610.1286 | -0.252892676 | 0.103 | -2.455 | 0.01409 | 0.0681 |
| ENSG00000145416.13 | 778.976 | -0.253563439 | 0.1033 | -2.454 | 0.01411 | 0.0682 |
| ENSG00000068137.14 | 228.3325 | 0.239909862 | 0.0978 | 2.4542 | 0.01412 | 0.0682 |
| ENSG00000240764.3 | 2009.299 | -0.468688055 | 0.1911 | -2.453 | 0.01416 | 0.0684 |
| ENSG00000167969.12 | 407.3259 | 0.201182272 | 0.082 | 2.4527 | 0.01418 | 0.0684 |
| ENSG00000224223.1 | 169.019 | -0.307846963 | 0.1255 | -2.452 | 0.0142 | 0.0684 |
| ENSG00000227036.6 | 274.1033 | 0.204309879 | 0.0833 | 2.4522 | 0.0142 | 0.0684 |
| ENSG00000266852.2 | 9.109959 | -0.546215443 | 0.2227 | -2.452 | 0.01419 | 0.0684 |
| ENSG00000089818.16 | 4264.358 | -0.243625116 | 0.0994 | -2.451 | 0.01423 | 0.0686 |
| ENSG00000172037.13 | 797.198 | 0.35416114 | 0.1445 | 2.4508 | 0.01425 | 0.0686 |
| ENSG00000185811.16 | 55.16655 | -0.750514365 | 0.3063 | -2.45 | 0.01427 | 0.0687 |
| ENSG00000169116.11 | 2568.486 | -0.365644272 | 0.1492 | -2.45 | 0.01429 | 0.0687 |
| ENSG00000205488.8 | 23.97425 | -0.495388069 | 0.2022 | -2.45 | 0.01429 | 0.0687 |
| ENSG00000110975.8 | 144.5499 | -0.477367933 | 0.1949 | -2.45 | 0.0143 | 0.0687 |
| ENSG00000260804.3 | 1631.886 | -0.232385404 | 0.0949 | -2.45 | 0.0143 | 0.0687 |
| ENSG00000169891.17 | 3962.798 | -0.278332847 | 0.1136 | -2.449 | 0.01432 | 0.0687 |
| ENSG00000100614.17 | 3740.063 | -0.224656153 | 0.0918 | -2.448 | 0.01436 | 0.0689 |
| ENSG00000103056.11 | 977.2852 | -0.240592382 | 0.0983 | -2.448 | 0.01437 | 0.0689 |
| ENSG00000173457.10 | 205.6503 | 0.276954751 | 0.1132 | 2.4473 | 0.01439 | 0.069 |
| ENSG00000175497.16 | 2449.382 | -0.270516028 | 0.1106 | -2.445 | 0.01447 | 0.0693 |
| ENSG00000139946.9 | 289.3336 | 0.240855523 | 0.0985 | 2.4451 | 0.01448 | 0.0693 |
| ENSG00000250770.3 | 125.7261 | -0.263382492 | 0.1077 | -2.445 | 0.01448 | 0.0693 |
| ENSG00000108018.15 | 1055.625 | -0.257829593 | 0.1055 | -2.444 | 0.01451 | 0.0694 |
| ENSG00000154102.10 | 45.24595 | 0.354967517 | 0.1452 | 2.444 | 0.01453 | 0.0694 |
| ENSG00000172568.4 | 431.8666 | -0.266267335 | 0.1089 | -2.444 | 0.01452 | 0.0694 |
| ENSG00000180535.3 | 16.4084 | -0.421616729 | 0.1725 | -2.444 | 0.01452 | 0.0694 |
| ENSG00000210144.1 | 10.54738 | 0.938549134 | 0.384 | 2.4444 | 0.01451 | 0.0694 |
| ENSG00000078237.6 | 257.2705 | -0.231465945 | 0.0948 | -2.443 | 0.01458 | 0.0696 |
| ENSG00000165804.15 | 927.3202 | 0.290817317 | 0.1191 | 2.4418 | 0.01462 | 0.0698 |
| ENSG00000048540.14 | 10918.36 | -0.239640052 | 0.0982 | -2.441 | 0.01463 | 0.0698 |
| ENSG00000258684.2 | 38.08319 | -0.394979255 | 0.1618 | -2.441 | 0.01466 | 0.0699 |
| ENSG00000119125.16 | 7685.348 | -0.365637301 | 0.1499 | -2.44 | 0.0147 | 0.07 |
| ENSG00000132975.7 | 591.9676 | -0.286745204 | 0.1175 | -2.44 | 0.0147 | 0.07 |
| ENSG00000196872.11 | 2725.142 | -0.203335183 | 0.0833 | -2.44 | 0.0147 | 0.07 |
| ENSG00000123500.9 | 49.64644 | -0.388971775 | 0.1595 | -2.439 | 0.01471 | 0.07 |
| ENSG00000254260.1 | 9.296684 | -0.649665059 | 0.2663 | -2.439 | 0.01472 | 0.07 |
| ENSG00000152932.7 | 8368.068 | -0.386062893 | 0.1583 | -2.439 | 0.01472 | 0.07 |
| ENSG00000164796.17 | 962.7931 | -0.174565832 | 0.0716 | -2.439 | 0.01474 | 0.0701 |
| ENSG00000143499.13 | 1208.942 | -0.285810664 | 0.1172 | -2.438 | 0.01475 | 0.0701 |
| ENSG00000089639.10 | 195.5714 | -0.238758825 | 0.0979 | -2.438 | 0.01478 | 0.0701 |
| ENSG00000188674.10 | 904.4185 | -0.242534535 | 0.0995 | -2.438 | 0.01478 | 0.0701 |
| ENSG00000138378.17 | 494.6876 | -0.322062447 | 0.1321 | -2.437 | 0.0148 | 0.0702 |
| ENSG00000144285.16 | 2046.317 | -0.330423413 | 0.1356 | -2.437 | 0.01482 | 0.0703 |
| ENSG00000234851.4 | 981.3982 | 0.223396775 | 0.0917 | 2.4366 | 0.01483 | 0.0703 |
| ENSG00000180440.3 | 1188.84 | -0.4195002 | 0.1722 | -2.436 | 0.01484 | 0.0703 |
| ENSG00000083857.13 | 1259.177 | 0.342571799 | 0.1407 | 2.4352 | 0.01488 | 0.0704 |
| ENSG00000149654.9 | 1711.76 | -0.281104578 | 0.1154 | -2.435 | 0.01488 | 0.0704 |
| ENSG00000163637.11 | 2844.924 | -0.257221779 | 0.1056 | -2.435 | 0.01488 | 0.0704 |
| ENSG00000080824.18 | 36272.72 | -0.256537521 | 0.1054 | -2.435 | 0.0149 | 0.0704 |
| ENSG00000129993.14 | 498.286 | -0.206655342 | 0.0849 | -2.434 | 0.01493 | 0.0705 |
| ENSG00000163898.9 | 47.47457 | -0.275045309 | 0.113 | -2.434 | 0.01493 | 0.0705 |
| ENSG00000111344.11 | 1504.01 | -0.26525239 | 0.109 | -2.434 | 0.01495 | 0.0706 |
| ENSG00000165899.10 | 292.501 | -0.333693869 | 0.1371 | -2.434 | 0.01495 | 0.0706 |
| ENSG00000023171.15 | 3525.406 | -0.233448048 | 0.096 | -2.433 | 0.01499 | 0.0707 |
| ENSG00000226752.8 | 176.6065 | -0.309443714 | 0.1272 | -2.432 | 0.015 | 0.0707 |
| ENSG00000102524.11 | 213.0584 | 0.402040067 | 0.1653 | 2.4322 | 0.01501 | 0.0707 |
| ENSG00000114455.13 | 10.47586 | 0.410912176 | 0.169 | 2.4319 | 0.01502 | 0.0708 |
| ENSG00000185567.6 | 1666.163 | -0.346561994 | 0.1425 | -2.432 | 0.01504 | 0.0708 |
| ENSG00000112137.17 | 6386.615 | -0.249430265 | 0.1026 | -2.431 | 0.01504 | 0.0708 |
| ENSG00000125458.6 | 176.5974 | 0.222189378 | 0.0914 | 2.4304 | 0.01508 | 0.0709 |
| ENSG00000237399.7 | 52.12606 | -0.299181109 | 0.1232 | -2.429 | 0.01513 | 0.0711 |
| ENSG00000124602.9 | 17.5074 | 0.414365574 | 0.1706 | 2.4289 | 0.01514 | 0.0712 |
| ENSG00000081320.10 | 214.8753 | 0.478110201 | 0.1969 | 2.4276 | 0.0152 | 0.0714 |
| ENSG00000100441.9 | 494.977 | 0.25290948 | 0.1042 | 2.4274 | 0.01521 | 0.0714 |
| ENSG00000145864.12 | 6742.14 | -0.293225287 | 0.1208 | -2.427 | 0.01521 | 0.0714 |
| ENSG00000198416.9 | 140.8036 | -0.299017009 | 0.1232 | -2.428 | 0.01519 | 0.0714 |
| ENSG00000164742.14 | 9167.25 | -0.325000853 | 0.1339 | -2.427 | 0.01524 | 0.0715 |
| ENSG00000196167.9 | 148.5302 | 0.203115096 | 0.0837 | 2.4261 | 0.01526 | 0.0715 |
| ENSG00000251637.6 | 268.5613 | 0.269713187 | 0.1112 | 2.4261 | 0.01526 | 0.0715 |
| ENSG00000253161.5 | 20.66463 | -0.318809604 | 0.1314 | -2.426 | 0.01527 | 0.0715 |
| ENSG00000174680.9 | 11.41146 | 0.491254386 | 0.2026 | 2.4253 | 0.0153 | 0.0716 |
| ENSG00000280776.1 | 92.92357 | -0.30956116 | 0.1276 | -2.425 | 0.01529 | 0.0716 |
| ENSG00000233176.3 | 9.92578 | -0.765852018 | 0.3159 | -2.424 | 0.01534 | 0.0717 |
| ENSG00000131153.8 | 45.29991 | 0.282784533 | 0.1167 | 2.4228 | 0.0154 | 0.072 |
| ENSG00000109654.14 | 9027.864 | -0.220525574 | 0.0911 | -2.422 | 0.01544 | 0.0721 |
| ENSG00000278616.1 | 103.8758 | -0.392825319 | 0.1622 | -2.421 | 0.01546 | 0.0722 |
| ENSG00000050438.16 | 2760.587 | -0.256298955 | 0.1059 | -2.42 | 0.01554 | 0.0724 |
| ENSG00000102181.20 | 5009.475 | -0.238115733 | 0.0984 | -2.42 | 0.01554 | 0.0724 |
| ENSG00000123636.17 | 1218.006 | 0.212091284 | 0.0877 | 2.4197 | 0.01553 | 0.0724 |
| ENSG00000177570.13 | 1599.473 | -0.326516924 | 0.1349 | -2.42 | 0.01554 | 0.0724 |
| ENSG00000148143.12 | 713.0801 | 0.199244394 | 0.0824 | 2.4187 | 0.01557 | 0.0725 |
| ENSG00000153233.12 | 1012.269 | -0.323386905 | 0.1337 | -2.419 | 0.01557 | 0.0725 |
| ENSG00000259577.1 | 73.89282 | -0.253660029 | 0.1049 | -2.418 | 0.0156 | 0.0726 |
| ENSG00000198242.13 | 2415.137 | 0.192720314 | 0.0797 | 2.4176 | 0.01562 | 0.0727 |
| ENSG00000121989.14 | 758.5215 | -0.218484824 | 0.0904 | -2.417 | 0.01564 | 0.0728 |
| ENSG00000106477.18 | 622.9891 | -0.19736721 | 0.0817 | -2.416 | 0.0157 | 0.073 |
| ENSG00000160766.14 | 464.3604 | -0.253971208 | 0.1051 | -2.416 | 0.01571 | 0.073 |
| ENSG00000103485.17 | 394.558 | 0.189605633 | 0.0785 | 2.415 | 0.01574 | 0.0731 |
| ENSG00000132481.6 | 359.8002 | 0.407419885 | 0.1687 | 2.4148 | 0.01574 | 0.0731 |
| ENSG00000207650.1 | 9.27368 | 0.527211425 | 0.2183 | 2.4146 | 0.01575 | 0.0731 |
| ENSG00000184388.5 | 380.3852 | -0.27631246 | 0.1145 | -2.413 | 0.01581 | 0.0733 |
| ENSG00000251621.1 | 26.31988 | -0.455899494 | 0.1889 | -2.413 | 0.01581 | 0.0733 |
| ENSG00000140988.15 | 3305.595 | 0.219935342 | 0.0911 | 2.4129 | 0.01583 | 0.0733 |
| ENSG00000015676.17 | 5770.497 | -0.237361032 | 0.0984 | -2.412 | 0.01588 | 0.0735 |
| ENSG00000189143.9 | 30.48384 | -0.328076922 | 0.1361 | -2.411 | 0.01589 | 0.0735 |
| ENSG00000230051.1 | 18.65751 | -0.685651045 | 0.2843 | -2.412 | 0.01588 | 0.0735 |
| ENSG00000243806.1 | 32.81646 | 0.4792854 | 0.1987 | 2.4118 | 0.01587 | 0.0735 |
| ENSG00000008283.15 | 1390.266 | -0.245755256 | 0.1019 | -2.411 | 0.0159 | 0.0735 |
| ENSG00000260328.1 | 510.4964 | -0.318818961 | 0.1322 | -2.411 | 0.01591 | 0.0735 |
| ENSG00000164031.16 | 2687.038 | -0.231550563 | 0.0961 | -2.411 | 0.01592 | 0.0736 |
| ENSG00000170442.11 | 38.98336 | -0.353289455 | 0.1466 | -2.41 | 0.01597 | 0.0737 |
| ENSG00000267462.1 | 67.91849 | 0.26145663 | 0.1085 | 2.4091 | 0.01599 | 0.0738 |
| ENSG00000170776.20 | 1278.274 | -0.167585294 | 0.0696 | -2.409 | 0.01601 | 0.0739 |
| ENSG00000226435.10 | 83.56024 | -0.273990195 | 0.1138 | -2.408 | 0.01603 | 0.0739 |
| ENSG00000187922.13 | 13.65545 | 0.691879921 | 0.2874 | 2.4075 | 0.01606 | 0.074 |
| ENSG00000148600.14 | 866.1674 | -0.297143025 | 0.1234 | -2.407 | 0.01607 | 0.074 |
| ENSG00000084444.13 | 2466.708 | -0.300132071 | 0.1247 | -2.406 | 0.01611 | 0.0741 |
| ENSG00000136531.14 | 8497.94 | -0.320089103 | 0.133 | -2.407 | 0.0161 | 0.0741 |
| ENSG00000185070.10 | 2802.834 | -0.178534699 | 0.0742 | -2.406 | 0.01611 | 0.0741 |
| ENSG00000250506.6 | 72.04143 | 0.288794677 | 0.12 | 2.4062 | 0.01612 | 0.0741 |
| ENSG00000164112.12 | 1321.41 | -0.386798551 | 0.1608 | -2.405 | 0.01616 | 0.0743 |
| ENSG00000128268.11 | 3078.172 | -0.226077302 | 0.094 | -2.405 | 0.01619 | 0.0744 |
| ENSG00000176076.7 | 30.00213 | 0.462530776 | 0.1924 | 2.4045 | 0.0162 | 0.0744 |
| ENSG00000086289.11 | 2230.279 | -0.26490697 | 0.1102 | -2.404 | 0.01623 | 0.0745 |
| ENSG00000166340.15 | 2745.352 | 0.266151353 | 0.1107 | 2.4037 | 0.01623 | 0.0745 |
| ENSG00000134709.10 | 867.9704 | -0.234486267 | 0.0976 | -2.403 | 0.01627 | 0.0746 |
| ENSG00000257151.1 | 6937.99 | -0.235057488 | 0.0978 | -2.403 | 0.01627 | 0.0746 |
| ENSG00000066739.11 | 1926.701 | -0.203550427 | 0.0847 | -2.402 | 0.01628 | 0.0746 |
| ENSG00000071553.16 | 4490.992 | -0.22007624 | 0.0916 | -2.402 | 0.01629 | 0.0746 |
| ENSG00000117519.15 | 2551.717 | 0.370817564 | 0.1544 | 2.4023 | 0.01629 | 0.0746 |
| ENSG00000163794.6 | 91.75336 | 0.259899063 | 0.1082 | 2.4019 | 0.01631 | 0.0746 |
| ENSG00000279917.1 | 30.7949 | 0.389444047 | 0.1622 | 2.4016 | 0.01632 | 0.0747 |
| ENSG00000166317.11 | 38.73618 | 0.234677217 | 0.0978 | 2.4007 | 0.01636 | 0.0748 |
| ENSG00000105483.16 | 353.6329 | 0.217657526 | 0.0907 | 2.4003 | 0.01638 | 0.0749 |
| ENSG00000081189.14 | 11846.29 | -0.290201915 | 0.121 | -2.399 | 0.01646 | 0.0751 |
| ENSG00000140284.10 | 115.5388 | -0.261073573 | 0.1089 | -2.398 | 0.01648 | 0.0751 |
| ENSG00000170153.10 | 1753.685 | -0.254153511 | 0.106 | -2.398 | 0.01647 | 0.0751 |
| ENSG00000222018.1 | 73.14953 | -0.301074438 | 0.1255 | -2.398 | 0.01647 | 0.0751 |
| ENSG00000224881.1 | 91.68402 | -0.291231671 | 0.1214 | -2.398 | 0.01648 | 0.0751 |
| ENSG00000254852.8 | 29.66207 | 0.681457576 | 0.2841 | 2.3985 | 0.01646 | 0.0751 |
| ENSG00000186369.10 | 817.5436 | -0.275446327 | 0.1149 | -2.398 | 0.0165 | 0.0751 |
| ENSG00000278134.1 | 39.36696 | 0.400236979 | 0.1669 | 2.3977 | 0.0165 | 0.0751 |
| ENSG00000071859.14 | 1855.003 | 0.439864189 | 0.1835 | 2.3973 | 0.01652 | 0.0752 |
| ENSG00000110693.16 | 563.6399 | 0.299439564 | 0.1249 | 2.3968 | 0.01654 | 0.0753 |
| ENSG00000198626.15 | 3315.9 | -0.308305488 | 0.1286 | -2.397 | 0.01654 | 0.0753 |
| ENSG00000113805.8 | 1251.721 | -0.234404652 | 0.0978 | -2.396 | 0.01657 | 0.0753 |
| ENSG00000119737.5 | 294.1744 | 0.289078149 | 0.1207 | 2.3955 | 0.0166 | 0.0753 |
| ENSG00000158966.13 | 987.8765 | 0.205936305 | 0.086 | 2.3957 | 0.01659 | 0.0753 |
| ENSG00000227473.1 | 44.6849 | -0.370463272 | 0.1546 | -2.396 | 0.0166 | 0.0753 |
| ENSG00000248485.1 | 809.7439 | -0.310380594 | 0.1296 | -2.396 | 0.01659 | 0.0753 |
| ENSG00000119771.14 | 823.0165 | -0.230542046 | 0.0963 | -2.395 | 0.01662 | 0.0753 |
| ENSG00000163635.17 | 676.3649 | -0.140454415 | 0.0587 | -2.395 | 0.01663 | 0.0753 |
| ENSG00000166006.12 | 3861.912 | -0.314175225 | 0.1312 | -2.395 | 0.01663 | 0.0753 |
| ENSG00000279026.1 | 172.2988 | 0.253170089 | 0.1057 | 2.3949 | 0.01663 | 0.0753 |
| ENSG00000105851.10 | 46.7267 | -0.368314886 | 0.1539 | -2.394 | 0.01667 | 0.0754 |
| ENSG00000113638.13 | 622.4085 | -0.269073985 | 0.1124 | -2.394 | 0.01667 | 0.0754 |
| ENSG00000135318.11 | 520.9579 | 0.414572599 | 0.1732 | 2.3937 | 0.01668 | 0.0754 |
| ENSG00000182109.7 | 146.2611 | 0.256469154 | 0.1072 | 2.3935 | 0.01669 | 0.0754 |
| ENSG00000183762.12 | 662.1529 | 0.213081918 | 0.089 | 2.3938 | 0.01668 | 0.0754 |
| ENSG00000228238.1 | 18.54065 | -0.42554593 | 0.1777 | -2.394 | 0.01666 | 0.0754 |
| ENSG00000274272.1 | 184.6787 | 0.261118002 | 0.1091 | 2.3936 | 0.01668 | 0.0754 |
| ENSG00000174516.14 | 1058.755 | -0.259497577 | 0.1084 | -2.393 | 0.0167 | 0.0754 |
| ENSG00000104899.6 | 93.54637 | 0.594503949 | 0.2486 | 2.3913 | 0.01679 | 0.0757 |
| ENSG00000186918.13 | 881.1164 | 0.175932652 | 0.0736 | 2.3914 | 0.01678 | 0.0757 |
| ENSG00000127325.18 | 91.72172 | 0.265412581 | 0.111 | 2.3909 | 0.01681 | 0.0758 |
| ENSG00000090975.12 | 2529.158 | -0.225893891 | 0.0945 | -2.39 | 0.01684 | 0.0759 |
| ENSG00000122966.14 | 4497.24 | -0.193107738 | 0.0808 | -2.39 | 0.01684 | 0.0759 |
| ENSG00000139192.11 | 111.1402 | 0.29750976 | 0.1245 | 2.3899 | 0.01685 | 0.0759 |
| ENSG00000272589.1 | 34.73368 | 0.324469798 | 0.1358 | 2.3887 | 0.01691 | 0.0761 |
| ENSG00000130287.13 | 6111.559 | 0.238876841 | 0.1 | 2.3885 | 0.01692 | 0.0761 |
| ENSG00000180113.15 | 227.4847 | -0.232431118 | 0.0973 | -2.388 | 0.01696 | 0.0763 |
| ENSG00000256407.2 | 13.36931 | 0.396441687 | 0.1661 | 2.3875 | 0.01697 | 0.0763 |
| ENSG00000166448.14 | 8410.922 | -0.267904775 | 0.1122 | -2.387 | 0.01699 | 0.0763 |
| ENSG00000255082.1 | 2070.904 | -0.18021028 | 0.0755 | -2.386 | 0.01701 | 0.0764 |
| ENSG00000279398.1 | 12.14684 | 0.539458939 | 0.2261 | 2.386 | 0.01703 | 0.0765 |
| ENSG00000164076.16 | 7375.014 | -0.267370342 | 0.1121 | -2.386 | 0.01705 | 0.0765 |
| ENSG00000283229.1 | 329.0048 | -0.782407484 | 0.328 | -2.386 | 0.01705 | 0.0765 |
| ENSG00000280332.1 | 96.14778 | 0.236864859 | 0.0993 | 2.3854 | 0.01706 | 0.0765 |
| ENSG00000280385.1 | 311.2865 | -0.324729575 | 0.1361 | -2.385 | 0.01707 | 0.0765 |
| ENSG00000097021.19 | 3534.308 | -0.258888354 | 0.1086 | -2.384 | 0.01711 | 0.0765 |
| ENSG00000100321.14 | 11577.72 | -0.274071632 | 0.115 | -2.384 | 0.01712 | 0.0765 |
| ENSG00000100916.13 | 1033.993 | -0.252077109 | 0.1057 | -2.384 | 0.01711 | 0.0765 |
| ENSG00000146006.7 | 2124.459 | -0.256325052 | 0.1075 | -2.384 | 0.0171 | 0.0765 |
| ENSG00000185339.8 | 258.129 | 0.274752566 | 0.1152 | 2.3841 | 0.01712 | 0.0765 |
| ENSG00000239775.1 | 13.02783 | 0.433083363 | 0.1816 | 2.3846 | 0.0171 | 0.0765 |
| ENSG00000114279.13 | 4254.174 | -0.281554927 | 0.1181 | -2.383 | 0.01716 | 0.0766 |
| ENSG00000171346.14 | 18.04954 | 0.455945779 | 0.1913 | 2.3835 | 0.01715 | 0.0766 |
| ENSG00000214894.6 | 16.08694 | 0.380559423 | 0.1597 | 2.3832 | 0.01716 | 0.0766 |
| ENSG00000197121.14 | 2307.534 | -0.260148308 | 0.1092 | -2.383 | 0.01718 | 0.0767 |
| ENSG00000147485.12 | 25.89914 | -0.292505359 | 0.1228 | -2.382 | 0.0172 | 0.0767 |
| ENSG00000245213.6 | 41.56087 | 0.266052945 | 0.1117 | 2.3825 | 0.0172 | 0.0767 |
| ENSG00000116329.10 | 849.7701 | -0.241114433 | 0.1013 | -2.381 | 0.01726 | 0.0769 |
| ENSG00000138653.9 | 15.72832 | -0.446028227 | 0.1873 | -2.381 | 0.01725 | 0.0769 |
| ENSG00000196353.11 | 1856.963 | -0.296401747 | 0.1245 | -2.381 | 0.01727 | 0.0769 |
| ENSG00000133687.15 | 1928.8 | -0.258351812 | 0.1086 | -2.38 | 0.01732 | 0.0771 |
| ENSG00000268055.1 | 15.59707 | 0.664729776 | 0.2793 | 2.3796 | 0.01733 | 0.0771 |
| ENSG00000216863.9 | 492.5114 | -0.292571109 | 0.123 | -2.379 | 0.01737 | 0.0772 |
| ENSG00000095203.14 | 2256.286 | -0.244005683 | 0.1026 | -2.378 | 0.01739 | 0.0773 |
| ENSG00000120549.17 | 1251.609 | -0.250196481 | 0.1052 | -2.378 | 0.01739 | 0.0773 |
| ENSG00000256538.1 | 22.70591 | -0.436767936 | 0.1837 | -2.378 | 0.01741 | 0.0773 |
| ENSG00000279487.1 | 12.13495 | 0.640230155 | 0.2693 | 2.3777 | 0.01742 | 0.0773 |
| ENSG00000175352.10 | 1694.616 | -0.289662278 | 0.1219 | -2.377 | 0.01746 | 0.0774 |
| ENSG00000223572.9 | 1652.941 | -0.259131535 | 0.1091 | -2.376 | 0.0175 | 0.0776 |
| ENSG00000272040.1 | 30.64698 | 0.307131523 | 0.1293 | 2.3761 | 0.0175 | 0.0776 |
| ENSG00000150627.15 | 1775.025 | -0.234021345 | 0.0985 | -2.376 | 0.01752 | 0.0776 |
| ENSG00000153933.9 | 1318.176 | -0.23451903 | 0.0987 | -2.376 | 0.01752 | 0.0776 |
| ENSG00000106211.8 | 2037.048 | 0.547560706 | 0.2305 | 2.3754 | 0.01753 | 0.0776 |
| ENSG00000139988.9 | 48.68339 | -0.389012466 | 0.1639 | -2.373 | 0.01763 | 0.078 |
| ENSG00000232040.2 | 102.9718 | -0.29109603 | 0.1227 | -2.373 | 0.01764 | 0.078 |
| ENSG00000255686.1 | 118.1729 | 0.341347922 | 0.1438 | 2.3732 | 0.01764 | 0.078 |
| ENSG00000269998.1 | 94.76025 | -0.273739388 | 0.1153 | -2.373 | 0.01762 | 0.078 |
| ENSG00000006534.15 | 102.5084 | 0.278914617 | 0.1176 | 2.3713 | 0.01772 | 0.0783 |
| ENSG00000249971.1 | 8.861657 | 2.167278806 | 0.914 | 2.3712 | 0.01773 | 0.0783 |
| ENSG00000266486.1 | 8.655496 | -0.555994376 | 0.2346 | -2.37 | 0.01778 | 0.0785 |
| ENSG00000143318.12 | 366.5249 | -0.259279275 | 0.1094 | -2.37 | 0.0178 | 0.0786 |
| ENSG00000248599.1 | 13.29679 | -0.623962202 | 0.2634 | -2.369 | 0.01783 | 0.0786 |
| ENSG00000151320.10 | 6048.245 | -0.247104237 | 0.1044 | -2.368 | 0.0179 | 0.0787 |
| ENSG00000166780.10 | 4734.701 | -0.232433854 | 0.0982 | -2.368 | 0.01789 | 0.0787 |
| ENSG00000167306.19 | 261.1266 | -0.343084132 | 0.1449 | -2.368 | 0.0179 | 0.0787 |
| ENSG00000169894.17 | 73.46762 | 0.397753926 | 0.1679 | 2.3684 | 0.01786 | 0.0787 |
| ENSG00000213424.8 | 884.912 | -0.296532338 | 0.1252 | -2.368 | 0.01789 | 0.0787 |
| ENSG00000242615.1 | 50.37023 | 0.233378509 | 0.0985 | 2.3688 | 0.01785 | 0.0787 |
| ENSG00000254726.2 | 41.82199 | 0.318470111 | 0.1345 | 2.3683 | 0.01787 | 0.0787 |
| ENSG00000258376.2 | 15.40813 | 0.473956187 | 0.2002 | 2.3678 | 0.01789 | 0.0787 |
| ENSG00000244257.5 | 1010.834 | -0.388928867 | 0.1644 | -2.366 | 0.01797 | 0.079 |
| ENSG00000081985.10 | 195.4649 | -0.367969999 | 0.1555 | -2.366 | 0.01799 | 0.079 |
| ENSG00000183682.7 | 70.18013 | -0.297388786 | 0.1257 | -2.365 | 0.01802 | 0.0791 |
| ENSG00000223443.2 | 22.13771 | 0.437684612 | 0.1851 | 2.3648 | 0.01804 | 0.0791 |
| ENSG00000239642.5 | 11.1452 | 0.416822811 | 0.1763 | 2.3648 | 0.01804 | 0.0791 |
| ENSG00000271275.1 | 31.57949 | -0.38662588 | 0.1636 | -2.364 | 0.01808 | 0.0793 |
| ENSG00000227811.2 | 46.56017 | -0.384760484 | 0.1628 | -2.363 | 0.01814 | 0.0795 |
| ENSG00000261426.1 | 16.80711 | -0.468430854 | 0.1984 | -2.361 | 0.01824 | 0.0799 |
| ENSG00000079950.13 | 2925.498 | -0.17146536 | 0.0727 | -2.36 | 0.01827 | 0.08 |
| ENSG00000187840.4 | 55.2759 | 0.279418687 | 0.1184 | 2.36 | 0.01827 | 0.08 |
| ENSG00000095932.6 | 40.19141 | -0.540187931 | 0.2289 | -2.36 | 0.01828 | 0.08 |
| ENSG00000173705.8 | 515.0184 | -0.245211646 | 0.1039 | -2.36 | 0.0183 | 0.08 |
| ENSG00000091831.21 | 73.67554 | -0.266371061 | 0.113 | -2.358 | 0.01837 | 0.0803 |
| ENSG00000022567.9 | 1039.534 | -0.23587072 | 0.1 | -2.358 | 0.01839 | 0.0803 |
| ENSG00000128596.16 | 1739.563 | -0.271911456 | 0.1153 | -2.357 | 0.01841 | 0.0804 |
| ENSG00000084207.15 | 1548.898 | 0.196476686 | 0.0834 | 2.3566 | 0.01844 | 0.0805 |
| ENSG00000126860.11 | 370.3236 | -0.466148901 | 0.1978 | -2.357 | 0.01845 | 0.0805 |
| ENSG00000148584.14 | 11.03224 | 0.392032434 | 0.1664 | 2.3566 | 0.01844 | 0.0805 |
| ENSG00000111052.7 | 1200.264 | -0.249062068 | 0.1057 | -2.356 | 0.01846 | 0.0805 |
| ENSG00000256725.1 | 29.13342 | -0.424821564 | 0.1803 | -2.356 | 0.01846 | 0.0805 |
| ENSG00000166689.15 | 182.0831 | 0.319103637 | 0.1355 | 2.3551 | 0.01852 | 0.0807 |
| ENSG00000147231.13 | 382.8776 | -0.237838675 | 0.101 | -2.355 | 0.01854 | 0.0807 |
| ENSG00000164219.9 | 424.6007 | 0.140604818 | 0.0597 | 2.3548 | 0.01854 | 0.0807 |
| ENSG00000263878.1 | 250.3886 | -0.323432855 | 0.1374 | -2.355 | 0.01854 | 0.0807 |
| ENSG00000277035.1 | 10.19159 | 0.476408658 | 0.2025 | 2.353 | 0.01862 | 0.081 |
| ENSG00000176105.13 | 567.4921 | 0.284691694 | 0.121 | 2.3526 | 0.01864 | 0.081 |
| ENSG00000100345.20 | 4382.708 | -0.238324018 | 0.1014 | -2.351 | 0.0187 | 0.0813 |
| ENSG00000113966.9 | 797.2073 | -0.265309655 | 0.1129 | -2.351 | 0.01873 | 0.0814 |
| ENSG00000128250.5 | 35.77208 | -0.557435775 | 0.2371 | -2.351 | 0.01874 | 0.0814 |
| ENSG00000133247.13 | 207.3447 | 0.211956069 | 0.0902 | 2.3502 | 0.01876 | 0.0814 |
| ENSG00000167772.11 | 272.9839 | 0.80308591 | 0.3417 | 2.3502 | 0.01876 | 0.0814 |
| ENSG00000263020.6 | 22.18167 | 1.002964434 | 0.4268 | 2.3501 | 0.01877 | 0.0814 |
| ENSG00000198077.10 | 24.64182 | 0.316252245 | 0.1346 | 2.3488 | 0.01883 | 0.0816 |
| ENSG00000272512.1 | 54.41744 | 0.630544106 | 0.2685 | 2.3486 | 0.01884 | 0.0816 |
| ENSG00000112419.14 | 712.338 | -0.22023292 | 0.0938 | -2.348 | 0.01885 | 0.0816 |
| ENSG00000102057.9 | 134.8662 | -0.20347946 | 0.0867 | -2.347 | 0.01894 | 0.0819 |
| ENSG00000137642.12 | 7243.871 | -0.231025872 | 0.0984 | -2.347 | 0.01894 | 0.0819 |
| ENSG00000260534.1 | 34.25026 | 0.857671703 | 0.3654 | 2.347 | 0.01893 | 0.0819 |
| ENSG00000265313.5 | 98.88334 | -0.389190361 | 0.1659 | -2.346 | 0.01896 | 0.082 |
| ENSG00000066468.21 | 1332.153 | 0.367996158 | 0.1569 | 2.3461 | 0.01897 | 0.082 |
| ENSG00000271723.5 | 169.805 | 0.269160844 | 0.1147 | 2.3459 | 0.01898 | 0.082 |
| ENSG00000112759.16 | 572.645 | -0.261452273 | 0.1115 | -2.345 | 0.019 | 0.0821 |
| ENSG00000175894.14 | 14.0215 | -0.476294803 | 0.2031 | -2.345 | 0.01903 | 0.0821 |
| ENSG00000100302.6 | 1192.501 | -0.256927416 | 0.1096 | -2.344 | 0.0191 | 0.0824 |
| ENSG00000140455.16 | 290.4067 | 0.247920006 | 0.1058 | 2.3431 | 0.01912 | 0.0825 |
| ENSG00000111371.15 | 4097.53 | -0.233137512 | 0.0995 | -2.343 | 0.01915 | 0.0825 |
| ENSG00000258733.5 | 24.48638 | 0.389479994 | 0.1663 | 2.3426 | 0.01915 | 0.0825 |
| ENSG00000279033.1 | 173.5864 | 0.480303351 | 0.205 | 2.3426 | 0.01915 | 0.0825 |
| ENSG00000232354.8 | 76.51766 | -0.25651177 | 0.1095 | -2.342 | 0.01917 | 0.0825 |
| ENSG00000276045.2 | 82.27953 | 0.259829695 | 0.1109 | 2.3422 | 0.01917 | 0.0825 |
| ENSG00000134333.13 | 6306.851 | -0.227367844 | 0.0971 | -2.342 | 0.01919 | 0.0825 |
| ENSG00000139874.5 | 1373.65 | -0.309517466 | 0.1322 | -2.342 | 0.01919 | 0.0825 |
| ENSG00000094804.9 | 125.8287 | -0.232760746 | 0.0994 | -2.341 | 0.01922 | 0.0825 |
| ENSG00000119042.16 | 2132.944 | -0.25292909 | 0.108 | -2.341 | 0.01924 | 0.0825 |
| ENSG00000135049.15 | 1791.365 | -0.26925538 | 0.115 | -2.341 | 0.01923 | 0.0825 |
| ENSG00000163412.12 | 2257.197 | -0.218089261 | 0.0932 | -2.341 | 0.01922 | 0.0825 |
| ENSG00000215218.3 | 2844.942 | -0.28810577 | 0.1231 | -2.341 | 0.01924 | 0.0825 |
| ENSG00000169255.13 | 1634.22 | -0.222796988 | 0.0953 | -2.339 | 0.01935 | 0.083 |
| ENSG00000275026.1 | 59.25647 | -0.315885035 | 0.1351 | -2.339 | 0.01935 | 0.083 |
| ENSG00000116690.12 | 14.23217 | 0.413619845 | 0.1769 | 2.3382 | 0.01938 | 0.083 |
| ENSG00000088881.20 | 391.2896 | 0.217357504 | 0.093 | 2.3379 | 0.01939 | 0.0831 |
| ENSG00000167291.15 | 1222.707 | 0.20790431 | 0.0889 | 2.3376 | 0.01941 | 0.0831 |
| ENSG00000128606.12 | 60.06941 | -0.267318192 | 0.1144 | -2.336 | 0.01947 | 0.0833 |
| ENSG00000134313.14 | 5041.074 | -0.191834779 | 0.0821 | -2.336 | 0.01947 | 0.0833 |
| ENSG00000157445.14 | 1158.906 | -0.255856362 | 0.1095 | -2.336 | 0.01947 | 0.0833 |
| ENSG00000013293.5 | 2882.503 | -0.287135438 | 0.1229 | -2.336 | 0.01949 | 0.0833 |
| ENSG00000103319.11 | 742.5065 | 0.201594288 | 0.0863 | 2.3358 | 0.0195 | 0.0833 |
| ENSG00000123342.15 | 11.60999 | -0.679484681 | 0.2909 | -2.336 | 0.01952 | 0.0834 |
| ENSG00000112559.13 | 47.49738 | 0.428463092 | 0.1835 | 2.3346 | 0.01957 | 0.0834 |
| ENSG00000143190.21 | 1320.485 | 0.176158017 | 0.0754 | 2.3353 | 0.01953 | 0.0834 |
| ENSG00000166473.17 | 153.1362 | 0.197822908 | 0.0847 | 2.3348 | 0.01955 | 0.0834 |
| ENSG00000185164.14 | 5037.815 | -0.254250702 | 0.1089 | -2.335 | 0.01956 | 0.0834 |
| ENSG00000214491.8 | 125.934 | 0.204329519 | 0.0875 | 2.3346 | 0.01956 | 0.0834 |
| ENSG00000231528.2 | 47.71287 | 0.626558418 | 0.2683 | 2.3349 | 0.01955 | 0.0834 |
| ENSG00000139190.16 | 1200.708 | -0.289783018 | 0.1242 | -2.334 | 0.01959 | 0.0834 |
| ENSG00000108825.17 | 40.27644 | -0.392685416 | 0.1683 | -2.334 | 0.0196 | 0.0834 |
| ENSG00000157992.12 | 15.04447 | 0.337492997 | 0.1446 | 2.3336 | 0.01962 | 0.0835 |
| ENSG00000062096.14 | 46.32337 | 0.299602881 | 0.1284 | 2.3328 | 0.01966 | 0.0836 |
| ENSG00000101152.10 | 7657.783 | -0.234623957 | 0.1006 | -2.333 | 0.01966 | 0.0836 |
| ENSG00000175426.10 | 1741.644 | -0.414411328 | 0.1776 | -2.333 | 0.01965 | 0.0836 |
| ENSG00000170759.10 | 4936.834 | 0.208903932 | 0.0896 | 2.3324 | 0.01968 | 0.0836 |
| ENSG00000174482.10 | 237.5976 | -0.236373987 | 0.1014 | -2.332 | 0.01969 | 0.0836 |
| ENSG00000176771.15 | 146.4986 | 0.303321162 | 0.1301 | 2.3319 | 0.0197 | 0.0836 |
| ENSG00000159904.11 | 65.83123 | -0.303103654 | 0.13 | -2.332 | 0.01971 | 0.0836 |
| ENSG00000168490.13 | 13907.73 | -0.241555776 | 0.1036 | -2.331 | 0.01974 | 0.0837 |
| ENSG00000246763.6 | 44.75807 | -0.269528152 | 0.1156 | -2.331 | 0.01974 | 0.0837 |
| ENSG00000129221.15 | 11.31777 | 0.425570256 | 0.1826 | 2.33 | 0.0198 | 0.0839 |
| ENSG00000133216.16 | 411.9307 | -0.218643612 | 0.0938 | -2.33 | 0.0198 | 0.0839 |
| ENSG00000279571.2 | 75.32814 | -0.361979977 | 0.1554 | -2.33 | 0.01982 | 0.0839 |
| ENSG00000165895.17 | 166.8117 | 0.32078652 | 0.1377 | 2.3292 | 0.01985 | 0.084 |
| ENSG00000237973.1 | 327.6493 | -0.57733621 | 0.2479 | -2.329 | 0.01986 | 0.084 |
| ENSG00000254092.1 | 20.65612 | -0.4269483 | 0.1833 | -2.329 | 0.01987 | 0.084 |
| ENSG00000124479.8 | 374.5923 | 0.252613977 | 0.1085 | 2.3285 | 0.01989 | 0.0841 |
| ENSG00000129646.14 | 222.5058 | -0.197367211 | 0.0848 | -2.328 | 0.01991 | 0.0841 |
| ENSG00000261759.1 | 55.6153 | 0.344458058 | 0.148 | 2.328 | 0.01991 | 0.0841 |
| ENSG00000213625.8 | 861.6057 | 0.283714109 | 0.1219 | 2.3278 | 0.01992 | 0.0841 |
| ENSG00000183458.13 | 1279.29 | -0.253710694 | 0.109 | -2.328 | 0.01994 | 0.0841 |
| ENSG00000267532.4 | 100.4477 | 0.332782183 | 0.143 | 2.3272 | 0.01995 | 0.0842 |
| ENSG00000104812.14 | 628.737 | 0.143658799 | 0.0617 | 2.3268 | 0.01997 | 0.0842 |
| ENSG00000149485.17 | 2005.64 | 0.254927013 | 0.1096 | 2.3266 | 0.01998 | 0.0842 |
| ENSG00000268903.1 | 38.70527 | 0.575717321 | 0.2475 | 2.3265 | 0.01999 | 0.0842 |
| ENSG00000081181.7 | 265.7145 | -0.254111145 | 0.1092 | -2.326 | 0.02 | 0.0842 |
| ENSG00000125848.9 | 484.3335 | -0.291486294 | 0.1253 | -2.326 | 0.02003 | 0.0843 |
| ENSG00000164066.12 | 2056.619 | 0.146625442 | 0.063 | 2.3259 | 0.02003 | 0.0843 |
| ENSG00000150175.13 | 250.5845 | -0.542631005 | 0.2333 | -2.325 | 0.02005 | 0.0843 |
| ENSG00000176601.12 | 15.16257 | 0.4448426 | 0.1913 | 2.3255 | 0.02004 | 0.0843 |
| ENSG00000116985.11 | 569.0405 | 0.297650683 | 0.1281 | 2.3245 | 0.0201 | 0.0845 |
| ENSG00000248441.6 | 195.7048 | -0.2112224 | 0.0909 | -2.324 | 0.02011 | 0.0845 |
| ENSG00000075303.12 | 859.9429 | -0.234341383 | 0.1009 | -2.323 | 0.0202 | 0.0848 |
| ENSG00000130518.16 | 80.23081 | -0.271094056 | 0.1167 | -2.323 | 0.02019 | 0.0848 |
| ENSG00000117400.15 | 26.59948 | -0.349950581 | 0.1507 | -2.322 | 0.02022 | 0.0848 |
| ENSG00000279765.3 | 120.9469 | 0.266572358 | 0.1148 | 2.3216 | 0.02026 | 0.085 |
| ENSG00000158445.9 | 5625.556 | -0.233792469 | 0.1007 | -2.321 | 0.02029 | 0.0851 |
| ENSG00000166402.8 | 3813.326 | -0.194534073 | 0.0839 | -2.319 | 0.02042 | 0.0856 |
| ENSG00000263400.6 | 19.93677 | 0.425266235 | 0.1835 | 2.3172 | 0.0205 | 0.0859 |
| ENSG00000137409.19 | 11199.08 | -0.219753577 | 0.0948 | -2.317 | 0.02051 | 0.0859 |
| ENSG00000233766.7 | 16.5046 | -0.32202802 | 0.139 | -2.316 | 0.02054 | 0.086 |
| ENSG00000117013.14 | 256.1568 | -0.240694739 | 0.1039 | -2.316 | 0.02057 | 0.0861 |
| ENSG00000228253.1 | 6829.03 | 0.299082211 | 0.1292 | 2.3155 | 0.02059 | 0.0861 |
| ENSG00000114480.12 | 336.6512 | -0.208061211 | 0.0899 | -2.315 | 0.0206 | 0.0861 |
| ENSG00000196549.10 | 47.34585 | -0.383661681 | 0.1658 | -2.314 | 0.02066 | 0.0863 |
| ENSG00000100842.12 | 467.4201 | 0.301923389 | 0.1305 | 2.3137 | 0.02069 | 0.0864 |
| ENSG00000270681.1 | 29.95894 | 0.308374539 | 0.1333 | 2.3135 | 0.0207 | 0.0864 |
| ENSG00000023516.8 | 9117.158 | -0.276566993 | 0.1196 | -2.313 | 0.02072 | 0.0865 |
| ENSG00000160999.10 | 187.408 | 0.257888355 | 0.1115 | 2.3126 | 0.02074 | 0.0866 |
| ENSG00000171246.5 | 11657.18 | -0.329711245 | 0.1426 | -2.312 | 0.02076 | 0.0866 |
| ENSG00000267023.5 | 201.5638 | 0.141979441 | 0.0614 | 2.3122 | 0.02077 | 0.0866 |
| ENSG00000111321.10 | 166.4522 | 0.312436265 | 0.1352 | 2.3117 | 0.02079 | 0.0867 |
| ENSG00000215146.4 | 207.9555 | -0.31081241 | 0.1345 | -2.311 | 0.02082 | 0.0867 |
| ENSG00000173227.13 | 1466.188 | -0.218470913 | 0.0945 | -2.311 | 0.02084 | 0.0868 |
| ENSG00000180667.10 | 505.4433 | -0.227258105 | 0.0984 | -2.31 | 0.02087 | 0.0868 |
| ENSG00000262873.1 | 31.11265 | -0.300332011 | 0.13 | -2.31 | 0.02087 | 0.0868 |
| ENSG00000185477.4 | 592.1216 | -0.269807205 | 0.1168 | -2.31 | 0.02089 | 0.0869 |
| ENSG00000165985.9 | 2049.976 | -0.471921511 | 0.2043 | -2.31 | 0.0209 | 0.0869 |
| ENSG00000161270.19 | 39.13211 | -0.333169241 | 0.1443 | -2.309 | 0.02096 | 0.0871 |
| ENSG00000173801.16 | 309.4592 | 0.25308984 | 0.1096 | 2.3086 | 0.02096 | 0.0871 |
| ENSG00000261390.5 | 13.133 | 0.381589239 | 0.1653 | 2.3078 | 0.02101 | 0.0872 |
| ENSG00000276231.4 | 19.83581 | -0.384486248 | 0.1666 | -2.308 | 0.02102 | 0.0872 |
| ENSG00000067992.12 | 283.8846 | -0.355228812 | 0.154 | -2.307 | 0.02103 | 0.0873 |
| ENSG00000198857.3 | 37.95339 | -0.336715609 | 0.1459 | -2.307 | 0.02104 | 0.0873 |
| ENSG00000092036.17 | 149.5358 | 0.219941022 | 0.0954 | 2.3063 | 0.02109 | 0.0874 |
| ENSG00000172375.12 | 2144.681 | -0.200383659 | 0.0869 | -2.306 | 0.02108 | 0.0874 |
| ENSG00000104324.15 | 334.1492 | 0.263922812 | 0.1145 | 2.3055 | 0.02114 | 0.0876 |
| ENSG00000121871.3 | 578.879 | -0.230602197 | 0.1001 | -2.305 | 0.02119 | 0.0877 |
| ENSG00000272779.1 | 312.7667 | -0.307644119 | 0.1336 | -2.303 | 0.02126 | 0.088 |
| ENSG00000280350.1 | 14.62421 | 0.406354792 | 0.1765 | 2.3029 | 0.02128 | 0.0881 |
| ENSG00000132819.16 | 242.2218 | 0.287522349 | 0.1249 | 2.302 | 0.02134 | 0.0882 |
| ENSG00000137693.13 | 742.2893 | 0.410303494 | 0.1783 | 2.3012 | 0.02138 | 0.0884 |
| ENSG00000100146.16 | 868.6393 | 0.373811483 | 0.1625 | 2.3007 | 0.02141 | 0.0885 |
| ENSG00000164236.11 | 1438.077 | -0.261698703 | 0.1138 | -2.3 | 0.02145 | 0.0885 |
| ENSG00000205060.10 | 1483.967 | -0.213415247 | 0.0928 | -2.3 | 0.02144 | 0.0885 |
| ENSG00000277744.1 | 18.33953 | 0.373444221 | 0.1624 | 2.3 | 0.02145 | 0.0885 |
| ENSG00000122877.14 | 186.9733 | -0.538944649 | 0.2344 | -2.299 | 0.02148 | 0.0886 |
| ENSG00000164611.12 | 74.30513 | -0.28679683 | 0.1247 | -2.299 | 0.02149 | 0.0886 |
| ENSG00000135541.20 | 1908.524 | -0.185823208 | 0.0809 | -2.298 | 0.02158 | 0.0887 |
| ENSG00000140545.14 | 1156.151 | 0.426546864 | 0.1856 | 2.2978 | 0.02157 | 0.0887 |
| ENSG00000162706.12 | 13501.42 | -0.223368495 | 0.0972 | -2.298 | 0.02158 | 0.0887 |
| ENSG00000164111.14 | 1175.697 | 0.321661811 | 0.14 | 2.298 | 0.02156 | 0.0887 |
| ENSG00000167434.9 | 363.2407 | -0.25404293 | 0.1105 | -2.298 | 0.02155 | 0.0887 |
| ENSG00000171608.15 | 400.591 | -0.192358411 | 0.0837 | -2.298 | 0.02154 | 0.0887 |
| ENSG00000177034.14 | 1482.192 | -0.19562117 | 0.0851 | -2.298 | 0.02158 | 0.0887 |
| ENSG00000205923.3 | 101.4889 | 0.403704504 | 0.1756 | 2.2987 | 0.02152 | 0.0887 |
| ENSG00000251442.5 | 184.712 | 0.571901113 | 0.2489 | 2.2981 | 0.02156 | 0.0887 |
| ENSG00000050030.13 | 1013.278 | -0.283628332 | 0.1235 | -2.297 | 0.02163 | 0.0888 |
| ENSG00000136750.11 | 2730.688 | -0.353394006 | 0.1539 | -2.296 | 0.02169 | 0.089 |
| ENSG00000174684.6 | 6283.055 | -0.244219839 | 0.1064 | -2.296 | 0.0217 | 0.089 |
| ENSG00000198740.8 | 733.2212 | 0.160063497 | 0.0697 | 2.2956 | 0.0217 | 0.089 |
| ENSG00000254685.6 | 198.7655 | -0.183947936 | 0.0801 | -2.296 | 0.02167 | 0.089 |
| ENSG00000133056.13 | 671.6667 | -0.169630938 | 0.0739 | -2.295 | 0.02176 | 0.0892 |
| ENSG00000106868.16 | 473.0217 | -0.250057749 | 0.109 | -2.294 | 0.0218 | 0.0893 |
| ENSG00000160791.13 | 12.19139 | -0.984727039 | 0.4293 | -2.294 | 0.02182 | 0.0894 |
| ENSG00000152137.6 | 1574.996 | 0.26405326 | 0.1152 | 2.2928 | 0.02186 | 0.0895 |
| ENSG00000073670.13 | 2714.419 | -0.236824497 | 0.1033 | -2.292 | 0.02188 | 0.0896 |
| ENSG00000135605.12 | 17.88222 | -0.355148826 | 0.1549 | -2.292 | 0.0219 | 0.0896 |
| ENSG00000148737.16 | 389.0085 | 0.286008176 | 0.1248 | 2.2921 | 0.0219 | 0.0896 |
| ENSG00000240253.5 | 52.04293 | -0.42172411 | 0.1841 | -2.291 | 0.02195 | 0.0897 |
| ENSG00000053108.16 | 598.4498 | -0.260117461 | 0.1135 | -2.291 | 0.02196 | 0.0897 |
| ENSG00000228021.7 | 84.98823 | 0.224014637 | 0.0978 | 2.291 | 0.02196 | 0.0897 |
| ENSG00000265458.1 | 19.37733 | 0.307343949 | 0.1342 | 2.2906 | 0.02199 | 0.0898 |
| ENSG00000280435.1 | 148.5079 | -0.321948459 | 0.1406 | -2.29 | 0.022 | 0.0898 |
| ENSG00000280893.1 | 14.54709 | -0.582936953 | 0.2545 | -2.29 | 0.02202 | 0.0898 |
| ENSG00000233639.5 | 165.2456 | 0.206472921 | 0.0902 | 2.2899 | 0.02202 | 0.0898 |
| ENSG00000064218.4 | 19.19397 | -0.623678079 | 0.2724 | -2.29 | 0.02203 | 0.0898 |
| ENSG00000233005.1 | 11.75857 | -0.535950888 | 0.2341 | -2.289 | 0.02207 | 0.0899 |
| ENSG00000182551.13 | 1094.882 | 0.203385391 | 0.0889 | 2.2885 | 0.02211 | 0.0901 |
| ENSG00000259205.2 | 32.75073 | -0.437929949 | 0.1914 | -2.288 | 0.02214 | 0.0901 |
| ENSG00000171450.5 | 3145.248 | -0.266091157 | 0.1163 | -2.288 | 0.02215 | 0.0902 |
| ENSG00000183793.13 | 413.0246 | 0.615908377 | 0.2693 | 2.2873 | 0.02218 | 0.0902 |
| ENSG00000186472.19 | 5382.459 | -0.298720013 | 0.1306 | -2.287 | 0.02219 | 0.0902 |
| ENSG00000183317.16 | 888.5141 | -0.238646443 | 0.1044 | -2.287 | 0.02222 | 0.0903 |
| ENSG00000135414.9 | 1176.657 | 0.172420463 | 0.0754 | 2.2862 | 0.02224 | 0.0904 |
| ENSG00000223711.1 | 12.93997 | -0.357832701 | 0.1566 | -2.286 | 0.02227 | 0.0905 |
| ENSG00000112339.14 | 1763.973 | -0.176753272 | 0.0774 | -2.285 | 0.02231 | 0.0906 |
| ENSG00000163914.4 | 9.019271 | -0.71341472 | 0.3123 | -2.284 | 0.02236 | 0.0908 |
| ENSG00000266489.1 | 43.5934 | 0.827046487 | 0.3621 | 2.2839 | 0.02238 | 0.0908 |
| ENSG00000226080.1 | 14.62568 | -0.506390176 | 0.2218 | -2.283 | 0.02241 | 0.0909 |
| ENSG00000253910.2 | 185.902 | 0.29387932 | 0.1287 | 2.2832 | 0.02242 | 0.0909 |
| ENSG00000134317.17 | 194.5531 | -0.195991249 | 0.0859 | -2.283 | 0.02244 | 0.0909 |
| ENSG00000236008.1 | 91.20509 | 0.295382311 | 0.1294 | 2.2829 | 0.02243 | 0.0909 |
| ENSG00000139133.6 | 50.36492 | -0.26705701 | 0.117 | -2.282 | 0.02246 | 0.091 |
| ENSG00000113262.14 | 65.36598 | 0.224807758 | 0.0985 | 2.2813 | 0.02253 | 0.0912 |
| ENSG00000188549.12 | 242.0553 | 0.222602769 | 0.0976 | 2.2812 | 0.02254 | 0.0912 |
| ENSG00000137573.13 | 562.8936 | -0.418682945 | 0.1836 | -2.28 | 0.02258 | 0.0912 |
| ENSG00000144893.12 | 878.1218 | -0.184493562 | 0.0809 | -2.281 | 0.02256 | 0.0912 |
| ENSG00000213713.3 | 365.1595 | -0.27280477 | 0.1196 | -2.28 | 0.02258 | 0.0912 |
| ENSG00000271980.1 | 23.3316 | 0.509206225 | 0.2233 | 2.2805 | 0.02258 | 0.0912 |
| ENSG00000111647.12 | 1560.82 | -0.22761055 | 0.0998 | -2.28 | 0.02259 | 0.0912 |
| ENSG00000139835.13 | 44.21001 | 0.320554432 | 0.1406 | 2.2799 | 0.02261 | 0.0912 |
| ENSG00000140548.9 | 1164.851 | 0.235227514 | 0.1032 | 2.28 | 0.02261 | 0.0912 |
| ENSG00000139998.14 | 4222.165 | -0.249509451 | 0.1095 | -2.28 | 0.02263 | 0.0913 |
| ENSG00000123119.11 | 7863.716 | -0.228334608 | 0.1002 | -2.279 | 0.02269 | 0.0915 |
| ENSG00000130600.16 | 22.80824 | 0.405628246 | 0.178 | 2.2784 | 0.02271 | 0.0915 |
| ENSG00000152102.17 | 6053.468 | -0.200150393 | 0.0879 | -2.278 | 0.02273 | 0.0916 |
| ENSG00000184984.9 | 46.00942 | -0.259204905 | 0.1138 | -2.278 | 0.02274 | 0.0916 |
| ENSG00000113361.12 | 648.0968 | -0.234937553 | 0.1032 | -2.277 | 0.02276 | 0.0916 |
| ENSG00000006432.15 | 2600.767 | -0.251390049 | 0.1104 | -2.277 | 0.0228 | 0.0917 |
| ENSG00000120738.7 | 2718.083 | -0.497813979 | 0.2187 | -2.277 | 0.0228 | 0.0917 |
| ENSG00000234769.7 | 29.40131 | -0.331342218 | 0.1456 | -2.276 | 0.02284 | 0.0918 |
| ENSG00000236627.1 | 9.786204 | 0.566483374 | 0.2489 | 2.2756 | 0.02287 | 0.0919 |
| ENSG00000138115.13 | 110.2683 | 0.316507836 | 0.1391 | 2.2754 | 0.02288 | 0.0919 |
| ENSG00000100234.11 | 2675.638 | 0.38771133 | 0.1704 | 2.2751 | 0.0229 | 0.092 |
| ENSG00000177301.13 | 3824.09 | -0.335511131 | 0.1475 | -2.275 | 0.02292 | 0.092 |
| ENSG00000134755.14 | 102.2363 | 0.272792234 | 0.1199 | 2.2746 | 0.02293 | 0.092 |
| ENSG00000196155.12 | 63.95459 | 0.250894318 | 0.1103 | 2.2743 | 0.02295 | 0.0921 |
| ENSG00000082293.12 | 133.5349 | -0.331702711 | 0.1459 | -2.274 | 0.02297 | 0.0921 |
| ENSG00000254266.5 | 10.41323 | -0.449828814 | 0.1979 | -2.273 | 0.023 | 0.0922 |
| ENSG00000104722.13 | 10490.95 | -0.317117946 | 0.1395 | -2.273 | 0.02303 | 0.0923 |
| ENSG00000259168.1 | 59.33482 | 0.578561846 | 0.2546 | 2.2727 | 0.02304 | 0.0923 |
| ENSG00000134757.4 | 10.06228 | -0.582044351 | 0.2561 | -2.272 | 0.02306 | 0.0923 |
| ENSG00000143507.17 | 93.25437 | -0.318820222 | 0.1404 | -2.271 | 0.02313 | 0.0925 |
| ENSG00000123836.14 | 1106.717 | -0.193254006 | 0.0851 | -2.271 | 0.02314 | 0.0926 |
| ENSG00000157470.11 | 1686.684 | -0.241506451 | 0.1064 | -2.271 | 0.02317 | 0.0926 |
| ENSG00000164142.15 | 23.5512 | -0.392557816 | 0.1729 | -2.27 | 0.02319 | 0.0927 |
| ENSG00000196542.8 | 284.0673 | -0.302632402 | 0.1333 | -2.27 | 0.02322 | 0.0928 |
| ENSG00000149289.10 | 396.3648 | 0.210855191 | 0.0929 | 2.2695 | 0.02324 | 0.0928 |
| ENSG00000131196.17 | 91.27603 | 0.441255445 | 0.1945 | 2.269 | 0.02327 | 0.0928 |
| ENSG00000154654.14 | 4913.756 | -0.237330045 | 0.1046 | -2.269 | 0.02327 | 0.0928 |
| ENSG00000280304.1 | 60.63117 | -0.339444886 | 0.1496 | -2.269 | 0.02326 | 0.0928 |
| ENSG00000242428.5 | 40.63817 | -0.270519172 | 0.1192 | -2.269 | 0.02328 | 0.0929 |
| ENSG00000135824.12 | 625.3334 | -0.353227244 | 0.1557 | -2.268 | 0.02332 | 0.093 |
| ENSG00000165072.9 | 48.87724 | -0.255012529 | 0.1125 | -2.267 | 0.02337 | 0.0931 |
| ENSG00000158125.9 | 10.12806 | 0.379095925 | 0.1672 | 2.2671 | 0.02339 | 0.0932 |
| ENSG00000188013.6 | 22.08288 | -0.52069776 | 0.2298 | -2.266 | 0.02346 | 0.0934 |
| ENSG00000225953.2 | 232.8708 | -0.281955876 | 0.1245 | -2.266 | 0.02348 | 0.0935 |
| ENSG00000206432.4 | 452.4994 | -0.194373892 | 0.0858 | -2.265 | 0.02351 | 0.0936 |
| ENSG00000163590.13 | 2957.814 | -0.265008583 | 0.117 | -2.265 | 0.02354 | 0.0936 |
| ENSG00000213689.10 | 156.4642 | 0.266311281 | 0.1176 | 2.2643 | 0.02355 | 0.0936 |
| ENSG00000281383.1 | 577.2989 | 0.282573177 | 0.1248 | 2.2641 | 0.02357 | 0.0937 |
| ENSG00000110400.10 | 2944.664 | -0.227784067 | 0.1006 | -2.264 | 0.02358 | 0.0937 |
| ENSG00000198794.11 | 9987.953 | -0.233133227 | 0.103 | -2.264 | 0.02359 | 0.0937 |
| ENSG00000013016.15 | 3887.575 | -0.241644422 | 0.1068 | -2.264 | 0.0236 | 0.0937 |
| ENSG00000269694.1 | 44.86098 | 0.296127124 | 0.1309 | 2.263 | 0.02364 | 0.0938 |
| ENSG00000117298.14 | 1009.1 | 0.217632736 | 0.0962 | 2.2628 | 0.02365 | 0.0938 |
| ENSG00000048140.17 | 1149.321 | -0.204297836 | 0.0903 | -2.262 | 0.02368 | 0.0939 |
| ENSG00000268555.1 | 9.270582 | 0.50536267 | 0.2235 | 2.2615 | 0.02373 | 0.094 |
| ENSG00000081059.19 | 61.62206 | 0.267023195 | 0.1181 | 2.2609 | 0.02376 | 0.0941 |
| ENSG00000162512.15 | 7351.406 | 0.174854857 | 0.0774 | 2.2599 | 0.02383 | 0.0943 |
| ENSG00000223930.5 | 55.38058 | -0.312971304 | 0.1385 | -2.26 | 0.02383 | 0.0943 |
| ENSG00000278000.1 | 48.27845 | 0.372619208 | 0.1649 | 2.2599 | 0.02383 | 0.0943 |
| ENSG00000140521.12 | 1047.569 | -0.14983713 | 0.0663 | -2.259 | 0.02386 | 0.0943 |
| ENSG00000162817.6 | 5444.262 | -0.260193054 | 0.1152 | -2.259 | 0.02388 | 0.0943 |
| ENSG00000165379.13 | 972.9659 | -0.184303771 | 0.0816 | -2.259 | 0.02387 | 0.0943 |
| ENSG00000187630.15 | 177.0814 | 0.304349948 | 0.1347 | 2.2595 | 0.02385 | 0.0943 |
| ENSG00000104044.15 | 151.4714 | -0.230639294 | 0.1021 | -2.259 | 0.0239 | 0.0944 |
| ENSG00000165685.8 | 46.09493 | -0.295798548 | 0.131 | -2.259 | 0.02389 | 0.0944 |
| ENSG00000227082.1 | 138.6079 | -0.250865514 | 0.1111 | -2.258 | 0.02393 | 0.0944 |
| ENSG00000264809.2 | 14.12816 | 0.411615543 | 0.1823 | 2.2584 | 0.02392 | 0.0944 |
| ENSG00000104888.9 | 29069.33 | -0.229630088 | 0.1017 | -2.258 | 0.02395 | 0.0944 |
| ENSG00000224272.2 | 84.95638 | 0.420250044 | 0.1862 | 2.2575 | 0.02398 | 0.0945 |
| ENSG00000041353.9 | 2027.385 | -0.341952251 | 0.1515 | -2.257 | 0.024 | 0.0946 |
| ENSG00000103316.10 | 3965.322 | -0.262494655 | 0.1163 | -2.257 | 0.02403 | 0.0946 |
| ENSG00000238276.5 | 22.7659 | 0.411358914 | 0.1823 | 2.2567 | 0.02403 | 0.0946 |
| ENSG00000272668.1 | 41.33234 | 0.386691983 | 0.1714 | 2.2566 | 0.02403 | 0.0946 |
| ENSG00000244482.10 | 20.92094 | -0.376833694 | 0.1671 | -2.256 | 0.0241 | 0.0948 |
| ENSG00000141622.13 | 1396.889 | -0.227404496 | 0.1009 | -2.255 | 0.02416 | 0.095 |
| ENSG00000171502.14 | 875.0764 | -0.220086869 | 0.0976 | -2.254 | 0.02418 | 0.095 |
| ENSG00000248677.1 | 10.37043 | -0.357975585 | 0.1588 | -2.255 | 0.02416 | 0.095 |
| ENSG00000267909.2 | 258.9196 | -0.290297885 | 0.1288 | -2.255 | 0.02416 | 0.095 |
| ENSG00000271533.1 | 456.7064 | -0.206067647 | 0.0914 | -2.254 | 0.02417 | 0.095 |
| ENSG00000108852.14 | 1718.737 | -0.207946074 | 0.0923 | -2.254 | 0.02421 | 0.095 |
| ENSG00000242261.1 | 26.28307 | -0.354890129 | 0.1575 | -2.254 | 0.02421 | 0.095 |
| ENSG00000111424.10 | 22.42742 | -0.367055516 | 0.1629 | -2.253 | 0.02424 | 0.095 |
| ENSG00000213190.3 | 7521.978 | -0.251357987 | 0.1115 | -2.253 | 0.02423 | 0.095 |
| ENSG00000113721.13 | 860.7796 | 0.410354579 | 0.1821 | 2.253 | 0.02426 | 0.0951 |
| ENSG00000272068.1 | 17.81004 | 0.416544206 | 0.1849 | 2.2529 | 0.02427 | 0.0951 |
| ENSG00000123908.11 | 1464.187 | -0.13077142 | 0.0581 | -2.252 | 0.02433 | 0.0953 |
| ENSG00000206344.6 | 63.71152 | 0.292322879 | 0.1298 | 2.2517 | 0.02434 | 0.0953 |
| ENSG00000047597.5 | 557.9111 | -0.270374483 | 0.1201 | -2.251 | 0.02438 | 0.0954 |
| ENSG00000080839.11 | 96.25016 | 0.194879233 | 0.0866 | 2.2498 | 0.02446 | 0.0955 |
| ENSG00000105426.15 | 5245.07 | -0.201510258 | 0.0896 | -2.25 | 0.02447 | 0.0955 |
| ENSG00000132570.14 | 183.681 | 0.157451835 | 0.07 | 2.2497 | 0.02447 | 0.0955 |
| ENSG00000151726.13 | 890.0009 | -0.243680816 | 0.1083 | -2.25 | 0.02442 | 0.0955 |
| ENSG00000153132.12 | 117.2843 | -0.271162249 | 0.1205 | -2.25 | 0.02446 | 0.0955 |
| ENSG00000155760.2 | 153.8081 | 0.362654878 | 0.1612 | 2.2494 | 0.02449 | 0.0955 |
| ENSG00000164241.13 | 81.01725 | 0.224583886 | 0.0998 | 2.2503 | 0.02443 | 0.0955 |
| ENSG00000182168.14 | 688.383 | -0.236463399 | 0.1051 | -2.249 | 0.02448 | 0.0955 |
| ENSG00000272568.5 | 132.8592 | 0.288124802 | 0.1281 | 2.2495 | 0.02448 | 0.0955 |
| ENSG00000119699.7 | 337.5765 | 0.264408092 | 0.1176 | 2.2474 | 0.02461 | 0.096 |
| ENSG00000156011.16 | 13631 | -0.254378003 | 0.1132 | -2.247 | 0.02465 | 0.096 |
| ENSG00000242265.5 | 7144.064 | -0.248482304 | 0.1106 | -2.247 | 0.02464 | 0.096 |
| ENSG00000206535.7 | 263.0639 | -0.252906466 | 0.1126 | -2.246 | 0.02468 | 0.0961 |
| ENSG00000118276.11 | 1835.605 | -0.221681065 | 0.0987 | -2.245 | 0.02474 | 0.0963 |
| ENSG00000158467.16 | 1966.311 | 0.18266618 | 0.0813 | 2.2456 | 0.02473 | 0.0963 |
| ENSG00000278068.1 | 21.89693 | 0.406737301 | 0.1811 | 2.2454 | 0.02475 | 0.0963 |
| ENSG00000130035.6 | 1379.923 | -0.249085659 | 0.111 | -2.245 | 0.02477 | 0.0963 |
| ENSG00000260038.1 | 27.54885 | 0.304361513 | 0.1356 | 2.2449 | 0.02477 | 0.0963 |
| ENSG00000273136.6 | 264.4451 | 0.52337862 | 0.2331 | 2.2451 | 0.02476 | 0.0963 |
| ENSG00000117407.16 | 17.98248 | 0.32329241 | 0.1441 | 2.2434 | 0.02487 | 0.0963 |
| ENSG00000130762.14 | 31.50439 | 0.356051985 | 0.1587 | 2.2436 | 0.02486 | 0.0963 |
| ENSG00000144857.14 | 231.7206 | 0.237157721 | 0.1057 | 2.2438 | 0.02485 | 0.0963 |
| ENSG00000156049.6 | 158.7536 | 0.419006039 | 0.1867 | 2.2442 | 0.02482 | 0.0963 |
| ENSG00000164082.14 | 943.3984 | -0.246030265 | 0.1097 | -2.243 | 0.02487 | 0.0963 |
| ENSG00000166922.8 | 4286.361 | -0.243808046 | 0.1087 | -2.244 | 0.02484 | 0.0963 |
| ENSG00000240747.7 | 17.92845 | 0.398110168 | 0.1774 | 2.2443 | 0.02482 | 0.0963 |
| ENSG00000280129.1 | 9.694871 | 0.413246552 | 0.1841 | 2.2441 | 0.02483 | 0.0963 |
| ENSG00000225569.1 | 28.06232 | -0.554660025 | 0.2473 | -2.243 | 0.02491 | 0.0965 |
| ENSG00000078401.6 | 185.5395 | 0.553391146 | 0.2468 | 2.2424 | 0.02494 | 0.0965 |
| ENSG00000210196.2 | 1769.812 | 0.343126746 | 0.153 | 2.242 | 0.02496 | 0.0966 |
| ENSG00000108375.12 | 115.9674 | 0.501541014 | 0.2237 | 2.2418 | 0.02497 | 0.0966 |
| ENSG00000204965.8 | 276.9495 | -0.302972258 | 0.1352 | -2.242 | 0.02499 | 0.0966 |
| ENSG00000133321.10 | 112.2649 | 0.374415086 | 0.1671 | 2.241 | 0.02503 | 0.0967 |
| ENSG00000272372.1 | 73.41219 | 0.336613907 | 0.1502 | 2.2404 | 0.02506 | 0.0968 |
| ENSG00000135298.13 | 2651.302 | -0.196744173 | 0.0878 | -2.24 | 0.02507 | 0.0968 |
| ENSG00000248115.1 | 17.94231 | -0.477937271 | 0.2134 | -2.24 | 0.02509 | 0.0969 |
| ENSG00000151948.11 | 1209.627 | -0.274965981 | 0.1228 | -2.239 | 0.02517 | 0.0971 |
| ENSG00000228824.6 | 162.3817 | -0.264038086 | 0.1179 | -2.239 | 0.02518 | 0.0971 |
| ENSG00000233008.5 | 22.05111 | 0.358620749 | 0.1602 | 2.2387 | 0.02517 | 0.0971 |
| ENSG00000154478.3 | 764.1006 | -0.45680509 | 0.2041 | -2.238 | 0.02519 | 0.0971 |
| ENSG00000104808.7 | 47.93675 | -0.369896118 | 0.1653 | -2.238 | 0.02523 | 0.0972 |
| ENSG00000170629.14 | 482.1884 | -0.342372823 | 0.1531 | -2.237 | 0.02529 | 0.0974 |
| ENSG00000168032.8 | 453.7485 | -0.268682131 | 0.1202 | -2.236 | 0.02536 | 0.0977 |
| ENSG00000283417.1 | 32.69208 | -0.328294389 | 0.1469 | -2.235 | 0.02539 | 0.0977 |
| ENSG00000067182.7 | 350.6271 | 0.382365908 | 0.1711 | 2.2343 | 0.02546 | 0.0978 |
| ENSG00000070190.12 | 16.73079 | -0.422286717 | 0.189 | -2.234 | 0.02545 | 0.0978 |
| ENSG00000125945.14 | 834.3393 | -0.224181546 | 0.1003 | -2.235 | 0.02543 | 0.0978 |
| ENSG00000185610.6 | 80.71627 | 0.326731456 | 0.1462 | 2.2344 | 0.02546 | 0.0978 |
| ENSG00000203709.10 | 450.0646 | -0.206623551 | 0.0925 | -2.234 | 0.02546 | 0.0978 |
| ENSG00000164818.15 | 283.281 | 0.134070855 | 0.06 | 2.234 | 0.02548 | 0.0978 |
| ENSG00000166326.6 | 5998.042 | -0.193197398 | 0.0865 | -2.234 | 0.02549 | 0.0978 |
| ENSG00000001461.16 | 2806.405 | -0.217945959 | 0.0976 | -2.234 | 0.0255 | 0.0979 |
| ENSG00000228400.1 | 10.11431 | -0.455038204 | 0.2037 | -2.234 | 0.02551 | 0.0979 |
| ENSG00000149403.11 | 668.9086 | -0.165224193 | 0.074 | -2.232 | 0.02561 | 0.0982 |
| ENSG00000163249.10 | 469.5451 | -0.208405097 | 0.0934 | -2.232 | 0.02561 | 0.0982 |
| ENSG00000174808.11 | 52.02962 | -0.220858398 | 0.099 | -2.231 | 0.02565 | 0.0983 |
| ENSG00000215244.2 | 29.65482 | 0.438076857 | 0.1963 | 2.2315 | 0.02565 | 0.0983 |
| ENSG00000145349.16 | 4485.754 | -0.22682159 | 0.1017 | -2.231 | 0.02569 | 0.0984 |
| ENSG00000119121.21 | 85.10995 | -0.307352106 | 0.1378 | -2.23 | 0.02577 | 0.0986 |
| ENSG00000171435.13 | 1730.968 | -0.235846268 | 0.1058 | -2.23 | 0.02577 | 0.0986 |
| ENSG00000100092.20 | 246.1309 | -0.204792025 | 0.0919 | -2.229 | 0.02583 | 0.0988 |
| ENSG00000132141.13 | 71.84244 | 0.198927 | 0.0893 | 2.2281 | 0.02588 | 0.0989 |
| ENSG00000155629.14 | 115.1217 | -0.360536658 | 0.1618 | -2.228 | 0.02588 | 0.0989 |
| ENSG00000248184.1 | 12.76412 | 0.469160315 | 0.2106 | 2.228 | 0.02588 | 0.0989 |
| ENSG00000100300.17 | 171.0362 | 0.326637145 | 0.1467 | 2.227 | 0.02595 | 0.0989 |
| ENSG00000163376.11 | 128.3633 | -0.248351991 | 0.1115 | -2.227 | 0.02592 | 0.0989 |
| ENSG00000166450.12 | 80.53426 | 0.278474656 | 0.125 | 2.227 | 0.02595 | 0.0989 |
| ENSG00000171722.12 | 19.9928 | -0.389407178 | 0.1748 | -2.227 | 0.02594 | 0.0989 |
| ENSG00000225975.6 | 80.02693 | -0.250268926 | 0.1124 | -2.227 | 0.02592 | 0.0989 |
| ENSG00000019582.14 | 2435.377 | -0.630869082 | 0.2833 | -2.227 | 0.02597 | 0.099 |
| ENSG00000183775.10 | 3360.693 | -0.286033076 | 0.1285 | -2.226 | 0.02599 | 0.0991 |
| ENSG00000173276.13 | 600.2033 | -0.188321721 | 0.0846 | -2.225 | 0.02605 | 0.0991 |
| ENSG00000185274.11 | 4578.269 | -0.284951039 | 0.128 | -2.225 | 0.02605 | 0.0991 |
| ENSG00000186439.12 | 27.99042 | 0.314921303 | 0.1415 | 2.2256 | 0.02604 | 0.0991 |
| ENSG00000203772.7 | 1415.922 | -0.268559051 | 0.1206 | -2.226 | 0.02602 | 0.0991 |
| ENSG00000231711.2 | 71.49915 | 0.277775778 | 0.1248 | 2.2254 | 0.02605 | 0.0991 |
| ENSG00000118946.11 | 3176.141 | -0.232112323 | 0.1043 | -2.225 | 0.02607 | 0.0991 |
| ENSG00000012211.12 | 34.89058 | 0.303144349 | 0.1363 | 2.2248 | 0.02609 | 0.0992 |
| ENSG00000215458.8 | 58.11348 | -0.235650863 | 0.1059 | -2.225 | 0.02611 | 0.0992 |
| ENSG00000078902.15 | 7939.543 | -0.217374196 | 0.0977 | -2.224 | 0.02615 | 0.0993 |
| ENSG00000153707.16 | 4016.224 | -0.22899815 | 0.103 | -2.224 | 0.02615 | 0.0993 |
| ENSG00000272273.1 | 22.96303 | 0.384289328 | 0.1728 | 2.2241 | 0.02614 | 0.0993 |
| ENSG00000254758.1 | 14.71107 | -0.393771221 | 0.1771 | -2.223 | 0.02621 | 0.0994 |
| ENSG00000256943.1 | 17.78831 | -0.444346783 | 0.2 | -2.222 | 0.02627 | 0.0996 |
| ENSG00000237984.3 | 76.9533 | -0.223261379 | 0.1005 | -2.221 | 0.02632 | 0.0998 |
| ENSG00000140416.19 | 2716.889 | -0.238970024 | 0.1076 | -2.221 | 0.02635 | 0.0999 |
| ENSG00000168297.15 | 1171.595 | -0.245122741 | 0.1104 | -2.221 | 0.02636 | 0.0999 |
| ENSG00000132386.10 | 962.023 | -0.265786974 | 0.1197 | -2.22 | 0.02639 | 0.0999 |
| ENSG00000164171.10 | 166.2902 | -0.304401996 | 0.1371 | -2.22 | 0.02641 | 0.1 |
| ENSG00000078269.13 | 2083.579 | -0.215622017 | 0.0971 | -2.219 | 0.02645 | 0.1 |
| ENSG00000100311.16 | 519.7264 | -0.200935497 | 0.0905 | -2.22 | 0.02645 | 0.1 |
| ENSG00000118898.15 | 947.0337 | -0.305013294 | 0.1374 | -2.22 | 0.02645 | 0.1 |
| ENSG00000276291.5 | 194.2114 | -0.250190808 | 0.1127 | -2.22 | 0.02643 | 0.1 |

**Supplementary Table 4 Low Axis-1 Comorbidity vs High Axis-1 Comorbidity**

| Ensemble ID | baseMean | log2FoldChange | lfcSE | stat | pvalue | padj |
| --- | --- | --- | --- | --- | --- | --- |
| ENSG00000211974.3 | 1.20907159 | -29.9998116 | 3.000034 | -9.99982 | 1.53E-23 | 1.10E-19 |
| ENSG00000244116.3 | 1.30879004 | -29.9997267 | 2.9970796 | -10.0097 | 1.38E-23 | 1.10E-19 |
| ENSG00000099139.13 | 8665.6939 | -3.42602044 | 0.5002031 | -6.84926 | 7.42E-12 | 3.57E-08 |
| ENSG00000225217.1 | 74.9021577 | -2.69420718 | 0.4222419 | -6.38072 | 1.76E-10 | 6.35E-07 |
| ENSG00000229344.1 | 12.9807485 | 2.10792157 | 0.3418062 | 6.16701 | 6.96E-10 | 2.01E-06 |
| ENSG00000275302.1 | 32.4254428 | -1.83281676 | 0.347617 | -5.27252 | 1.35E-07 | 0.00032 |
| ENSG00000007908.15 | 61.9222933 | -1.86472688 | 0.429941 | -4.33717 | 1.44E-05 | 0.02971 |
| ENSG00000169248.12 | 16.7269605 | -1.79350166 | 0.4176254 | -4.29452 | 1.75E-05 | 0.03153 |
| ENSG00000266146.1 | 2.23146825 | -1.48679544 | 0.3511469 | -4.23411 | 2.29E-05 | 0.03306 |
| ENSG00000273679.1 | 3.17546247 | 1.18656433 | 0.2787054 | 4.25741 | 2.07E-05 | 0.03306 |
| ENSG00000277632.1 | 33.2066802 | -1.92628297 | 0.4752081 | -4.05356 | 5.04E-05 | 0.06608 |
| ENSG00000276070.4 | 24.5961015 | -1.11735361 | 0.2788789 | -4.00659 | 6.16E-05 | 0.07397 |
